# From neglecting to including cultivar-specific *per se* temperature responses: Extending the concept of thermal time for plant development modeling

**DOI:** 10.1101/2023.08.29.555271

**Authors:** Lukas Roth, Martina Binder, Norbert Kirchgessner, Flavian Tschurr, Steven Yates, Andreas Hund, Lukas Kronenberg, Achim Walter

## Abstract

Predicting plant development, a longstanding goal in plant physiology, involves two interwoven components: continuous growth and the progression of growth stages (phenology). Current models, like thermal time, assume species-level growth responses to temperature. We challenge this assumption, suggesting that cultivar-specific temperature responses significantly affect phenology. To investigate, we collected field-based growth and phenology data in winter wheat and soybean over multiple years. We used diverse models, from linear to neural networks, to assess growth responses to temperature at various trait and covariate levels. Cultivar-specific non-linear models best explained phenology-related cultivar-environment interactions. With cultivar-specific models, additional relations to other stressors than temperature were found. The availability of the presented field phenotyping tools allows incorporating cultivar-specific temperature response functions in future plant physiology studies, which will deepen our understanding of key factors that influence plant development. Consequently, this work has implications for crop breeding and cultivation under adverse climatic conditions.

## 1. Introduction

To mitigate the effects of global environmental change on crop production, a profound understanding of its influence on plant growth is required (Ramirez-Villegas, Watson, and Challinor 2015). Crop models promise to be a versatile tool in analyzing and predicting plant growth (Pauli et al. 2016), in particular for future climate scenarios (White et al. 2011; Tardieu et al. 2020). Yet, the model choice represents a challenging trade-off between biological realism and the principle of parsimony (Hammer et al. 2019).

From a temporal (i.e., growth process based) perspective, plant growth appears non-linear (Figure 1a). Rapid short-term changes in environmental conditions result in related short-term growth patterns (Nagelmüller et al. 2016). These patterns are superimposed on seasonal changes of environmental conditions. On top of these relations, stress conditions may result in yet another superimposed (and potentially negative) growth pattern (Tschurr et al. 2023).

**Figure 1:**
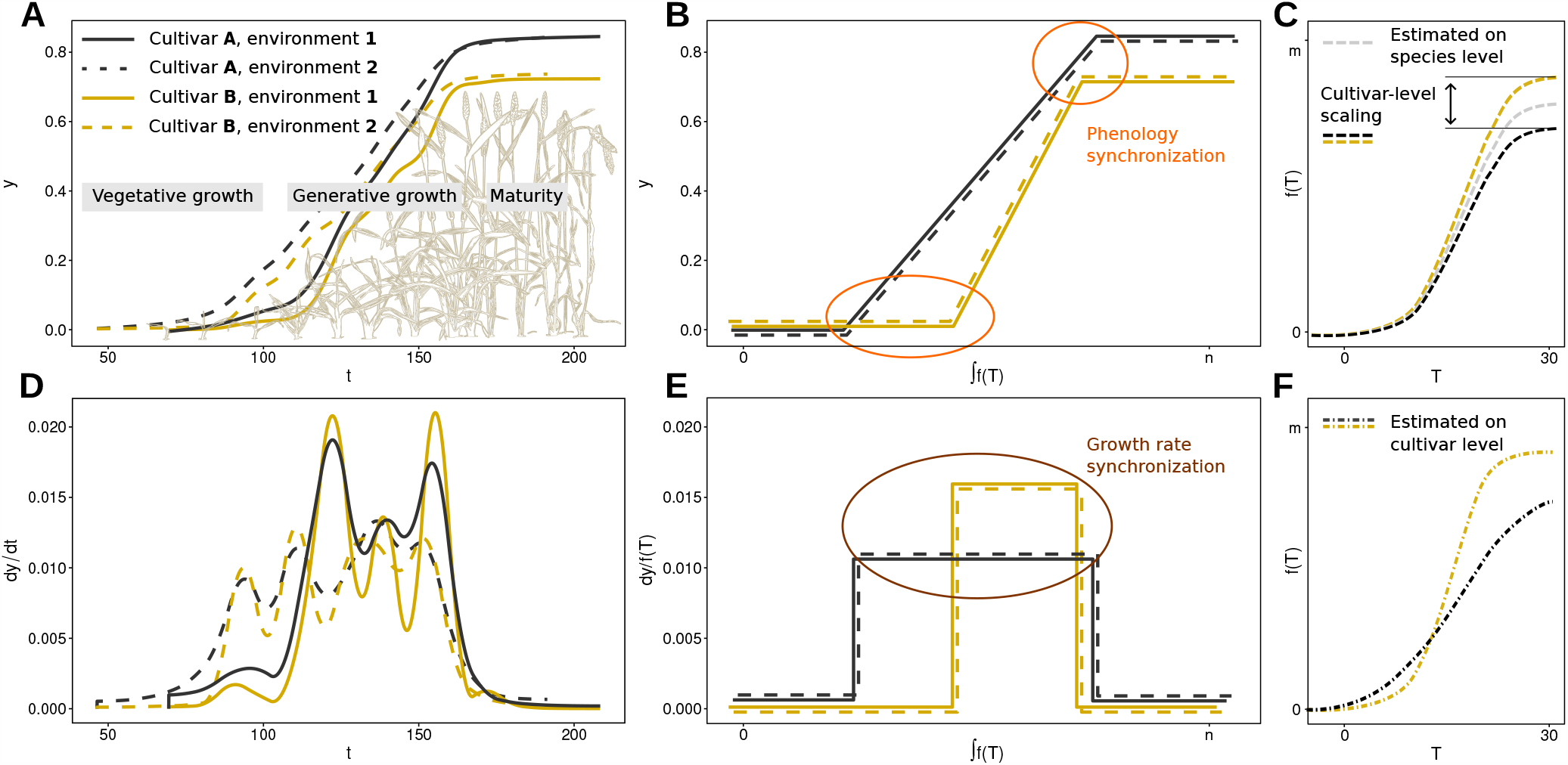
Schematic visualization of strategies in crop modeling to compensate for fluctuating temperatures on the example of generative growth in winter wheat. Plant growth over time appears non-linear (a) and reveals irregular, potentially cultivar and environment specific growth rate patterns (d). By replacing time *t* with the area under the curve of a temperature-compensation function, ∫*f* (*T*_*t*_), phenology stages (b) and growth rates (e) may be synchronized in respect to the independent variable. Temperature-compensation functions can be based on species-level dose-response curves scaled to cultivar-level intrinsic growth rates (c), or on cultivar-specific dose-response curves (f).

Finally, temporal patterns are also caused by advancing plant development, known as phenology. Fundamental influencing factors in cultivar-specific phenology include photoperiod sensitivity (Steinberg and Garner 1936), and, in the case of winter cereals, vernalization requirements (Slafer 1996). If modeling phenology on a rather small scale in environments with neglectable differences in photoperiod and vernalization, temperature remains as a dominant driver of phenology (Bogard et al. 2014; Ochagavía et al. 2019).

Consequently, a common modeling approach is to temperature-compensate time, thus ‘linearize’ growth (Figure 1e) and phenology (Figure 1b) using a species-specific *per se* temperature growth response (Figure 1c) (Bonhomme 2000). Differences in growth rates and phenology between cultivars are then modeled using cultivar-specific factors that scale the predicted growth rate to measured (i.e., cultivar ‘intrinsic’) growth rates (Parent and Tardieu 2012). While it was shown that extending linear temperature responses (i.e., thermal time) to non-linear functions can further improve predictions (Wang et al. 2017), Parent and Tardieu (2012) provided evidence that modeling a *per se* temperature response at species-level is sufficient. They speculated that evolutionary processes may have fixed the response for lower plant systematic levels. Hence, using linear and non-linear temperature compensation functions with fixed, literature-based parameters seems justified.

Nevertheless, there is evidence that phenology is related to cultivar-specific temperature responses (Kronenberg et al. 2020a). Consequently, one may assume that selecting for phenology traits in breeding—such as earlier flowering in winter wheat—co-selected for temperature response (Roth et al. 2022b). Indeed, we have repeatedly observed cultivar-specific temperature responses in our outdoor, high-throughput phenotyping site at ETH Zurich. We found cultivar-specific differences in the temperature response in the early canopy development of winter wheat (Grieder, Hund, and Walter 2015; Nagelmüller et al. 2016), as well as in the stem elongation phase of winter wheat (Kronenberg et al. 2020a; Roth, Piepho, and Hund 2022) and soybean (Friedli et al. 2016). Furthermore, we found that the differences in the stem elongation phase of winter wheat were related to the breeding origin of cultivars (Roth et al. 2022b) and allow a ‘phenomic prediction/selection’ for yield (Roth et al. 2023).

Similar observations have been made in crop modeling for other crops than wheat and soybean. Wallach et al. (2018) could demonstrate that it is feasible to include a cultivar-specific temperature response parameter (*T*_opt_) for flowering time predictions in common bean. Viswanathan et al. (2022) were able to optimize two temperature response parameters (*T*_min_ and *T*_opt_) for two growth stages in maize.

Given these evidences, it is striking how rarely temperature response parameters are included in the optimization process in crop growth and phenology models. Reasons may be found in the state-of-the-art use of so-called multi-environmental trials (MET) where the phenology of cultivars is measured in different environments. The heterogeneity of the environments often requires including other factors such as photoperiod sensitivity and vernalization requirements (White et al. 2008). The additional inclusion of temperature response parameters will bring the number of parameters that require optimization close to the degrees of freedom of the data. One suggestion to overcome this limitation is to incorporate the genetic relatedness of cultivars, e.g, using QTLs (Wallach et al. 2018) or whole genome predictions (Messina et al. 2018).

Another way to address the problem is to massively increase the number of data points per cultivar and environment. As phenology consists of single events, this can only be achieved by measuring continuous growth instead. Field-based plant organ tracking devices (Mielewczik et al. 2013; Nagelmüller et al. 2016) and field-phenotyping platforms (Kirchgessner et al. 2017) can provide such dense time series with tens to thousands of growth rate / temperature value pairs. Simulation data (Roth, Piepho, and Hund 2022) and real-world data analysis (Millet et al. 2019; Roth et al. 2022b) have shown that, provided the temporal density of the time series is high enough, a few environments are sufficient to reliably determine cultivar-specific responses.

The question arises, whether transferring such pre-calibrated cultivar-specific temperature responses— determined on dense time series in few environments—to crop models may improve phenology predictions. Studies reporting phenology stages in species-specific thermal time often found severe genotype-by-environment (G*×* E) interactions in their data (Sadras et al. 2009; Salazar-Gutierrez et al. 2013; Slafer et al. 2015). We suspect that large portions of the reported G *×* E interactions in phenology are artifacts of over-generalizing the *per se* temperature response on species-level. In other words, when explaining the observed performance of plants in different environments, thermal time is reificated— its abstract concept is by mistake treated as being a real, interpretable object.

If our assumption holds, using cultivar-specific linear or non-linear temperature responses will improve the estimation of growth rates (Figure 1e) and of phenology stages (Figure 1b). To test this hypothesis, we evaluated a unique, temporally very dense multi-environment (i.e., one site, multiple years) outdoor winter wheat and soybean data set. The setup corresponds to the specific situation where the prediction is to be improved for a clearly defined environment (in our case, Switzerland). Measurements and predictions are made under near-constant photoperiod and vernalization conditions, allowing to focus on temperature response only.

The data were collected with temporally-resolved leaf growth tracking devices as well as highthroughput field phenotyping devices (Figure 2). We evaluated the response of traits to temperature using models of increasing complexity (ranging from linear models to hierarchical splines and neural networks). Additionally, the trait level (leaf growth, canopy development, stem elongation), the covariate level (soil temperature, air temperature) and the covariate measurement level (below/inside canopies, at a reference weather station) were varied. Finally, phenology period estimations for the three main growth phases of wheat (vegetative growth, generative growth, maturity) were performed using the pre-parameterized temperature response models.

**Figure 2:**
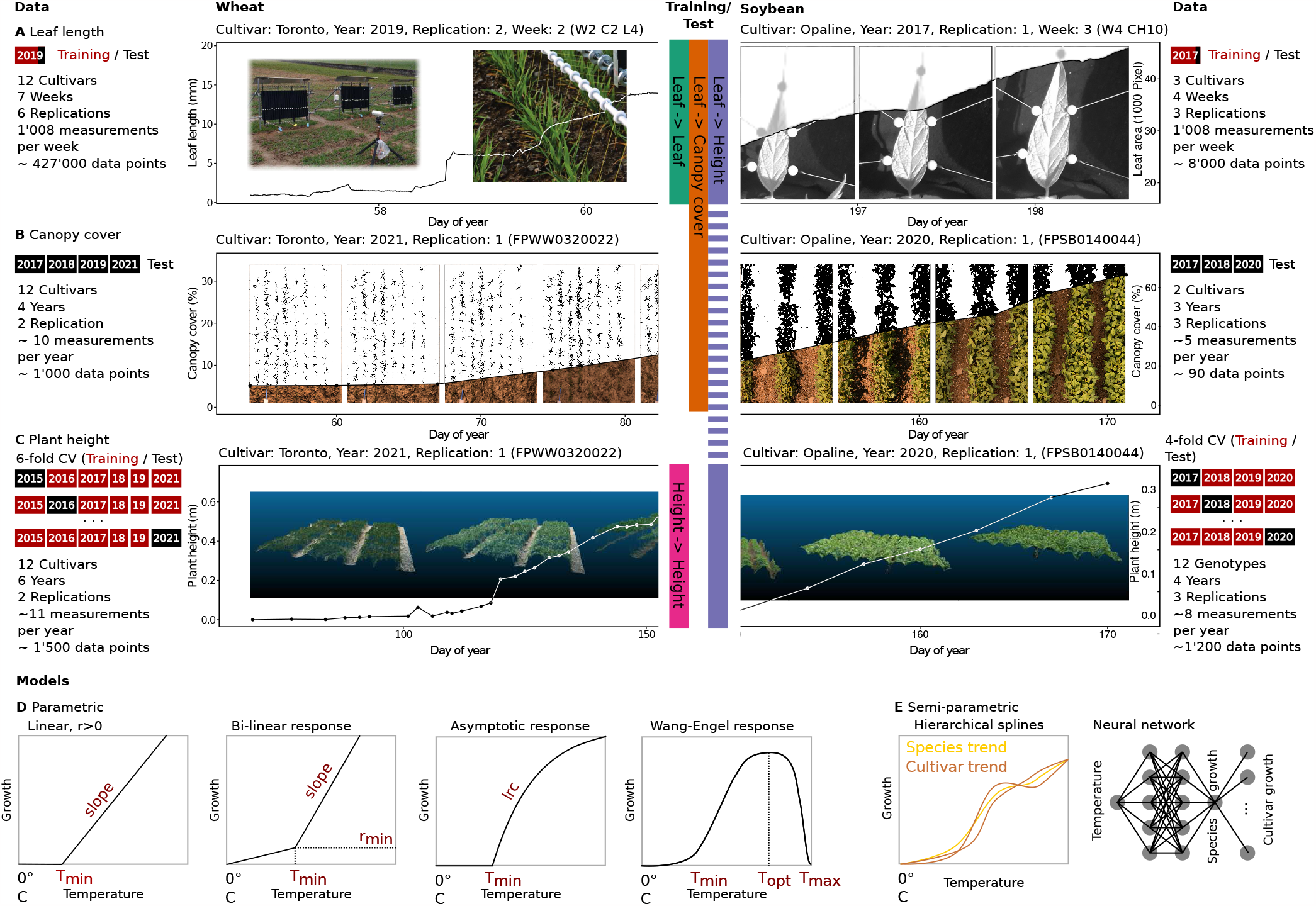
Evaluated data set for winter wheat and soybean. Leaf elongation and growth was measured using a leaf length tracker device for wheat and a leaf growth tracker device for soybean (a). Canopy cover observations were taken using a field phenotyping platform to collect RGB images, followed by segmenting them in pixels showing plants and soil (b). Plant height measurements were performed using a phenotyping platform based terrestrial laser scanner and drone-based Structure-from-Motion (SfM) techniques (c). Four parametric dose-response models (d) and two semi-parametric dose-response models (e) were evaluated. Model testing was performed on unseen data in test-train splits (a, b) and cross-validations (CV) (c).

## 2. Results

### 2.1 Growth rate prediction

A first aim of the study was to identify suitable temperature response models to predict continuous growth, and to test for the transfer ability of these trained models from one growth stage to others. This step will provide insight into the importance of model choice, explaining covariates, and variety versus species-level.

Parametric models (Equation 3–6), hierarchical splines, and a neural network model were trained on leaf and plant height growth data sets (Figure 2a and 2c). For all models, cultivar-specific (Equation 1) and species-level (Equation 2) variations were considered. Trained models were tested on unseen leaf, canopy, and plant height growth data sets (Figure 2a–c). Tests were based on measured and predicted differences between consecutive measurements using random regressions (Equation 7) that account for year effects (Equation 8) and, in case species-level models were fitted, for cultivar-specific scaling (Equation 9).

In summary, cultivar-level models and species-level models were equally well suited to model growth (Figure 3). Scaling species-level models such as thermal time to cultivar-specific intrinsic growth rates successfully predicted growth for unseen test sets (Figure 3b). However, fitted cultivar-level models showed clear *per se* temperature response characteristics, as indicated for example by cross-overs between cultivar-level response curves for the bi-linear model (Figure 3a). Using such cultivar-specific models to predict unseen growth data test sets resulted in similar performance as for species-level models (Figure 3b). More important than the model choice was the choice of covariate (soil or air temperature) in relation to the growth phases. In the following, detailed results will be reported for wheat and soybean.

**Figure 3:**
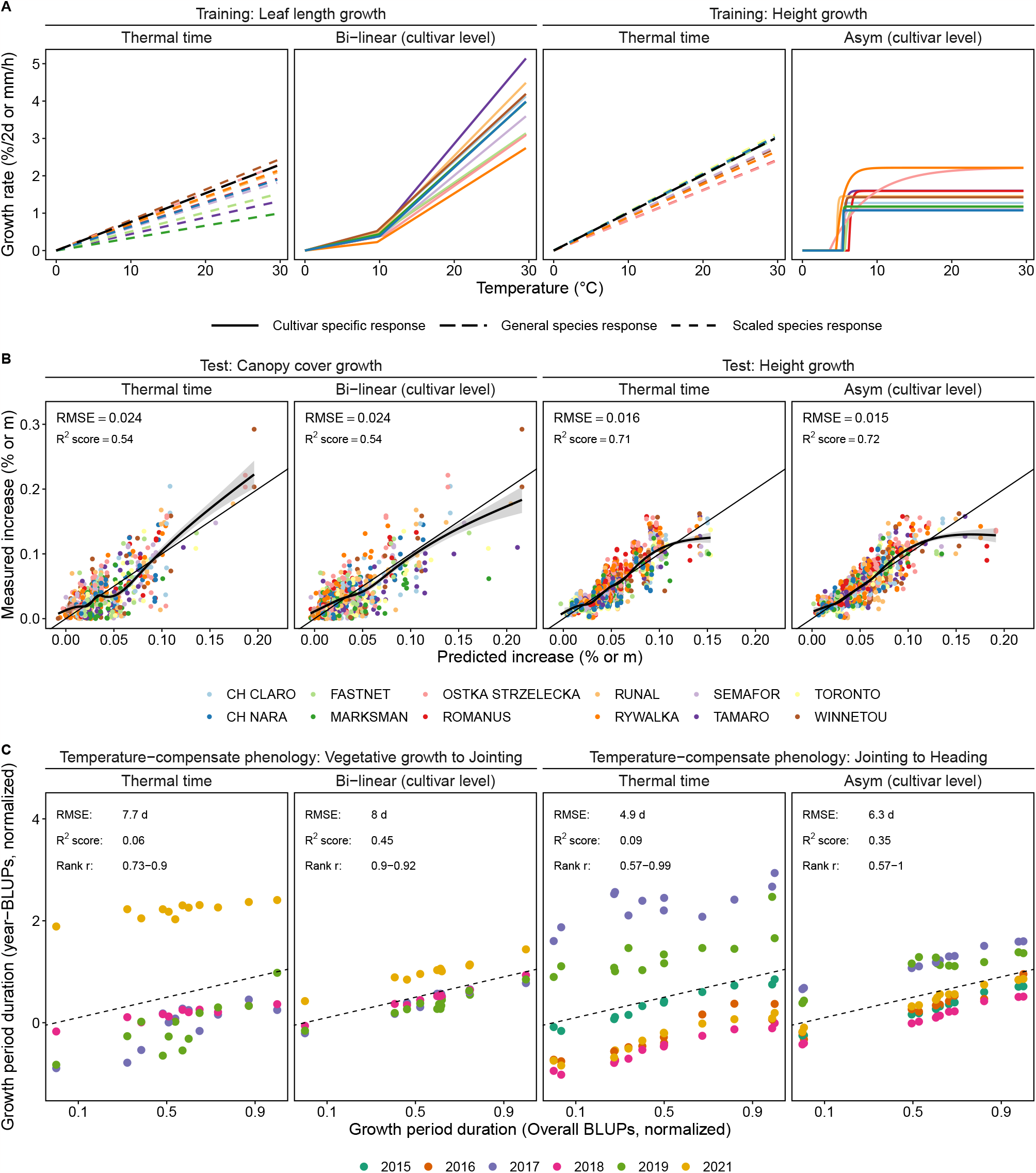
Overview of performance of the best species-level and cultivar-level models for winter wheat. While species-specific thermal time models (Equation 2) where scaled to cultivar-specific intrinsic growth rates (Equation 3), cultivar-level bi-linear (Equation 4) and asymptotic models (Equation 5) were fitted for each cultivar separately (Equation 1) (a). Models were tested on unseen growth data sets (b) and unseen phenology data sets (c) to indicate their potential to predict growth and to reduce estimated G*×* E in phenology. For a full comparison of model performance for both wheat and soybean, see Figure 4 for growth predictions and Figure 5 for phenology.

#### 2.1.1 Wheat growth rate predictions

Cultivar-specific response models outperformed species-level models if training and test sets were closely related (Figure 4, first row). If training and test sets originated from the leaf elongation measurements, the highest growth prediction accuracy was reached by three cultivar-specific models: The bi-linear model, the hierarchical splines, and the neural network (*R*^2^ = 0.22, root-mean-squared error (RMSE)=0.39 mm/h). Relying on plot-based temperature measurements outperformed reference station measurements (Δ*R*^2^ = 0.08). Using plot-based soil temperatures slightly outperformed using plot-based air temperatures (Δ*R*^2^ = 0.01).

**Figure 4:**
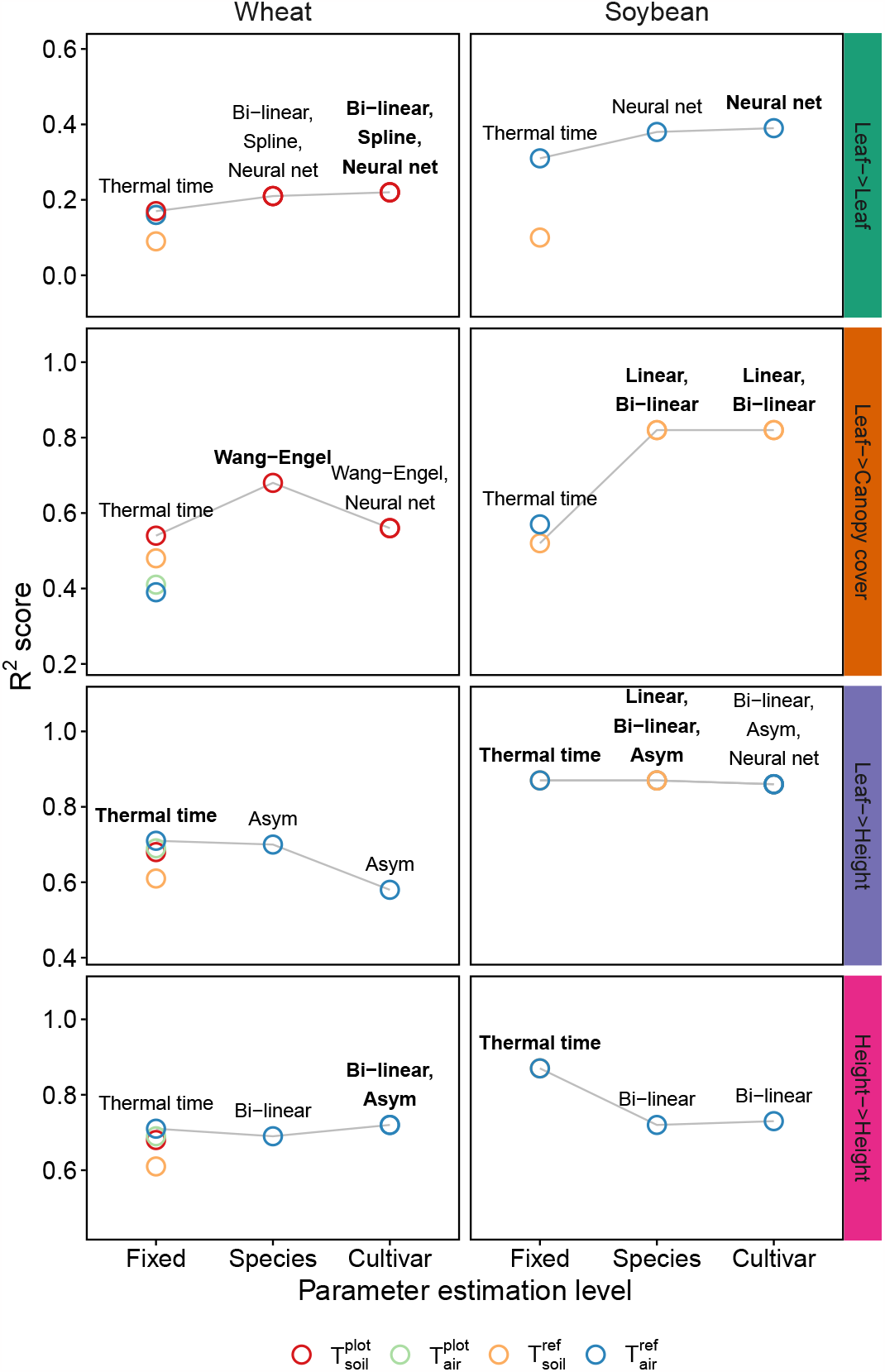
Performance of linear (Equation 3) and non-linear (Equation 4–6) models for growth rate predictions (Equation 7) in winter wheat and soybean. Models were trained on leaf length data (Leaf->) and plant height data (Height->). Predictions were tested on unseen leaf length data (->Leaf), canopy cover data (->Canopy cover), and plant height data (->Height). The covariate temperature was measured in air (T_air_) and soil (T_soil_) at plot level (T^plot^) and at a reference station (T^ref^). Model parameters were either estimated on species (Equation 9) or cultivar-level (Equation 8) or based on literature and therefore fixed. Indicated are the coefficients of determination (R^2^ score) of predictions of the best performing temperature response model per training/test combination and parameter estimation level. Overall best performing models per training/test combination are indicated in **bold**.

If training and test sets differed, species-level response models generalized better (Figure 4, second and third row). When applying models trained on leaf elongation measurements to whole-canopy measurements made in the same growth phase, the species-level Wang-Engel model (*R*^2^ = 0.68, RMSE = 0.44%/day) outperformed the cultivar-level Wang-Engel and neural network models (Δ*R*^2^ = −0.12) and thermal time (Δ*R*^2^ = −0.14). Again, using soil temperature at the plot level resulted in a higher accuracy than using soil temperature or air temperature measured at a reference station (Δ*R*^2^ ≥ 0.06).

When further reducing the relatedness of training and test sets by predicting plant height growth with leaf elongation models, thermal time (*R*^2^ = 0.71, RMSE = 0.18 mm/day) performed slightly better than the asymptotic species-level model (Δ*R*^2^ = −0.01) and the corresponding cultivar-level model (Δ*R*^2^ =− 0.13).

The advantage of cultivar-level models could be restored by training models directly on plant height data, thus using closely related training and test sets (Figure 4, last row). Two cultivar-level models were suggested, the asymptotic and bi-linear dose-response curves (*R*^2^ = 0.72, RSME: 0.16 mm/day). To predict plant height data, air temperature measured at the reference weather station was better suited than plot-based measurements (Δ*R*^2^ ≥ 0.02).

#### 2.1.2 Soybean growth rate predictions

As for the wheat data set, the model performance in soybean was dependent on the relatedness of training and test sets. If the training and test set both originated from the leaf growth measurements (Figure 4, first row), the cultivar-level neural network performed best (*R*^2^ = 0.39, RMSE = 0.46h). Using air temperature clearly outperformed soil temperature (Δ*R*^2^ = 0.21).

If training and test sets differed, species-level response models performed as good as cultivarlevel response models (Figure 4, second and third row). For canopy cover growth predictions, simple cultivar or species-specific models (linear and bi-linear) performed best (*R*^2^ = 0.82, RMSE = 1.0%/day). Strikingly, while air temperature performed better than soil temperature for thermal time (Δ*R*^2^ = 0.05), for species and cultivar-specific models, growth was best predicted by soil temperature (Δ*R*^2^ = 0.30).

If further reducing the relatedness of training and test sets by predicting plant height with leaf growth models, species-level models were more accurate than cultivar-level models. Literature based, linear thermal time performed equally well as the three best species-level models; i.e., the linear, bi-linear, and asymptotic model (*R*^2^ = 0.87, RMSE = 3.8–4.4 mm/day). Nevertheless, differences to the best cultivar-level model were very small (Δ*R*^2^ = 0.01).

In contrast to winter wheat, training models directly on soybean plant height data could not restore the advantage of cultivar-level models (Figure 4, last row). While the bi-linear model performed best on the species and cultivar-level (*R*^2^ = 0.73, RMSE = 5.1 mm/day), its performance was still worse than that of the thermal time model (*R*^2^ = 0.87).

### 2.2 Phenology prediction

A second aim of the study was to test the hypothesis whether phenology is driven by the previously extracted cultivar-specific temperature responses or not. Time periods between successive growth stages (e.g., jointing to heading) per cultivar and year were either expressed in thermal time, using species-level non-linear temperature response models, or using cultivar-level models (Equation 11). Then, G *×* E interactions were estimated using a linear mixed model (Equation 12).

Cultivar-level models showed a clear advantage over species-level models (Figure 3c). While the prediction error between models was comparable, using cultivar-level models resulted in higher cultivar rank correlations between environments, and better correspondences between overall BLUPs and year-BLUPs. The findings indicated that temperature-compensating with cultivar-level models decreases the estimate G*×*E effects for phenology, while using thermal time inflates these effects. In the following, detailed results and consequences for G*×*E analysis are provided.

#### 2.2.1 Severity of the estimated G*×*E interaction for different models

Using thermal time as temperature response model resulted in large estimated variances of G *×* E interactions (Figure 5 and Appendix Figure A.3). Depending on the growth stage, up to 73–82% of the total genotypic variance was related to G*×*E. Consequently, variety rank changes across years were frequent, and rank correlations between years and overall means varied widely (*r* =0.57– 0.99). RMSEs of predictions in calendar days were larger for earlier growth stages than for later growth stages (7.7 days for vegetative growth versus 1.8 days for maturity) (Figure 6).

**Figure 5:**
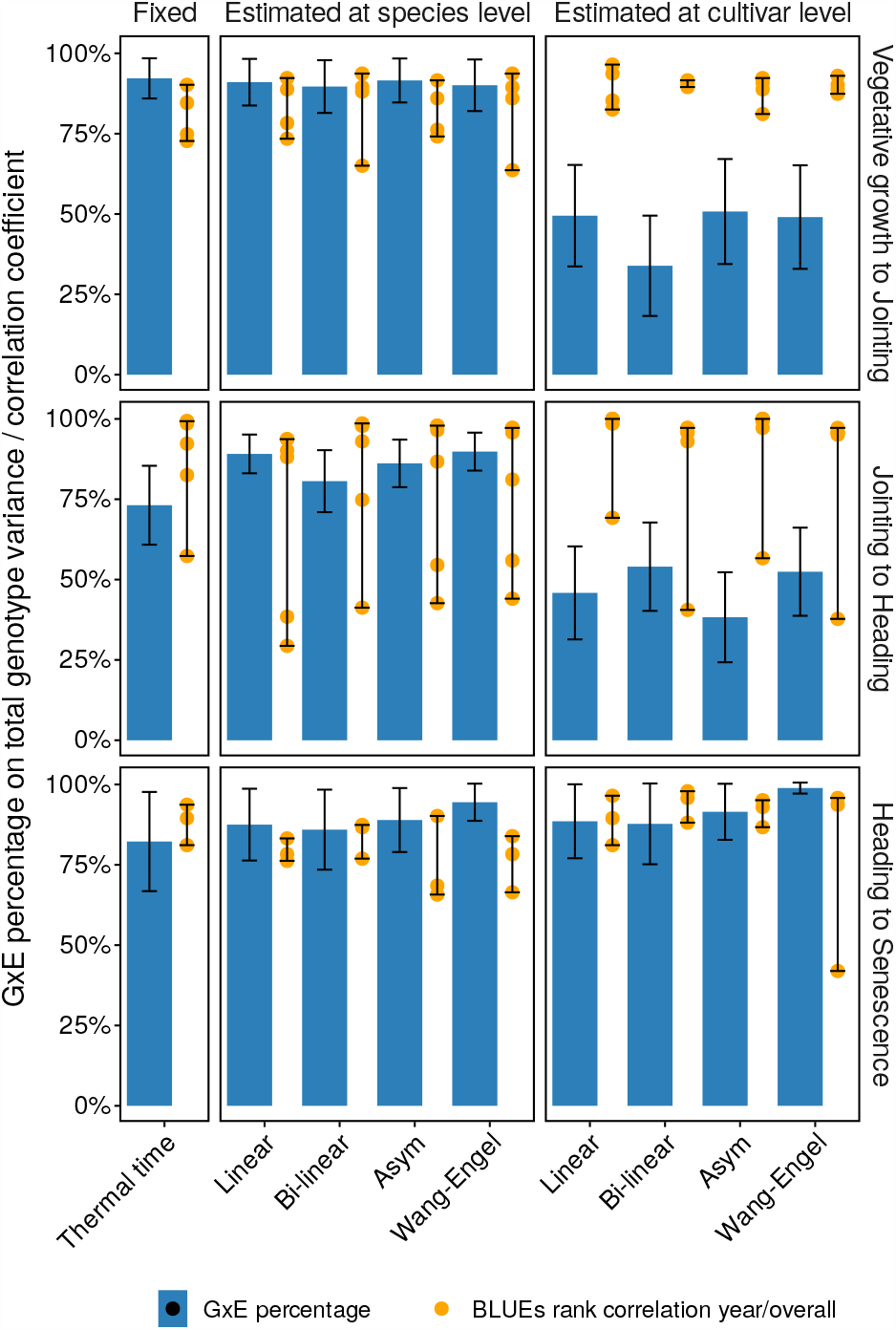
Performance of linear and non-linear models for temperature compensated phenology period duration predictions (Equation 11) in winter wheat. Predictions were based on cultivar-level best linear unbiased estimations (overall BLUPs) of a linear mixed model (Equation 12) that included effects for cultivars (*g*_*i*_), years (*v*_*j*_), and year-cultivar interactions ((*vg*)_*i j*_). Indicated are the percentage of estimated G*×*E variance 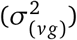 on the total genotypic variance 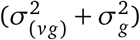, and Spearman’s rank correlations of year-specific (*v*_*j*_ + *g*_*i*_ + (*vg*)_*i j*_) versus overall (*g*_*i*_) phenology duration predictions (Equation 12).

**Figure 6:**
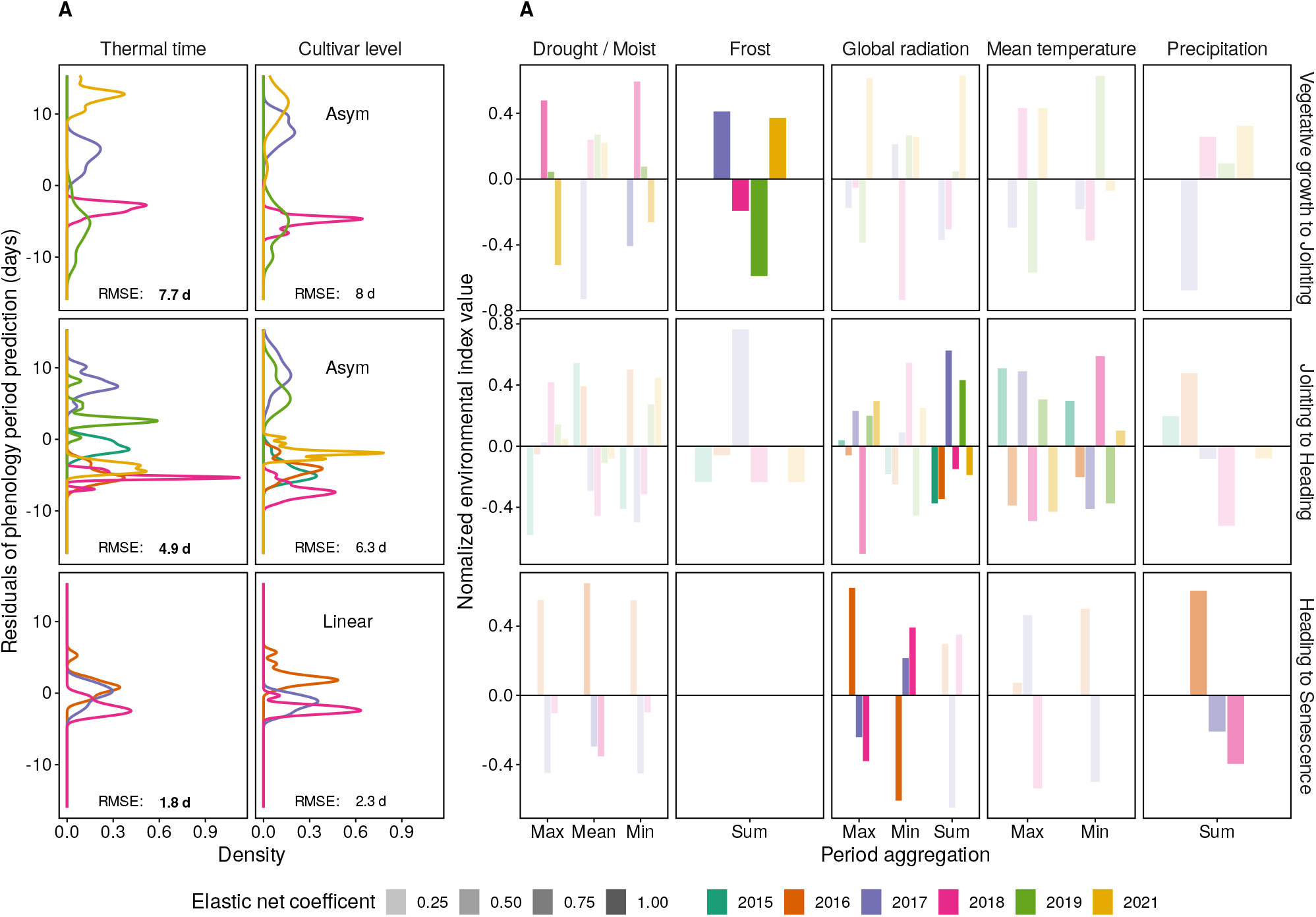
Temperature compensated phenology period duration predictions (Equation 11) for win-ter wheat, and corresponding environmental indices. Period predictions are based on genotype effects plus the mean of year effects (*g*_*i*_ + ^1^/_*j*_ Σ*v*_*j*_) estimated using a linear mixed model (Equation 12). Indicated are (a) residuals of predictions for the thermal time model and the best performing cultivar-level model (asymptotic and linear) and related root-mean-squared error (RMSE), and (b) environmental index values per year for indices that were suggested by an elastic net regression to best explain residuals of cultivar-level models. Residuals (a) and indices (b) with the same sign indicate a positive correlation (high index → extended period), differing signs indicate a negative correlation (high index → shortened period).

Using species-level dose-response curve models further increased estimated G *×* E variances (81– 95%) (Figure 5). Correspondingly, rank correlations between years and overall means did not improve (*r* =0.29–0.97).

In contrast, cultivar-level temperature response models resulted in the lowest G *×* E estimations for two of three phenology periods (vegetative growth, generative growth) (Figure 5a). Highest rank correlations and lowest G*×*E were found for the bi-linear model in the vegetative phase (34%, *r* =0.90–0.92) and the asymptotic model in the generative phase (38%, *r* =0.57–1.00). Differences in RMSEs to thermal time were small (≤ 1.4 days) (Figure 6).

#### 2.2.2 Explainability of G*×*E interactions with other environmental factors

To investigate the sources of the estimated G*×* E interactions after temperature-compensating time, the residuals of the phenology predictions based on genotype effects and the mean of year effects (Equation 12) were further decomposed in components related to environmental indices. For cultivar-level models, moist conditions and frost best explained differences in vegetative growth period duration values and delayed jointing (Figure 6). Extended generative growth and hence delayed heading was mainly related to high global radiation values. For maturity, wet conditions and/or extremes in global radiation best explained delayed senescence.

Using thermal time as temperature response model instead of cultivar-level models resulted in weaker relations of residuals to environmental indices. Although predictions were temperature-compensated, remaining relations to temperature indices were indicated. For the last growth period, only temperature related indices were found to be relevant. Links to drought and limited global radiation—as indicated by the cultivar-level models—were entirely missing.

## 3. Discussion

For both soybean and winter wheat, the results indicated an advantage of cultivar-specific nonlinear temperature response models if training and test sets were closely related. Using these cultivar-level models for wheat phenology predictions could reduce the observes G *×* E significantly.

The non-linearity of growth responses to temperature has long been suspected and investigated (Shaykewich 1995). Conclusively, the herein found best performing response models for winter wheat are well-known: The Wang-Engel model (Wang and Engel 1998) that performed best in predicting canopy growth is known for its ability to accurately model winter wheat growth (Wang et al. 2017). The asymptotic model that performed best in predicting plant height growth can be seen as simplified Wang-Engel model, given that temperatures do not exceed supra-optimal ranges (Roth, Piepho, and Hund 2022). In contrast, for soybean leaf growth, a neural network model performed best, indicating high degrees of freedom required to accurately predict responses. For measurements at the coarser canopy level, simpler linear and bi-linear models were more accurate.

Nevertheless, as other authors noted before (e.g., Parent, Millet, and Tardieu 2019), the superiority of cultivar-level and non-linear models was not given in all situations. In particular the transferability to other trait levels (i.e., plant organ versus canopy level) and to other growth phases (i.e., vegetative versus generative growth) appeared limited. Large confounding effects in the test sets are suspected. Physiological changes that are not directly related to plant organ growth, e.g., tillering/branching in the vegetative phase or lodging in the generative phase, may dilute cultivarspecific response signals (i.e., growth rates) on the canopy level. Consequently, simpler models such as species-level thermal time generalize better in such situations.

Not only the model choice, but also the covariate measurement level choice may enhance generalization: For winter wheat, the shoot apical meristem is located below the soil surface for half of the lifetime. In contrast, the growing tissue in soybean is above-ground. Consequently, vegetative growth for winter wheat was best predicted using soil temperature, confirming the findings of Jamieson et al. (1995). Surprisingly, for soybean, soil temperature could predict canopy growth more accurately than air temperature. We suspect that this is due to the fact that air temperature measured at a reference station is less representative of in-canopy temperature than soil temperature. Soil temperature courses may also better match the diurnal growth patterns commonly found in soybean (Kronenberg et al. 2020b).

Finally, changing cardinal temperatures with time (Porter and Gawith 1999) are an additional concern in temperature response modeling. Indeed, for the different growth periods, the choice of model and covariate level changed, and models performed inferior if not trained on the same growth period data. Nevertheless, for predictions, the division of the crop growth cycle into three consecutive growth phases—vegetative growth, generative growth, and maturity—was sufficient to accurately predict phenology as well as growth rates.

In this work, two approaches to derive cultivar-specific responses were proposes, (1) plant organ tracker devices for the vegetative growth phase, and (2) high-throughput plant height measurements for the generative growth phase. Both methods have been proven to provide reliable estimations for their respective growth phases (Mielewczik et al. 2013; Nagelmüller et al. 2016; Roth, Piepho, and Hund 2022), and this work could confirm their readiness for application. For plant organ tracker approaches, measurements in a few weeks per cultivar are sufficient, while for height data, measuring in multiple years is inevitable (Roth et al. 2022b). Unfortunately, no such method is yet available for the maturity phase, indicating that further research is needed to measure temperature responses in the late season.

Stress response related crop modeling may significantly profit from temperature response models that result in lower observed G *×* E. In our data, a clear clustering of environment means became visible when using asymptotic and linear, cultivar-level models (Figure 6a). For the vegetative growth phase, the years 2017 and 2021 were separated from other years, and differences were best explained by frost and moist conditions. Indeed, such frost events, followed by measurable leaf area reductions, were observed in the field in 2018 and 2021 (Tschurr et al. 2023). In the generative growth phase, the detected relations between heading date and global radiation are in accordance with Benaouda et al. (2022) who found temperature to be the main driver and high global radiation to be the main delayer of heading. Finally, for maturity, Anderegg et al. (2020) reported a delayed senescence in the year 2016 due to the extraordinary wet year with severe Septoria tritici blotch (STB) disease pressure, which was confirmed in our data by the relation of residuals to precipitation and global radiation.

The primary research question addressed in this study is whether to ignore or incorporate cultivarspecific *per se* temperature responses when modeling. Based on the growth rate predictions (Figure 3b), it appears reasonable to agree with Parent, Millet, and Tardieu (2019) that the use of specieslevel thermal time has a sound theoretical basis. Neglecting cultivar-specific *per se* temperature responses seems justified. However, further testing the thermal time concept on phenology data significantly weakened the soundness of thermal time—ignoring cultivar-level *per se* responses inflated the estimated G*×* E interactions (Figure 3c). Based on our results, we have to conclude that ignoring such cultivar differences creates bias in follow-up investigations of other G *×* E interactions, such as those induced by for example frost or drought stress (Figure 6). The tools for assessing cultivar-specific *per se* temperature responses in real-world field conditions are widely available now. With this study, we give implications on how and why those tools should be applied. Consequently, the theoretical concept of thermal time can be taken to the next level, which is cultivar-specific.

## 4. Material and Methods

The increase of a trait *y* related to genotype *i* with time *t* in a steady-state growth phase can be modeled using a dose-response function *r* of covariates 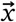 and cultivar-specific crop growth parameters 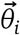 (Roth et al. 2021),

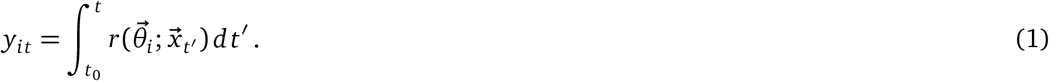

In this framework, complexity may be varied at four levels:

- The trait *y* can be measured at different scales, e.g., at plant organ level, or plant stand (canopy) level.
- The dose-response function *r* can vary in complexity, e.g., using linear regressions, nonlinear regressions, semi-parametric splines (Pérez-Valencia et al. 2022), or neural network regressions.
- The covariates 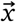 can be measured at different scale, e.g., close to the growing meristem, at the experimental unit (plot) level, or at a reference station.
- The crop growth parameters 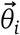 can be estimated at different scale, e.g., at variety/genotype level *i*, or at species-level.

In the following, we pursue this structure, describing how traits were measured, complexity varied, and growth and phenology modeled.

Note that in Equation 1, 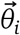was defined as cultivar-specific parameter set. As such, the doseresponse function *r* will both model response *per se* (e.g., the base temperature below growth stops) and intrinsic growth rate differences between cultivars (e.g., absolute growth at optimum temperature) (Figure 1f). When replacing 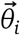 with a species-level parameter set 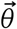, *r* reduces to a function that models relative growth rates. Thermal time is one example of such a function. To scale these relative growth rates to cultivar-specific intrinsic growth rates, one has to scale *r* to *y*_*it*_ using a cultivar-specific factor *g*_*i*_ (Figure 1c),

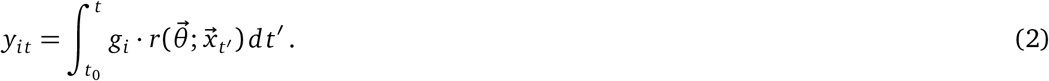

To allow a comparison of species-level and cultivar-level models, Equation 1 was used for cultivar-level models, and Equation 2 with the cultivar-level parameter *g*_*i*_ for species-level models. For further details, please see Equation 8, 9, and 12.

### 4.1 Material

All experiments were performed at the ETH research station for plant sciences Lindau-Eschikon, Switzerland (‘Eschikon’; 47.449 °N, 8.682 °E, 520 m a.s.l.) on the field of the field phenotyping platform ‘FIP’ (Kirchgessner et al. 2017). The wheat experiment comprised a set of 12 varieties (CH Claro, CH Nara, Fastnet, Marksman, Ostka Strzelecka, Romanus, Runal, Rywalka, Semafor, Tamaro, Toronto, Winnetou), replicated 2 times per year, and cultivated in 2015–2021 as subset of a larger experiment with approximately 350 genotypes (Kronenberg et al. 2017; Kronenberg et al. 2020a; Roth et al. 2020). For leaf elongation tracking, the 12 varieties were additionally grown in four small plots beside the main experiment in 2019. These plots (0.9 m x 1 m) contained three cultivars each, and were sown by hand in stripes of 0.3 m x 1 m.

The soybean experiment comprised a set of 3 varieties (Castetis, Gallec, Opaline), replicated 3 times per year, cultivated in 2017–2020 as subset of a larger experiment with 36 genotypes (Roth et al. 2022a). Leaf growth tracking was performed directly in the main experiment in 2017.

### 4.2 Trait measurements

#### 4.2.1 Leaf length tracking in winter wheat

Leaf elongation rates of 12 wheat cultivars were measured in the field from mid-February to beginning of April 2019 using the leaf length tracker (LLT) system described by Nagelmüller et al. (2016). The installation followed the principle of an auxanometer. Briefly, the youngest leaf was attached to a hairpin to which a thread was attached. The thread was guided over several rollers along the panel and held taut with a counter weight (20 g). At the other end, the thread was attached to a white bead that moved over the panel in accordance with the elongation of the leaf. A waterproof CCTV camera (Lupusnet HD LE934, CMOS sensor, maximal resolution of 1920 × 1080 pixels, Lupus-Electronics® GmbH, Germany) took images of the panel every 2 minutes. A custom software (https://sourceforge.net/projects/leaf-length-tracker/) evaluated the position of the beads from the pictures, as they were used as indirect artificial landmarks to measure leaf elongation rate. Measurements were performed on in average 6 replications over 7 weeks (Figure 2a).

Measured growth rates were corrected for weight-temperature interaction effects based on a calibration performed in a climate chamber. In this calibration setup, the growth of undisturbed leaves was compared with the growth of leaves where a force-equivalent of 20 g was applied. The differences in measured growth rates suggested a cultivar-unspecific correction of 0.004 mm/h per °C.

#### 4.2.2 Leaf growth tracking in soybean

Leaf growth rates of 3 soybean cultivars were measured in the field from the beginning of June to mid-July 2017 using the leaf growth tracker (MARTRACK) system described by Mielewczik et al. (2013). Briefly, beads connected to threads were glued to the emerging leaves and fixed in front of a camera using a wire frame. The same cameras as above were used to record images every 2 minutes. A custom software (https://sourceforge.net/projects/martrackleaf/) evaluated the position of the beads from the pictures. Leaf area was then calculated based on the convex hull of bead positions in the planar image space. Relative growth rates were calculated based on differences of logarithmic leaf areas of two successive time points, divided by the time difference. Measurements were performed on in average 3 replications over 4 weeks (Figure 2a).

#### 4.2.3 Canopy cover monitoring based on RGB imaging in winter wheat and soybean

Canopy cover increase was monitored using the high-throughput field phenotyping platform ‘FIP’. The FIP platform is–among other sensors—equipped with an RGB camera (EOS 5D Mark II, 35 mm lens, Canon Inc., Tokyo, Japan). Plots were monitored with this sensor from a distance of 3 m to the ground. This setting results in a ground sampling distance of 0.3 mm/pixel. In the early canopy growth phase, in average 10 measurements per year (2017–2019, 2021) were taken for winter wheat, and in average 5 measurements per year (2017–2018, 2020) for soybean. RGB images were segmented pixel-wise into a plant and a soil fraction using a deep convolutional neural network (Zenkl et al. 2022).

To enable a pixel-precise extraction of plot canopy cover values, image time series were first aligned using planar homography. Then, plot-specific shapes were projected to image time points. As feature detection algorithm, SIFT (Lowe 1999) and ORB (Rublee et al. 2011) were used. Feature matching was performed using RANSAC (Fischler and Bolles 1981). Subsequently, the segmented and cutout image parts showing individual plots were further rectified by rotating them step-wise (-1.5° to 1.5° in steps of 0.2°) to maximize the distance between the minimum and maximum of plant pixels in image columns. For canopy cover extraction in winter wheat, only the inner 7 rows (of 9 rows per meter) were considered. For soybean with larger row spacing (3 rows per meter), only the inner row and half of both outer rows were used for further processing.

Canopy cover was then calculated as plant pixel ratio per plot. For winter wheat, measurements between approximately the beginning of the year to mid-April were considered, for soybean, measurements between approximately mid-May and end of June. Only positive values, i.e., only canopy increase, was used for further processing. All processing was performed in Python using OpenCV and scipy (Virtanen et al. 2020).

#### 4.2.4 Plant height monitoring in winter wheat and soybean

Plant height increase was monitored using the high-throughput field phenotyping platform ‘FIP’ as well as drones. The FIP platform is–among other sensors—equipped with an terrestrial laser scanning device. The first three years of plant height measurements in winter wheat were collected with this device (Friedli et al. 2016; Kronenberg et al. 2017; Kronenberg et al. 2020a). From the resulting point clouds, the percentile best matching manual measurements (97*th* percentile, Kronenberg et al. 2017), was extracted per plot as plant height estimation per time point. For the subsequent years of winter wheat experiments and for all soybean experiments, drone-based Structure-from-Motion (SfM) was used (Roth and Streit 2018; Roth et al. 2022b). From the resulting point clouds, the percentile best matching manual measurements (90*th* percentile, Roth and Streit 2018) per plot was extracted as plant height estimation per time point.

For wheat, measurements were performed on 2 replications on in average 11 time points per year (2015–2019, 2021) that fell into the stem elongation phase (Figure 2c). For soybean, measurements were performed on 3 replications on in average 8 time points per year (2017–2020) that fell into the stem elongation phase (Figure 2c). This included measurements between approximately mid-April and end of May for wheat and mid-June to mid-July for soybean.

Plot time series were smoothed using P-splines with the R package *scam* (Pya 2019) before further processing to reduce prediction errors origin from autocorrelations of measurement errors.

#### 4.2.5 Phenology measurements and estimations in winter wheat

Heading and senescence measurements were performed manually by trained persons. Heading was defined as the time point when the inflorescence was fully emerged for ≥ 50% of all shoots (GS 59) (Meier 2018). Heading was measured for all years (2015–2019 and 2021) on 1–2 replications.

Senescence was defined as the time point where the senescence of the central plot area has reached its midpoint (Anderegg et al. 2020). Senescence was assessed on two replicates in 2016, 2017 and 2018.

The start of the stem elongation was estimated for genotypes in two replications based on plant height data using the quarter-of-maximum-elongation rate method (QMER) described in Roth et al. (2021) and Roth et al. (2022b) for the years 2017–2019 and 2021. For 2015 and 2016, no detailed plant height data for the early season were available, wherefore the start was approximated for all genotypes alike (2015-04-28 and 2016-04-15).

### 4.3 Covariate measurements

Reference air temperature 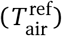 at the local weather station (in close proximity to the experimental field) was measured above a grass strip at 0.1 m above ground using Campbell CS215 sensors (Campbell Scientific Inc., U.S.A.). Air temperature inside the experiment 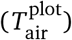 was measured in 2–4 wheat plots at 0.1 m above the ground, therefore above the plants before the start of the stem elongation, and inside the canopy for later growth stages, using Campbell CS215 sensors. Reference soil temperature 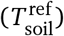 was measured 0.05 m below ground at three reference positions below grass strips using Sentek/Hydrolina soil sensors (Sentek Sensor Technologies, Australia). Soil temperature inside the experiment 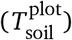 was measured 0.05 m below ground in 2–4 wheat plots using using Sentek/Hydrolina soil sensors. Values of measurements performed at multiple locations (plots or reference positions) were averaged.

### 4.4 Growth modeling

#### 4.4.1 Dose-response models

As baseline model, thermal time based on hourly temperature recordings was used,

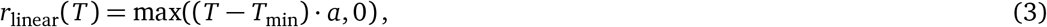

where *T*_min_ is the base temperature of growth, *a* the slope, and max(, 0) prevents negative growth rate predictions by replacing values lower than zero with zero. This model was called ‘thermal time’ if *a* = 1 and *T*_min_ was set to the literature based threshold temperature of 0 °C for winter wheat (Baker and Gallagher 1983) and 5 °C for soybean (Whigham and Minor 1978), and ‘linear model’ if *T*_min_ and *a* were estimated based on own data.

To account for lower growth rates at temperatures close to zero that were observed in LLT data, the linear model was extended to a bi-linear model,

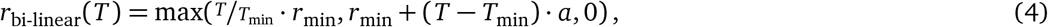

where *r*_min_ is a growth rate ≥ 0 at *T*_min_.

Plant height growth rate modeling has shown that an asymptotic model can approximate a WangEngel model given that temperatures do not exceed to supra-optimal growth ranges (Roth, Piepho, and Hund 2022). The asymptotic models is defined as

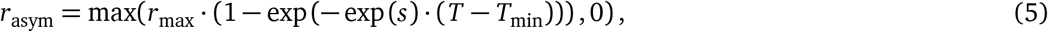

where *r*_max_ is the maximum absolute growth rate (and therefore the asymptote of the curve), *T*_min_ the base temperature where the growth rate is zero, and *s* characterizes the steepness of the response (natural logarithm of the rate constant, thus ‘*lrc*’) (Pinheiro and Bates 2000).

Finally, the original Wang-Engel model (Wang et al. 2017) is defined as

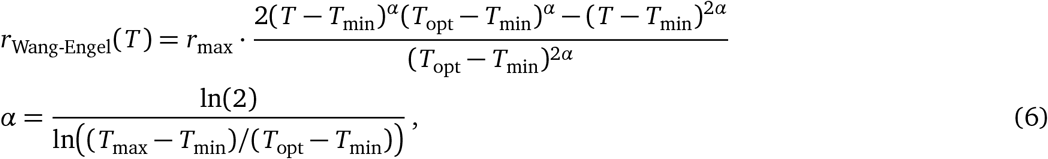

where *r*_max_ is the maximum absolute growth rate at the temperature optimum *T*_opt_, *T*_min_ the lower base temperature and *T*_max_ the upper base temperature of growth.

#### 4.4.2 Model fitting to temporally resolved leaf growth tracking device data

Before training models, the data were split in training and test sets using a ratio of 6.37:1 for wheat and 6.31:1 for soybean, taking care that time series of replications/leaves stayed together in either the training or test set. The linear and non-linear models defined above were then fitted to training data using maximum likelihood optimization. In contrast to previous attempts to process leaf growth tracking data (Mielewczik et al. 2013; Nagelmüller et al. 2016), the raw measurement data were not smoothed. Instead, the measurement error was estimated using nested models with residual autocorrelation of order 1–3. The best fitting model was selected based on the Bayesian Information Criterion (BIC). Models were fitted using the base R (R Core Team 2019) function *mle*. In addition to the parametric linear and non-linear models, two so called ‘semi-parametric’ models were trained. The first one was based on a hierarchical spline approach for longitudinal data (Durbán et al. 2005) that models a general population trend, a genotype trend, and a replication trend (Pérez-Valencia et al. 2022, R-code herein provided). We modified this approach in the way that time *t* was replaced by temperature *T* as ‘longitudinal dimension’, thus fitting hierarchical splines that represent dose-response curves.

The second ‘semi-parametric’ model approach was based on a multi-output neural network: A small network with two hidden layers of size 5 and sigmoid activation functions was trained to regress temperature on growth rates. An additional layer was then added to the network that transformed the single-output in a multi-output of size 12 (one for each genotype). This layer was not activated, thus representing a linear transformation only. Training and validation sets were split with a 9:1 ratio, loss was calculated as mean squared error (MSE). L1 regularization was applied to the last layer. Optimization was done using the Adam optimizer in Pytorch Lightning with 1500 epochs in pre-training and 800 epochs for fine-tuning with early stopping if the MSE did not improve by more than 0.0001 for 40 epochs. Initial learning rate was 0.05 with exponential decay with gamma 0.996, precision was 16 bit (half-precision), batch size 2000. Early stopping was always reached.

#### 4.4.3 Model fitting to canopy cover and height data

As for the leaf growth tracking data, canopy cover and height data were split in a training and test set. Here, the split was performed based on whole years, and repeated in a cross-validation (CV) scheme. For wheat, this resulted in a 5:1 split ratio for plan height data in a 6-fold CV. For soybean, this resulted in a 3:1 split ratio for plan height data in a 4-fold CV.

The parametric models were then fit to the training set using a maximum likelihood approach that can fit high-resolution (hours) temperature courses and low-resolution (days) trait measurements (Roth, Piepho, and Hund 2022).

An attempt to use canopy cover data to train models resulted in very poor estimates or failed convergence. As a consequence, canopy cover data were only used for model testing and not for fitting.

### 4.5 Model testing for growth predictions

Leaf growth tracking data originated from the same year. Consequently, the train/test split was used to calculate performance values by pooling all measurement values of the test set per cultivar. All other data were collected in differing years. Therefore, a random regression model was used for model testing. This approach was chosen based on the longitudinal character of plot-based time series, where one has to expect temporally and spatially correlated measurement errors (Roth et al. 2021).

For such measurements, a trait *y* is measured at repeated times *t* for genotypes *i* in the year *j* at the replication *k*. Time *t* is ‘linearized’ using the different dose-response models, 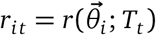. Consequently, the difference between two consecutive measurements can be expressed as

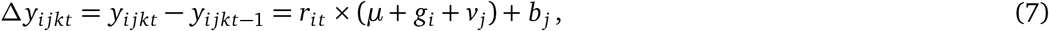

where *μ* is a fixed overall coefficient, *g*_*i*_ and *v*_*j*_ random coefficients related to genotypes and years, and *b*_*j*_ a year-specific offset. The random coefficient structure was estimated using a variancecovariance structure among genotype replications *g*_*i*_ and years *v*_*j*_ . The model was fitted in R using *ASReml-R* (Butler 2018).

The reported coefficients of determination of the predictions (R^2^ score) and root-mean-squared errors (RMSE) were based on the fixed overall coefficient *μ* for cultivar-level models,

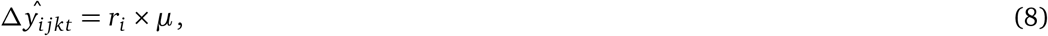

and on the fixed overall coefficient *μ* and random genotype coefficient *g*_*i*_ for species-level models and thermal time,

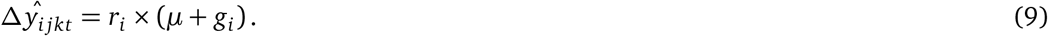

The R^2^ score was defined as

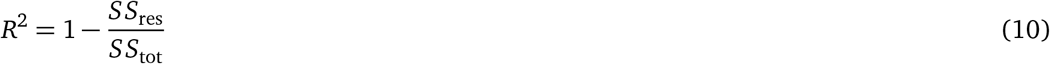

where *SS*_tot_ the total sum of squared and *SS*_res_ is the residual sum of squares in relation to the 1:1 line.

### 4.6 Model testing for phenology predictions

To test the prediction ability of phenology period duration, a linear mixed model was used. Such an approach can account for random sources of variation such as genotype effects and G *×* E interactions (Piepho, Büchse, and Emrich 2003). For phenology timing periods, two time points *t*_1_ and *t*_2_ are measured per genotype *i* in the year *j* at the replication *k*. Then, the time period in between is ‘linearized’ using the different dose-response models, resulting in a new trait *y*,

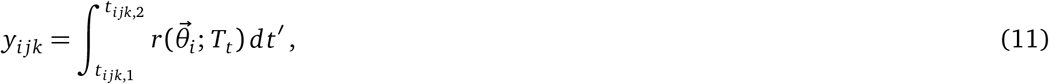

where *t*_*i jk*,1_ is the start of the period and *t*_*i jk*,2_ the end of the period. Note that the new trait *y*_*i jk*_ is on a genotype-specific scale. To allow comparison and variance decomposition, *y*_*i jk*_ values were scaled to one per genotype. After this time period transformation, overall best linear unbiased predictions (BLUPs) were estimated using the model

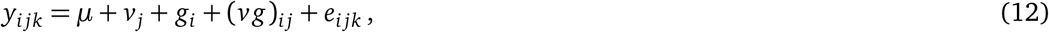

where *μ* is a global intercept, *v*_*j*_ a fixed year-effect, *g*_*i*_ a random genotype effect, and (*vg*)_*i j*_ random genotype-environment effects modeled using a diagonal variance structure, allowing for differing genotypic variances for years. The residual variance structure *e*_*i jk*_ was set to account for random row and range effects and random interactions of row and range effects, thus accounting for differing spatial gradients for years. The model was fitted in R using *ASReml-R* (Butler 2018).

#### 4.6.1 Residual analysis based on environmental indices

Environmental indices were calculated based on daily mean, minimum, and maximum temperature, precipitation sum and global radiation (Supplementary materials, Table B.1). Precipitation values were further transformed into the Standardised Precipitation Index (SPI) (McKee, Doesken, and Kleist 1993) and the Standardised Precipitation and Evapotranspiration Index (SPEI) (Vicente-Serrano, Beguería, and López-Moreno 2010) using the Thornthwaite transformation (Thronthwaite 1948) to estimate evapotranspiration. SPI and SPEI were calculated using the R package *SPEI* (Beguería et al. 2014). Both indices were calculated with a 30-day smoothing to account for effects within each phenological period rather than long-term effects. For each index, the minimum, maximum and cumulative values per phenological period were calculated. Furthermore, a cold stress index considering the temperature sum of minimum daily temperatures below 0 °C was added. Drought and moisture extreme indices were calculated using the sum of SPI and SPEI values above and below a threshold of 1 and 1.75, respectively -1 and -1.75. Negative values represent very moist periods and can therefore define wet seasons. Positive values indicate dry periods and can therefore correlate with periods of drought stress. In addition to the environmental indices, the mean growth period duration in days per cultivar was added to the list of features. A lasso regression was then applied to the residuals of the phenology prediction models, using a lasso and elastic-net regularized generalized linear model from the R package *glmnet* (Friedman et al. 2022). The model was fitted using the R package *caret* (Kuhn 2008) with a search grid for *λ* = 10^−8^ to 5 and *α* = 1 in a repeated CV with 10 repeats and 5 folds. Features were centered and scaled before fitting.

## A. Appendix

**Figure A.1:**
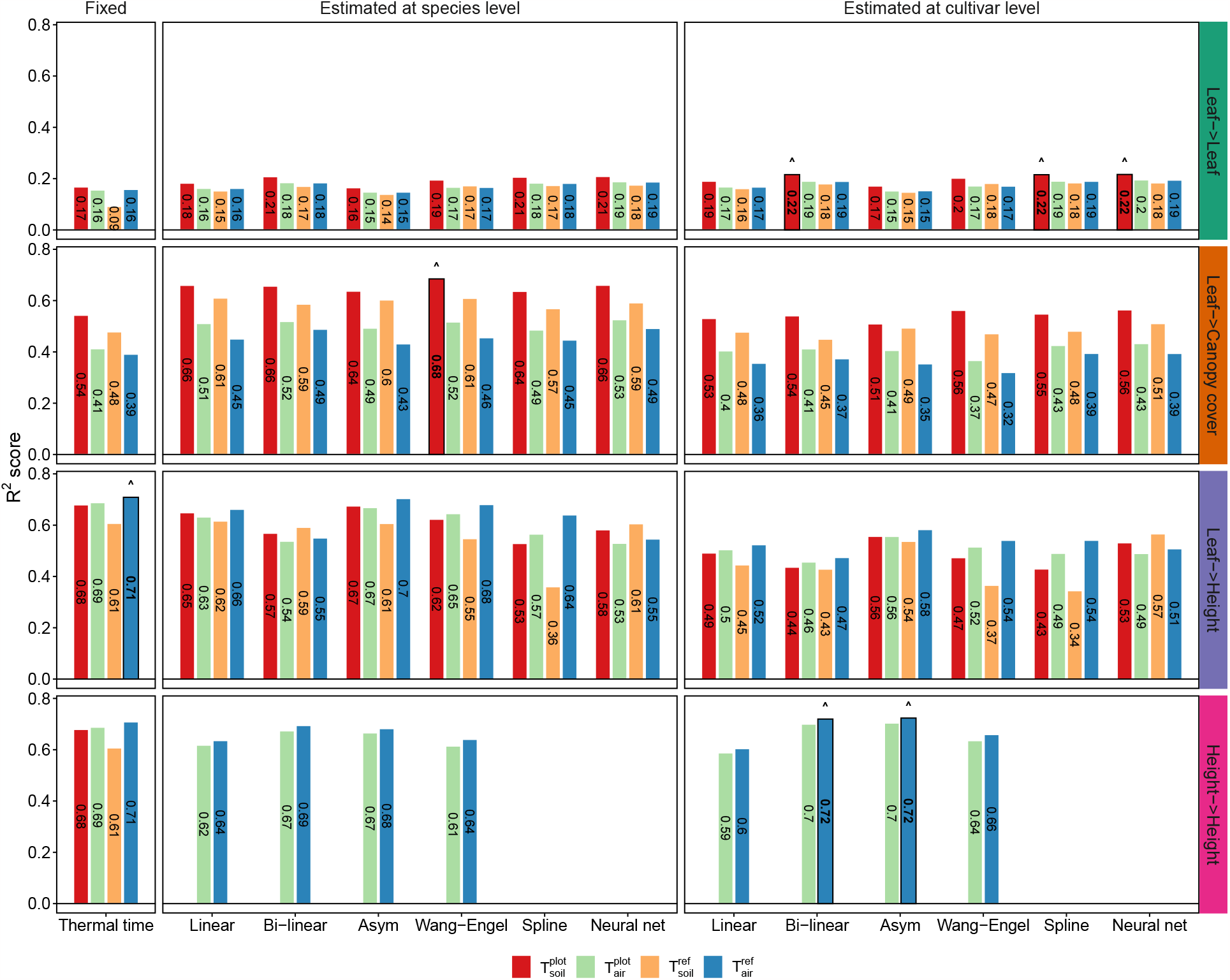
Performance of growth rate predictions for winter wheat. Models were trained on leaf length data (Leaf->) and plant height data (Height->). Predictions were tested on unseen leaf length data (->Leaf), canopy cover data (->Canopy cover), and plant height data (->Height). The covariate temperature was measured in air (T_air_) and soil (T_soil_) at plot level (T^plot^) and at a reference station (T^ref^). Indicated are the coefficients of determination (R^2^ score) of predictions based on the corresponding temperature response model. At the species-level, predictions were based on an overall coefficient and genotype coefficients (Equation 9). For cultivar-level models, the genotype specificity is already incorporated in the response model, and predictions therefore based on an overall coefficient only (Equation 8). Coefficients were fitted using a random regression model with random coefficients for years and plots and the mentioned fixed overall coefficient and genotype coefficients (Equation 7).

**Figure A.2:**
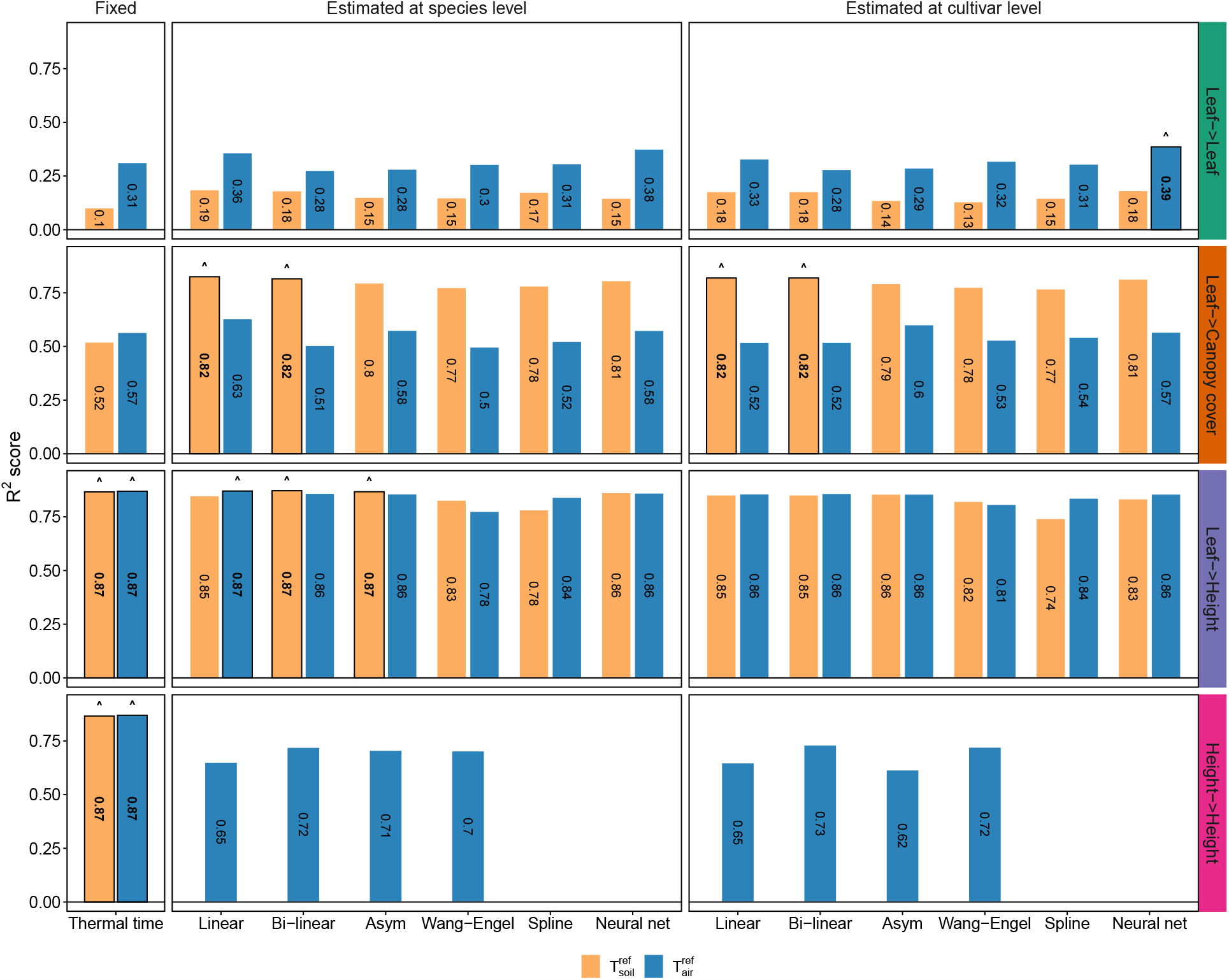
Performance of growth rate predictions for soybean. Models were trained on leaf area data (Leaf->) and plant height data (Height->). Predictions were tested on unseen leaf area data (->Leaf), canopy cover data (->Canopy cover), and plant height data (->Height). The covariate temperature was measured in air (T_air_) and soil (T_soil_) at a reference station (T^ref^). Indicated are the coefficients of determination (R^2^ score) of predictions based on the corresponding temperature response model. At the specieslevel, predictions were based on an overall coefficient and genotype coefficients (Equation 9). For cultivar-level models, the genotype specificity is already incorporated in the response model, and predictions therefore based on an overall coefficient only (Equation 8). Coefficients were fitted using a random regression model with random coefficients for years and plots and the mentioned fixed overall coefficient and genotype coefficients (Equation 7).

**Figure A.3:**
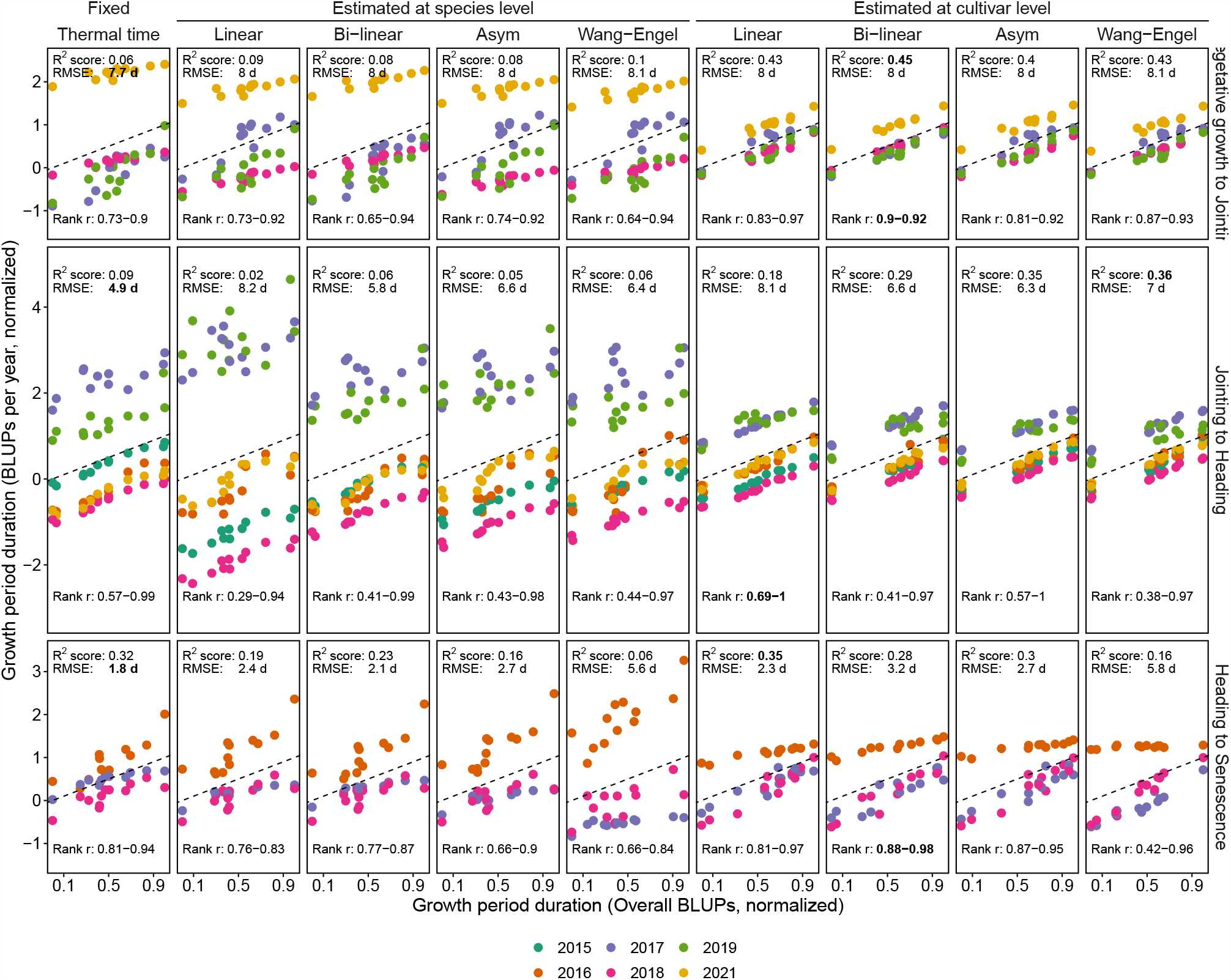
Performance of phenology period duration predictions for winter wheat. Predictions were based on cultivar-level best linear unbiased estimations (overall BLUPs) of a linear mixed model that included effects for cultivars, years, and year-cultivar interactions (Equation 12). The linear mixed model itself was fitted to thermal time, cultivar-level temperature response model, and species-level temperature response model outputs from growth rate fits.

## Acknowledgments

We acknowledge Hansueli Zellweger and Simon Corrado for field management at the FIP site. Furthermore, we acknowledge the support in data collection of Jonas Anderegg, Phillip Braun and Moritz Affentranger (ETH Zurich).

A.W. discloses support for the research of this work from Swiss National Science Foundation [grant number 169542] and Swiss National Science Foundation [grant number 200756].

All authors were involved in designing the research. LR, MB, NK, FT, and LK performed the research. LR performed the analysis and wrote the first manuscript draft. All authors contributed to the final manuscript.

The authors declare that they have no competing interests. All data needed to evaluate the conclusions in the paper are present in the paper, the Supplementary Materials, and on https://gitlab.ethz.ch/crop_phenotyping/phenoflow_early_growth.

