## Supplementary Figures and Tables for "From neglecting to including cultivar-specific *per se* temperature responses: Extending the concept of thermal time for plant development modeling"

### B. Supplementary Materials

#### B.1. Figures

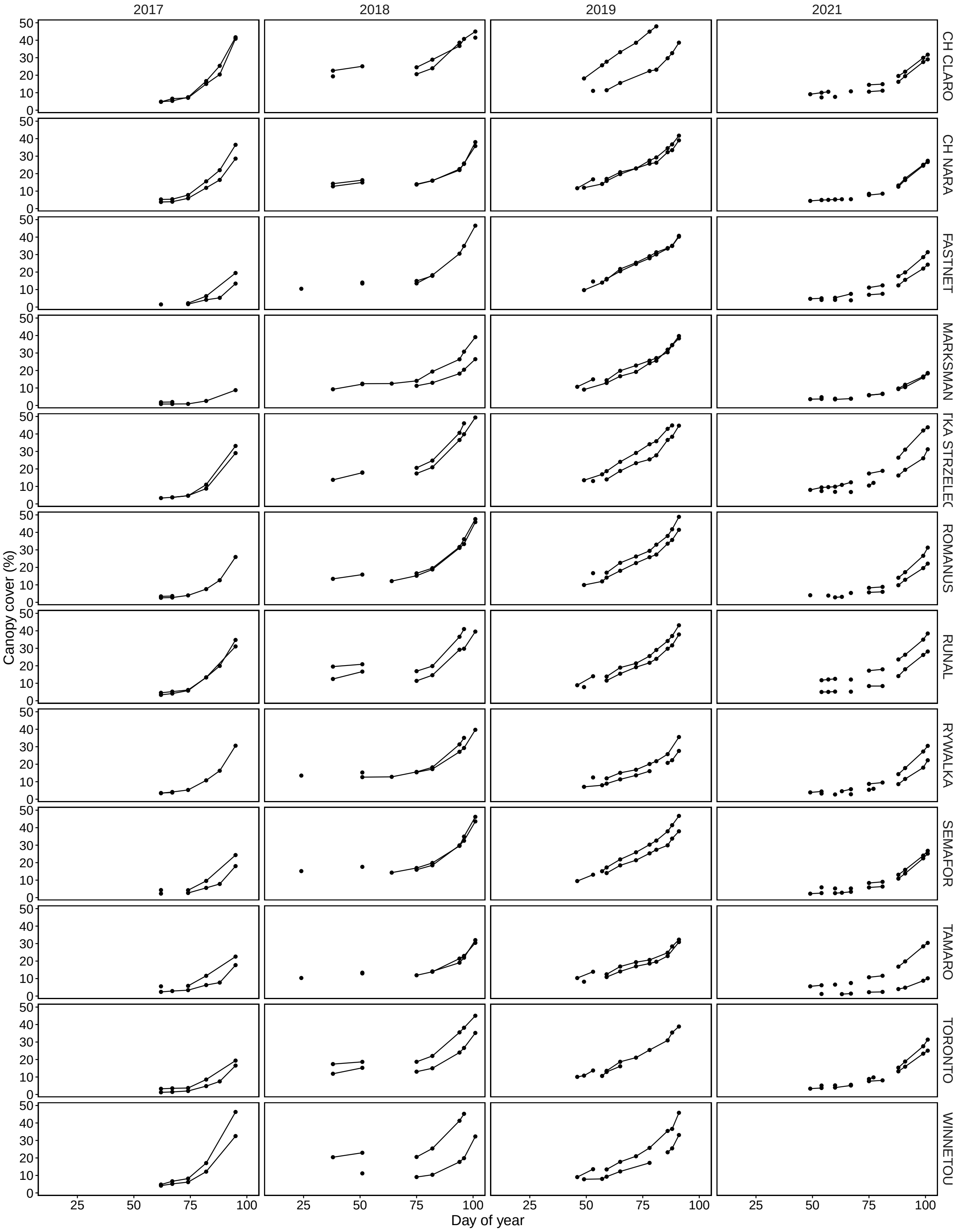


Figure B.1: Canopy cover measurements in wheat with the FIP (FIP RGB)

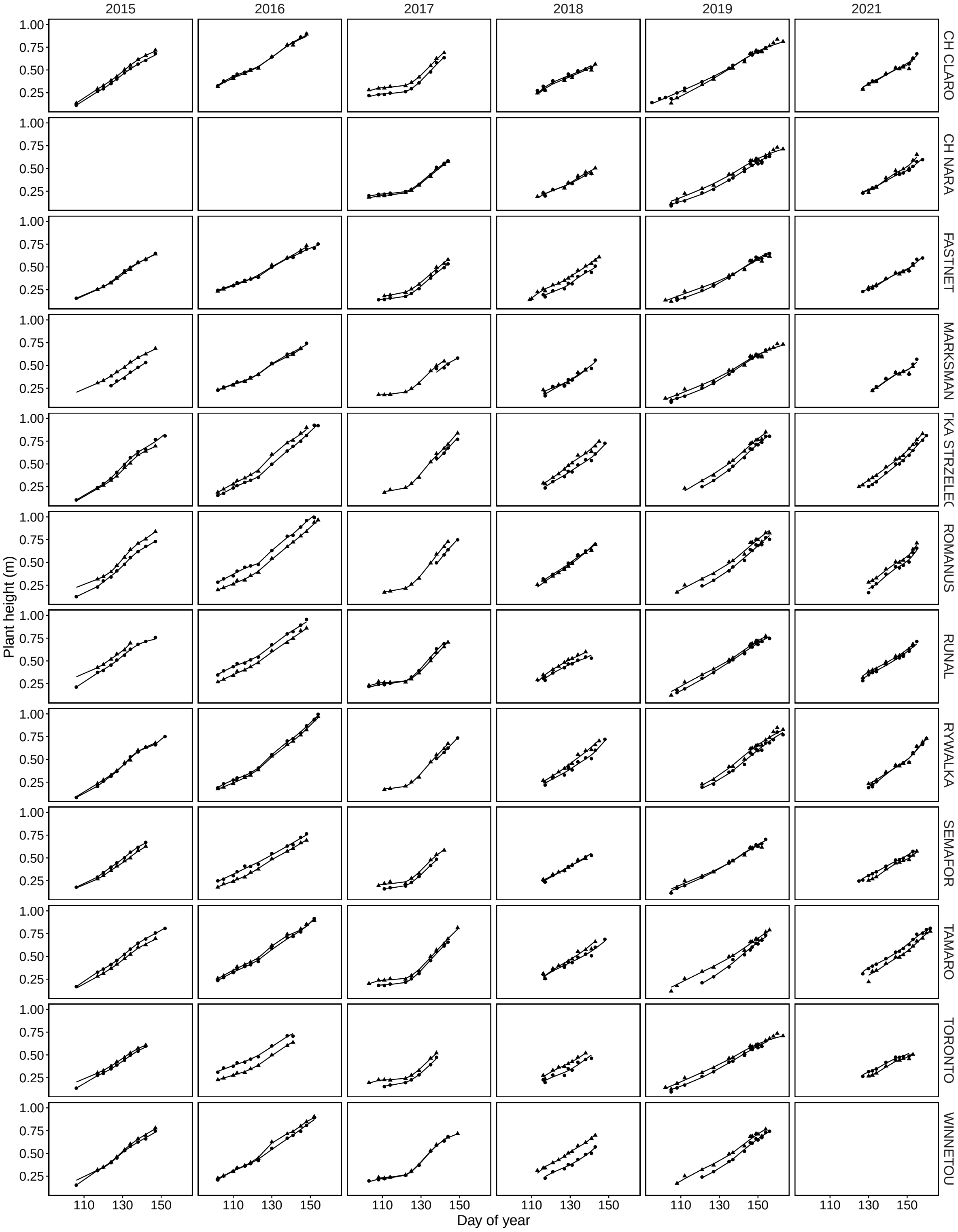


Figure B.2: Plant height measurements in wheat with the FIP (FIP TLS, 2015–2017) and drones (UAV SfM, 2018–2021), smoothed with a P-spline.

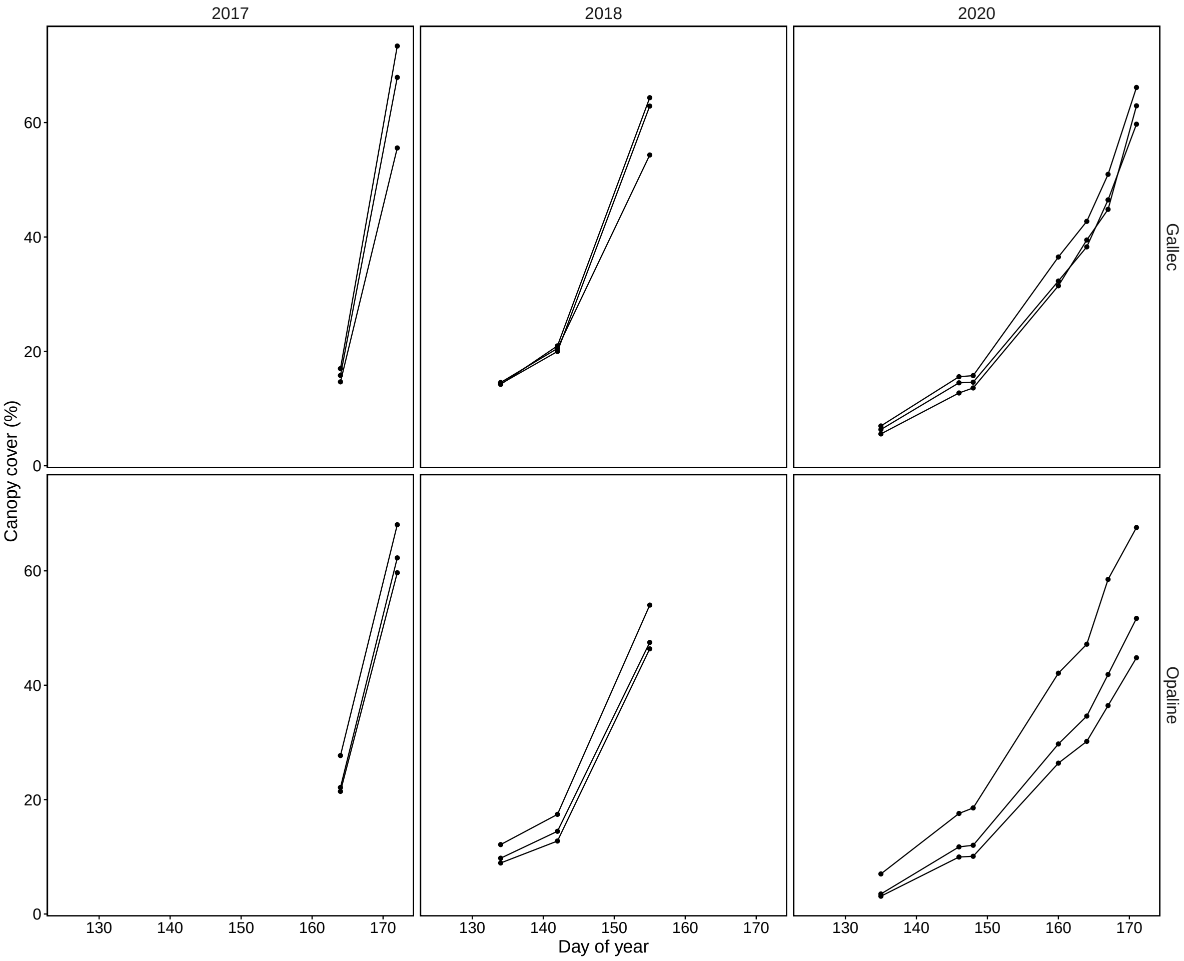


Figure B.3: Canopy cover measurements in soybean with the FIP (FIP RGB)

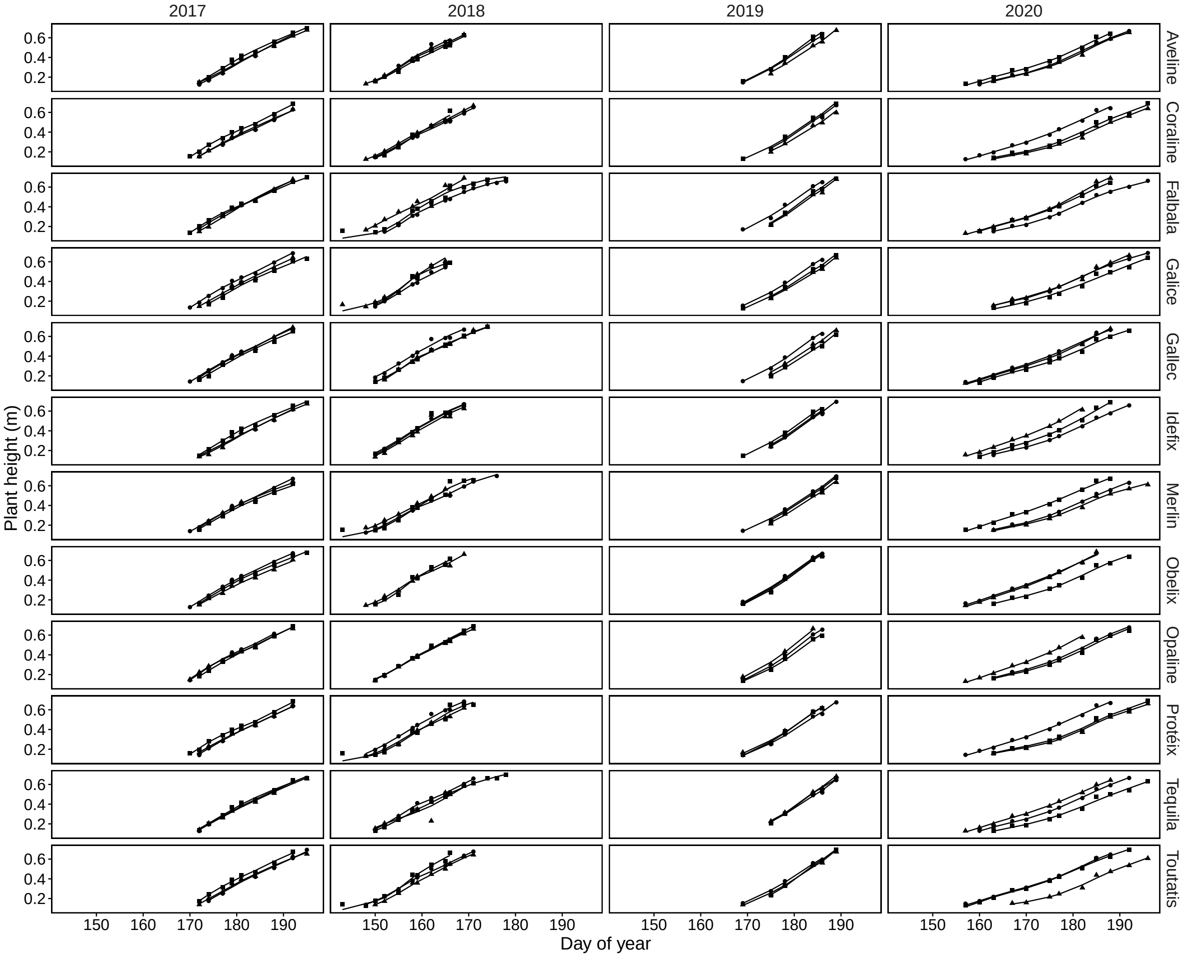


Figure B.4: Plant height measurements in soybean with drones (UAV SfM, 2017–2020), smoothed with a P-spline.

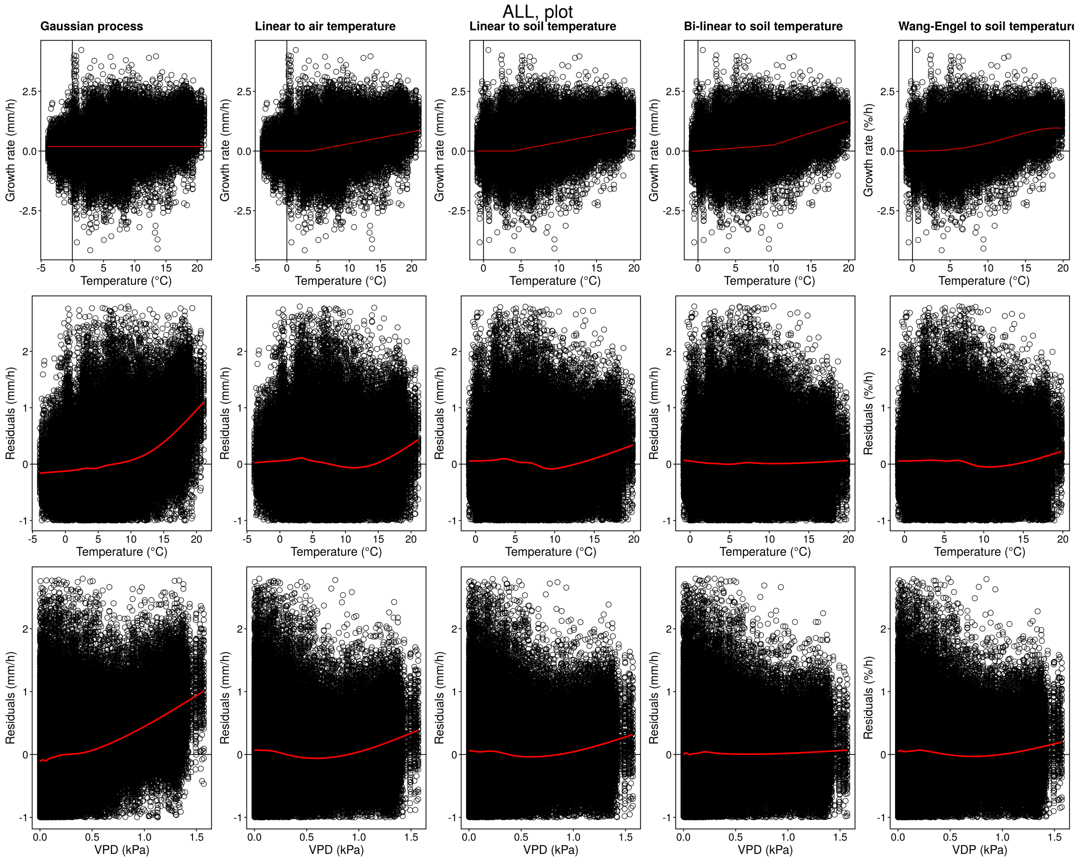


Figure B.5: Leaf length measurements in wheat with the LLT, all 12 genotypes, selected parametric models.

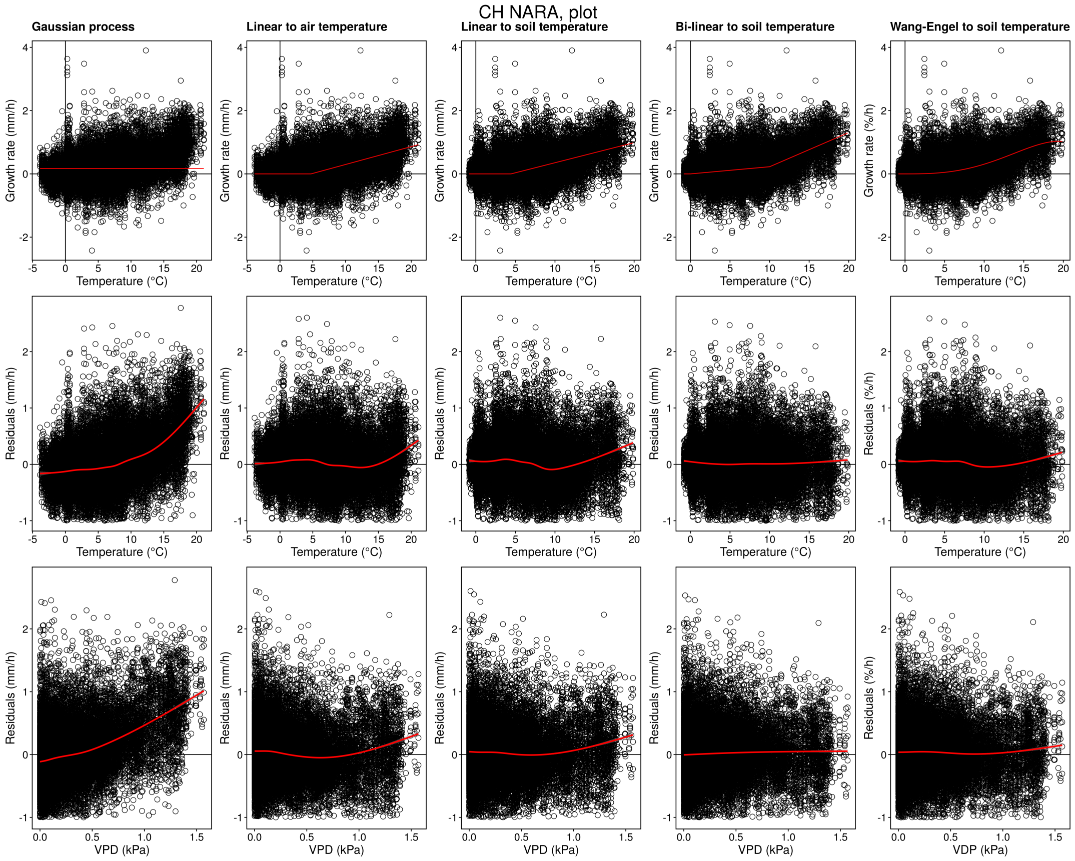


Figure B.6: Leaf length measurements in wheat with the LLT, selected genotype (CH Nara), selected parametric models.

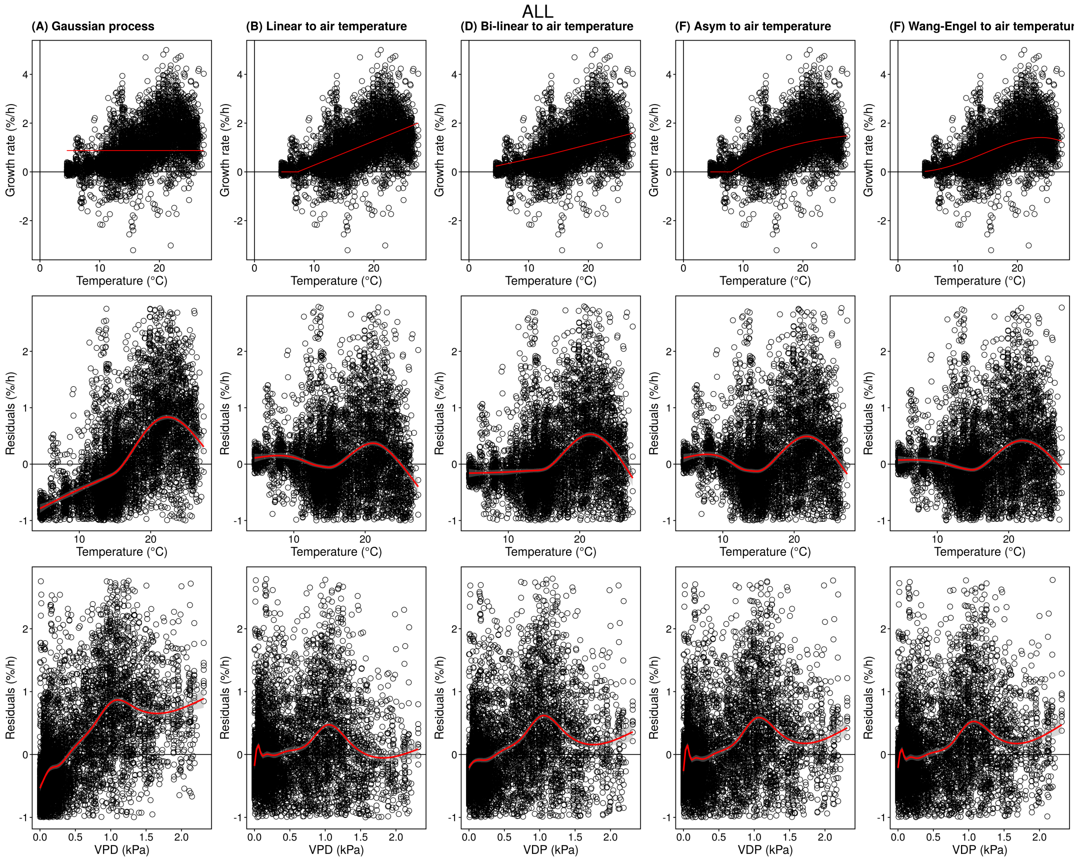


Figure B.7: Leaf growth measurements in soybean with MARTRACK, all 3 genotypes, selected parametric models.

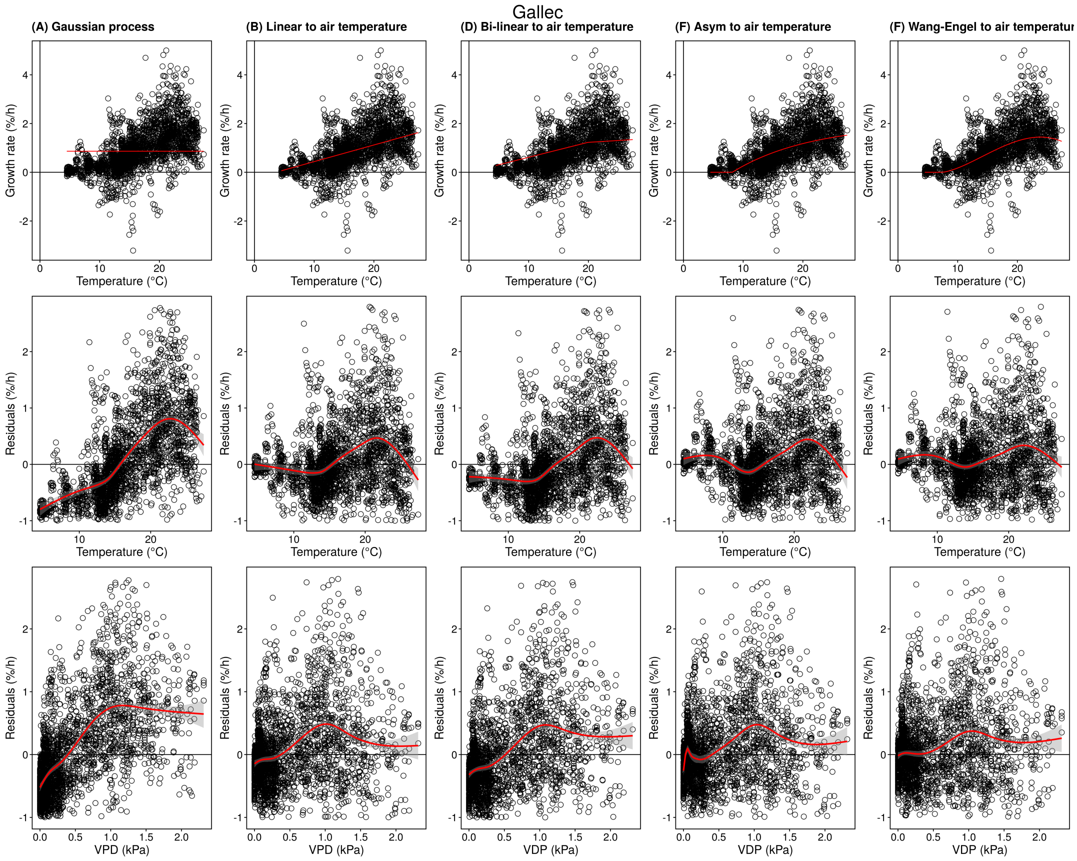


Figure B.8: Leaf growth measurements in soybean with MARTRACK, selected genotype (Gallec), selected parametric models.

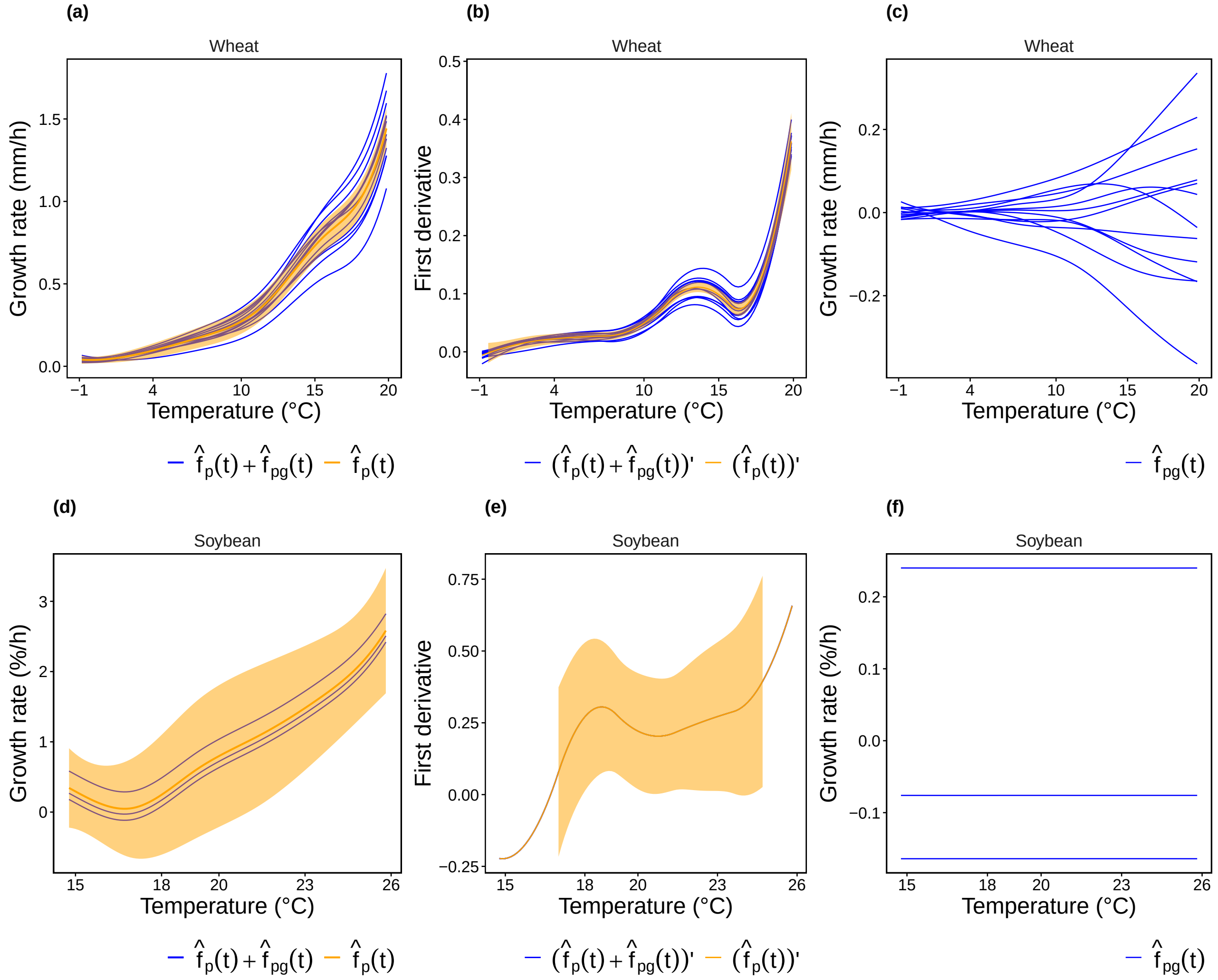


Figure B.9: Fitted hierarchical splines for leaf length tracker (LLT) measurements in wheat and leaf growth tracker (MARTRACK) measurements in soybean.

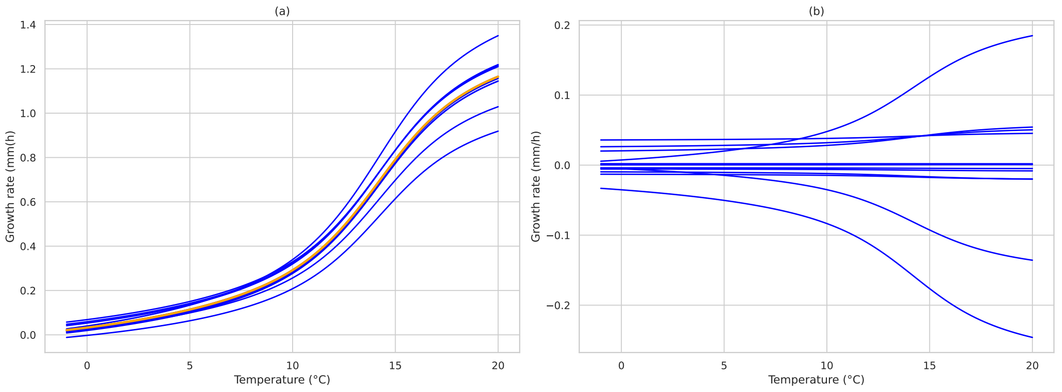


Figure B.10: Fitted neural network model for leaf length tracker (LLT) measurements in wheat, regression to plot soil temperature.

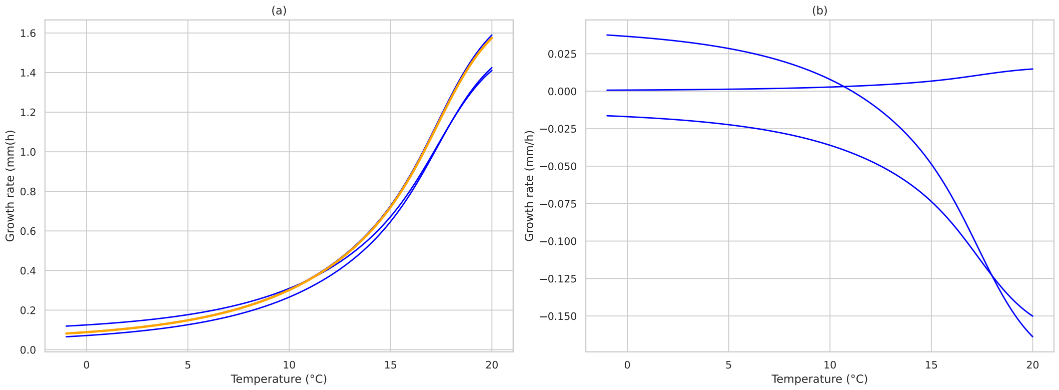


Figure B.11: Fitted neural network model for leaf growth tracker (MARTRACK) measurements in soybean, regression to reference air temperature.

#### B.2. Tables

Table B.1: Environmental indices and categories.

| | Environmental variable | Daily | Growth period | Category |
|---|---|---|---|---|
| pr_mean | Precipitation | Sum | Mean | Precipitation |
| pr_max | Precipitation | Sum | Maximum | |
| pr_cum | Precipitation | Sum | Sum | |
| tas_mean | Temp. | Mean | Mean | Mean temp. |
| tas_min | Temp. | Mean | Minimum | |
| tas_max | Temp. | Mean | Maximum | |
| tas_cum | Temp. | Mean | Sum | |
| tasmin_mean | Temp. | Minumum | Mean | Min temp. |
| tasmin_min | Temp. | Minumum | Minumum | |
| tasmin_cum | Temp. | Minumum | Sum | |
| tasmax_mean | Temp. | Maximum | Mean | Max temp. |
| tasmax_max | Temp. | Maximum | Max | |
| tasmax_cum | Temp. | Maximum | Sum | |
| global_radiation_mean | Global radiation | Sum | Mean | Global radiation |
| global_radiation_min | Global radiation | Sum | Min | |
| global_radiation_max | Global radiation | Sum | Maximum | |
| global_radiation_cum | Global radiation | Sum | Sum | |
| SPI_mean | Precipitation/Temp. | Sum/Mean | Mean | Drought / Moist |
| SPI_min | Precipitation/Temp. | Sum/Mean | Min | |
| SPI_max | Precipitation/Temp. | Sum/Mean | Maximum | |
| SPI_cum | Precipitation/Temp. | Sum/Mean | Sum | |
| SPEI_mean | Precipitation/Temp. | Sum/Mean | Mean | |
| SPEI_min | Precipitation/Temp. | Sum/Mean | Min | |
| SPEI_max | Precipitation/Temp. | Sum/Mean | Maximum | |
| SPEI_cum | Precipitation/Temp. | Sum/Mean | Sum | |
| tasmin_sum_below_0 | Temp. | Minimum | Sum(Abs) if $< 0$ °C | Frost |

Table B.2: Fitted non-linear models to leaf length tracker (LLT) wheat data. Genotype=ALL denotes the species-specific models.

| genotype_name | model | covariate_suffix | AIC | BIC | RMSE | Tmin | a | rmin | lrc | Asym | Topt | Tmax | rmax | mu | sigma | rho_1 | rho_2 | rho_3 |
|---|---|---|---|---|---|---|---|---|---|---|---|---|---|---|---|---|---|---|
| ALL | Asym response to air temperature | ^{plot} | 342621.60 | 342686.35 | 0.41 | 4.29 | | | 0.71 | 1.00 | | | | | 0.39 | 0.11 | 0.15 | 0.15 |
| ALL | Asym response to air temperature | ^{ref} | 342846.90 | 342911.71 | 0.41 | 3.60 | | | 0.67 | 1.00 | | | | | 0.39 | 0.11 | 0.15 | 0.15 |
| ALL | Asym response to soil temperature | ^{plot} | 339503.20 | 339567.95 | 0.40 | 4.41 | | | 0.94 | 1.00 | | | | | 0.39 | 0.10 | 0.14 | 0.14 |
| ALL | Asym response to soil temperature | ^{ref} | 343053.00 | 343117.80 | 0.41 | 5.83 | | | 1.53 | 1.00 | | | | | 0.39 | 0.11 | 0.16 | 0.15 |
| ALL | Bi-linear response to air temperature | ^{plot} | 336613.10 | 336688.64 | 0.40 | 12.57 | 0.10 | 0.31 | | | | | | | 0.39 | 0.09 | 0.13 | 0.13 |
| ALL | Bi-linear response to air temperature | ^{ref} | 336879.20 | 336954.71 | 0.40 | 12.31 | 0.11 | 0.32 | | | | | | | 0.39 | 0.09 | 0.13 | 0.13 |
| ALL | Bi-linear response to soil temperature | ^{plot} | 331876.10 | 331951.69 | 0.39 | 9.95 | 0.10 | 0.25 | | | | | | | 0.38 | 0.07 | 0.12 | 0.12 |
| ALL | Bi-linear response to soil temperature | ^{ref} | 338460.70 | 338536.25 | 0.40 | 7.83 | 0.17 | 0.15 | | | | | | | 0.39 | 0.10 | 0.14 | 0.14 |
| ALL | Gaussian process | ^{plot} | 358273.10 | 358327.04 | 0.44 | | | | | | | | | 0.19 | 0.40 | 0.16 | 0.21 | 0.20 |
| ALL | Gaussian process | ^{ref} | 358322.90 | 358376.85 | 0.44 | | | | | | | | | 0.19 | 0.40 | 0.16 | 0.21 | 0.20 |
| ALL | Linear response to air temperature | ^{plot} | 340399.10 | 340463.87 | 0.40 | 3.75 | 0.05 | | | | | | | | 0.39 | 0.10 | 0.15 | 0.14 |
| ALL | Linear response to air temperature | ^{ref} | 340626.70 | 340691.42 | 0.40 | 3.22 | 0.05 | | | | | | | | 0.39 | 0.10 | 0.15 | 0.14 |
| ALL | Linear response to soil temperature | ^{plot} | 336532.40 | 336597.13 | 0.40 | 3.88 | 0.06 | | | | | | | | 0.39 | 0.09 | 0.13 | 0.13 |
| ALL | Linear response to soil temperature | ^{ref} | 341125.00 | 341189.79 | 0.40 | 5.53 | 0.11 | | | | | | | | 0.39 | 0.10 | 0.15 | 0.15 |
| ALL | Thermal time to air temperature | ^{plot} | 342177.50 | 342231.49 | 0.41 | 0.00 | 0.03 | | | | | | | | 0.39 | 0.11 | 0.15 | 0.15 |
| ALL | Thermal time to air temperature | ^{ref} | 341963.90 | 342017.86 | 0.40 | 0.00 | 0.03 | | | | | | | | 0.39 | 0.11 | 0.15 | 0.15 |
| ALL | Thermal time to soil temperature | ^{plot} | 340023.80 | 340077.79 | 0.40 | 0.00 | 0.04 | | | | | | | | 0.39 | 0.10 | 0.15 | 0.14 |
| ALL | Thermal time to soil temperature | ^{ref} | 350097.90 | 350151.91 | 0.42 | 0.00 | 0.03 | | | | | | | | 0.39 | 0.13 | 0.18 | 0.17 |
| ALL | Wang-Engel response to air temperature | ^{plot} | 339730.40 | 339795.18 | 0.40 | 0.00 | | | | | 19.19 | 25.00 | 0.81 | | 0.39 | 0.10 | 0.14 | 0.14 |
| ALL | Wang-Engel response to air temperature | ^{ref} | 340008.70 | 340073.46 | 0.40 | 0.00 | | | | | 18.94 | 25.00 | 0.81 | | 0.39 | 0.10 | 0.14 | 0.14 |
| ALL | Wang-Engel response to soil temperature | ^{plot} | 334542.50 | 334607.28 | 0.40 | 0.00 | | | | | 19.07 | 25.00 | 0.96 | | 0.39 | 0.08 | 0.13 | 0.12 |
| ALL | Wang-Engel response to soil temperature | ^{ref} | 338024.40 | 338089.21 | 0.40 | 0.00 | | | | | 20.40 | 25.00 | 2.88 | | 0.39 | 0.09 | 0.14 | 0.14 |
| CH CLARO | Asym response to air temperature | ^{plot} | 29564.00 | 29613.63 | 0.44 | 3.20 | | | 0.78 | 1.00 | | | | | 0.41 | 0.18 | 0.17 | 0.15 |
| CH CLARO | Asym response to air temperature | ^{ref} | 29581.70 | 29631.28 | 0.44 | 3.71 | | | 0.93 | 1.00 | | | | | 0.41 | 0.18 | 0.17 | 0.15 |
| CH CLARO | Asym response to soil temperature | ^{plot} | 29252.00 | 29301.61 | 0.43 | 4.53 | | | 1.21 | 1.00 | | | | | 0.40 | 0.17 | 0.16 | 0.14 |
| CH CLARO | Asym response to soil temperature | ^{ref} | 29385.80 | 29435.39 | 0.43 | 6.04 | | | 1.83 | 1.00 | | | | | 0.40 | 0.18 | 0.17 | 0.15 |
| CH CLARO | Bi-linear response to air temperature | ^{plot} | 29128.20 | 29186.10 | 0.43 | 10.65 | 0.09 | 0.31 | | | | | | | 0.40 | 0.17 | 0.16 | 0.14 |
| CH CLARO | Bi-linear response to air temperature | ^{ref} | 29169.00 | 29226.88 | 0.43 | 10.09 | 0.09 | 0.31 | | | | | | | 0.40 | 0.17 | 0.16 | 0.14 |
| CH CLARO | Bi-linear response to soil temperature | ^{plot} | 28751.30 | 28809.18 | 0.42 | 9.64 | 0.11 | 0.30 | | | | | | | 0.40 | 0.15 | 0.14 | 0.13 |
| CH CLARO | Bi-linear response to soil temperature | ^{ref} | 28978.10 | 29036.01 | 0.42 | 7.81 | 0.20 | 0.17 | | | | | | | 0.40 | 0.16 | 0.15 | 0.14 |
| CH CLARO | Gaussian process | ^{plot} | 30612.30 | 30653.67 | 0.48 | | | | | | | | | 0.23 | 0.41 | 0.23 | 0.22 | 0.20 |
| CH CLARO | Gaussian process | ^{ref} | 30616.60 | 30657.93 | 0.48 | | | | | | | | | 0.23 | 0.41 | 0.23 | 0.22 | 0.20 |
| CH CLARO | Linear response to air temperature | ^{plot} | 29363.80 | 29413.39 | 0.43 | 2.35 | 0.05 | | | | | | | | 0.40 | 0.18 | 0.16 | 0.15 |
| CH CLARO | Linear response to air temperature | ^{ref} | 29391.00 | 29440.58 | 0.43 | 1.85 | 0.05 | | | | | | | | 0.40 | 0.18 | 0.16 | 0.15 |
| CH CLARO | Linear response to soil temperature | ^{plot} | 29006.40 | 29055.99 | 0.42 | 3.17 | 0.07 | | | | | | | | 0.40 | 0.16 | 0.15 | 0.13 |
| CH CLARO | Linear response to soil temperature | ^{ref} | 29186.40 | 29236.02 | 0.43 | 5.85 | 0.14 | | | | | | | | 0.40 | 0.17 | 0.16 | 0.14 |
| CH CLARO | Thermal time to air temperature | ^{plot} | 29435.10 | 29476.42 | 0.43 | 0.00 | 0.04 | | | | | | | | 0.40 | 0.18 | 0.17 | 0.15 |
| CH CLARO | Thermal time to air temperature | ^{ref} | 29436.40 | 29477.77 | 0.43 | 0.00 | 0.04 | | | | | | | | 0.40 | 0.18 | 0.17 | 0.15 |
| CH CLARO | Thermal time to soil temperature | ^{plot} | 29228.80 | 29270.16 | 0.43 | 0.00 | 0.04 | | | | | | | | 0.40 | 0.17 | 0.16 | 0.14 |
| CH CLARO | Thermal time to soil temperature | ^{ref} | 29985.00 | 30026.33 | 0.45 | 0.00 | 0.04 | | | | | | | | 0.41 | 0.20 | 0.19 | 0.17 |
| CH CLARO | Wang-Engel response to air temperature | ^{plot} | 29340.40 | 29390.00 | 0.43 | 0.00 | | | | | 18.85 | 25.00 | 0.91 | | 0.40 | 0.17 | 0.16 | 0.14 |
| CH CLARO | Wang-Engel response to air temperature | ^{ref} | 29374.10 | 29423.69 | 0.43 | 0.00 | | | | | 18.57 | 25.00 | 0.90 | | 0.40 | 0.17 | 0.16 | 0.15 |
| CH CLARO | Wang-Engel response to soil temperature | ^{plot} | 28900.70 | 28950.26 | 0.42 | 0.00 | | | | | 18.73 | 25.00 | 1.06 | | 0.40 | 0.16 | 0.15 | 0.13 |
| CH CLARO | Wang-Engel response to soil temperature | ^{ref} | 28972.10 | 29021.71 | 0.42 | 0.00 | | | | | 20.44 | 25.00 | 3.48 | | 0.40 | 0.16 | 0.15 | 0.14 |
| CH NARA | Asym response to air temperature | ^{plot} | 27009.60 | 27060.54 | 0.36 | 5.82 | | | 0.97 | 1.00 | | | | | 0.35 | 0.09 | 0.15 | 0.14 |
| CH NARA | Asym response to air temperature | ^{ref} | 27049.70 | 27100.65 | 0.36 | 5.21 | | | 0.93 | 1.00 | | | | | 0.35 | 0.09 | 0.15 | 0.14 |
| CH NARA | Asym response to soil temperature | ^{plot} | 27210.30 | 27261.28 | 0.37 | 4.78 | | | 0.94 | 1.00 | | | | | 0.35 | 0.09 | 0.16 | 0.15 |
| CH NARA | Asym response to soil temperature | ^{ref} | 27924.40 | 27975.33 | 0.38 | 5.28 | | | 1.24 | 1.00 | | | | | 0.36 | 0.12 | 0.18 | 0.18 |

| | | | | | | | | | | | | | | | | | | |
|---|---|---|---|---|---|---|---|---|---|---|---|---|---|---|---|---|---|---|
| CH NARA | Bi-linear response to air temperature | ^{plot} | 26324.70 | 26384.19 | 0.36 | 10.80 | 0.09 | 0.23 | | | | | | | 0.35 | 0.07 | 0.13 | 0.12 |
| CH NARA | Bi-linear response to air temperature | ^{ref} | 26364.20 | 26423.66 | 0.36 | 10.56 | 0.09 | 0.24 | | | | | | | 0.35 | 0.06 | 0.13 | 0.12 |
| CH NARA | Bi-linear response to soil temperature | ^{plot} | 26228.20 | 26287.64 | 0.36 | 10.01 | 0.11 | 0.23 | | | | | | | 0.35 | 0.06 | 0.12 | 0.12 |
| CH NARA | Bi-linear response to soil temperature | ^{ref} | 27516.20 | 27575.65 | 0.37 | 8.73 | 0.20 | 0.18 | | | | | | | 0.35 | 0.10 | 0.17 | 0.16 |
| CH NARA | Gaussian process | ^{plot} | 29118.60 | 29161.09 | 0.40 | | | | | | | | | 0.17 | 0.36 | 0.16 | 0.22 | 0.21 |
| CH NARA | Gaussian process | ^{ref} | 29126.00 | 29168.44 | 0.40 | | | | | | | | | 0.17 | 0.36 | 0.16 | 0.22 | 0.21 |
| CH NARA | Linear response to air temperature | ^{plot} | 26713.60 | 26764.59 | 0.36 | 4.73 | 0.06 | | | | | | | | 0.35 | 0.08 | 0.14 | 0.13 |
| CH NARA | Linear response to air temperature | ^{ref} | 26761.80 | 26812.78 | 0.36 | 4.44 | 0.06 | | | | | | | | 0.35 | 0.08 | 0.14 | 0.13 |
| CH NARA | Linear response to soil temperature | ^{plot} | 26858.80 | 26909.79 | 0.36 | 4.45 | 0.06 | | | | | | | | 0.35 | 0.08 | 0.14 | 0.14 |
| CH NARA | Linear response to soil temperature | ^{ref} | 27773.80 | 27824.77 | 0.38 | 5.13 | 0.09 | | | | | | | | 0.36 | 0.11 | 0.18 | 0.17 |
| CH NARA | Thermal time to air temperature | ^{plot} | 27190.00 | 27232.50 | 0.37 | 0.00 | 0.03 | | | | | | | | 0.35 | 0.09 | 0.16 | 0.15 |
| CH NARA | Thermal time to air temperature | ^{ref} | 27167.40 | 27209.84 | 0.37 | 0.00 | 0.03 | | | | | | | | 0.35 | 0.09 | 0.15 | 0.15 |
| CH NARA | Thermal time to soil temperature | ^{plot} | 27332.40 | 27374.84 | 0.37 | 0.00 | 0.03 | | | | | | | | 0.35 | 0.10 | 0.16 | 0.16 |
| CH NARA | Thermal time to soil temperature | ^{ref} | 28421.80 | 28464.25 | 0.39 | 0.00 | 0.03 | | | | | | | | 0.36 | 0.13 | 0.20 | 0.19 |
| CH NARA | Wang-Engel response to air temperature | ^{plot} | 26581.10 | 26632.10 | 0.36 | 0.00 | | | | | 19.48 | 25.00 | 0.86 | | 0.35 | 0.07 | 0.13 | 0.13 |
| CH NARA | Wang-Engel response to air temperature | ^{ref} | 26616.50 | 26667.48 | 0.36 | 0.00 | | | | | 19.28 | 25.00 | 0.87 | | 0.35 | 0.07 | 0.13 | 0.13 |
| CH NARA | Wang-Engel response to soil temperature | ^{plot} | 26538.00 | 26588.91 | 0.36 | 0.00 | | | | | 19.42 | 25.00 | 1.03 | | 0.35 | 0.07 | 0.13 | 0.13 |
| CH NARA | Wang-Engel response to soil temperature | ^{ref} | 27495.30 | 27546.28 | 0.37 | 0.00 | | | | | 20.32 | 25.00 | 2.52 | | 0.35 | 0.10 | 0.17 | 0.16 |
| FASTNET | Asym response to air temperature | ^{plot} | 28372.00 | 28420.77 | 0.43 | 4.98 | | | 0.69 | 1.00 | | | | | 0.43 | -0.02 | 0.12 | 0.12 |
| FASTNET | Asym response to air temperature | ^{ref} | 28397.30 | 28446.11 | 0.43 | 4.40 | | | 0.67 | 1.00 | | | | | 0.43 | -0.02 | 0.12 | 0.12 |
| FASTNET | Asym response to soil temperature | ^{plot} | 28131.60 | 28180.45 | 0.43 | 4.11 | | | 0.76 | 1.00 | | | | | 0.42 | -0.03 | 0.11 | 0.11 |
| FASTNET | Asym response to soil temperature | ^{ref} | 28364.20 | 28413.01 | 0.43 | 5.18 | | | 1.28 | 1.00 | | | | | 0.43 | -0.02 | 0.12 | 0.12 |
| FASTNET | Bi-linear response to air temperature | ^{plot} | 27954.10 | 28011.06 | 0.43 | 10.08 | 0.07 | 0.20 | | | | | | | 0.42 | -0.04 | 0.10 | 0.10 |
| FASTNET | Bi-linear response to air temperature | ^{ref} | 27974.50 | 28031.48 | 0.43 | 9.59 | 0.07 | 0.20 | | | | | | | 0.42 | -0.04 | 0.10 | 0.10 |
| FASTNET | Bi-linear response to soil temperature | ^{plot} | 27676.10 | 27733.02 | 0.42 | 9.65 | 0.08 | 0.23 | | | | | | | 0.42 | -0.05 | 0.08 | 0.09 |
| FASTNET | Bi-linear response to soil temperature | ^{ref} | 28172.30 | 28229.24 | 0.43 | 6.36 | 0.10 | 0.09 | | | | | | | 0.42 | -0.03 | 0.11 | 0.12 |
| FASTNET | Gaussian process | ^{plot} | 29772.10 | 29812.81 | 0.46 | | | | | | | | | 0.18 | 0.44 | 0.04 | 0.19 | 0.19 |
| FASTNET | Gaussian process | ^{ref} | 29776.80 | 29817.50 | 0.46 | | | | | | | | | 0.18 | 0.44 | 0.04 | 0.19 | 0.19 |
| FASTNET | Linear response to air temperature | ^{plot} | 28232.10 | 28280.93 | 0.43 | 4.35 | 0.05 | | | | | | | | 0.42 | -0.03 | 0.11 | 0.12 |
| FASTNET | Linear response to air temperature | ^{ref} | 28256.00 | 28304.79 | 0.43 | 3.86 | 0.05 | | | | | | | | 0.42 | -0.03 | 0.11 | 0.11 |
| FASTNET | Linear response to soil temperature | ^{plot} | 27949.20 | 27997.96 | 0.43 | 2.90 | 0.05 | | | | | | | | 0.42 | -0.04 | 0.10 | 0.11 |
| FASTNET | Linear response to soil temperature | ^{ref} | 28246.00 | 28294.86 | 0.43 | 4.89 | 0.09 | | | | | | | | 0.42 | -0.03 | 0.11 | 0.12 |
| FASTNET | Thermal time to air temperature | ^{plot} | 28387.60 | 28428.25 | 0.43 | 0.00 | 0.03 | | | | | | | | 0.43 | -0.02 | 0.12 | 0.12 |
| FASTNET | Thermal time to air temperature | ^{ref} | 28369.70 | 28410.33 | 0.43 | 0.00 | 0.03 | | | | | | | | 0.43 | -0.02 | 0.12 | 0.12 |
| FASTNET | Thermal time to soil temperature | ^{plot} | 28172.90 | 28213.57 | 0.43 | 0.00 | 0.03 | | | | | | | | 0.42 | -0.03 | 0.11 | 0.12 |
| FASTNET | Thermal time to soil temperature | ^{ref} | 28977.60 | 29018.27 | 0.44 | 0.00 | 0.03 | | | | | | | | 0.43 | 0.01 | 0.15 | 0.15 |
| FASTNET | Wang-Engel response to air temperature | ^{plot} | 28165.70 | 28214.56 | 0.43 | 0.00 | | | | | 19.11 | 25.00 | 0.71 | | 0.42 | -0.03 | 0.11 | 0.11 |
| FASTNET | Wang-Engel response to air temperature | ^{ref} | 28183.10 | 28231.88 | 0.43 | 0.00 | | | | | 18.90 | 25.00 | 0.72 | | 0.42 | -0.03 | 0.11 | 0.11 |
| FASTNET | Wang-Engel response to soil temperature | ^{plot} | 27869.30 | 27918.12 | 0.43 | 0.00 | | | | | 18.75 | 25.00 | 0.80 | | 0.42 | -0.04 | 0.09 | 0.10 |
| FASTNET | Wang-Engel response to soil temperature | ^{ref} | 28111.00 | 28159.82 | 0.43 | 0.00 | | | | | 19.84 | 25.00 | 1.90 | | 0.42 | -0.03 | 0.11 | 0.11 |
| MARKSMAN | Asym response to air temperature | ^{plot} | 27898.50 | 27949.39 | 0.37 | 4.76 | | | 0.82 | 1.00 | | | | | 0.36 | 0.03 | 0.18 | 0.19 |
| MARKSMAN | Asym response to air temperature | ^{ref} | 27940.40 | 27991.27 | 0.37 | 3.59 | | | 0.70 | 1.00 | | | | | 0.36 | 0.03 | 0.18 | 0.19 |
| MARKSMAN | Asym response to soil temperature | ^{plot} | 27340.40 | 27391.31 | 0.37 | 4.38 | | | 0.98 | 1.00 | | | | | 0.36 | 0.01 | 0.17 | 0.18 |
| MARKSMAN | Asym response to soil temperature | ^{ref} | 27822.90 | 27873.75 | 0.37 | 5.78 | | | 1.57 | 1.00 | | | | | 0.36 | 0.03 | 0.19 | 0.19 |
| MARKSMAN | Bi-linear response to air temperature | ^{plot} | 26961.60 | 27020.99 | 0.36 | 10.98 | 0.09 | 0.26 | | | | | | | 0.35 | -0.00 | 0.15 | 0.16 |
| MARKSMAN | Bi-linear response to air temperature | ^{ref} | 27030.70 | 27090.11 | 0.36 | 10.57 | 0.09 | 0.26 | | | | | | | 0.35 | -0.00 | 0.15 | 0.16 |
| MARKSMAN | Bi-linear response to soil temperature | ^{plot} | 26188.00 | 26247.40 | 0.36 | 9.93 | 0.11 | 0.26 | | | | | | | 0.35 | -0.02 | 0.13 | 0.14 |
| MARKSMAN | Bi-linear response to soil temperature | ^{ref} | 27276.00 | 27335.32 | 0.37 | 7.67 | 0.17 | 0.15 | | | | | | | 0.36 | 0.01 | 0.17 | 0.17 |
| MARKSMAN | Gaussian process | ^{plot} | 29711.80 | 29754.26 | 0.41 | | | | | | | | | 0.19 | 0.37 | 0.09 | 0.26 | 0.25 |
| MARKSMAN | Gaussian process | ^{ref} | 29718.30 | 29760.74 | 0.41 | | | | | | | | | 0.19 | 0.37 | 0.09 | 0.26 | 0.25 |
| MARKSMAN | Linear response to air temperature | ^{plot} | 27573.90 | 27624.79 | 0.37 | 3.55 | 0.05 | | | | | | | | 0.36 | 0.02 | 0.17 | 0.18 |
| MARKSMAN | Linear response to air temperature | ^{ref} | 27611.60 | 27662.46 | 0.37 | 3.28 | 0.05 | | | | | | | | 0.36 | 0.02 | 0.17 | 0.18 |

| | | | | | | | | | | | | | | | | | |
|---|---|---|---|---|---|---|---|---|---|---|---|---|---|---|---|---|---|
| MARKSMAN | Linear response to soil temperature | ^{plot} | 26896.10 | 26946.96 | 0.36 | 4.01 | 0.06 | | | | | | | | 0.35 | -0.00 | 0.15 | 0.16 |
| MARKSMAN | Linear response to soil temperature | ^{ref} | 27550.10 | 27600.98 | 0.37 | 5.45 | 0.11 | | | | | | | | 0.36 | 0.02 | 0.18 | 0.18 |
| MARKSMAN | Thermal time to air temperature | ^{plot} | 27816.10 | 27858.55 | 0.37 | 0.00 | 0.03 | | | | | | | | 0.36 | 0.03 | 0.18 | 0.19 |
| MARKSMAN | Thermal time to air temperature | ^{ref} | 27797.10 | 27839.48 | 0.37 | 0.00 | 0.03 | | | | | | | | 0.36 | 0.03 | 0.18 | 0.18 |
| MARKSMAN | Thermal time to soil temperature | ^{plot} | 27405.50 | 27447.89 | 0.37 | 0.00 | 0.04 | | | | | | | | 0.36 | 0.02 | 0.17 | 0.18 |
| MARKSMAN | Thermal time to soil temperature | ^{ref} | 28753.60 | 28796.01 | 0.39 | 0.00 | 0.03 | | | | | | | | 0.36 | 0.06 | 0.22 | 0.22 |
| MARKSMAN | Wang-Engel response to air temperature | ^{plot} | 27417.90 | 27468.78 | 0.37 | 0.00 | | | | | 19.34 | 25.00 | 0.86 | | 0.36 | 0.01 | 0.16 | 0.17 |
| MARKSMAN | Wang-Engel response to air temperature | ^{ref} | 27459.90 | 27510.80 | 0.37 | 0.00 | | | | | 19.09 | 25.00 | 0.86 | | 0.36 | 0.01 | 0.17 | 0.17 |
| MARKSMAN | Wang-Engel response to soil temperature | ^{plot} | 26586.30 | 26637.21 | 0.36 | 0.00 | | | | | 19.06 | 25.00 | 1.00 | | 0.35 | -0.01 | 0.14 | 0.15 |
| MARKSMAN | Wang-Engel response to soil temperature | ^{ref} | 27166.90 | 27217.80 | 0.37 | 0.00 | | | | | 20.37 | 25.00 | 2.94 | | 0.35 | 0.01 | 0.16 | 0.17 |
| OSTKA STRZELECKA | Asym response to air temperature | ^{plot} | 36334.70 | 36385.16 | 0.42 | 2.21 | | | 0.14 | 1.00 | | | | | 0.42 | -0.01 | 0.08 | 0.10 |
| OSTKA STRZELECKA | Asym response to air temperature | ^{ref} | 36327.50 | 36377.96 | 0.42 | 1.37 | | | 0.09 | 1.00 | | | | | 0.42 | -0.01 | 0.08 | 0.10 |
| OSTKA STRZELECKA | Asym response to soil temperature | ^{plot} | 35746.60 | 35797.14 | 0.42 | 2.63 | | | 0.39 | 1.00 | | | | | 0.41 | -0.03 | 0.06 | 0.08 |
| OSTKA STRZELECKA | Asym response to soil temperature | ^{ref} | 35741.80 | 35792.35 | 0.42 | 5.10 | | | 1.10 | 1.00 | | | | | 0.41 | -0.03 | 0.06 | 0.08 |
| OSTKA STRZELECKA | Bi-linear response to air temperature | ^{plot} | 35715.20 | 35774.10 | 0.42 | 13.60 | 0.10 | 0.28 | | | | | | | 0.41 | -0.03 | 0.06 | 0.08 |
| OSTKA STRZELECKA | Bi-linear response to air temperature | ^{ref} | 35725.80 | 35784.68 | 0.42 | 13.22 | 0.09 | 0.29 | | | | | | | 0.41 | -0.03 | 0.06 | 0.08 |
| OSTKA STRZELECKA | Bi-linear response to soil temperature | ^{plot} | 34994.00 | 35052.89 | 0.41 | 10.65 | 0.08 | 0.24 | | | | | | | 0.41 | -0.06 | 0.04 | 0.06 |
| OSTKA STRZELECKA | Bi-linear response to soil temperature | ^{ref} | 35325.60 | 35384.53 | 0.41 | 7.42 | 0.12 | 0.12 | | | | | | | 0.41 | -0.05 | 0.05 | 0.07 |
| OSTKA STRZELECKA | Gaussian process | ^{plot} | 37807.40 | 37849.54 | 0.44 | | | | | | | | | 0.16 | 0.43 | 0.04 | 0.14 | 0.15 |
| OSTKA STRZELECKA | Gaussian process | ^{ref} | 37811.80 | 37853.85 | 0.44 | | | | | | | | | 0.16 | 0.43 | 0.04 | 0.14 | 0.15 |
| OSTKA STRZELECKA | Linear response to air temperature | ^{plot} | 36173.30 | 36223.77 | 0.42 | 1.72 | 0.03 | | | | | | | | 0.42 | -0.02 | 0.08 | 0.10 |
| OSTKA STRZELECKA | Linear response to air temperature | ^{ref} | 36166.40 | 36216.89 | 0.42 | 1.12 | 0.03 | | | | | | | | 0.42 | -0.02 | 0.08 | 0.10 |
| OSTKA STRZELECKA | Linear response to soil temperature | ^{plot} | 35502.40 | 35552.94 | 0.41 | 2.23 | 0.04 | | | | | | | | 0.41 | -0.04 | 0.05 | 0.08 |
| OSTKA STRZELECKA | Linear response to soil temperature | ^{ref} | 35538.70 | 35589.24 | 0.41 | 4.92 | 0.08 | | | | | | | | 0.41 | -0.04 | 0.06 | 0.08 |
| OSTKA STRZELECKA | Thermal time to air temperature | ^{plot} | 36209.90 | 36251.97 | 0.42 | 0.00 | 0.03 | | | | | | | | 0.42 | -0.02 | 0.08 | 0.10 |
| OSTKA STRZELECKA | Thermal time to air temperature | ^{ref} | 36180.40 | 36222.45 | 0.42 | 0.00 | 0.03 | | | | | | | | 0.42 | -0.02 | 0.08 | 0.10 |
| OSTKA STRZELECKA | Thermal time to soil temperature | ^{plot} | 35697.50 | 35739.56 | 0.42 | 0.00 | 0.03 | | | | | | | | 0.41 | -0.03 | 0.06 | 0.08 |
| OSTKA STRZELECKA | Thermal time to soil temperature | ^{ref} | 36650.60 | 36692.73 | 0.42 | 0.00 | 0.03 | | | | | | | | 0.42 | -0.00 | 0.10 | 0.11 |
| OSTKA STRZELECKA | Wang-Engel response to air temperature | ^{plot} | 36199.20 | 36249.66 | 0.42 | 0.00 | | | | | 18.95 | 25.00 | 0.60 | | 0.42 | -0.02 | 0.08 | 0.10 |
| OSTKA STRZELECKA | Wang-Engel response to air temperature | ^{ref} | 36216.50 | 36266.97 | 0.42 | 0.00 | | | | | 18.68 | 25.00 | 0.60 | | 0.42 | -0.02 | 0.08 | 0.10 |
| OSTKA STRZELECKA | Wang-Engel response to soil temperature | ^{plot} | 35420.90 | 35471.42 | 0.41 | 0.00 | | | | | 18.77 | 25.00 | 0.73 | | 0.41 | -0.04 | 0.05 | 0.07 |
| OSTKA STRZELECKA | Wang-Engel response to soil temperature | ^{ref} | 35197.80 | 35248.31 | 0.41 | 0.00 | | | | | 20.13 | 25.00 | 2.04 | | 0.41 | -0.05 | 0.04 | 0.07 |
| ROMANUS | Asym response to air temperature | ^{plot} | 13478.10 | 13527.41 | 0.33 | 2.55 | | | 0.33 | 1.00 | | | | | 0.31 | 0.20 | 0.15 | 0.15 |
| ROMANUS | Asym response to air temperature | ^{ref} | 13484.40 | 13533.74 | 0.33 | 2.53 | | | 0.39 | 1.00 | | | | | 0.31 | 0.20 | 0.15 | 0.15 |
| ROMANUS | Asym response to soil temperature | ^{plot} | 13058.20 | 13107.58 | 0.32 | 4.39 | | | 0.82 | 1.00 | | | | | 0.31 | 0.19 | 0.13 | 0.13 |
| ROMANUS | Asym response to soil temperature | ^{ref} | 13318.50 | 13367.81 | 0.33 | 6.33 | | | 1.58 | 1.00 | | | | | 0.31 | 0.20 | 0.14 | 0.14 |
| ROMANUS | Bi-linear response to air temperature | ^{plot} | 13136.30 | 13193.92 | 0.32 | 10.80 | 0.07 | 0.23 | | | | | | | 0.31 | 0.19 | 0.13 | 0.13 |
| ROMANUS | Bi-linear response to air temperature | ^{ref} | 13146.10 | 13203.67 | 0.33 | 14.24 | 0.12 | 0.36 | | | | | | | 0.31 | 0.19 | 0.13 | 0.13 |
| ROMANUS | Bi-linear response to soil temperature | ^{plot} | 12667.60 | 12725.22 | 0.32 | 9.35 | 0.08 | 0.21 | | | | | | | 0.30 | 0.17 | 0.12 | 0.11 |
| ROMANUS | Bi-linear response to soil temperature | ^{ref} | 12929.40 | 12986.95 | 0.32 | 7.67 | 0.15 | 0.11 | | | | | | | 0.31 | 0.18 | 0.13 | 0.13 |
| ROMANUS | Gaussian process | ^{plot} | 14595.20 | 14636.34 | 0.36 | | | | | | | | | 0.17 | 0.32 | 0.25 | 0.19 | 0.19 |
| ROMANUS | Gaussian process | ^{ref} | 14599.90 | 14640.99 | 0.36 | | | | | | | | | 0.17 | 0.32 | 0.25 | 0.19 | 0.19 |
| ROMANUS | Linear response to air temperature | ^{plot} | 13347.40 | 13396.76 | 0.33 | 2.43 | 0.04 | | | | | | | | 0.31 | 0.20 | 0.14 | 0.14 |
| ROMANUS | Linear response to air temperature | ^{ref} | 13362.00 | 13411.37 | 0.33 | 2.11 | 0.04 | | | | | | | | 0.31 | 0.20 | 0.14 | 0.14 |
| ROMANUS | Linear response to soil temperature | ^{plot} | 12883.30 | 12932.66 | 0.32 | 3.99 | 0.06 | | | | | | | | 0.31 | 0.18 | 0.13 | 0.12 |
| ROMANUS | Linear response to soil temperature | ^{ref} | 13172.50 | 13221.84 | 0.33 | 6.01 | 0.11 | | | | | | | | 0.31 | 0.19 | 0.14 | 0.14 |
| ROMANUS | Thermal time to air temperature | ^{plot} | 13431.80 | 13472.89 | 0.33 | 0.00 | 0.03 | | | | | | | | 0.31 | 0.20 | 0.15 | 0.15 |
| ROMANUS | Thermal time to air temperature | ^{ref} | 13424.90 | 13466.05 | 0.33 | 0.00 | 0.03 | | | | | | | | 0.31 | 0.20 | 0.15 | 0.15 |
| ROMANUS | Thermal time to soil temperature | ^{plot} | 13196.20 | 13237.29 | 0.33 | 0.00 | 0.03 | | | | | | | | 0.31 | 0.19 | 0.14 | 0.14 |
| ROMANUS | Thermal time to soil temperature | ^{ref} | 13992.50 | 14033.66 | 0.34 | 0.00 | 0.03 | | | | | | | | 0.31 | 0.23 | 0.17 | 0.17 |
| ROMANUS | Wang-Engel response to air temperature | ^{plot} | 13371.80 | 13421.17 | 0.33 | 0.00 | | | | | 18.87 | 25.00 | 0.67 | | 0.31 | 0.20 | 0.14 | 0.14 |
| ROMANUS | Wang-Engel response to air temperature | ^{ref} | 13399.60 | 13448.95 | 0.33 | 0.00 | | | | | 18.56 | 25.00 | 0.66 | | 0.31 | 0.20 | 0.14 | 0.14 |

| | | | | | | | | | | | | | | | | | |
|---|---|---|---|---|---|---|---|---|---|---|---|---|---|---|---|---|---|
| ROMANUS | Wang-Engel response to soil temperature | ^{plot} | 12773.10 | 12822.48 | 0.32 | 0.00 | | | | | 18.86 | 25.00 | 0.82 | | 0.31 | 0.18 | 0.12 | 0.12 |
| ROMANUS | Wang-Engel response to soil temperature | ^{ref} | 12904.90 | 12954.30 | 0.32 | 0.00 | | | | | 20.67 | 25.00 | 3.16 | | 0.31 | 0.18 | 0.13 | 0.13 |
| RUNAL | Asym response to air temperature | ^{plot} | 16955.10 | 17004.05 | 0.37 | 3.18 | | | 0.65 | 1.00 | | | | | 0.34 | 0.25 | 0.14 | 0.14 |
| RUNAL | Asym response to air temperature | ^{ref} | 16961.80 | 17010.75 | 0.37 | 2.31 | | | 0.59 | 1.00 | | | | | 0.34 | 0.25 | 0.14 | 0.14 |
| RUNAL | Asym response to soil temperature | ^{plot} | 16669.30 | 16718.34 | 0.36 | 3.02 | | | 0.85 | 1.00 | | | | | 0.33 | 0.23 | 0.13 | 0.13 |
| RUNAL | Asym response to soil temperature | ^{ref} | 16919.00 | 16968.02 | 0.36 | 5.66 | | | 1.64 | 1.00 | | | | | 0.34 | 0.24 | 0.14 | 0.14 |
| RUNAL | Bi-linear response to air temperature | ^{plot} | 16477.60 | 16534.71 | 0.35 | 13.37 | 0.13 | 0.38 | | | | | | | 0.33 | 0.23 | 0.12 | 0.12 |
| RUNAL | Bi-linear response to air temperature | ^{ref} | 16499.30 | 16556.50 | 0.35 | 13.19 | 0.14 | 0.40 | | | | | | | 0.33 | 0.23 | 0.12 | 0.12 |
| RUNAL | Bi-linear response to soil temperature | ^{plot} | 15943.40 | 16000.58 | 0.34 | 10.04 | 0.12 | 0.30 | | | | | | | 0.33 | 0.20 | 0.10 | 0.10 |
| RUNAL | Bi-linear response to soil temperature | ^{ref} | 16352.50 | 16409.68 | 0.35 | 8.06 | 0.22 | 0.18 | | | | | | | 0.33 | 0.22 | 0.12 | 0.12 |
| RUNAL | Gaussian process | ^{plot} | 17967.30 | 18008.16 | 0.40 | | | | | | | | | 0.20 | 0.34 | 0.29 | 0.18 | 0.19 |
| RUNAL | Gaussian process | ^{ref} | 17970.80 | 18011.67 | 0.40 | | | | | | | | | 0.21 | 0.34 | 0.29 | 0.18 | 0.19 |
| RUNAL | Linear response to air temperature | ^{plot} | 16774.30 | 16823.31 | 0.36 | 2.51 | 0.05 | | | | | | | | 0.33 | 0.24 | 0.13 | 0.13 |
| RUNAL | Linear response to air temperature | ^{ref} | 16788.20 | 16837.24 | 0.36 | 2.12 | 0.05 | | | | | | | | 0.33 | 0.24 | 0.13 | 0.14 |
| RUNAL | Linear response to soil temperature | ^{plot} | 16372.10 | 16421.05 | 0.35 | 2.90 | 0.06 | | | | | | | | 0.33 | 0.22 | 0.12 | 0.12 |
| RUNAL | Linear response to soil temperature | ^{ref} | 16706.40 | 16755.40 | 0.36 | 5.36 | 0.12 | | | | | | | | 0.33 | 0.24 | 0.13 | 0.13 |
| RUNAL | Thermal time to air temperature | ^{plot} | 16861.90 | 16902.75 | 0.36 | 0.00 | 0.03 | | | | | | | | 0.34 | 0.24 | 0.14 | 0.14 |
| RUNAL | Thermal time to air temperature | ^{ref} | 16851.60 | 16892.38 | 0.36 | 0.00 | 0.04 | | | | | | | | 0.33 | 0.24 | 0.14 | 0.14 |
| RUNAL | Thermal time to soil temperature | ^{plot} | 16600.10 | 16640.94 | 0.36 | 0.00 | 0.04 | | | | | | | | 0.33 | 0.23 | 0.13 | 0.13 |
| RUNAL | Thermal time to soil temperature | ^{ref} | 17382.90 | 17423.70 | 0.38 | 0.00 | 0.04 | | | | | | | | 0.34 | 0.27 | 0.16 | 0.16 |
| RUNAL | Wang-Engel response to air temperature | ^{plot} | 16773.90 | 16822.91 | 0.36 | 0.00 | | | | | 18.91 | 25.00 | 0.85 | | 0.33 | 0.24 | 0.13 | 0.13 |
| RUNAL | Wang-Engel response to air temperature | ^{ref} | 16807.00 | 16856.02 | 0.36 | 0.00 | | | | | 18.61 | 25.00 | 0.84 | | 0.33 | 0.24 | 0.13 | 0.13 |
| RUNAL | Wang-Engel response to soil temperature | ^{plot} | 16222.10 | 16271.12 | 0.35 | 0.00 | | | | | 18.93 | 25.00 | 1.09 | | 0.33 | 0.22 | 0.11 | 0.11 |
| RUNAL | Wang-Engel response to soil temperature | ^{ref} | 16360.10 | 16409.07 | 0.35 | 0.00 | | | | | 20.48 | 25.00 | 3.62 | | 0.33 | 0.22 | 0.12 | 0.12 |
| RYWALKA | Asym response to air temperature | ^{plot} | 29473.10 | 29523.82 | 0.38 | 5.51 | | | 0.40 | 1.00 | | | | | 0.37 | 0.02 | 0.10 | 0.10 |
| RYWALKA | Asym response to air temperature | ^{ref} | 29499.90 | 29550.59 | 0.38 | 5.22 | | | 0.43 | 1.00 | | | | | 0.37 | 0.02 | 0.10 | 0.10 |
| RYWALKA | Asym response to soil temperature | ^{plot} | 29289.10 | 29339.81 | 0.37 | 5.71 | | | 0.66 | 1.00 | | | | | 0.37 | 0.01 | 0.10 | 0.10 |
| RYWALKA | Asym response to soil temperature | ^{ref} | 29909.40 | 29960.14 | 0.38 | 6.11 | | | 1.06 | 1.00 | | | | | 0.37 | 0.03 | 0.12 | 0.12 |
| RYWALKA | Bi-linear response to air temperature | ^{plot} | 29213.50 | 29272.73 | 0.37 | 12.49 | 0.08 | 0.19 | | | | | | | 0.37 | 0.01 | 0.09 | 0.10 |
| RYWALKA | Bi-linear response to air temperature | ^{ref} | 29216.30 | 29275.47 | 0.37 | 12.16 | 0.08 | 0.20 | | | | | | | 0.37 | 0.01 | 0.09 | 0.10 |
| RYWALKA | Bi-linear response to soil temperature | ^{plot} | 28850.90 | 28910.07 | 0.37 | 9.69 | 0.07 | 0.13 | | | | | | | 0.37 | -0.00 | 0.08 | 0.09 |
| RYWALKA | Bi-linear response to soil temperature | ^{ref} | 29623.20 | 29682.42 | 0.38 | 8.04 | 0.12 | 0.09 | | | | | | | 0.37 | 0.02 | 0.11 | 0.11 |
| RYWALKA | Gaussian process | ^{plot} | 31064.10 | 31106.34 | 0.39 | | | | | | | | | 0.11 | 0.38 | 0.07 | 0.16 | 0.16 |
| RYWALKA | Gaussian process | ^{ref} | 31068.30 | 31110.56 | 0.39 | | | | | | | | | 0.11 | 0.38 | 0.07 | 0.16 | 0.16 |
| RYWALKA | Linear response to air temperature | ^{plot} | 29374.20 | 29424.95 | 0.37 | 5.18 | 0.04 | | | | | | | | 0.37 | 0.01 | 0.10 | 0.10 |
| RYWALKA | Linear response to air temperature | ^{ref} | 29401.40 | 29452.09 | 0.37 | 4.62 | 0.04 | | | | | | | | 0.37 | 0.01 | 0.10 | 0.10 |
| RYWALKA | Linear response to soil temperature | ^{plot} | 29149.80 | 29200.49 | 0.37 | 5.25 | 0.05 | | | | | | | | 0.37 | 0.01 | 0.09 | 0.10 |
| RYWALKA | Linear response to soil temperature | ^{ref} | 29836.10 | 29886.80 | 0.38 | 6.00 | 0.08 | | | | | | | | 0.37 | 0.03 | 0.12 | 0.12 |
| RYWALKA | Thermal time to air temperature | ^{plot} | 29727.50 | 29769.79 | 0.38 | 0.00 | 0.02 | | | | | | | | 0.37 | 0.03 | 0.11 | 0.11 |
| RYWALKA | Thermal time to air temperature | ^{ref} | 29687.60 | 29729.84 | 0.38 | 0.00 | 0.02 | | | | | | | | 0.37 | 0.02 | 0.11 | 0.11 |
| RYWALKA | Thermal time to soil temperature | ^{plot} | 29668.40 | 29710.72 | 0.38 | 0.00 | 0.02 | | | | | | | | 0.37 | 0.02 | 0.11 | 0.11 |
| RYWALKA | Thermal time to soil temperature | ^{ref} | 30469.70 | 30511.96 | 0.39 | 0.00 | 0.02 | | | | | | | | 0.38 | 0.05 | 0.14 | 0.14 |
| RYWALKA | Wang-Engel response to air temperature | ^{plot} | 29288.90 | 29339.61 | 0.37 | 0.00 | | | | | 19.62 | 25.00 | 0.59 | | 0.37 | 0.01 | 0.10 | 0.10 |
| RYWALKA | Wang-Engel response to air temperature | ^{ref} | 29309.10 | 29359.85 | 0.37 | 0.00 | | | | | 19.38 | 25.00 | 0.59 | | 0.37 | 0.01 | 0.10 | 0.10 |
| RYWALKA | Wang-Engel response to soil temperature | ^{plot} | 28961.90 | 29012.66 | 0.37 | 0.00 | | | | | 19.62 | 25.00 | 0.74 | | 0.37 | 0.00 | 0.09 | 0.09 |
| RYWALKA | Wang-Engel response to soil temperature | ^{ref} | 29614.80 | 29665.57 | 0.38 | 0.00 | | | | | 20.66 | 25.00 | 2.21 | | 0.37 | 0.02 | 0.11 | 0.11 |
| SEMAFOR | Asym response to air temperature | ^{plot} | 21929.10 | 21977.89 | 0.40 | 2.44 | | | 0.36 | 1.00 | | | | | 0.38 | 0.26 | 0.09 | 0.09 |
| SEMAFOR | Asym response to air temperature | ^{ref} | 21929.00 | 21977.72 | 0.40 | 2.28 | | | 0.40 | 1.00 | | | | | 0.38 | 0.26 | 0.09 | 0.09 |
| SEMAFOR | Asym response to soil temperature | ^{plot} | 22008.00 | 22056.79 | 0.40 | 3.03 | | | 0.55 | 1.00 | | | | | 0.38 | 0.27 | 0.10 | 0.09 |
| SEMAFOR | Asym response to soil temperature | ^{ref} | 22297.30 | 22346.03 | 0.41 | 4.91 | | | 1.02 | 1.00 | | | | | 0.38 | 0.28 | 0.11 | 0.11 |
| SEMAFOR | Bi-linear response to air temperature | ^{plot} | 21691.40 | 21748.28 | 0.39 | 13.68 | 0.11 | 0.34 | | | | | | | 0.37 | 0.25 | 0.08 | 0.08 |
| SEMAFOR | Bi-linear response to air temperature | ^{ref} | 21691.30 | 21748.20 | 0.39 | 14.07 | 0.12 | 0.38 | | | | | | | 0.37 | 0.25 | 0.08 | 0.08 |

| | | | | | | | | | | | | | | | | | | |
|---|---|---|---|---|---|---|---|---|---|---|---|---|---|---|---|---|---|---|
| SEMAFOR | Bi-linear response to soil temperature | ^{plot} | 21637.70 | 21694.58 | 0.39 | 10.31 | 0.10 | 0.24 | | | | | | | 0.37 | 0.25 | 0.08 | 0.08 |
| SEMAFOR | Bi-linear response to soil temperature | ^{ref} | 22046.20 | 22103.07 | 0.40 | 8.55 | 0.17 | 0.17 | | | | | | | 0.38 | 0.27 | 0.10 | 0.10 |
| SEMAFOR | Gaussian process | ^{plot} | 22834.70 | 22875.32 | 0.43 | | | | | | | | | 0.17 | 0.38 | 0.30 | 0.13 | 0.13 |
| SEMAFOR | Gaussian process | ^{ref} | 22836.70 | 22877.32 | 0.43 | | | | | | | | | 0.17 | 0.38 | 0.30 | 0.13 | 0.13 |
| SEMAFOR | Linear response to air temperature | ^{plot} | 21846.40 | 21895.15 | 0.40 | 1.91 | 0.04 | | | | | | | | 0.37 | 0.26 | 0.09 | 0.08 |
| SEMAFOR | Linear response to air temperature | ^{ref} | 21847.60 | 21896.39 | 0.40 | 1.40 | 0.04 | | | | | | | | 0.37 | 0.26 | 0.09 | 0.08 |
| SEMAFOR | Linear response to soil temperature | ^{plot} | 21893.30 | 21942.04 | 0.40 | 2.92 | 0.05 | | | | | | | | 0.38 | 0.26 | 0.09 | 0.09 |
| SEMAFOR | Linear response to soil temperature | ^{ref} | 22235.00 | 22283.75 | 0.41 | 4.74 | 0.08 | | | | | | | | 0.38 | 0.28 | 0.11 | 0.10 |
| SEMAFOR | Thermal time to air temperature | ^{plot} | 21888.50 | 21929.11 | 0.40 | 0.00 | 0.03 | | | | | | | | 0.38 | 0.26 | 0.09 | 0.09 |
| SEMAFOR | Thermal time to air temperature | ^{ref} | 21874.70 | 21915.29 | 0.40 | 0.00 | 0.03 | | | | | | | | 0.38 | 0.26 | 0.09 | 0.09 |
| SEMAFOR | Thermal time to soil temperature | ^{plot} | 21976.10 | 22016.71 | 0.40 | 0.00 | 0.03 | | | | | | | | 0.38 | 0.27 | 0.10 | 0.09 |
| SEMAFOR | Thermal time to soil temperature | ^{ref} | 22463.60 | 22504.25 | 0.41 | 0.00 | 0.03 | | | | | | | | 0.38 | 0.29 | 0.11 | 0.11 |
| SEMAFOR | Wang-Engel response to air temperature | ^{plot} | 21887.20 | 21936.01 | 0.40 | 0.00 | | | | | 18.49 | 25.00 | 0.65 | | 0.38 | 0.26 | 0.09 | 0.08 |
| SEMAFOR | Wang-Engel response to air temperature | ^{ref} | 21887.20 | 21935.95 | 0.40 | 0.00 | | | | | 18.25 | 25.00 | 0.65 | | 0.38 | 0.26 | 0.09 | 0.09 |
| SEMAFOR | Wang-Engel response to soil temperature | ^{plot} | 21806.50 | 21855.23 | 0.40 | 0.00 | | | | | 19.06 | 25.00 | 0.87 | | 0.37 | 0.26 | 0.09 | 0.08 |
| SEMAFOR | Wang-Engel response to soil temperature | ^{ref} | 22084.30 | 22133.06 | 0.40 | 0.00 | | | | | 20.20 | 25.00 | 2.16 | | 0.38 | 0.27 | 0.10 | 0.10 |
| TAMARO | Asym response to air temperature | ^{plot} | 30921.10 | 30971.30 | 0.43 | 4.83 | | | 1.04 | 1.00 | | | | | 0.39 | 0.15 | 0.20 | 0.18 |
| TAMARO | Asym response to air temperature | ^{ref} | 30928.80 | 30979.04 | 0.43 | 4.38 | | | 1.03 | 1.00 | | | | | 0.39 | 0.15 | 0.20 | 0.18 |
| TAMARO | Asym response to soil temperature | ^{plot} | 30568.90 | 30619.14 | 0.42 | 4.99 | | | 1.30 | 1.00 | | | | | 0.39 | 0.14 | 0.19 | 0.17 |
| TAMARO | Asym response to soil temperature | ^{ref} | 30696.60 | 30746.85 | 0.42 | 6.26 | | | 1.95 | 1.00 | | | | | 0.39 | 0.14 | 0.20 | 0.17 |
| TAMARO | Bi-linear response to air temperature | ^{plot} | 30101.20 | 30159.79 | 0.41 | 13.00 | 0.16 | 0.37 | | | | | | | 0.39 | 0.12 | 0.17 | 0.15 |
| TAMARO | Bi-linear response to air temperature | ^{ref} | 30139.50 | 30198.07 | 0.41 | 12.27 | 0.15 | 0.37 | | | | | | | 0.39 | 0.12 | 0.17 | 0.15 |
| TAMARO | Bi-linear response to soil temperature | ^{plot} | 29513.40 | 29571.99 | 0.40 | 9.99 | 0.14 | 0.28 | | | | | | | 0.38 | 0.10 | 0.15 | 0.13 |
| TAMARO | Bi-linear response to soil temperature | ^{ref} | 30093.30 | 30151.86 | 0.41 | 8.04 | 0.24 | 0.16 | | | | | | | 0.39 | 0.12 | 0.18 | 0.15 |
| TAMARO | Gaussian process | ^{plot} | 32152.60 | 32194.47 | 0.47 | | | | | | | | | 0.22 | 0.40 | 0.20 | 0.25 | 0.23 |
| TAMARO | Gaussian process | ^{ref} | 32155.30 | 32197.16 | 0.47 | | | | | | | | | 0.22 | 0.40 | 0.20 | 0.25 | 0.23 |
| TAMARO | Linear response to air temperature | ^{plot} | 30590.50 | 30640.73 | 0.42 | 4.16 | 0.06 | | | | | | | | 0.39 | 0.14 | 0.19 | 0.17 |
| TAMARO | Linear response to air temperature | ^{ref} | 30604.40 | 30654.66 | 0.42 | 4.11 | 0.07 | | | | | | | | 0.39 | 0.14 | 0.19 | 0.17 |
| TAMARO | Linear response to soil temperature | ^{plot} | 30073.20 | 30123.45 | 0.41 | 4.76 | 0.09 | | | | | | | | 0.39 | 0.12 | 0.17 | 0.15 |
| TAMARO | Linear response to soil temperature | ^{ref} | 30358.80 | 30409.01 | 0.41 | 6.10 | 0.16 | | | | | | | | 0.39 | 0.13 | 0.19 | 0.16 |
| TAMARO | Thermal time to air temperature | ^{plot} | 30862.60 | 30904.42 | 0.43 | 0.00 | 0.04 | | | | | | | | 0.39 | 0.15 | 0.20 | 0.18 |
| TAMARO | Thermal time to air temperature | ^{ref} | 30841.80 | 30883.60 | 0.42 | 0.00 | 0.04 | | | | | | | | 0.39 | 0.15 | 0.20 | 0.18 |
| TAMARO | Thermal time to soil temperature | ^{plot} | 30628.70 | 30670.56 | 0.42 | 0.00 | 0.04 | | | | | | | | 0.39 | 0.14 | 0.20 | 0.17 |
| TAMARO | Thermal time to soil temperature | ^{ref} | 31511.20 | 31553.02 | 0.45 | 0.00 | 0.04 | | | | | | | | 0.40 | 0.18 | 0.23 | 0.20 |
| TAMARO | Wang-Engel response to air temperature | ^{plot} | 30463.10 | 30513.32 | 0.41 | 0.00 | | | | | 19.71 | 25.00 | 1.12 | | 0.39 | 0.13 | 0.18 | 0.16 |
| TAMARO | Wang-Engel response to air temperature | ^{ref} | 30472.60 | 30522.82 | 0.42 | 0.00 | | | | | 19.50 | 25.00 | 1.14 | | 0.39 | 0.13 | 0.19 | 0.16 |
| TAMARO | Wang-Engel response to soil temperature | ^{plot} | 29776.90 | 29827.08 | 0.40 | 0.00 | | | | | 19.54 | 25.00 | 1.36 | | 0.39 | 0.11 | 0.16 | 0.14 |
| TAMARO | Wang-Engel response to soil temperature | ^{ref} | 30013.90 | 30064.12 | 0.41 | 0.00 | | | | | 20.91 | 25.00 | 5.36 | | 0.39 | 0.12 | 0.17 | 0.15 |
| TORONTO | Asym response to air temperature | ^{plot} | 24084.30 | 24132.42 | 0.43 | 5.36 | | | 1.01 | 1.00 | | | | | 0.41 | 0.17 | 0.12 | 0.12 |
| TORONTO | Asym response to air temperature | ^{ref} | 24092.20 | 24140.29 | 0.43 | 5.02 | | | 1.03 | 1.00 | | | | | 0.41 | 0.17 | 0.12 | 0.13 |
| TORONTO | Asym response to soil temperature | ^{plot} | 23858.60 | 23906.74 | 0.43 | 4.53 | | | 1.04 | 1.00 | | | | | 0.41 | 0.16 | 0.11 | 0.11 |
| TORONTO | Asym response to soil temperature | ^{ref} | 23948.90 | 23997.04 | 0.43 | 6.38 | | | 1.77 | 1.00 | | | | | 0.41 | 0.16 | 0.11 | 0.12 |
| TORONTO | Bi-linear response to air temperature | ^{plot} | 23639.10 | 23695.23 | 0.42 | 10.97 | 0.10 | 0.26 | | | | | | | 0.41 | 0.15 | 0.10 | 0.10 |
| TORONTO | Bi-linear response to air temperature | ^{ref} | 23655.50 | 23711.59 | 0.42 | 10.59 | 0.10 | 0.27 | | | | | | | 0.41 | 0.15 | 0.10 | 0.10 |
| TORONTO | Bi-linear response to soil temperature | ^{plot} | 23335.00 | 23391.15 | 0.42 | 9.98 | 0.11 | 0.26 | | | | | | | 0.41 | 0.13 | 0.08 | 0.09 |
| TORONTO | Bi-linear response to soil temperature | ^{ref} | 23654.20 | 23710.35 | 0.42 | 7.65 | 0.17 | 0.13 | | | | | | | 0.41 | 0.15 | 0.10 | 0.11 |
| TORONTO | Gaussian process | ^{plot} | 25115.60 | 25155.68 | 0.47 | | | | | | | | | 0.20 | 0.42 | 0.22 | 0.17 | 0.18 |
| TORONTO | Gaussian process | ^{ref} | 25119.00 | 25159.07 | 0.47 | | | | | | | | | 0.20 | 0.42 | 0.22 | 0.17 | 0.18 |
| TORONTO | Linear response to air temperature | ^{plot} | 23916.10 | 23964.23 | 0.43 | 4.02 | 0.06 | | | | | | | | 0.41 | 0.16 | 0.11 | 0.12 |
| TORONTO | Linear response to air temperature | ^{ref} | 23920.20 | 23968.32 | 0.43 | 3.99 | 0.06 | | | | | | | | 0.41 | 0.16 | 0.11 | 0.12 |
| TORONTO | Linear response to soil temperature | ^{plot} | 23629.20 | 23677.26 | 0.42 | 4.28 | 0.07 | | | | | | | | 0.41 | 0.15 | 0.10 | 0.10 |
| TORONTO | Linear response to soil temperature | ^{ref} | 23802.80 | 23850.87 | 0.43 | 6.01 | 0.13 | | | | | | | | 0.41 | 0.16 | 0.11 | 0.11 |

| | | | | | | | | | | | | | | | | | | |
|---|---|---|---|---|---|---|---|---|---|---|---|---|---|---|---|---|---|---|
| TORONTO | Thermal time to air temperature | ^{plot} | 24084.70 | 24124.80 | 0.43 | 0.00 | 0.03 | | | | | | | | 0.41 | 0.17 | 0.12 | 0.13 |
| TORONTO | Thermal time to air temperature | ^{ref} | 24069.20 | 24109.26 | 0.43 | 0.00 | 0.04 | | | | | | | | 0.41 | 0.17 | 0.12 | 0.13 |
| TORONTO | Thermal time to soil temperature | ^{plot} | 23897.60 | 23937.70 | 0.43 | 0.00 | 0.04 | | | | | | | | 0.41 | 0.16 | 0.11 | 0.12 |
| TORONTO | Thermal time to soil temperature | ^{ref} | 24569.10 | 24609.16 | 0.45 | 0.00 | 0.03 | | | | | | | | 0.42 | 0.20 | 0.14 | 0.15 |
| TORONTO | Wang-Engel response to air temperature | ^{plot} | 23822.90 | 23870.97 | 0.43 | 0.00 | | | | | 19.40 | 25.00 | 0.93 | | 0.41 | 0.16 | 0.10 | 0.11 |
| TORONTO | Wang-Engel response to air temperature | ^{ref} | 23831.90 | 23880.00 | 0.43 | 0.00 | | | | | 19.17 | 25.00 | 0.94 | | 0.41 | 0.16 | 0.11 | 0.11 |
| TORONTO | Wang-Engel response to soil temperature | ^{plot} | 23504.60 | 23552.66 | 0.42 | 0.00 | | | | | 19.10 | 25.00 | 1.02 | | 0.41 | 0.14 | 0.09 | 0.10 |
| TORONTO | Wang-Engel response to soil temperature | ^{ref} | 23617.40 | 23665.52 | 0.42 | 0.00 | | | | | 20.65 | 25.00 | 3.52 | | 0.41 | 0.15 | 0.10 | 0.10 |
| WINNETOU | Asym response to air temperature | ^{plot} | 42769.90 | 42820.42 | 0.47 | 2.70 | | | 0.73 | 1.00 | | | | | 0.46 | 0.06 | 0.13 | 0.14 |
| WINNETOU | Asym response to air temperature | ^{ref} | 42766.50 | 42817.01 | 0.47 | 2.45 | | | 0.76 | 1.00 | | | | | 0.46 | 0.05 | 0.12 | 0.13 |
| WINNETOU | Asym response to soil temperature | ^{plot} | 42476.80 | 42527.30 | 0.47 | 3.45 | | | 1.02 | 1.00 | | | | | 0.46 | 0.05 | 0.12 | 0.13 |
| WINNETOU | Asym response to soil temperature | ^{ref} | 42952.60 | 43003.13 | 0.47 | 4.98 | | | 1.51 | 1.00 | | | | | 0.46 | 0.06 | 0.13 | 0.14 |
| WINNETOU | Bi-linear response to air temperature | ^{plot} | 41890.20 | 41949.08 | 0.46 | 10.74 | 0.10 | 0.31 | | | | | | | 0.45 | 0.03 | 0.09 | 0.11 |
| WINNETOU | Bi-linear response to air temperature | ^{ref} | 41908.40 | 41967.34 | 0.46 | 11.63 | 0.11 | 0.38 | | | | | | | 0.45 | 0.03 | 0.09 | 0.11 |
| WINNETOU | Bi-linear response to soil temperature | ^{plot} | 41423.30 | 41482.22 | 0.45 | 9.66 | 0.11 | 0.31 | | | | | | | 0.45 | 0.01 | 0.08 | 0.09 |
| WINNETOU | Bi-linear response to soil temperature | ^{ref} | 42399.40 | 42458.31 | 0.47 | 7.33 | 0.17 | 0.17 | | | | | | | 0.46 | 0.04 | 0.11 | 0.13 |
| WINNETOU | Gaussian process | ^{plot} | 44666.20 | 44708.23 | 0.51 | | | | | | | | | 0.24 | 0.47 | 0.12 | 0.20 | 0.20 |
| WINNETOU | Gaussian process | ^{ref} | 44670.20 | 44712.24 | 0.51 | | | | | | | | | 0.24 | 0.47 | 0.12 | 0.20 | 0.20 |
| WINNETOU | Linear response to air temperature | ^{plot} | 42373.50 | 42424.03 | 0.47 | 2.57 | 0.05 | | | | | | | | 0.46 | 0.04 | 0.11 | 0.12 |
| WINNETOU | Linear response to air temperature | ^{ref} | 42374.90 | 42425.43 | 0.46 | 2.08 | 0.05 | | | | | | | | 0.46 | 0.04 | 0.11 | 0.12 |
| WINNETOU | Linear response to soil temperature | ^{plot} | 41977.70 | 42028.18 | 0.46 | 2.72 | 0.06 | | | | | | | | 0.45 | 0.03 | 0.10 | 0.11 |
| WINNETOU | Linear response to soil temperature | ^{ref} | 42648.40 | 42698.86 | 0.47 | 4.83 | 0.11 | | | | | | | | 0.46 | 0.05 | 0.12 | 0.13 |
| WINNETOU | Thermal time to air temperature | ^{plot} | 42554.70 | 42596.76 | 0.47 | 0.00 | 0.04 | | | | | | | | 0.46 | 0.05 | 0.12 | 0.13 |
| WINNETOU | Thermal time to air temperature | ^{ref} | 42501.60 | 42543.70 | 0.47 | 0.00 | 0.04 | | | | | | | | 0.46 | 0.05 | 0.12 | 0.13 |
| WINNETOU | Thermal time to soil temperature | ^{plot} | 42305.30 | 42347.34 | 0.46 | 0.00 | 0.04 | | | | | | | | 0.46 | 0.04 | 0.11 | 0.12 |
| WINNETOU | Thermal time to soil temperature | ^{ref} | 43634.50 | 43676.58 | 0.49 | 0.00 | 0.04 | | | | | | | | 0.47 | 0.09 | 0.16 | 0.17 |
| WINNETOU | Wang-Engel response to air temperature | ^{plot} | 42353.70 | 42404.23 | 0.46 | 0.00 | | | | | 18.95 | 25.00 | 0.93 | | 0.46 | 0.04 | 0.11 | 0.12 |
| WINNETOU | Wang-Engel response to air temperature | ^{ref} | 42368.00 | 42418.48 | 0.46 | 0.00 | | | | | 18.67 | 25.00 | 0.93 | | 0.46 | 0.04 | 0.11 | 0.12 |
| WINNETOU | Wang-Engel response to soil temperature | ^{plot} | 41802.00 | 41852.51 | 0.46 | 0.00 | | | | | 18.72 | 25.00 | 1.07 | | 0.45 | 0.02 | 0.09 | 0.10 |
| WINNETOU | Wang-Engel response to soil temperature | ^{ref} | 42332.60 | 42383.11 | 0.46 | 0.00 | | | | | 20.00 | 25.00 | 2.69 | | 0.46 | 0.04 | 0.11 | 0.12 |

Table B.3: Fitted non-linear models to leaf growth tracker (MARTRACK) soybean data. Genotype=ALL denotes the species-specific models.

| genotype_name | model | covariate_ | AIC | BIC | RMSE | Tmin | a | rmin | lrc | Asym | Topt | Tmax | rmax | mu | sigma | rho_1 | rho_2 | rho_3 |
|---|---|---|---|---|---|---|---|---|---|---|---|---|---|---|---|---|---|---|
| ALL | Asym response to air temperature | T[air]^{ref} | 2422.50 | 2463.24 | 0.70 | 7.97 | | | 1.04 | 1.74 | | | | | 0.29 | 1.00 | -0.11 | 0.00 |
| ALL | Asym response to soil temperature | T[soil]^{ref} | 2450.90 | 2491.55 | 0.73 | 15.20 | | | 1.44 | 1.88 | | | | | 0.29 | 1.01 | -0.11 | 0.00 |
| ALL | Bi-linear response to air temperature | T[air]^{ref} | 2441.60 | 2482.29 | 0.70 | 6.79 | 0.06 | 0.29 | | | | | | | 0.29 | 1.00 | -0.11 | 0.00 |
| ALL | Bi-linear response to soil temperature | T[soil]^{ref} | 2467.00 | 2514.42 | 0.73 | 9.39 | 0.14 | -0.62 | | | | | | | 0.29 | 1.01 | -0.11 | 0.00 |
| ALL | Gaussian process | | 2575.90 | 2602.99 | 0.85 | | | | | | | | | 0.87 | 0.30 | 1.02 | -0.10 | 0.00 |
| ALL | Linear response to air temperature | T[air]^{ref} | 13203.60 | 13223.97 | 0.69 | 6.84 | 0.10 | | | | | | | | 0.67 | 0.00 | 0.00 | 0.00 |
| ALL | Linear response to soil temperature | T[soil]^{ref} | 13952.40 | 13972.73 | 0.72 | 14.76 | 0.17 | | | | | | | | 0.71 | 0.00 | 0.00 | 0.00 |
| ALL | Thermal time to air temperature | T[air]^{ref} | 2446.20 | 2473.34 | 0.70 | 5.00 | 0.07 | | | | | | | | 0.29 | 1.00 | -0.11 | 0.00 |
| ALL | Thermal time to soil temperature | T[soil]^{ref} | 2502.80 | 2529.97 | 0.77 | 5.00 | 0.06 | | | | | | | | 0.29 | 1.01 | -0.10 | 0.00 |
| ALL | Wang-Engel response to air temperature | T[air]^{ref} | 2398.30 | 2445.77 | 0.68 | 2.65 | | | | | 23.72 | 32.83 | 1.40 | | 0.29 | 1.00 | -0.11 | 0.00 |
| ALL | Wang-Engel response to soil temperature | T[soil]^{ref} | 2448.50 | 2495.99 | 0.72 | 15.41 | | | | | 24.18 | 30.91 | 1.33 | | 0.29 | 1.01 | -0.11 | 0.00 |
| Castetis | Asym response to air temperature | T[air]^{ref} | 457.40 | 494.19 | 0.74 | 7.41 | | | 1.30 | 1.67 | | | | | 0.28 | 1.09 | -0.20 | 0.01 |
| Castetis | Asym response to soil temperature | T[soil]^{ref} | 481.30 | 512.84 | 0.81 | 14.65 | | | 1.92 | 1.48 | | | | | 0.29 | 1.09 | -0.17 | 0.00 |
| Castetis | Bi-linear response to air temperature | T[air]^{ref} | 448.90 | 480.43 | 0.74 | 22.28 | -0.05 | 1.51 | | | | | | | 0.28 | 1.08 | -0.18 | 0.00 |
| Castetis | Bi-linear response to soil temperature | T[soil]^{ref} | 486.80 | 523.60 | 0.81 | 11.09 | 0.09 | 0.14 | | | | | | | 0.29 | 1.09 | -0.19 | 0.01 |
| Castetis | Gaussian process | | 491.60 | 512.60 | 0.91 | | | | | | | | | 0.99 | 0.29 | 1.09 | -0.16 | 0.00 |
| Castetis | Linear response to air temperature | T[air]^{ref} | 2967.50 | 2983.30 | 0.72 | 5.80 | 0.10 | | | | | | | | 0.70 | 0.00 | 0.00 | 0.00 |
| Castetis | Linear response to soil temperature | T[soil]^{ref} | 483.20 | 509.41 | 0.81 | 9.67 | 0.09 | | | | | | | | 0.29 | 1.08 | -0.17 | 0.00 |
| Castetis | Thermal time to air temperature | T[air]^{ref} | 465.60 | 486.63 | 0.74 | 5.00 | 0.08 | | | | | | | | 0.29 | 1.09 | -0.19 | 0.00 |
| Castetis | Thermal time to soil temperature | T[soil]^{ref} | 482.20 | 503.18 | 0.83 | 5.00 | 0.07 | | | | | | | | 0.29 | 1.09 | -0.17 | 0.00 |
| Castetis | Wang-Engel response to air temperature | T[air]^{ref} | 449.90 | 486.66 | 0.71 | -0.54 | | | | | 23.03 | 31.29 | 1.53 | | 0.28 | 1.08 | -0.19 | 0.00 |
| Castetis | Wang-Engel response to soil temperature | T[soil]^{ref} | 477.20 | 514.00 | 0.80 | -29.92 | | | | | 23.36 | 26.51 | 1.39 | | 0.29 | 1.09 | -0.18 | 0.00 |
| Gallec | Asym response to air temperature | T[air]^{ref} | 280.20 | 315.57 | 0.65 | 8.32 | | | 0.90 | 1.87 | | | | | 0.25 | 1.06 | -0.17 | 0.00 |
| Gallec | Asym response to soil temperature | T[soil]^{ref} | 290.40 | 325.78 | 0.69 | 15.42 | | | 1.85 | 1.52 | | | | | 0.26 | 1.07 | -0.16 | 0.00 |
| Gallec | Bi-linear response to air temperature | T[air]^{ref} | 288.20 | 323.56 | 0.66 | 11.25 | 0.07 | 0.48 | | | | | | | 0.25 | 1.06 | -0.16 | 0.00 |
| Gallec | Bi-linear response to soil temperature | T[soil]^{ref} | 303.70 | 339.07 | 0.68 | 11.65 | 0.14 | -0.38 | | | | | | | 0.26 | 1.07 | -0.16 | 0.00 |
| Gallec | Gaussian process | | 356.70 | 380.31 | 0.80 | | | | | | | | | 0.85 | 0.26 | 1.09 | -0.15 | 0.00 |
| Gallec | Linear response to air temperature | T[air]^{ref} | 287.80 | 317.21 | 0.66 | 3.39 | 0.07 | | | | | | | | 0.25 | 1.06 | -0.16 | 0.00 |
| Gallec | Linear response to soil temperature | T[soil]^{ref} | 303.60 | 338.95 | 0.68 | 14.31 | 0.14 | | | | | | | | 0.26 | 1.07 | -0.16 | -0.01 |
| Gallec | Thermal time to air temperature | T[air]^{ref} | 287.20 | 310.79 | 0.65 | 5.00 | 0.07 | | | | | | | | 0.26 | 1.06 | -0.16 | 0.00 |
| Gallec | Thermal time to soil temperature | T[soil]^{ref} | 322.20 | 345.80 | 0.74 | 5.00 | 0.06 | | | | | | | | 0.26 | 1.08 | -0.16 | 0.00 |
| Gallec | Wang-Engel response to air temperature | T[air]^{ref} | 263.40 | 304.65 | 0.63 | 6.93 | | | | | 23.64 | 32.75 | 1.42 | | 0.25 | 1.06 | -0.17 | 0.00 |
| Gallec | Wang-Engel response to soil temperature | T[soil]^{ref} | 291.10 | 332.35 | 0.69 | 15.86 | | | | | 25.10 | 36.37 | 1.27 | | 0.25 | 1.07 | -0.16 | 0.00 |
| Opaline | Asym response to air temperature | T[air]^{ref} | 1498.40 | 1527.37 | 0.71 | 7.97 | | | 1.01 | 1.71 | | | | | 0.33 | 0.88 | 0.00 | 0.00 |
| Opaline | Asym response to soil temperature | T[soil]^{ref} | 1492.30 | 1521.28 | 0.72 | 15.73 | | | 1.04 | 2.60 | | | | | 0.33 | 0.88 | 0.00 | 0.00 |
| Opaline | Bi-linear response to air temperature | T[air]^{ref} | 1502.60 | 1531.57 | 0.72 | 10.09 | 0.06 | 0.45 | | | | | | | 0.33 | 0.88 | 0.00 | 0.00 |
| Opaline | Bi-linear response to soil temperature | T[soil]^{ref} | 1494.70 | 1523.65 | 0.72 | 13.27 | 0.16 | -0.23 | | | | | | | 0.33 | 0.88 | 0.00 | 0.00 |
| Opaline | Gaussian process | | 1544.00 | 1561.34 | 0.84 | | | | | | | | | 0.83 | 0.33 | 0.91 | 0.00 | 0.00 |
| Opaline | Linear response to air temperature | T[air]^{ref} | 1501.20 | 1524.34 | 0.72 | 2.29 | 0.06 | | | | | | | | 0.33 | 0.88 | 0.00 | 0.00 |
| Opaline | Linear response to soil temperature | T[soil]^{ref} | 1492.70 | 1515.85 | 0.72 | 14.76 | 0.16 | | | | | | | | 0.33 | 0.88 | 0.00 | 0.00 |
| Opaline | Thermal time to air temperature | T[air]^{ref} | 1501.70 | 1519.06 | 0.71 | 5.00 | 0.07 | | | | | | | | 0.33 | 0.88 | 0.00 | 0.00 |
| Opaline | Thermal time to soil temperature | T[soil]^{ref} | 1514.30 | 1531.71 | 0.77 | 5.00 | 0.06 | | | | | | | | 0.33 | 0.90 | 0.00 | 0.00 |
| Opaline | Wang-Engel response to air temperature | T[air]^{ref} | 1497.30 | 1532.04 | 0.71 | 2.80 | | | | | 24.86 | 35.86 | 1.34 | | 0.33 | 0.87 | 0.00 | 0.00 |
| Opaline | Wang-Engel response to soil temperature | T[soil]^{ref} | 1493.50 | 1528.26 | 0.72 | 13.22 | | | | | 24.43 | 28.95 | 1.42 | | 0.33 | 0.88 | 0.00 | 0.00 |

Table B.4: Fitted non-linear models to plant height data (SfM, TLS) wheat and soybean data Genotype=ALL denotes the species-specific models.

| crop | genotype.name | model | covariate_ | model_ | skipped_year | AIC | BIC | RMSE | Tmin | a | rmin | lrc | Asym | Topt | rmax | mu | sigma_error | rho_error |
|---|---|---|---|---|---|---|---|---|---|---|---|---|---|---|---|---|---|---|
| Soybean | ALL | asym | T[air]^{plot} | Asymptotic | 2017.00 | 6665.57 | 6689.43 | | 6.37 | | | 0.00 | 2.84 | | | | 13.21 | 0.51 |
| Soybean | ALL | asym | T[air]^{plot} | Asymptotic | 2018.00 | 6294.67 | 6318.30 | | 7.99 | | | 0.86 | 1.69 | | | | 12.79 | 0.49 |
| Soybean | ALL | asym | T[air]^{plot} | Asymptotic | 2019.00 | 7357.61 | 7382.12 | 12.98 | 9.67 | | | 2.67 | 0.96 | | | | 9.75 | 0.66 |
| Soybean | ALL | asym | T[air]^{plot} | Asymptotic | 2020.00 | 6287.73 | 6311.35 | | 0.00 | | | 0.12 | 2.02 | | | | 12.73 | 0.47 |
| Soybean | ALL | asym | T[air]^{ref} | Asymptotic | 2017.00 | 6665.57 | 6689.43 | | 6.37 | | | 0.00 | 2.84 | | | | 13.21 | 0.51 |
| Soybean | ALL | asym | T[air]^{ref} | Asymptotic | 2018.00 | 6294.67 | 6318.30 | | 7.99 | | | 0.86 | 1.69 | | | | 12.79 | 0.49 |
| Soybean | ALL | asym | T[air]^{ref} | Asymptotic | 2019.00 | 7357.61 | 7382.12 | 12.98 | 9.67 | | | 2.67 | 0.96 | | | | 9.75 | 0.66 |
| Soybean | ALL | asym | T[air]^{ref} | Asymptotic | 2020.00 | 6287.73 | 6311.35 | | 0.00 | | | 0.12 | 2.02 | | | | 12.73 | 0.47 |
| Soybean | ALL | bilinear | T[air]^{plot} | Bi-linear | 2017.00 | 6668.06 | 6691.92 | | 11.46 | 0.06 | 0.49 | | | | | | 13.23 | 0.53 |
| Soybean | ALL | bilinear | T[air]^{plot} | Bi-linear | 2018.00 | 6314.86 | 6338.48 | | 29.91 | -0.02 | 1.46 | | | | | | 12.95 | 0.57 |
| Soybean | ALL | bilinear | T[air]^{plot} | Bi-linear | 2019.00 | 7400.20 | 7424.72 | 13.70 | 17.87 | -0.00 | 0.98 | | | | | | 9.96 | 0.69 |
| Soybean | ALL | bilinear | T[air]^{plot} | Bi-linear | 2020.00 | 6296.65 | 6320.27 | | 29.87 | -0.24 | 1.61 | | | | | | 12.80 | 0.45 |
| Soybean | ALL | bilinear | T[air]^{ref} | Bi-linear | 2017.00 | 6668.06 | 6691.92 | | 11.46 | 0.06 | 0.49 | | | | | | 13.23 | 0.53 |
| Soybean | ALL | bilinear | T[air]^{ref} | Bi-linear | 2018.00 | 6314.86 | 6338.48 | | 29.91 | -0.02 | 1.46 | | | | | | 12.95 | 0.57 |
| Soybean | ALL | bilinear | T[air]^{ref} | Bi-linear | 2019.00 | 7400.20 | 7424.72 | 13.70 | 17.87 | -0.00 | 0.98 | | | | | | 9.96 | 0.69 |
| Soybean | ALL | bilinear | T[air]^{ref} | Bi-linear | 2020.00 | 6296.65 | 6320.27 | | 29.87 | -0.24 | 1.61 | | | | | | 12.80 | 0.45 |
| Soybean | ALL | gauss | ^{plot} | Gaussian | 2017.00 | 8836.45 | 8850.77 | 38.56 | | | | | | | | 68.34 | 38.26 | 0.13 |
| Soybean | ALL | gauss | ^{plot} | Gaussian | 2018.00 | 8330.93 | 8345.11 | 36.18 | | | | | | | | 73.48 | 36.02 | 0.10 |
| Soybean | ALL | gauss | ^{plot} | Gaussian | 2019.00 | 9002.85 | 9017.56 | 22.34 | | | | | | | | 59.52 | 22.35 | 0.02 |
| Soybean | ALL | gauss | ^{plot} | Gaussian | 2020.00 | 8376.36 | 8390.53 | 37.37 | | | | | | | | 71.89 | 37.02 | 0.14 |
| Soybean | ALL | gauss | ^{ref} | Gaussian | 2017.00 | 8836.45 | 8850.77 | 38.56 | | | | | | | | 68.34 | 38.26 | 0.13 |
| Soybean | ALL | gauss | ^{ref} | Gaussian | 2018.00 | 8330.93 | 8345.11 | 36.18 | | | | | | | | 73.48 | 36.02 | 0.10 |
| Soybean | ALL | gauss | ^{ref} | Gaussian | 2019.00 | 9002.85 | 9017.56 | 22.34 | | | | | | | | 59.52 | 22.35 | 0.02 |
| Soybean | ALL | gauss | ^{ref} | Gaussian | 2020.00 | 8376.36 | 8390.53 | 37.37 | | | | | | | | 71.89 | 37.02 | 0.14 |
| Soybean | ALL | linear | T[air]^{plot} | Linear | 2017.00 | 6964.41 | 6983.49 | | -45.89 | 0.01 | | | | | | | 15.96 | 0.69 |
| Soybean | ALL | linear | T[air]^{plot} | Linear | 2018.00 | 6572.55 | 6591.45 | | -46.28 | 0.01 | | | | | | | 15.39 | 0.69 |
| Soybean | ALL | linear | T[air]^{plot} | Linear | 2019.00 | 7465.38 | 7485.00 | 14.06 | -17.95 | 0.02 | | | | | | | 10.30 | 0.68 |
| Soybean | ALL | linear | T[air]^{plot} | Linear | 2020.00 | 6334.16 | 6353.06 | | -47.80 | 0.02 | | | | | | | 13.19 | 0.49 |
| Soybean | ALL | linear | T[air]^{ref} | Linear | 2017.00 | 6964.41 | 6983.49 | | -45.89 | 0.01 | | | | | | | 15.96 | 0.69 |
| Soybean | ALL | linear | T[air]^{ref} | Linear | 2018.00 | 6572.55 | 6591.45 | | -46.28 | 0.01 | | | | | | | 15.39 | 0.69 |
| Soybean | ALL | linear | T[air]^{ref} | Linear | 2019.00 | 7465.38 | 7485.00 | 14.06 | -17.95 | 0.02 | | | | | | | 10.30 | 0.68 |
| Soybean | ALL | linear | T[air]^{ref} | Linear | 2020.00 | 6334.16 | 6353.06 | | -47.80 | 0.02 | | | | | | | 13.19 | 0.49 |
| Soybean | ALL | thermal | T[air]^{plot} | Thermal time | 2017.00 | 6680.51 | 6694.83 | | | | | | | | | | | |
| Soybean | ALL | thermal | T[air]^{plot} | Thermal time | 2018.00 | 6312.92 | 6327.10 | | | | | | | | | | | |
| Soybean | ALL | thermal | T[air]^{plot} | Thermal time | 2019.00 | 7557.34 | 7572.04 | 12.99 | | | | | | | | | | |
| Soybean | ALL | thermal | T[air]^{plot} | Thermal time | 2020.00 | 6330.86 | 6345.04 | | | | | | | | | | | |
| Soybean | ALL | thermal | T[air]^{ref} | Thermal time | 2017.00 | 6680.51 | 6694.83 | | | | | | | | | | | |
| Soybean | ALL | thermal | T[air]^{ref} | Thermal time | 2018.00 | 6312.92 | 6327.10 | | | | | | | | | | | |
| Soybean | ALL | thermal | T[air]^{ref} | Thermal time | 2019.00 | 7557.34 | 7572.04 | 12.99 | | | | | | | | | | |
| Soybean | ALL | thermal | T[air]^{ref} | Thermal time | 2020.00 | 6330.86 | 6345.04 | | | | | | | | | | | |
| Soybean | ALL | wang | T[air]^{plot} | Wang-Engel | 2017.00 | 6729.78 | 6748.87 | | | | | | | 26.39 | 1.49 | | 13.74 | 0.45 |
| Soybean | ALL | wang | T[air]^{plot} | Wang-Engel | 2018.00 | 6319.43 | 6338.33 | | | | | | | 26.24 | 1.44 | | 13.01 | 0.47 |
| Soybean | ALL | wang | T[air]^{plot} | Wang-Engel | 2019.00 | 7394.56 | 7414.17 | 13.31 | | | | | | 22.54 | 1.06 | | 9.94 | 0.67 |
| Soybean | ALL | wang | T[air]^{plot} | Wang-Engel | 2020.00 | 6369.39 | 6388.29 | | | | | | | 25.08 | 1.40 | | 13.42 | 0.40 |
| Soybean | ALL | wang | T[air]^{ref} | Wang-Engel | 2017.00 | 6729.78 | 6748.87 | | | | | | | 26.39 | 1.49 | | 13.74 | 0.45 |
| Soybean | ALL | wang | T[air]^{ref} | Wang-Engel | 2018.00 | 6319.43 | 6338.33 | | | | | | | 26.24 | 1.44 | | 13.01 | 0.47 |
| Soybean | ALL | wang | T[air]^{ref} | Wang-Engel | 2019.00 | 7394.56 | 7414.17 | 13.31 | | | | | | 22.54 | 1.06 | | 9.94 | 0.67 |

| | | | | | | | | | | | | | | | | | | |
|---|---|---|---|---|---|---|---|---|---|---|---|---|---|---|---|---|---|---|
| Soybean | ALL | wang | T[air]^{ref} | Wang-Engel | 2020.00 | 6369.39 | 6388.29 | | | | | | | 25.08 | 1.40 | | 13.42 | 0.40 |
| Soybean | Aveline | asym | T[air]^{plot} | Asymptotic | 2017.00 | 523.26 | 534.57 | | 7.85 | | | 0.68 | 1.94 | | | | 11.82 | 0.47 |
| Soybean | Aveline | asym | T[air]^{plot} | Asymptotic | 2018.00 | 513.63 | 524.87 | | 9.24 | | | 1.55 | 1.32 | | | | 11.64 | 0.59 |
| Soybean | Aveline | asym | T[air]^{plot} | Asymptotic | 2019.00 | 595.86 | 607.89 | 12.10 | 9.72 | | | 2.86 | 0.94 | | | | 9.00 | 0.67 |
| Soybean | Aveline | asym | T[air]^{plot} | Asymptotic | 2020.00 | 503.28 | 514.38 | | 4.01 | | | 3.90 | 1.09 | | | | 12.13 | 0.53 |
| Soybean | Aveline | asym | T[air]^{ref} | Asymptotic | 2017.00 | 523.26 | 534.57 | | 7.85 | | | 0.68 | 1.94 | | | | 11.82 | 0.47 |
| Soybean | Aveline | asym | T[air]^{ref} | Asymptotic | 2018.00 | 513.63 | 524.87 | | 9.24 | | | 1.55 | 1.32 | | | | 11.64 | 0.59 |
| Soybean | Aveline | asym | T[air]^{ref} | Asymptotic | 2019.00 | 595.86 | 607.89 | 12.10 | 9.72 | | | 2.86 | 0.94 | | | | 9.00 | 0.67 |
| Soybean | Aveline | asym | T[air]^{ref} | Asymptotic | 2020.00 | 503.28 | 514.38 | | 4.01 | | | 3.90 | 1.09 | | | | 12.13 | 0.53 |
| Soybean | Aveline | bilinear | T[air]^{plot} | Bi-linear | 2017.00 | 524.51 | 535.82 | | 8.20 | 0.06 | 0.30 | | | | | | 11.95 | 0.51 |
| Soybean | Aveline | bilinear | T[air]^{plot} | Bi-linear | 2018.00 | 518.41 | 529.65 | | 15.21 | 0.04 | 0.78 | | | | | | 12.08 | 0.64 |
| Soybean | Aveline | bilinear | T[air]^{plot} | Bi-linear | 2019.00 | 599.13 | 611.17 | 13.13 | 17.93 | -0.02 | 1.01 | | | | | | 9.19 | 0.72 |
| Soybean | Aveline | bilinear | T[air]^{plot} | Bi-linear | 2020.00 | 492.25 | 503.35 | | 5.00 | 0.03 | 0.69 | | | | | | 11.00 | 0.55 |
| Soybean | Aveline | bilinear | T[air]^{ref} | Bi-linear | 2017.00 | 524.51 | 535.82 | | 8.20 | 0.06 | 0.30 | | | | | | 11.95 | 0.51 |
| Soybean | Aveline | bilinear | T[air]^{ref} | Bi-linear | 2018.00 | 518.41 | 529.65 | | 15.21 | 0.04 | 0.78 | | | | | | 12.08 | 0.64 |
| Soybean | Aveline | bilinear | T[air]^{ref} | Bi-linear | 2019.00 | 599.13 | 611.17 | 13.13 | 17.93 | -0.02 | 1.01 | | | | | | 9.19 | 0.72 |
| Soybean | Aveline | bilinear | T[air]^{ref} | Bi-linear | 2020.00 | 492.25 | 503.35 | | 5.00 | 0.03 | 0.69 | | | | | | 11.00 | 0.55 |
| Soybean | Aveline | gauss | ^{plot} | Gaussian | 2017.00 | 715.31 | 722.10 | 38.54 | | | | | | | | 69.43 | 38.38 | 0.11 |
| Soybean | Aveline | gauss | ^{plot} | Gaussian | 2018.00 | 697.48 | 704.22 | 36.47 | | | | | | | | 73.46 | 36.30 | 0.11 |
| Soybean | Aveline | gauss | ^{plot} | Gaussian | 2019.00 | 735.30 | 742.52 | 21.80 | | | | | | | | 59.84 | 21.82 | -0.01 |
| Soybean | Aveline | gauss | ^{plot} | Gaussian | 2020.00 | 680.34 | 687.00 | 37.30 | | | | | | | | 72.86 | 37.10 | 0.12 |
| Soybean | Aveline | gauss | ^{ref} | Gaussian | 2017.00 | 715.31 | 722.10 | 38.54 | | | | | | | | 69.43 | 38.38 | 0.11 |
| Soybean | Aveline | gauss | ^{ref} | Gaussian | 2018.00 | 697.48 | 704.22 | 36.47 | | | | | | | | 73.46 | 36.30 | 0.11 |
| Soybean | Aveline | gauss | ^{ref} | Gaussian | 2019.00 | 735.30 | 742.52 | 21.80 | | | | | | | | 59.84 | 21.82 | -0.01 |
| Soybean | Aveline | gauss | ^{ref} | Gaussian | 2020.00 | 680.34 | 687.00 | 37.30 | | | | | | | | 72.86 | 37.10 | 0.12 |
| Soybean | Aveline | linear | T[air]^{plot} | Linear | 2017.00 | 550.09 | 559.14 | | -46.66 | 0.01 | | | | | | | 14.83 | 0.72 |
| Soybean | Aveline | linear | T[air]^{plot} | Linear | 2018.00 | 536.41 | 545.40 | | -47.09 | 0.01 | | | | | | | 14.21 | 0.74 |
| Soybean | Aveline | linear | T[air]^{plot} | Linear | 2019.00 | 606.75 | 616.38 | 14.24 | -50.16 | 0.01 | | | | | | | 9.76 | 0.73 |
| Soybean | Aveline | linear | T[air]^{plot} | Linear | 2020.00 | 492.01 | 500.89 | | -49.31 | 0.02 | | | | | | | 11.29 | 0.55 |
| Soybean | Aveline | linear | T[air]^{ref} | Linear | 2017.00 | 550.09 | 559.14 | | -46.66 | 0.01 | | | | | | | 14.83 | 0.72 |
| Soybean | Aveline | linear | T[air]^{ref} | Linear | 2018.00 | 536.41 | 545.40 | | -47.09 | 0.01 | | | | | | | 14.21 | 0.74 |
| Soybean | Aveline | linear | T[air]^{ref} | Linear | 2019.00 | 606.75 | 616.38 | 14.24 | -50.16 | 0.01 | | | | | | | 9.76 | 0.73 |
| Soybean | Aveline | linear | T[air]^{ref} | Linear | 2020.00 | 492.01 | 500.89 | | -49.31 | 0.02 | | | | | | | 11.29 | 0.55 |
| Soybean | Aveline | thermal | T[air]^{plot} | Thermal time | 2017.00 | 521.85 | 528.64 | | | | | | | | | | | |
| Soybean | Aveline | thermal | T[air]^{plot} | Thermal time | 2018.00 | 514.87 | 521.62 | | | | | | | | | | | |
| Soybean | Aveline | thermal | T[air]^{plot} | Thermal time | 2019.00 | 617.03 | 624.25 | 12.43 | | | | | | | | | | |
| Soybean | Aveline | thermal | T[air]^{plot} | Thermal time | 2020.00 | 500.12 | 506.78 | | | | | | | | | | | |
| Soybean | Aveline | thermal | T[air]^{ref} | Thermal time | 2017.00 | 521.85 | 528.64 | | | | | | | | | | | |
| Soybean | Aveline | thermal | T[air]^{ref} | Thermal time | 2018.00 | 514.87 | 521.62 | | | | | | | | | | | |
| Soybean | Aveline | thermal | T[air]^{ref} | Thermal time | 2019.00 | 617.03 | 624.25 | 12.43 | | | | | | | | | | |
| Soybean | Aveline | thermal | T[air]^{ref} | Thermal time | 2020.00 | 500.12 | 506.78 | | | | | | | | | | | |
| Soybean | Aveline | wang | T[air]^{plot} | Wang-Engel | 2017.00 | 526.87 | 535.92 | | | | | | | 26.32 | 1.49 | | 12.33 | 0.39 |
| Soybean | Aveline | wang | T[air]^{plot} | Wang-Engel | 2018.00 | 515.62 | 524.61 | | | | | | | 25.69 | 1.38 | | 12.01 | 0.57 |
| Soybean | Aveline | wang | T[air]^{plot} | Wang-Engel | 2019.00 | 596.62 | 606.24 | 12.68 | | | | | | 21.78 | 1.02 | | 9.16 | 0.70 |
| Soybean | Aveline | wang | T[air]^{plot} | Wang-Engel | 2020.00 | 502.61 | 511.49 | | | | | | | 24.29 | 1.35 | | 12.26 | 0.44 |
| Soybean | Aveline | wang | T[air]^{ref} | Wang-Engel | 2017.00 | 526.87 | 535.92 | | | | | | | 26.32 | 1.49 | | 12.33 | 0.39 |
| Soybean | Aveline | wang | T[air]^{ref} | Wang-Engel | 2018.00 | 515.62 | 524.61 | | | | | | | 25.69 | 1.38 | | 12.01 | 0.57 |
| Soybean | Aveline | wang | T[air]^{ref} | Wang-Engel | 2019.00 | 596.62 | 606.24 | 12.68 | | | | | | 21.78 | 1.02 | | 9.16 | 0.70 |
| Soybean | Aveline | wang | T[air]^{ref} | Wang-Engel | 2020.00 | 502.61 | 511.49 | | | | | | | 24.29 | 1.35 | | 12.26 | 0.44 |
| Soybean | Coraline | asym | T[air]^{plot} | Asymptotic | 2017.00 | 592.38 | 603.97 | | 8.96 | | | 0.87 | 1.77 | | | | 13.81 | 0.45 |
| Soybean | Coraline | asym | T[air]^{plot} | Asymptotic | 2018.00 | 553.54 | 564.85 | | 9.65 | | | 1.21 | 1.51 | | | | 13.17 | 0.42 |
| Soybean | Coraline | asym | T[air]^{plot} | Asymptotic | 2019.00 | 615.76 | 627.91 | 12.10 | 9.91 | | | 2.49 | 0.98 | | | | 9.30 | 0.64 |

| | | | | | | | | | | | | | | | | | | |
|---|---|---|---|---|---|---|---|---|---|---|---|---|---|---|---|---|---|---|
| Soybean | Coraline | asym | T[air]^{plot} | Asymptotic | 2020.00 | 551.14 | 562.38 | | 4.01 | | | 3.89 | 1.04 | | | | 13.73 | 0.38 |
| Soybean | Coraline | asym | T[air]^{ref} | Asymptotic | 2017.00 | 592.38 | 603.97 | | 8.96 | | | 0.87 | 1.77 | | | | 13.81 | 0.45 |
| Soybean | Coraline | asym | T[air]^{ref} | Asymptotic | 2018.00 | 553.54 | 564.85 | | 9.65 | | | 1.21 | 1.51 | | | | 13.17 | 0.42 |
| Soybean | Coraline | asym | T[air]^{ref} | Asymptotic | 2019.00 | 615.76 | 627.91 | 12.10 | 9.91 | | | 2.49 | 0.98 | | | | 9.30 | 0.64 |
| Soybean | Coraline | asym | T[air]^{ref} | Asymptotic | 2020.00 | 551.14 | 562.38 | | 4.01 | | | 3.89 | 1.04 | | | | 13.73 | 0.38 |
| Soybean | Coraline | bilinear | T[air]^{plot} | Bi-linear | 2017.00 | 595.46 | 607.05 | | 16.19 | 0.06 | 0.71 | | | | | | 14.11 | 0.50 |
| Soybean | Coraline | bilinear | T[air]^{plot} | Bi-linear | 2018.00 | 557.70 | 569.01 | | 9.33 | 0.06 | 0.30 | | | | | | 13.57 | 0.42 |
| Soybean | Coraline | bilinear | T[air]^{plot} | Bi-linear | 2019.00 | 620.97 | 633.13 | 12.94 | 19.89 | -0.00 | 1.00 | | | | | | 9.60 | 0.67 |
| Soybean | Coraline | bilinear | T[air]^{plot} | Bi-linear | 2020.00 | 539.86 | 551.10 | | 29.81 | -0.25 | 1.57 | | | | | | 12.62 | 0.36 |
| Soybean | Coraline | bilinear | T[air]^{ref} | Bi-linear | 2017.00 | 595.46 | 607.05 | | 16.19 | 0.06 | 0.71 | | | | | | 14.11 | 0.50 |
| Soybean | Coraline | bilinear | T[air]^{ref} | Bi-linear | 2018.00 | 557.70 | 569.01 | | 9.33 | 0.06 | 0.30 | | | | | | 13.57 | 0.42 |
| Soybean | Coraline | bilinear | T[air]^{ref} | Bi-linear | 2019.00 | 620.97 | 633.13 | 12.94 | 19.89 | -0.00 | 1.00 | | | | | | 9.60 | 0.67 |
| Soybean | Coraline | bilinear | T[air]^{ref} | Bi-linear | 2020.00 | 539.86 | 551.10 | | 29.81 | -0.25 | 1.57 | | | | | | 12.62 | 0.36 |
| Soybean | Coraline | gauss | ^{plot} | Gaussian | 2017.00 | 745.37 | 752.32 | 35.73 | | | | | | | | 67.25 | 35.76 | 0.03 |
| Soybean | Coraline | gauss | ^{plot} | Gaussian | 2018.00 | 699.86 | 706.65 | 34.34 | | | | | | | | 71.94 | 34.37 | 0.09 |
| Soybean | Coraline | gauss | ^{plot} | Gaussian | 2019.00 | 754.12 | 761.41 | 21.82 | | | | | | | | 58.95 | 21.93 | 0.03 |
| Soybean | Coraline | gauss | ^{plot} | Gaussian | 2020.00 | 692.59 | 699.34 | 34.87 | | | | | | | | 71.36 | 35.04 | 0.02 |
| Soybean | Coraline | gauss | ^{ref} | Gaussian | 2017.00 | 745.37 | 752.32 | 35.73 | | | | | | | | 67.25 | 35.76 | 0.03 |
| Soybean | Coraline | gauss | ^{ref} | Gaussian | 2018.00 | 699.86 | 706.65 | 34.34 | | | | | | | | 71.94 | 34.37 | 0.09 |
| Soybean | Coraline | gauss | ^{ref} | Gaussian | 2019.00 | 754.12 | 761.41 | 21.82 | | | | | | | | 58.95 | 21.93 | 0.03 |
| Soybean | Coraline | gauss | ^{ref} | Gaussian | 2020.00 | 692.59 | 699.34 | 34.87 | | | | | | | | 71.36 | 35.04 | 0.02 |
| Soybean | Coraline | linear | T[air]^{plot} | Linear | 2017.00 | 619.20 | 628.47 | | -45.16 | 0.01 | | | | | | | 17.01 | 0.62 |
| Soybean | Coraline | linear | T[air]^{plot} | Linear | 2018.00 | 583.54 | 592.59 | | -45.30 | 0.01 | | | | | | | 16.81 | 0.62 |
| Soybean | Coraline | linear | T[air]^{plot} | Linear | 2019.00 | 630.42 | 640.15 | 14.87 | -49.78 | 0.01 | | | | | | | 10.31 | 0.72 |
| Soybean | Coraline | linear | T[air]^{plot} | Linear | 2020.00 | 539.99 | 548.99 | | -48.27 | 0.02 | | | | | | | 12.86 | 0.39 |
| Soybean | Coraline | linear | T[air]^{ref} | Linear | 2017.00 | 619.20 | 628.47 | | -45.16 | 0.01 | | | | | | | 17.01 | 0.62 |
| Soybean | Coraline | linear | T[air]^{ref} | Linear | 2018.00 | 583.54 | 592.59 | | -45.30 | 0.01 | | | | | | | 16.81 | 0.62 |
| Soybean | Coraline | linear | T[air]^{ref} | Linear | 2019.00 | 630.42 | 640.15 | 14.87 | -49.78 | 0.01 | | | | | | | 10.31 | 0.72 |
| Soybean | Coraline | linear | T[air]^{ref} | Linear | 2020.00 | 539.99 | 548.99 | | -48.27 | 0.02 | | | | | | | 12.86 | 0.39 |
| Soybean | Coraline | thermal | T[air]^{plot} | Thermal time | 2017.00 | 593.11 | 600.06 | | | | | | | | | | | |
| Soybean | Coraline | thermal | T[air]^{plot} | Thermal time | 2018.00 | 556.48 | 563.27 | | | | | | | | | | | |
| Soybean | Coraline | thermal | T[air]^{plot} | Thermal time | 2019.00 | 625.29 | 632.58 | 12.16 | | | | | | | | | | |
| Soybean | Coraline | thermal | T[air]^{plot} | Thermal time | 2020.00 | 539.61 | 546.36 | | | | | | | | | | | |
| Soybean | Coraline | thermal | T[air]^{ref} | Thermal time | 2017.00 | 593.11 | 600.06 | | | | | | | | | | | |
| Soybean | Coraline | thermal | T[air]^{ref} | Thermal time | 2018.00 | 556.48 | 563.27 | | | | | | | | | | | |
| Soybean | Coraline | thermal | T[air]^{ref} | Thermal time | 2019.00 | 625.29 | 632.58 | 12.16 | | | | | | | | | | |
| Soybean | Coraline | thermal | T[air]^{ref} | Thermal time | 2020.00 | 539.61 | 546.36 | | | | | | | | | | | |
| Soybean | Coraline | wang | T[air]^{plot} | Wang-Engel | 2017.00 | 592.56 | 601.83 | | | | | | | 26.62 | 1.47 | | 14.02 | 0.41 |
| Soybean | Coraline | wang | T[air]^{plot} | Wang-Engel | 2018.00 | 551.76 | 560.81 | | | | | | | 26.66 | 1.45 | | 13.19 | 0.41 |
| Soybean | Coraline | wang | T[air]^{plot} | Wang-Engel | 2019.00 | 617.52 | 627.25 | 12.45 | | | | | | 23.42 | 1.09 | | 9.52 | 0.65 |
| Soybean | Coraline | wang | T[air]^{plot} | Wang-Engel | 2020.00 | 543.89 | 552.88 | | | | | | | 24.93 | 1.36 | | 13.20 | 0.31 |
| Soybean | Coraline | wang | T[air]^{ref} | Wang-Engel | 2017.00 | 592.56 | 601.83 | | | | | | | 26.62 | 1.47 | | 14.02 | 0.41 |
| Soybean | Coraline | wang | T[air]^{ref} | Wang-Engel | 2018.00 | 551.76 | 560.81 | | | | | | | 26.66 | 1.45 | | 13.19 | 0.41 |
| Soybean | Coraline | wang | T[air]^{ref} | Wang-Engel | 2019.00 | 617.52 | 627.25 | 12.45 | | | | | | 23.42 | 1.09 | | 9.52 | 0.65 |
| Soybean | Coraline | wang | T[air]^{ref} | Wang-Engel | 2020.00 | 543.89 | 552.88 | | | | | | | 24.93 | 1.36 | | 13.20 | 0.31 |
| Soybean | Falbala | asym | T[air]^{plot} | Asymptotic | 2017.00 | 620.78 | 632.63 | | 7.63 | | | 0.00 | 3.14 | | | | 13.46 | 0.49 |
| Soybean | Falbala | asym | T[air]^{plot} | Asymptotic | 2018.00 | 544.61 | 555.85 | | 7.58 | | | 0.39 | 2.28 | | | | 13.07 | 0.45 |
| Soybean | Falbala | asym | T[air]^{plot} | Asymptotic | 2019.00 | 683.03 | 695.53 | 14.29 | 9.36 | | | 2.64 | 0.99 | | | | 10.62 | 0.67 |
| Soybean | Falbala | asym | T[air]^{plot} | Asymptotic | 2020.00 | 597.66 | 609.31 | | 3.15 | | | 0.00 | 2.48 | | | | 13.55 | 0.53 |
| Soybean | Falbala | asym | T[air]^{ref} | Asymptotic | 2017.00 | 620.78 | 632.63 | | 7.63 | | | 0.00 | 3.14 | | | | 13.46 | 0.49 |
| Soybean | Falbala | asym | T[air]^{ref} | Asymptotic | 2018.00 | 544.61 | 555.85 | | 7.58 | | | 0.39 | 2.28 | | | | 13.07 | 0.45 |
| Soybean | Falbala | asym | T[air]^{ref} | Asymptotic | 2019.00 | 683.03 | 695.53 | 14.29 | 9.36 | | | 2.64 | 0.99 | | | | 10.62 | 0.67 |

| | | | | | | | | | | | | | | | | | | |
|---|---|---|---|---|---|---|---|---|---|---|---|---|---|---|---|---|---|---|
| Soybean | Falbala | asym | T[air]^{ref} | Asymptotic | 2020.00 | 597.66 | 609.31 | | 3.15 | | | 0.00 | 2.48 | | | | 13.55 | 0.53 |
| Soybean | Falbala | bilinear | T[air]^{plot} | Bi-linear | 2017.00 | 621.17 | 633.02 | | 11.44 | 0.07 | 0.42 | | | | | | 13.49 | 0.50 |
| Soybean | Falbala | bilinear | T[air]^{plot} | Bi-linear | 2018.00 | 545.86 | 557.10 | | 16.80 | 0.07 | 0.76 | | | | | | 13.20 | 0.48 |
| Soybean | Falbala | bilinear | T[air]^{plot} | Bi-linear | 2019.00 | 686.62 | 699.12 | 14.97 | 16.04 | 0.01 | 0.93 | | | | | | 10.83 | 0.69 |
| Soybean | Falbala | bilinear | T[air]^{plot} | Bi-linear | 2020.00 | 597.07 | 608.72 | | 12.51 | 0.05 | 0.71 | | | | | | 13.49 | 0.54 |
| Soybean | Falbala | bilinear | T[air]^{ref} | Bi-linear | 2017.00 | 621.17 | 633.02 | | 11.44 | 0.07 | 0.42 | | | | | | 13.49 | 0.50 |
| Soybean | Falbala | bilinear | T[air]^{ref} | Bi-linear | 2018.00 | 545.86 | 557.10 | | 16.80 | 0.07 | 0.76 | | | | | | 13.20 | 0.48 |
| Soybean | Falbala | bilinear | T[air]^{ref} | Bi-linear | 2019.00 | 686.62 | 699.12 | 14.97 | 16.04 | 0.01 | 0.93 | | | | | | 10.83 | 0.69 |
| Soybean | Falbala | bilinear | T[air]^{ref} | Bi-linear | 2020.00 | 597.07 | 608.72 | | 12.51 | 0.05 | 0.71 | | | | | | 13.49 | 0.54 |
| Soybean | Falbala | gauss | ^{plot} | Gaussian | 2017.00 | 804.25 | 811.36 | 40.74 | | | | | | | | 66.09 | 40.37 | 0.15 |
| Soybean | Falbala | gauss | ^{plot} | Gaussian | 2018.00 | 704.44 | 711.18 | 38.12 | | | | | | | | 74.87 | 38.18 | 0.10 |
| Soybean | Falbala | gauss | ^{plot} | Gaussian | 2019.00 | 828.50 | 836.00 | 24.60 | | | | | | | | 57.60 | 24.58 | 0.11 |
| Soybean | Falbala | gauss | ^{plot} | Gaussian | 2020.00 | 769.48 | 776.48 | 39.74 | | | | | | | | 69.38 | 39.29 | 0.19 |
| Soybean | Falbala | gauss | ^{ref} | Gaussian | 2017.00 | 804.25 | 811.36 | 40.74 | | | | | | | | 66.09 | 40.37 | 0.15 |
| Soybean | Falbala | gauss | ^{ref} | Gaussian | 2018.00 | 704.44 | 711.18 | 38.12 | | | | | | | | 74.87 | 38.18 | 0.10 |
| Soybean | Falbala | gauss | ^{ref} | Gaussian | 2019.00 | 828.50 | 836.00 | 24.60 | | | | | | | | 57.60 | 24.58 | 0.11 |
| Soybean | Falbala | gauss | ^{ref} | Gaussian | 2020.00 | 769.48 | 776.48 | 39.74 | | | | | | | | 69.38 | 39.29 | 0.19 |
| Soybean | Falbala | linear | T[air]^{plot} | Linear | 2017.00 | 655.43 | 664.90 | | -44.97 | 0.02 | | | | | | | 17.29 | 0.66 |
| Soybean | Falbala | linear | T[air]^{plot} | Linear | 2018.00 | 573.42 | 582.41 | | -45.43 | 0.02 | | | | | | | 16.63 | 0.65 |
| Soybean | Falbala | linear | T[air]^{plot} | Linear | 2019.00 | 690.98 | 700.98 | 16.40 | -49.14 | 0.01 | | | | | | | 11.24 | 0.73 |
| Soybean | Falbala | linear | T[air]^{plot} | Linear | 2020.00 | 606.76 | 616.08 | | -47.04 | 0.02 | | | | | | | 14.69 | 0.57 |
| Soybean | Falbala | linear | T[air]^{ref} | Linear | 2017.00 | 655.43 | 664.90 | | -44.97 | 0.02 | | | | | | | 17.29 | 0.66 |
| Soybean | Falbala | linear | T[air]^{ref} | Linear | 2018.00 | 573.42 | 582.41 | | -45.43 | 0.02 | | | | | | | 16.63 | 0.65 |
| Soybean | Falbala | linear | T[air]^{ref} | Linear | 2019.00 | 690.98 | 700.98 | 16.40 | -49.14 | 0.01 | | | | | | | 11.24 | 0.73 |
| Soybean | Falbala | linear | T[air]^{ref} | Linear | 2020.00 | 606.76 | 616.08 | | -47.04 | 0.02 | | | | | | | 14.69 | 0.57 |
| Soybean | Falbala | thermal | T[air]^{plot} | Thermal time | 2017.00 | 622.48 | 629.59 | | | | | | | | | | | |
| Soybean | Falbala | thermal | T[air]^{plot} | Thermal time | 2018.00 | 543.74 | 550.49 | | | | | | | | | | | |
| Soybean | Falbala | thermal | T[air]^{plot} | Thermal time | 2019.00 | 692.28 | 699.78 | 13.84 | | | | | | | | | | |
| Soybean | Falbala | thermal | T[air]^{plot} | Thermal time | 2020.00 | 593.60 | 600.59 | | | | | | | | | | | |
| Soybean | Falbala | thermal | T[air]^{ref} | Thermal time | 2017.00 | 622.48 | 629.59 | | | | | | | | | | | |
| Soybean | Falbala | thermal | T[air]^{ref} | Thermal time | 2018.00 | 543.74 | 550.49 | | | | | | | | | | | |
| Soybean | Falbala | thermal | T[air]^{ref} | Thermal time | 2019.00 | 692.28 | 699.78 | 13.84 | | | | | | | | | | |
| Soybean | Falbala | thermal | T[air]^{ref} | Thermal time | 2020.00 | 593.60 | 600.59 | | | | | | | | | | | |
| Soybean | Falbala | wang | T[air]^{plot} | Wang-Engel | 2017.00 | 624.21 | 633.69 | | | | | | | 26.92 | 1.60 | | 13.94 | 0.47 |
| Soybean | Falbala | wang | T[air]^{plot} | Wang-Engel | 2018.00 | 544.76 | 553.76 | | | | | | | 26.69 | 1.54 | | 13.29 | 0.44 |
| Soybean | Falbala | wang | T[air]^{plot} | Wang-Engel | 2019.00 | 686.00 | 695.99 | 14.44 | | | | | | 22.65 | 1.10 | | 10.91 | 0.65 |
| Soybean | Falbala | wang | T[air]^{plot} | Wang-Engel | 2020.00 | 601.92 | 611.25 | | | | | | | 26.09 | 1.49 | | 14.14 | 0.49 |
| Soybean | Falbala | wang | T[air]^{ref} | Wang-Engel | 2017.00 | 624.21 | 633.69 | | | | | | | 26.92 | 1.60 | | 13.94 | 0.47 |
| Soybean | Falbala | wang | T[air]^{ref} | Wang-Engel | 2018.00 | 544.76 | 553.76 | | | | | | | 26.69 | 1.54 | | 13.29 | 0.44 |
| Soybean | Falbala | wang | T[air]^{ref} | Wang-Engel | 2019.00 | 686.00 | 695.99 | 14.44 | | | | | | 22.65 | 1.10 | | 10.91 | 0.65 |
| Soybean | Falbala | wang | T[air]^{ref} | Wang-Engel | 2020.00 | 601.92 | 611.25 | | | | | | | 26.09 | 1.49 | | 14.14 | 0.49 |
| Soybean | Galice | asym | T[air]^{plot} | Asymptotic | 2017.00 | 542.11 | 553.28 | | 9.91 | | | 1.57 | 1.37 | | | | 15.46 | 0.51 |
| Soybean | Galice | asym | T[air]^{plot} | Asymptotic | 2018.00 | 515.11 | 526.35 | | 9.91 | | | 1.63 | 1.27 | | | | 11.78 | 0.56 |
| Soybean | Galice | asym | T[air]^{plot} | Asymptotic | 2019.00 | 621.30 | 633.08 | 15.98 | 9.96 | | | 2.56 | 1.00 | | | | 12.81 | 0.60 |
| Soybean | Galice | asym | T[air]^{plot} | Asymptotic | 2020.00 | 497.59 | 508.46 | | 0.00 | | | 1.00 | 1.32 | | | | 14.07 | 0.51 |
| Soybean | Galice | asym | T[air]^{ref} | Asymptotic | 2017.00 | 542.11 | 553.28 | | 9.91 | | | 1.57 | 1.37 | | | | 15.46 | 0.51 |
| Soybean | Galice | asym | T[air]^{ref} | Asymptotic | 2018.00 | 515.11 | 526.35 | | 9.91 | | | 1.63 | 1.27 | | | | 11.78 | 0.56 |
| Soybean | Galice | asym | T[air]^{ref} | Asymptotic | 2019.00 | 621.30 | 633.08 | 15.98 | 9.96 | | | 2.56 | 1.00 | | | | 12.81 | 0.60 |
| Soybean | Galice | asym | T[air]^{ref} | Asymptotic | 2020.00 | 497.59 | 508.46 | | 0.00 | | | 1.00 | 1.32 | | | | 14.07 | 0.51 |
| Soybean | Galice | bilinear | T[air]^{plot} | Bi-linear | 2017.00 | 545.45 | 556.62 | | 8.00 | 0.06 | 0.30 | | | | | | 15.87 | 0.55 |
| Soybean | Galice | bilinear | T[air]^{plot} | Bi-linear | 2018.00 | 521.91 | 533.16 | | 14.88 | 0.04 | 0.70 | | | | | | 12.42 | 0.61 |
| Soybean | Galice | bilinear | T[air]^{plot} | Bi-linear | 2019.00 | 623.79 | 635.58 | 17.16 | 19.95 | -0.02 | 1.05 | | | | | | 13.02 | 0.66 |

| | | | | | | | | | | | | | | | | | |
|---|---|---|---|---|---|---|---|---|---|---|---|---|---|---|---|---|---|
| Soybean | Galice | bilinear | T[air]^{plot} | Bi-linear | 2020.00 | 497.30 | 508.17 | | 12.69 | 0.02 | 0.95 | | | | | | 14.04 | 0.52 |
| Soybean | Galice | bilinear | T[air]^{ref} | Bi-linear | 2017.00 | 545.45 | 556.62 | | 8.00 | 0.06 | 0.30 | | | | | | 15.87 | 0.55 |
| Soybean | Galice | bilinear | T[air]^{ref} | Bi-linear | 2018.00 | 521.91 | 533.16 | | 14.88 | 0.04 | 0.70 | | | | | | 12.42 | 0.61 |
| Soybean | Galice | bilinear | T[air]^{ref} | Bi-linear | 2019.00 | 623.79 | 635.58 | 17.16 | 19.95 | -0.02 | 1.05 | | | | | | 13.02 | 0.66 |
| Soybean | Galice | bilinear | T[air]^{ref} | Bi-linear | 2020.00 | 497.30 | 508.17 | | 12.69 | 0.02 | 0.95 | | | | | | 14.04 | 0.52 |
| Soybean | Galice | gauss | ^{plot} | Gaussian | 2017.00 | 692.85 | 699.55 | 37.74 | | | | | | | | 70.11 | 37.77 | 0.03 |
| Soybean | Galice | gauss | ^{plot} | Gaussian | 2018.00 | 686.70 | 693.44 | 33.40 | | | | | | | | 72.52 | 33.57 | 0.01 |
| Soybean | Galice | gauss | ^{plot} | Gaussian | 2019.00 | 713.08 | 720.15 | 23.97 | | | | | | | | 60.93 | 23.86 | 0.04 |
| Soybean | Galice | gauss | ^{plot} | Gaussian | 2020.00 | 648.60 | 655.13 | 36.43 | | | | | | | | 74.18 | 36.65 | 0.02 |
| Soybean | Galice | gauss | ^{ref} | Gaussian | 2017.00 | 692.85 | 699.55 | 37.74 | | | | | | | | 70.11 | 37.77 | 0.03 |
| Soybean | Galice | gauss | ^{ref} | Gaussian | 2018.00 | 686.70 | 693.44 | 33.40 | | | | | | | | 72.52 | 33.57 | 0.01 |
| Soybean | Galice | gauss | ^{ref} | Gaussian | 2019.00 | 713.08 | 720.15 | 23.97 | | | | | | | | 60.93 | 23.86 | 0.04 |
| Soybean | Galice | gauss | ^{ref} | Gaussian | 2020.00 | 648.60 | 655.13 | 36.43 | | | | | | | | 74.18 | 36.65 | 0.02 |
| Soybean | Galice | linear | T[air]^{plot} | Linear | 2017.00 | 559.56 | 568.49 | | -44.39 | 0.02 | | | | | | | 18.14 | 0.68 |
| Soybean | Galice | linear | T[air]^{plot} | Linear | 2018.00 | 541.08 | 550.07 | | -46.73 | 0.01 | | | | | | | 14.73 | 0.71 |
| Soybean | Galice | linear | T[air]^{plot} | Linear | 2019.00 | 627.78 | 637.20 | 18.58 | -47.54 | 0.01 | | | | | | | 13.56 | 0.69 |
| Soybean | Galice | linear | T[air]^{plot} | Linear | 2020.00 | 495.63 | 504.33 | | -48.00 | 0.02 | | | | | | | 14.09 | 0.53 |
| Soybean | Galice | linear | T[air]^{ref} | Linear | 2017.00 | 559.56 | 568.49 | | -44.39 | 0.02 | | | | | | | 18.14 | 0.68 |
| Soybean | Galice | linear | T[air]^{ref} | Linear | 2018.00 | 541.08 | 550.07 | | -46.73 | 0.01 | | | | | | | 14.73 | 0.71 |
| Soybean | Galice | linear | T[air]^{ref} | Linear | 2019.00 | 627.78 | 637.20 | 18.58 | -47.54 | 0.01 | | | | | | | 13.56 | 0.69 |
| Soybean | Galice | linear | T[air]^{ref} | Linear | 2020.00 | 495.63 | 504.33 | | -48.00 | 0.02 | | | | | | | 14.09 | 0.53 |
| Soybean | Galice | thermal | T[air]^{plot} | Thermal time | 2017.00 | 542.04 | 548.74 | | | | | | | | | | | |
| Soybean | Galice | thermal | T[air]^{plot} | Thermal time | 2018.00 | 517.94 | 524.68 | | | | | | | | | | | |
| Soybean | Galice | thermal | T[air]^{plot} | Thermal time | 2019.00 | 627.36 | 634.43 | 16.38 | | | | | | | | | | |
| Soybean | Galice | thermal | T[air]^{plot} | Thermal time | 2020.00 | 504.78 | 511.30 | | | | | | | | | | | |
| Soybean | Galice | thermal | T[air]^{ref} | Thermal time | 2017.00 | 542.04 | 548.74 | | | | | | | | | | | |
| Soybean | Galice | thermal | T[air]^{ref} | Thermal time | 2018.00 | 517.94 | 524.68 | | | | | | | | | | | |
| Soybean | Galice | thermal | T[air]^{ref} | Thermal time | 2019.00 | 627.36 | 634.43 | 16.38 | | | | | | | | | | |
| Soybean | Galice | thermal | T[air]^{ref} | Thermal time | 2020.00 | 504.78 | 511.30 | | | | | | | | | | | |
| Soybean | Galice | wang | T[air]^{plot} | Wang-Engel | 2017.00 | 541.58 | 550.52 | | | | | | | 26.02 | 1.41 | | 15.64 | 0.50 |
| Soybean | Galice | wang | T[air]^{plot} | Wang-Engel | 2018.00 | 514.16 | 523.16 | | | | | | | 25.95 | 1.34 | | 11.88 | 0.58 |
| Soybean | Galice | wang | T[air]^{plot} | Wang-Engel | 2019.00 | 620.44 | 629.87 | 16.49 | | | | | | 23.00 | 1.09 | | 12.91 | 0.63 |
| Soybean | Galice | wang | T[air]^{plot} | Wang-Engel | 2020.00 | 498.65 | 507.35 | | | | | | | 23.55 | 1.33 | | 14.44 | 0.41 |
| Soybean | Galice | wang | T[air]^{ref} | Wang-Engel | 2017.00 | 541.58 | 550.52 | | | | | | | 26.02 | 1.41 | | 15.64 | 0.50 |
| Soybean | Galice | wang | T[air]^{ref} | Wang-Engel | 2018.00 | 514.16 | 523.16 | | | | | | | 25.95 | 1.34 | | 11.88 | 0.58 |
| Soybean | Galice | wang | T[air]^{ref} | Wang-Engel | 2019.00 | 620.44 | 629.87 | 16.49 | | | | | | 23.00 | 1.09 | | 12.91 | 0.63 |
| Soybean | Galice | wang | T[air]^{ref} | Wang-Engel | 2020.00 | 498.65 | 507.35 | | | | | | | 23.55 | 1.33 | | 14.44 | 0.41 |
| Soybean | Gallec | asym | T[air]^{plot} | Asymptotic | 2017.00 | 576.44 | 588.03 | | 6.60 | | | 0.00 | 2.88 | | | | 12.36 | 0.34 |
| Soybean | Gallec | asym | T[air]^{plot} | Asymptotic | 2018.00 | 539.08 | 550.32 | | 8.05 | | | 0.89 | 1.68 | | | | 12.55 | 0.31 |
| Soybean | Gallec | asym | T[air]^{plot} | Asymptotic | 2019.00 | 602.93 | 615.14 | 10.68 | 9.72 | | | 2.66 | 0.98 | | | | 8.25 | 0.65 |
| Soybean | Gallec | asym | T[air]^{plot} | Asymptotic | 2020.00 | 533.77 | 545.01 | | 0.00 | | | 0.03 | 2.14 | | | | 12.06 | 0.31 |
| Soybean | Gallec | asym | T[air]^{ref} | Asymptotic | 2017.00 | 576.44 | 588.03 | | 6.60 | | | 0.00 | 2.88 | | | | 12.36 | 0.34 |
| Soybean | Gallec | asym | T[air]^{ref} | Asymptotic | 2018.00 | 539.08 | 550.32 | | 8.05 | | | 0.89 | 1.68 | | | | 12.55 | 0.31 |
| Soybean | Gallec | asym | T[air]^{ref} | Asymptotic | 2019.00 | 602.93 | 615.14 | 10.68 | 9.72 | | | 2.66 | 0.98 | | | | 8.25 | 0.65 |
| Soybean | Gallec | asym | T[air]^{ref} | Asymptotic | 2020.00 | 533.77 | 545.01 | | 0.00 | | | 0.03 | 2.14 | | | | 12.06 | 0.31 |
| Soybean | Gallec | bilinear | T[air]^{plot} | Bi-linear | 2017.00 | 578.84 | 590.43 | | 17.85 | 0.07 | 0.83 | | | | | | 12.57 | 0.37 |
| Soybean | Gallec | bilinear | T[air]^{plot} | Bi-linear | 2018.00 | 540.93 | 552.18 | | 8.27 | 0.06 | 0.31 | | | | | | 12.78 | 0.31 |
| Soybean | Gallec | bilinear | T[air]^{plot} | Bi-linear | 2019.00 | 609.46 | 621.67 | 11.34 | 19.63 | -0.01 | 1.04 | | | | | | 8.58 | 0.67 |
| Soybean | Gallec | bilinear | T[air]^{plot} | Bi-linear | 2020.00 | 533.47 | 544.71 | | 12.52 | 0.03 | 0.79 | | | | | | 12.03 | 0.32 |
| Soybean | Gallec | bilinear | T[air]^{ref} | Bi-linear | 2017.00 | 578.84 | 590.43 | | 17.85 | 0.07 | 0.83 | | | | | | 12.57 | 0.37 |
| Soybean | Gallec | bilinear | T[air]^{ref} | Bi-linear | 2018.00 | 540.93 | 552.18 | | 8.27 | 0.06 | 0.31 | | | | | | 12.78 | 0.31 |
| Soybean | Gallec | bilinear | T[air]^{ref} | Bi-linear | 2019.00 | 609.46 | 621.67 | 11.34 | 19.63 | -0.01 | 1.04 | | | | | | 8.58 | 0.67 |

| | | | | | | | | | | | | | | | | | | |
|---|---|---|---|---|---|---|---|---|---|---|---|---|---|---|---|---|---|---|
| Soybean | Gallec | bilinear | T[air]^{ref} | Bi-linear | 2020.00 | 533.47 | 544.71 | | 12.52 | 0.03 | 0.79 | | | | | | 12.03 | 0.32 |
| Soybean | Gallec | gauss | ^{plot} | Gaussian | 2017.00 | 747.74 | 754.69 | 36.27 | | | | | | | | 68.93 | 36.34 | 0.05 |
| Soybean | Gallec | gauss | ^{plot} | Gaussian | 2018.00 | 691.66 | 698.41 | 34.62 | | | | | | | | 73.84 | 34.80 | 0.04 |
| Soybean | Gallec | gauss | ^{plot} | Gaussian | 2019.00 | 758.03 | 765.35 | 21.41 | | | | | | | | 60.74 | 21.27 | -0.07 |
| Soybean | Gallec | gauss | ^{plot} | Gaussian | 2020.00 | 694.58 | 701.32 | 35.40 | | | | | | | | 72.91 | 35.54 | 0.06 |
| Soybean | Gallec | gauss | ^{ref} | Gaussian | 2017.00 | 747.74 | 754.69 | 36.27 | | | | | | | | 68.93 | 36.34 | 0.05 |
| Soybean | Gallec | gauss | ^{ref} | Gaussian | 2018.00 | 691.66 | 698.41 | 34.62 | | | | | | | | 73.84 | 34.80 | 0.04 |
| Soybean | Gallec | gauss | ^{ref} | Gaussian | 2019.00 | 758.03 | 765.35 | 21.41 | | | | | | | | 60.74 | 21.27 | -0.07 |
| Soybean | Gallec | gauss | ^{ref} | Gaussian | 2020.00 | 694.58 | 701.32 | 35.40 | | | | | | | | 72.91 | 35.54 | 0.06 |
| Soybean | Gallec | linear | T[air]^{plot} | Linear | 2017.00 | 608.49 | 617.76 | | -46.00 | 0.01 | | | | | | | 15.80 | 0.55 |
| Soybean | Gallec | linear | T[air]^{plot} | Linear | 2018.00 | 566.60 | 575.60 | | -46.01 | 0.01 | | | | | | | 15.78 | 0.55 |
| Soybean | Gallec | linear | T[air]^{plot} | Linear | 2019.00 | 619.48 | 629.25 | 13.30 | -50.52 | 0.01 | | | | | | | 9.24 | 0.74 |
| Soybean | Gallec | linear | T[air]^{plot} | Linear | 2020.00 | 537.78 | 546.77 | | -48.55 | 0.02 | | | | | | | 12.65 | 0.32 |
| Soybean | Gallec | linear | T[air]^{ref} | Linear | 2017.00 | 608.49 | 617.76 | | -46.00 | 0.01 | | | | | | | 15.80 | 0.55 |
| Soybean | Gallec | linear | T[air]^{ref} | Linear | 2018.00 | 566.60 | 575.60 | | -46.01 | 0.01 | | | | | | | 15.78 | 0.55 |
| Soybean | Gallec | linear | T[air]^{ref} | Linear | 2019.00 | 619.48 | 629.25 | 13.30 | -50.52 | 0.01 | | | | | | | 9.24 | 0.74 |
| Soybean | Gallec | linear | T[air]^{ref} | Linear | 2020.00 | 537.78 | 546.77 | | -48.55 | 0.02 | | | | | | | 12.65 | 0.32 |
| Soybean | Gallec | thermal | T[air]^{plot} | Thermal time | 2017.00 | 576.55 | 583.51 | | | | | | | | | | | |
| Soybean | Gallec | thermal | T[air]^{plot} | Thermal time | 2018.00 | 538.25 | 545.00 | | | | | | | | | | | |
| Soybean | Gallec | thermal | T[air]^{plot} | Thermal time | 2019.00 | 618.48 | 625.81 | 10.63 | | | | | | | | | | |
| Soybean | Gallec | thermal | T[air]^{plot} | Thermal time | 2020.00 | 533.64 | 540.38 | | | | | | | | | | | |
| Soybean | Gallec | thermal | T[air]^{ref} | Thermal time | 2017.00 | 576.55 | 583.51 | | | | | | | | | | | |
| Soybean | Gallec | thermal | T[air]^{ref} | Thermal time | 2018.00 | 538.25 | 545.00 | | | | | | | | | | | |
| Soybean | Gallec | thermal | T[air]^{ref} | Thermal time | 2019.00 | 618.48 | 625.81 | 10.63 | | | | | | | | | | |
| Soybean | Gallec | thermal | T[air]^{ref} | Thermal time | 2020.00 | 533.64 | 540.38 | | | | | | | | | | | |
| Soybean | Gallec | wang | T[air]^{plot} | Wang-Engel | 2017.00 | 577.36 | 586.63 | | | | | | | 26.29 | 1.49 | | 12.62 | 0.31 |
| Soybean | Gallec | wang | T[air]^{plot} | Wang-Engel | 2018.00 | 539.89 | 548.88 | | | | | | | 26.10 | 1.44 | | 12.81 | 0.32 |
| Soybean | Gallec | wang | T[air]^{plot} | Wang-Engel | 2019.00 | 606.10 | 615.87 | 11.03 | | | | | | 22.95 | 1.11 | | 8.51 | 0.65 |
| Soybean | Gallec | wang | T[air]^{plot} | Wang-Engel | 2020.00 | 535.24 | 544.23 | | | | | | | 25.13 | 1.42 | | 12.38 | 0.20 |
| Soybean | Gallec | wang | T[air]^{ref} | Wang-Engel | 2017.00 | 577.36 | 586.63 | | | | | | | 26.29 | 1.49 | | 12.62 | 0.31 |
| Soybean | Gallec | wang | T[air]^{ref} | Wang-Engel | 2018.00 | 539.89 | 548.88 | | | | | | | 26.10 | 1.44 | | 12.81 | 0.32 |
| Soybean | Gallec | wang | T[air]^{ref} | Wang-Engel | 2019.00 | 606.10 | 615.87 | 11.03 | | | | | | 22.95 | 1.11 | | 8.51 | 0.65 |
| Soybean | Gallec | wang | T[air]^{ref} | Wang-Engel | 2020.00 | 535.24 | 544.23 | | | | | | | 25.13 | 1.42 | | 12.38 | 0.20 |
| Soybean | Idefix | asym | T[air]^{plot} | Asymptotic | 2017.00 | 505.08 | 516.10 | | 6.12 | | | 0.00 | 2.99 | | | | 11.58 | 0.47 |
| Soybean | Idefix | asym | T[air]^{plot} | Asymptotic | 2018.00 | 508.09 | 519.04 | | 8.72 | | | 1.33 | 1.47 | | | | 12.60 | 0.52 |
| Soybean | Idefix | asym | T[air]^{plot} | Asymptotic | 2019.00 | 569.36 | 581.21 | 12.06 | 9.91 | | | 3.01 | 1.02 | | | | 8.73 | 0.70 |
| Soybean | Idefix | asym | T[air]^{plot} | Asymptotic | 2020.00 | 518.33 | 529.35 | | 4.01 | | | 3.89 | 1.10 | | | | 12.84 | 0.51 |
| Soybean | Idefix | asym | T[air]^{ref} | Asymptotic | 2017.00 | 505.08 | 516.10 | | 6.12 | | | 0.00 | 2.99 | | | | 11.58 | 0.47 |
| Soybean | Idefix | asym | T[air]^{ref} | Asymptotic | 2018.00 | 508.09 | 519.04 | | 8.72 | | | 1.33 | 1.47 | | | | 12.60 | 0.52 |
| Soybean | Idefix | asym | T[air]^{ref} | Asymptotic | 2019.00 | 569.36 | 581.21 | 12.06 | 9.91 | | | 3.01 | 1.02 | | | | 8.73 | 0.70 |
| Soybean | Idefix | asym | T[air]^{ref} | Asymptotic | 2020.00 | 518.33 | 529.35 | | 4.01 | | | 3.89 | 1.10 | | | | 12.84 | 0.51 |
| Soybean | Idefix | bilinear | T[air]^{plot} | Bi-linear | 2017.00 | 505.74 | 516.77 | | 11.41 | 0.06 | 0.53 | | | | | | 11.64 | 0.49 |
| Soybean | Idefix | bilinear | T[air]^{plot} | Bi-linear | 2018.00 | 511.41 | 522.36 | | 13.69 | 0.05 | 0.72 | | | | | | 12.95 | 0.53 |
| Soybean | Idefix | bilinear | T[air]^{plot} | Bi-linear | 2019.00 | 576.84 | 588.68 | 12.76 | 17.87 | -0.01 | 1.10 | | | | | | 9.16 | 0.70 |
| Soybean | Idefix | bilinear | T[air]^{plot} | Bi-linear | 2020.00 | 502.74 | 513.76 | | 11.90 | 0.03 | 0.83 | | | | | | 11.36 | 0.56 |
| Soybean | Idefix | bilinear | T[air]^{ref} | Bi-linear | 2017.00 | 505.74 | 516.77 | | 11.41 | 0.06 | 0.53 | | | | | | 11.64 | 0.49 |
| Soybean | Idefix | bilinear | T[air]^{ref} | Bi-linear | 2018.00 | 511.41 | 522.36 | | 13.69 | 0.05 | 0.72 | | | | | | 12.95 | 0.53 |
| Soybean | Idefix | bilinear | T[air]^{ref} | Bi-linear | 2019.00 | 576.84 | 588.68 | 12.76 | 17.87 | -0.01 | 1.10 | | | | | | 9.16 | 0.70 |
| Soybean | Idefix | bilinear | T[air]^{ref} | Bi-linear | 2020.00 | 502.74 | 513.76 | | 11.90 | 0.03 | 0.83 | | | | | | 11.36 | 0.56 |
| Soybean | Idefix | gauss | ^{plot} | Gaussian | 2017.00 | 680.27 | 686.88 | 39.99 | | | | | | | | 73.92 | 40.01 | 0.02 |
| Soybean | Idefix | gauss | ^{plot} | Gaussian | 2018.00 | 663.44 | 670.01 | 38.01 | | | | | | | | 76.07 | 38.03 | 0.05 |
| Soybean | Idefix | gauss | ^{plot} | Gaussian | 2019.00 | 717.19 | 724.30 | 23.28 | | | | | | | | 62.85 | 23.10 | -0.13 |

| | | | | | | | | | | | | | | | | | | |
|---|---|---|---|---|---|---|---|---|---|---|---|---|---|---|---|---|---|---|
| Soybean | Idefix | gauss | ^{plot} | Gaussian | 2020.00 | 671.99 | 678.60 | 37.65 | | | | | | | | 74.03 | 37.58 | 0.09 |
| Soybean | Idefix | gauss | ^{ref} | Gaussian | 2017.00 | 680.27 | 686.88 | 39.99 | | | | | | | | 73.92 | 40.01 | 0.02 |
| Soybean | Idefix | gauss | ^{ref} | Gaussian | 2018.00 | 663.44 | 670.01 | 38.01 | | | | | | | | 76.07 | 38.03 | 0.05 |
| Soybean | Idefix | gauss | ^{ref} | Gaussian | 2019.00 | 717.19 | 724.30 | 23.28 | | | | | | | | 62.85 | 23.10 | -0.13 |
| Soybean | Idefix | gauss | ^{ref} | Gaussian | 2020.00 | 671.99 | 678.60 | 37.65 | | | | | | | | 74.03 | 37.58 | 0.09 |
| Soybean | Idefix | linear | T[air]^{plot} | Linear | 2017.00 | 537.77 | 546.59 | | -46.31 | 0.02 | | | | | | | 15.35 | 0.65 |
| Soybean | Idefix | linear | T[air]^{plot} | Linear | 2018.00 | 509.51 | 518.27 | | 0.19 | 0.05 | | | | | | | 12.95 | 0.52 |
| Soybean | Idefix | linear | T[air]^{plot} | Linear | 2019.00 | 586.69 | 596.17 | 14.21 | -50.06 | 0.01 | | | | | | | 9.90 | 0.72 |
| Soybean | Idefix | linear | T[air]^{plot} | Linear | 2020.00 | 504.83 | 513.65 | | -49.05 | 0.02 | | | | | | | 11.77 | 0.55 |
| Soybean | Idefix | linear | T[air]^{ref} | Linear | 2017.00 | 537.77 | 546.59 | | -46.31 | 0.02 | | | | | | | 15.35 | 0.65 |
| Soybean | Idefix | linear | T[air]^{ref} | Linear | 2018.00 | 509.51 | 518.27 | | 0.19 | 0.05 | | | | | | | 12.95 | 0.52 |
| Soybean | Idefix | linear | T[air]^{ref} | Linear | 2019.00 | 586.69 | 596.17 | 14.21 | -50.06 | 0.01 | | | | | | | 9.90 | 0.72 |
| Soybean | Idefix | linear | T[air]^{ref} | Linear | 2020.00 | 504.83 | 513.65 | | -49.05 | 0.02 | | | | | | | 11.77 | 0.55 |
| Soybean | Idefix | thermal | T[air]^{plot} | Thermal time | 2017.00 | 503.53 | 510.15 | | | | | | | | | | | |
| Soybean | Idefix | thermal | T[air]^{plot} | Thermal time | 2018.00 | 507.51 | 514.08 | | | | | | | | | | | |
| Soybean | Idefix | thermal | T[air]^{plot} | Thermal time | 2019.00 | 596.42 | 603.53 | 12.88 | | | | | | | | | | |
| Soybean | Idefix | thermal | T[air]^{plot} | Thermal time | 2020.00 | 506.82 | 513.44 | | | | | | | | | | | |
| Soybean | Idefix | thermal | T[air]^{ref} | Thermal time | 2017.00 | 503.53 | 510.15 | | | | | | | | | | | |
| Soybean | Idefix | thermal | T[air]^{ref} | Thermal time | 2018.00 | 507.51 | 514.08 | | | | | | | | | | | |
| Soybean | Idefix | thermal | T[air]^{ref} | Thermal time | 2019.00 | 596.42 | 603.53 | 12.88 | | | | | | | | | | |
| Soybean | Idefix | thermal | T[air]^{ref} | Thermal time | 2020.00 | 506.82 | 513.44 | | | | | | | | | | | |
| Soybean | Idefix | wang | T[air]^{plot} | Wang-Engel | 2017.00 | 507.86 | 516.68 | | | | | | | 26.25 | 1.58 | | 12.02 | 0.40 |
| Soybean | Idefix | wang | T[air]^{plot} | Wang-Engel | 2018.00 | 507.42 | 516.18 | | | | | | | 25.76 | 1.46 | | 12.74 | 0.50 |
| Soybean | Idefix | wang | T[air]^{plot} | Wang-Engel | 2019.00 | 572.34 | 581.82 | 11.83 | | | | | | 22.16 | 1.14 | | 9.01 | 0.66 |
| Soybean | Idefix | wang | T[air]^{plot} | Wang-Engel | 2020.00 | 512.18 | 521.00 | | | | | | | 24.89 | 1.42 | | 12.43 | 0.46 |
| Soybean | Idefix | wang | T[air]^{ref} | Wang-Engel | 2017.00 | 507.86 | 516.68 | | | | | | | 26.25 | 1.58 | | 12.02 | 0.40 |
| Soybean | Idefix | wang | T[air]^{ref} | Wang-Engel | 2018.00 | 507.42 | 516.18 | | | | | | | 25.76 | 1.46 | | 12.74 | 0.50 |
| Soybean | Idefix | wang | T[air]^{ref} | Wang-Engel | 2019.00 | 572.34 | 581.82 | 11.83 | | | | | | 22.16 | 1.14 | | 9.01 | 0.66 |
| Soybean | Idefix | wang | T[air]^{ref} | Wang-Engel | 2020.00 | 512.18 | 521.00 | | | | | | | 24.89 | 1.42 | | 12.43 | 0.46 |
| Soybean | Merlin | asym | T[air]^{plot} | Asymptotic | 2017.00 | 605.82 | 617.54 | | 8.45 | | | 0.89 | 1.68 | | | | 13.55 | 0.45 |
| Soybean | Merlin | asym | T[air]^{plot} | Asymptotic | 2018.00 | 551.52 | 562.76 | | 9.00 | | | 1.23 | 1.39 | | | | 13.77 | 0.47 |
| Soybean | Merlin | asym | T[air]^{plot} | Asymptotic | 2019.00 | 640.19 | 652.46 | 13.14 | 9.19 | | | 2.64 | 0.89 | | | | 9.86 | 0.67 |
| Soybean | Merlin | asym | T[air]^{plot} | Asymptotic | 2020.00 | 572.83 | 584.28 | | 0.00 | | | 0.35 | 1.66 | | | | 13.48 | 0.46 |
| Soybean | Merlin | asym | T[air]^{ref} | Asymptotic | 2017.00 | 605.82 | 617.54 | | 8.45 | | | 0.89 | 1.68 | | | | 13.55 | 0.45 |
| Soybean | Merlin | asym | T[air]^{ref} | Asymptotic | 2018.00 | 551.52 | 562.76 | | 9.00 | | | 1.23 | 1.39 | | | | 13.77 | 0.47 |
| Soybean | Merlin | asym | T[air]^{ref} | Asymptotic | 2019.00 | 640.19 | 652.46 | 13.14 | 9.19 | | | 2.64 | 0.89 | | | | 9.86 | 0.67 |
| Soybean | Merlin | asym | T[air]^{ref} | Asymptotic | 2020.00 | 572.83 | 584.28 | | 0.00 | | | 0.35 | 1.66 | | | | 13.48 | 0.46 |
| Soybean | Merlin | bilinear | T[air]^{plot} | Bi-linear | 2017.00 | 607.34 | 619.06 | | 8.69 | 0.06 | 0.30 | | | | | | 13.70 | 0.47 |
| Soybean | Merlin | bilinear | T[air]^{plot} | Bi-linear | 2018.00 | 554.31 | 565.56 | | 17.36 | 0.05 | 0.77 | | | | | | 14.05 | 0.51 |
| Soybean | Merlin | bilinear | T[air]^{plot} | Bi-linear | 2019.00 | 642.09 | 654.37 | 13.67 | 16.05 | -0.00 | 0.87 | | | | | | 9.97 | 0.69 |
| Soybean | Merlin | bilinear | T[air]^{plot} | Bi-linear | 2020.00 | 572.47 | 583.92 | | 12.51 | 0.03 | 0.78 | | | | | | 13.45 | 0.47 |
| Soybean | Merlin | bilinear | T[air]^{ref} | Bi-linear | 2017.00 | 607.34 | 619.06 | | 8.69 | 0.06 | 0.30 | | | | | | 13.70 | 0.47 |
| Soybean | Merlin | bilinear | T[air]^{ref} | Bi-linear | 2018.00 | 554.31 | 565.56 | | 17.36 | 0.05 | 0.77 | | | | | | 14.05 | 0.51 |
| Soybean | Merlin | bilinear | T[air]^{ref} | Bi-linear | 2019.00 | 642.09 | 654.37 | 13.67 | 16.05 | -0.00 | 0.87 | | | | | | 9.97 | 0.69 |
| Soybean | Merlin | bilinear | T[air]^{ref} | Bi-linear | 2020.00 | 572.47 | 583.92 | | 12.51 | 0.03 | 0.78 | | | | | | 13.45 | 0.47 |
| Soybean | Merlin | gauss | ^{plot} | Gaussian | 2017.00 | 765.79 | 772.83 | 36.01 | | | | | | | | 65.26 | 35.86 | 0.13 |
| Soybean | Merlin | gauss | ^{plot} | Gaussian | 2018.00 | 687.31 | 694.06 | 33.60 | | | | | | | | 71.15 | 33.72 | 0.07 |
| Soybean | Merlin | gauss | ^{plot} | Gaussian | 2019.00 | 761.65 | 769.01 | 20.69 | | | | | | | | 56.30 | 20.62 | -0.00 |
| Soybean | Merlin | gauss | ^{plot} | Gaussian | 2020.00 | 721.05 | 727.92 | 34.97 | | | | | | | | 68.43 | 34.70 | 0.17 |
| Soybean | Merlin | gauss | ^{ref} | Gaussian | 2017.00 | 765.79 | 772.83 | 36.01 | | | | | | | | 65.26 | 35.86 | 0.13 |
| Soybean | Merlin | gauss | ^{ref} | Gaussian | 2018.00 | 687.31 | 694.06 | 33.60 | | | | | | | | 71.15 | 33.72 | 0.07 |
| Soybean | Merlin | gauss | ^{ref} | Gaussian | 2019.00 | 761.65 | 769.01 | 20.69 | | | | | | | | 56.30 | 20.62 | -0.00 |

| | | | | | | | | | | | | | | | | | | |
|---|---|---|---|---|---|---|---|---|---|---|---|---|---|---|---|---|---|---|
| Soybean | Merlin | gauss | ^{ref} | Gaussian | 2020.00 | 721.05 | 727.92 | 34.97 | | | | | | | | 68.43 | 34.70 | 0.17 |
| Soybean | Merlin | linear | T[air]^{plot} | Linear | 2017.00 | 605.31 | 614.69 | | 3.56 | 0.06 | | | | | | | 13.69 | 0.47 |
| Soybean | Merlin | linear | T[air]^{plot} | Linear | 2018.00 | 552.01 | 561.01 | | 1.93 | 0.05 | | | | | | | 14.03 | 0.50 |
| Soybean | Merlin | linear | T[air]^{plot} | Linear | 2019.00 | 644.95 | 654.77 | 14.47 | -49.81 | 0.01 | | | | | | | 10.26 | 0.71 |
| Soybean | Merlin | linear | T[air]^{plot} | Linear | 2020.00 | 573.17 | 582.34 | | -47.56 | 0.01 | | | | | | | 13.74 | 0.47 |
| Soybean | Merlin | linear | T[air]^{ref} | Linear | 2017.00 | 605.31 | 614.69 | | 3.56 | 0.06 | | | | | | | 13.69 | 0.47 |
| Soybean | Merlin | linear | T[air]^{ref} | Linear | 2018.00 | 552.01 | 561.01 | | 1.93 | 0.05 | | | | | | | 14.03 | 0.50 |
| Soybean | Merlin | linear | T[air]^{ref} | Linear | 2019.00 | 644.95 | 654.77 | 14.47 | -49.81 | 0.01 | | | | | | | 10.26 | 0.71 |
| Soybean | Merlin | linear | T[air]^{ref} | Linear | 2020.00 | 573.17 | 582.34 | | -47.56 | 0.01 | | | | | | | 13.74 | 0.47 |
| Soybean | Merlin | thermal | T[air]^{plot} | Thermal time | 2017.00 | 604.95 | 611.98 | | | | | | | | | | | |
| Soybean | Merlin | thermal | T[air]^{plot} | Thermal time | 2018.00 | 550.39 | 557.14 | | | | | | | | | | | |
| Soybean | Merlin | thermal | T[air]^{plot} | Thermal time | 2019.00 | 652.88 | 660.24 | 12.94 | | | | | | | | | | |
| Soybean | Merlin | thermal | T[air]^{plot} | Thermal time | 2020.00 | 573.59 | 580.46 | | | | | | | | | | | |
| Soybean | Merlin | thermal | T[air]^{ref} | Thermal time | 2017.00 | 604.95 | 611.98 | | | | | | | | | | | |
| Soybean | Merlin | thermal | T[air]^{ref} | Thermal time | 2018.00 | 550.39 | 557.14 | | | | | | | | | | | |
| Soybean | Merlin | thermal | T[air]^{ref} | Thermal time | 2019.00 | 652.88 | 660.24 | 12.94 | | | | | | | | | | |
| Soybean | Merlin | thermal | T[air]^{ref} | Thermal time | 2020.00 | 573.59 | 580.46 | | | | | | | | | | | |
| Soybean | Merlin | wang | T[air]^{plot} | Wang-Engel | 2017.00 | 606.72 | 616.10 | | | | | | | 26.22 | 1.39 | | 13.83 | 0.43 |
| Soybean | Merlin | wang | T[air]^{plot} | Wang-Engel | 2018.00 | 549.16 | 558.15 | | | | | | | 26.14 | 1.35 | | 13.73 | 0.47 |
| Soybean | Merlin | wang | T[air]^{plot} | Wang-Engel | 2019.00 | 640.31 | 650.13 | 13.48 | | | | | | 21.88 | 0.97 | | 9.98 | 0.68 |
| Soybean | Merlin | wang | T[air]^{plot} | Wang-Engel | 2020.00 | 575.25 | 584.42 | | | | | | | 24.68 | 1.30 | | 13.91 | 0.40 |
| Soybean | Merlin | wang | T[air]^{ref} | Wang-Engel | 2017.00 | 606.72 | 616.10 | | | | | | | 26.22 | 1.39 | | 13.83 | 0.43 |
| Soybean | Merlin | wang | T[air]^{ref} | Wang-Engel | 2018.00 | 549.16 | 558.15 | | | | | | | 26.14 | 1.35 | | 13.73 | 0.47 |
| Soybean | Merlin | wang | T[air]^{ref} | Wang-Engel | 2019.00 | 640.31 | 650.13 | 13.48 | | | | | | 21.88 | 0.97 | | 9.98 | 0.68 |
| Soybean | Merlin | wang | T[air]^{ref} | Wang-Engel | 2020.00 | 575.25 | 584.42 | | | | | | | 24.68 | 1.30 | | 13.91 | 0.40 |
| Soybean | Obelix | asym | T[air]^{plot} | Asymptotic | 2017.00 | 418.66 | 429.13 | | 8.40 | 0.07 | 0.30 | | | | | | 11.43 | 0.41 |
| Soybean | Obelix | asym | T[air]^{plot} | Asymptotic | 2017.00 | 416.35 | 426.82 | | 8.49 | | | 0.94 | 1.86 | | | | 11.18 | 0.46 |
| Soybean | Obelix | asym | T[air]^{plot} | Asymptotic | 2018.00 | 479.15 | 490.24 | | 8.43 | | | 1.27 | 1.47 | | | | 11.32 | 0.54 |
| Soybean | Obelix | asym | T[air]^{plot} | Asymptotic | 2019.00 | 530.85 | 542.16 | 12.06 | 9.96 | | | 2.82 | 0.98 | | | | 10.00 | 0.57 |
| Soybean | Obelix | asym | T[air]^{plot} | Asymptotic | 2020.00 | 417.69 | 428.08 | | 22.23 | -0.01 | 1.36 | | | | | | 12.20 | 0.59 |
| Soybean | Obelix | asym | T[air]^{plot} | Asymptotic | 2020.00 | 428.73 | 439.12 | | 4.00 | | | 3.93 | 1.18 | | | | 13.56 | 0.63 |
| Soybean | Obelix | asym | T[air]^{ref} | Asymptotic | 2017.00 | 418.66 | 429.13 | | 8.40 | 0.07 | 0.30 | | | | | | 11.43 | 0.41 |
| Soybean | Obelix | asym | T[air]^{ref} | Asymptotic | 2017.00 | 416.35 | 426.82 | | 8.49 | | | 0.94 | 1.86 | | | | 11.18 | 0.46 |
| Soybean | Obelix | asym | T[air]^{ref} | Asymptotic | 2018.00 | 479.15 | 490.24 | | 8.43 | | | 1.27 | 1.47 | | | | 11.32 | 0.54 |
| Soybean | Obelix | asym | T[air]^{ref} | Asymptotic | 2019.00 | 530.85 | 542.16 | 12.06 | 9.96 | | | 2.82 | 0.98 | | | | 10.00 | 0.57 |
| Soybean | Obelix | asym | T[air]^{ref} | Asymptotic | 2020.00 | 417.69 | 428.08 | | 22.23 | -0.01 | 1.36 | | | | | | 12.20 | 0.59 |
| Soybean | Obelix | asym | T[air]^{ref} | Asymptotic | 2020.00 | 428.73 | 439.12 | | 4.00 | | | 3.93 | 1.18 | | | | 13.56 | 0.63 |
| Soybean | Obelix | bilinear | T[air]^{plot} | Bi-linear | 2018.00 | 482.26 | 493.36 | | 13.87 | 0.05 | 0.70 | | | | | | 11.61 | 0.53 |
| Soybean | Obelix | bilinear | T[air]^{plot} | Bi-linear | 2019.00 | 532.97 | 544.29 | 12.41 | 22.23 | -0.05 | 1.17 | | | | | | 10.14 | 0.59 |
| Soybean | Obelix | bilinear | T[air]^{ref} | Bi-linear | 2018.00 | 482.26 | 493.36 | | 13.87 | 0.05 | 0.70 | | | | | | 11.61 | 0.53 |
| Soybean | Obelix | bilinear | T[air]^{ref} | Bi-linear | 2019.00 | 532.97 | 544.29 | 12.41 | 22.23 | -0.05 | 1.17 | | | | | | 10.14 | 0.59 |
| Soybean | Obelix | gauss | ^{plot} | Gaussian | 2017.00 | 617.97 | 624.25 | 43.22 | | | | | | | | 75.60 | 43.26 | 0.09 |
| Soybean | Obelix | gauss | ^{plot} | Gaussian | 2018.00 | 685.33 | 691.98 | 38.49 | | | | | | | | 75.71 | 38.50 | 0.11 |
| Soybean | Obelix | gauss | ^{plot} | Gaussian | 2019.00 | 635.00 | 641.79 | 21.81 | | | | | | | | 61.60 | 21.63 | -0.12 |
| Soybean | Obelix | gauss | ^{plot} | Gaussian | 2020.00 | 601.87 | 608.10 | 41.10 | | | | | | | | 76.95 | 41.18 | 0.11 |
| Soybean | Obelix | gauss | ^{ref} | Gaussian | 2017.00 | 617.97 | 624.25 | 43.22 | | | | | | | | 75.60 | 43.26 | 0.09 |
| Soybean | Obelix | gauss | ^{ref} | Gaussian | 2018.00 | 685.33 | 691.98 | 38.49 | | | | | | | | 75.71 | 38.50 | 0.11 |
| Soybean | Obelix | gauss | ^{ref} | Gaussian | 2019.00 | 635.00 | 641.79 | 21.81 | | | | | | | | 61.60 | 21.63 | -0.12 |
| Soybean | Obelix | gauss | ^{ref} | Gaussian | 2020.00 | 601.87 | 608.10 | 41.10 | | | | | | | | 76.95 | 41.18 | 0.11 |
| Soybean | Obelix | linear | T[air]^{plot} | Linear | 2017.00 | 449.30 | 457.68 | | -46.31 | 0.02 | | | | | | | 15.74 | 0.72 |
| Soybean | Obelix | linear | T[air]^{plot} | Linear | 2018.00 | 503.93 | 512.81 | | -47.07 | 0.02 | | | | | | | 14.24 | 0.71 |
| Soybean | Obelix | linear | T[air]^{plot} | Linear | 2019.00 | 541.79 | 550.85 | 13.99 | -49.32 | 0.01 | | | | | | | 10.97 | 0.63 |

| | | | | | | | | | | | | | | | | | | |
|---|---|---|---|---|---|---|---|---|---|---|---|---|---|---|---|---|---|---|
| Soybean | Obelix | linear | T[air]^{plot} | Linear | 2020.00 | 417.75 | 426.07 | | -48.77 | 0.02 | | | | | | | 12.47 | 0.65 |
| Soybean | Obelix | linear | T[air]^{ref} | Linear | 2017.00 | 449.30 | 457.68 | | -46.31 | 0.02 | | | | | | | 15.74 | 0.72 |
| Soybean | Obelix | linear | T[air]^{ref} | Linear | 2018.00 | 503.93 | 512.81 | | -47.07 | 0.02 | | | | | | | 14.24 | 0.71 |
| Soybean | Obelix | linear | T[air]^{ref} | Linear | 2019.00 | 541.79 | 550.85 | 13.99 | -49.32 | 0.01 | | | | | | | 10.97 | 0.63 |
| Soybean | Obelix | linear | T[air]^{ref} | Linear | 2020.00 | 417.75 | 426.07 | | -48.77 | 0.02 | | | | | | | 12.47 | 0.65 |
| Soybean | Obelix | thermal | T[air]^{plot} | Thermal time | 2017.00 | 417.30 | 423.59 | | | | | | | | | | | |
| Soybean | Obelix | thermal | T[air]^{plot} | Thermal time | 2018.00 | 478.29 | 484.94 | | | | | | | | | | | |
| Soybean | Obelix | thermal | T[air]^{plot} | Thermal time | 2019.00 | 544.76 | 551.55 | 12.67 | | | | | | | | | | |
| Soybean | Obelix | thermal | T[air]^{plot} | Thermal time | 2020.00 | 421.97 | 428.21 | | | | | | | | | | | |
| Soybean | Obelix | thermal | T[air]^{ref} | Thermal time | 2017.00 | 417.30 | 423.59 | | | | | | | | | | | |
| Soybean | Obelix | thermal | T[air]^{ref} | Thermal time | 2018.00 | 478.29 | 484.94 | | | | | | | | | | | |
| Soybean | Obelix | thermal | T[air]^{ref} | Thermal time | 2019.00 | 544.76 | 551.55 | 12.67 | | | | | | | | | | |
| Soybean | Obelix | thermal | T[air]^{ref} | Thermal time | 2020.00 | 421.97 | 428.21 | | | | | | | | | | | |
| Soybean | Obelix | wang | T[air]^{plot} | Wang-Engel | 2017.00 | 418.46 | 426.84 | | | | | | | 26.21 | 1.60 | | 11.63 | 0.44 |
| Soybean | Obelix | wang | T[air]^{plot} | Wang-Engel | 2018.00 | 479.76 | 488.64 | | | | | | | 25.71 | 1.44 | | 11.56 | 0.51 |
| Soybean | Obelix | wang | T[air]^{plot} | Wang-Engel | 2019.00 | 529.04 | 538.09 | 12.03 | | | | | | 22.64 | 1.10 | | 10.00 | 0.57 |
| Soybean | Obelix | wang | T[air]^{plot} | Wang-Engel | 2020.00 | 419.12 | 427.43 | | | | | | | 24.63 | 1.46 | | 12.61 | 0.52 |
| Soybean | Obelix | wang | T[air]^{ref} | Wang-Engel | 2017.00 | 418.46 | 426.84 | | | | | | | 26.21 | 1.60 | | 11.63 | 0.44 |
| Soybean | Obelix | wang | T[air]^{ref} | Wang-Engel | 2018.00 | 479.76 | 488.64 | | | | | | | 25.71 | 1.44 | | 11.56 | 0.51 |
| Soybean | Obelix | wang | T[air]^{ref} | Wang-Engel | 2019.00 | 529.04 | 538.09 | 12.03 | | | | | | 22.64 | 1.10 | | 10.00 | 0.57 |
| Soybean | Obelix | wang | T[air]^{ref} | Wang-Engel | 2020.00 | 419.12 | 427.43 | | | | | | | 24.63 | 1.46 | | 12.61 | 0.52 |
| Soybean | Opaline | asym | T[air]^{plot} | Asymptotic | 2017.00 | 474.98 | 486.15 | | 7.67 | | | 0.00 | 3.41 | | | | 10.29 | 0.46 |
| Soybean | Opaline | asym | T[air]^{plot} | Asymptotic | 2018.00 | 458.93 | 469.80 | | 9.38 | 0.07 | 0.30 | | | | | | 11.60 | 0.38 |
| Soybean | Opaline | asym | T[air]^{plot} | Asymptotic | 2018.00 | 456.01 | 466.89 | | 9.19 | | | 0.95 | 1.91 | | | | 11.31 | 0.47 |
| Soybean | Opaline | asym | T[air]^{plot} | Asymptotic | 2019.00 | 562.92 | 574.83 | 10.21 | 9.38 | | | 2.22 | 1.10 | | | | 8.01 | 0.62 |
| Soybean | Opaline | asym | T[air]^{plot} | Asymptotic | 2020.00 | 490.04 | 501.14 | | 4.01 | | | 3.88 | 1.20 | | | | 12.38 | 0.57 |
| Soybean | Opaline | asym | T[air]^{ref} | Asymptotic | 2017.00 | 474.98 | 486.15 | | 7.67 | | | 0.00 | 3.41 | | | | 10.29 | 0.46 |
| Soybean | Opaline | asym | T[air]^{ref} | Asymptotic | 2018.00 | 458.93 | 469.80 | | 9.38 | 0.07 | 0.30 | | | | | | 11.60 | 0.38 |
| Soybean | Opaline | asym | T[air]^{ref} | Asymptotic | 2018.00 | 456.01 | 466.89 | | 9.19 | | | 0.95 | 1.91 | | | | 11.31 | 0.47 |
| Soybean | Opaline | asym | T[air]^{ref} | Asymptotic | 2019.00 | 562.92 | 574.83 | 10.21 | 9.38 | | | 2.22 | 1.10 | | | | 8.01 | 0.62 |
| Soybean | Opaline | asym | T[air]^{ref} | Asymptotic | 2020.00 | 490.04 | 501.14 | | 4.01 | | | 3.88 | 1.20 | | | | 12.38 | 0.57 |
| Soybean | Opaline | bilinear | T[air]^{plot} | Bi-linear | 2017.00 | 475.55 | 486.72 | | 11.45 | 0.08 | 0.44 | | | | | | 10.34 | 0.46 |
| Soybean | Opaline | bilinear | T[air]^{plot} | Bi-linear | 2019.00 | 572.92 | 584.83 | 10.83 | 9.43 | 0.04 | 0.59 | | | | | | 8.54 | 0.62 |
| Soybean | Opaline | bilinear | T[air]^{plot} | Bi-linear | 2020.00 | 452.33 | 463.43 | | 30.00 | -0.13 | 1.76 | | | | | | 9.09 | 0.52 |
| Soybean | Opaline | bilinear | T[air]^{ref} | Bi-linear | 2017.00 | 475.55 | 486.72 | | 11.45 | 0.08 | 0.44 | | | | | | 10.34 | 0.46 |
| Soybean | Opaline | bilinear | T[air]^{ref} | Bi-linear | 2019.00 | 572.92 | 584.83 | 10.83 | 9.43 | 0.04 | 0.59 | | | | | | 8.54 | 0.62 |
| Soybean | Opaline | bilinear | T[air]^{ref} | Bi-linear | 2020.00 | 452.33 | 463.43 | | 30.00 | -0.13 | 1.76 | | | | | | 9.09 | 0.52 |
| Soybean | Opaline | gauss | ^{plot} | Gaussian | 2017.00 | 707.11 | 713.82 | 42.13 | | | | | | | | 72.93 | 41.94 | 0.11 |
| Soybean | Opaline | gauss | ^{plot} | Gaussian | 2018.00 | 662.74 | 669.27 | 40.94 | | | | | | | | 78.12 | 40.93 | 0.11 |
| Soybean | Opaline | gauss | ^{plot} | Gaussian | 2019.00 | 722.43 | 729.57 | 22.93 | | | | | | | | 62.10 | 22.54 | -0.12 |
| Soybean | Opaline | gauss | ^{plot} | Gaussian | 2020.00 | 691.97 | 698.62 | 40.71 | | | | | | | | 74.83 | 40.46 | 0.15 |
| Soybean | Opaline | gauss | ^{ref} | Gaussian | 2017.00 | 707.11 | 713.82 | 42.13 | | | | | | | | 72.93 | 41.94 | 0.11 |
| Soybean | Opaline | gauss | ^{ref} | Gaussian | 2018.00 | 662.74 | 669.27 | 40.94 | | | | | | | | 78.12 | 40.93 | 0.11 |
| Soybean | Opaline | gauss | ^{ref} | Gaussian | 2019.00 | 722.43 | 729.57 | 22.93 | | | | | | | | 62.10 | 22.54 | -0.12 |
| Soybean | Opaline | gauss | ^{ref} | Gaussian | 2020.00 | 691.97 | 698.62 | 40.71 | | | | | | | | 74.83 | 40.46 | 0.15 |
| Soybean | Opaline | linear | T[air]^{plot} | Linear | 2017.00 | 524.02 | 532.95 | | -46.04 | 0.02 | | | | | | | 15.73 | 0.75 |
| Soybean | Opaline | linear | T[air]^{plot} | Linear | 2018.00 | 496.97 | 505.67 | | -45.43 | 0.02 | | | | | | | 16.61 | 0.74 |
| Soybean | Opaline | linear | T[air]^{plot} | Linear | 2019.00 | 584.08 | 593.61 | 13.30 | -50.47 | 0.01 | | | | | | | 9.31 | 0.72 |
| Soybean | Opaline | linear | T[air]^{plot} | Linear | 2020.00 | 447.93 | 456.80 | | -6.57 | 0.04 | | | | | | | 8.91 | 0.59 |
| Soybean | Opaline | linear | T[air]^{ref} | Linear | 2017.00 | 524.02 | 532.95 | | -46.04 | 0.02 | | | | | | | 15.73 | 0.75 |
| Soybean | Opaline | linear | T[air]^{ref} | Linear | 2018.00 | 496.97 | 505.67 | | -45.43 | 0.02 | | | | | | | 16.61 | 0.74 |
| Soybean | Opaline | linear | T[air]^{ref} | Linear | 2019.00 | 584.08 | 593.61 | 13.30 | -50.47 | 0.01 | | | | | | | 9.31 | 0.72 |

| | | | | | | | | | | | | | | | | | | |
|---|---|---|---|---|---|---|---|---|---|---|---|---|---|---|---|---|---|---|
| Soybean | Opaline | linear | T[air]^{ref} | Linear | 2020.00 | 447.93 | 456.80 | | -6.57 | 0.04 | | | | | | | 8.91 | 0.59 |
| Soybean | Opaline | thermal | T[air]^{plot} | Thermal time | 2017.00 | 479.42 | 486.12 | | | | | | | | | | | |
| Soybean | Opaline | thermal | T[air]^{plot} | Thermal time | 2018.00 | 459.94 | 466.47 | | | | | | | | | | | |
| Soybean | Opaline | thermal | T[air]^{plot} | Thermal time | 2019.00 | 572.25 | 579.40 | 10.14 | | | | | | | | | | |
| Soybean | Opaline | thermal | T[air]^{plot} | Thermal time | 2020.00 | 450.53 | 457.19 | | | | | | | | | | | |
| Soybean | Opaline | thermal | T[air]^{ref} | Thermal time | 2017.00 | 479.42 | 486.12 | | | | | | | | | | | |
| Soybean | Opaline | thermal | T[air]^{ref} | Thermal time | 2018.00 | 459.94 | 466.47 | | | | | | | | | | | |
| Soybean | Opaline | thermal | T[air]^{ref} | Thermal time | 2019.00 | 572.25 | 579.40 | 10.14 | | | | | | | | | | |
| Soybean | Opaline | thermal | T[air]^{ref} | Thermal time | 2020.00 | 450.53 | 457.19 | | | | | | | | | | | |
| Soybean | Opaline | wang | T[air]^{plot} | Wang-Engel | 2017.00 | 483.58 | 492.51 | | | | | | | 26.96 | 1.74 | | 11.20 | 0.39 |
| Soybean | Opaline | wang | T[air]^{plot} | Wang-Engel | 2018.00 | 457.50 | 466.20 | | | | | | | 26.82 | 1.67 | | 11.66 | 0.42 |
| Soybean | Opaline | wang | T[air]^{plot} | Wang-Engel | 2019.00 | 569.39 | 578.92 | 10.41 | | | | | | 23.75 | 1.20 | | 8.45 | 0.59 |
| Soybean | Opaline | wang | T[air]^{plot} | Wang-Engel | 2020.00 | 467.68 | 476.56 | | | | | | | 25.80 | 1.61 | | 10.47 | 0.35 |
| Soybean | Opaline | wang | T[air]^{ref} | Wang-Engel | 2017.00 | 483.58 | 492.51 | | | | | | | 26.96 | 1.74 | | 11.20 | 0.39 |
| Soybean | Opaline | wang | T[air]^{ref} | Wang-Engel | 2018.00 | 457.50 | 466.20 | | | | | | | 26.82 | 1.67 | | 11.66 | 0.42 |
| Soybean | Opaline | wang | T[air]^{ref} | Wang-Engel | 2019.00 | 569.39 | 578.92 | 10.41 | | | | | | 23.75 | 1.20 | | 8.45 | 0.59 |
| Soybean | Opaline | wang | T[air]^{ref} | Wang-Engel | 2020.00 | 467.68 | 476.56 | | | | | | | 25.80 | 1.61 | | 10.47 | 0.35 |
| Soybean | Protéix | asym | T[air]^{plot} | Asymptotic | 2017.00 | 563.38 | 575.10 | | 9.62 | | | 1.41 | 1.46 | | | | 12.60 | 0.60 |
| Soybean | Protéix | asym | T[air]^{plot} | Asymptotic | 2018.00 | 501.63 | 512.94 | | 9.62 | | | 1.40 | 1.43 | | | | 11.27 | 0.58 |
| Soybean | Protéix | asym | T[air]^{plot} | Asymptotic | 2019.00 | 643.28 | 655.55 | 13.49 | 9.91 | | | 2.59 | 0.97 | | | | 10.04 | 0.67 |
| Soybean | Protéix | asym | T[air]^{plot} | Asymptotic | 2020.00 | 498.99 | 510.38 | | 0.00 | | | 0.55 | 1.65 | | | | 10.41 | 0.62 |
| Soybean | Protéix | asym | T[air]^{ref} | Asymptotic | 2017.00 | 563.38 | 575.10 | | 9.62 | | | 1.41 | 1.46 | | | | 12.60 | 0.60 |
| Soybean | Protéix | asym | T[air]^{ref} | Asymptotic | 2018.00 | 501.63 | 512.94 | | 9.62 | | | 1.40 | 1.43 | | | | 11.27 | 0.58 |
| Soybean | Protéix | asym | T[air]^{ref} | Asymptotic | 2019.00 | 643.28 | 655.55 | 13.49 | 9.91 | | | 2.59 | 0.97 | | | | 10.04 | 0.67 |
| Soybean | Protéix | asym | T[air]^{ref} | Asymptotic | 2020.00 | 498.99 | 510.38 | | 0.00 | | | 0.55 | 1.65 | | | | 10.41 | 0.62 |
| Soybean | Protéix | bilinear | T[air]^{plot} | Bi-linear | 2017.00 | 567.70 | 579.42 | | 8.51 | 0.06 | 0.30 | | | | | | 13.00 | 0.60 |
| Soybean | Protéix | bilinear | T[air]^{plot} | Bi-linear | 2018.00 | 509.05 | 520.36 | | 29.90 | -0.03 | 1.46 | | | | | | 11.94 | 0.67 |
| Soybean | Protéix | bilinear | T[air]^{plot} | Bi-linear | 2019.00 | 650.61 | 662.88 | 14.54 | 17.34 | 0.00 | 0.93 | | | | | | 10.48 | 0.70 |
| Soybean | Protéix | bilinear | T[air]^{plot} | Bi-linear | 2020.00 | 504.09 | 515.47 | | 29.81 | -0.29 | 1.67 | | | | | | 10.82 | 0.55 |
| Soybean | Protéix | bilinear | T[air]^{ref} | Bi-linear | 2017.00 | 567.70 | 579.42 | | 8.51 | 0.06 | 0.30 | | | | | | 13.00 | 0.60 |
| Soybean | Protéix | bilinear | T[air]^{ref} | Bi-linear | 2018.00 | 509.05 | 520.36 | | 29.90 | -0.03 | 1.46 | | | | | | 11.94 | 0.67 |
| Soybean | Protéix | bilinear | T[air]^{ref} | Bi-linear | 2019.00 | 650.61 | 662.88 | 14.54 | 17.34 | 0.00 | 0.93 | | | | | | 10.48 | 0.70 |
| Soybean | Protéix | bilinear | T[air]^{ref} | Bi-linear | 2020.00 | 504.09 | 515.47 | | 29.81 | -0.29 | 1.67 | | | | | | 10.82 | 0.55 |
| Soybean | Protéix | gauss | ^{plot} | Gaussian | 2017.00 | 771.64 | 778.68 | 37.43 | | | | | | | | 66.60 | 37.27 | 0.14 |
| Soybean | Protéix | gauss | ^{plot} | Gaussian | 2018.00 | 704.51 | 711.30 | 35.82 | | | | | | | | 72.37 | 35.53 | 0.16 |
| Soybean | Protéix | gauss | ^{plot} | Gaussian | 2019.00 | 777.74 | 785.10 | 22.61 | | | | | | | | 58.58 | 22.66 | 0.08 |
| Soybean | Protéix | gauss | ^{plot} | Gaussian | 2020.00 | 719.25 | 726.08 | 36.86 | | | | | | | | 71.37 | 36.74 | 0.13 |
| Soybean | Protéix | gauss | ^{ref} | Gaussian | 2017.00 | 771.64 | 778.68 | 37.43 | | | | | | | | 66.60 | 37.27 | 0.14 |
| Soybean | Protéix | gauss | ^{ref} | Gaussian | 2018.00 | 704.51 | 711.30 | 35.82 | | | | | | | | 72.37 | 35.53 | 0.16 |
| Soybean | Protéix | gauss | ^{ref} | Gaussian | 2019.00 | 777.74 | 785.10 | 22.61 | | | | | | | | 58.58 | 22.66 | 0.08 |
| Soybean | Protéix | gauss | ^{ref} | Gaussian | 2020.00 | 719.25 | 726.08 | 36.86 | | | | | | | | 71.37 | 36.74 | 0.13 |
| Soybean | Protéix | linear | T[air]^{plot} | Linear | 2017.00 | 593.21 | 602.58 | | -45.88 | 0.01 | | | | | | | 15.96 | 0.73 |
| Soybean | Protéix | linear | T[air]^{plot} | Linear | 2018.00 | 535.39 | 544.44 | | -46.51 | 0.01 | | | | | | | 15.05 | 0.75 |
| Soybean | Protéix | linear | T[air]^{plot} | Linear | 2019.00 | 660.48 | 670.29 | 16.02 | -49.13 | 0.01 | | | | | | | 11.26 | 0.72 |
| Soybean | Protéix | linear | T[air]^{plot} | Linear | 2020.00 | 500.43 | 509.54 | | -49.63 | 0.02 | | | | | | | 10.72 | 0.63 |
| Soybean | Protéix | linear | T[air]^{ref} | Linear | 2017.00 | 593.21 | 602.58 | | -45.88 | 0.01 | | | | | | | 15.96 | 0.73 |
| Soybean | Protéix | linear | T[air]^{ref} | Linear | 2018.00 | 535.39 | 544.44 | | -46.51 | 0.01 | | | | | | | 15.05 | 0.75 |
| Soybean | Protéix | linear | T[air]^{ref} | Linear | 2019.00 | 660.48 | 670.29 | 16.02 | -49.13 | 0.01 | | | | | | | 11.26 | 0.72 |
| Soybean | Protéix | linear | T[air]^{ref} | Linear | 2020.00 | 500.43 | 509.54 | | -49.63 | 0.02 | | | | | | | 10.72 | 0.63 |
| Soybean | Protéix | thermal | T[air]^{plot} | Thermal time | 2017.00 | 565.47 | 572.50 | | | | | | | | | | | |
| Soybean | Protéix | thermal | T[air]^{plot} | Thermal time | 2018.00 | 505.31 | 512.10 | | | | | | | | | | | |
| Soybean | Protéix | thermal | T[air]^{plot} | Thermal time | 2019.00 | 656.04 | 663.40 | 13.49 | | | | | | | | | | |

| | | | | | | | | | | | | | | | | | | |
|---|---|---|---|---|---|---|---|---|---|---|---|---|---|---|---|---|---|---|
| Soybean | Protéix | thermal | T[air]^{plot} | Thermal time | 2020.00 | 506.67 | 513.50 | | | | | | | | | | | |
| Soybean | Protéix | thermal | T[air]^{ref} | Thermal time | 2017.00 | 565.47 | 572.50 | | | | | | | | | | | |
| Soybean | Protéix | thermal | T[air]^{ref} | Thermal time | 2018.00 | 505.31 | 512.10 | | | | | | | | | | | |
| Soybean | Protéix | thermal | T[air]^{ref} | Thermal time | 2019.00 | 656.04 | 663.40 | 13.49 | | | | | | | | | | |
| Soybean | Protéix | thermal | T[air]^{ref} | Thermal time | 2020.00 | 506.67 | 513.50 | | | | | | | | | | | |
| Soybean | Protéix | wang | T[air]^{plot} | Wang-Engel | 2017.00 | 568.10 | 577.47 | | | | | | | 26.17 | 1.46 | | 13.22 | 0.56 |
| Soybean | Protéix | wang | T[air]^{plot} | Wang-Engel | 2018.00 | 503.03 | 512.08 | | | | | | | 26.34 | 1.44 | | 11.57 | 0.56 |
| Soybean | Protéix | wang | T[air]^{plot} | Wang-Engel | 2019.00 | 648.52 | 658.33 | 14.09 | | | | | | 23.01 | 1.06 | | 10.47 | 0.67 |
| Soybean | Protéix | wang | T[air]^{plot} | Wang-Engel | 2020.00 | 507.81 | 516.92 | | | | | | | 24.35 | 1.40 | | 11.31 | 0.51 |
| Soybean | Protéix | wang | T[air]^{ref} | Wang-Engel | 2017.00 | 568.10 | 577.47 | | | | | | | 26.17 | 1.46 | | 13.22 | 0.56 |
| Soybean | Protéix | wang | T[air]^{ref} | Wang-Engel | 2018.00 | 503.03 | 512.08 | | | | | | | 26.34 | 1.44 | | 11.57 | 0.56 |
| Soybean | Protéix | wang | T[air]^{ref} | Wang-Engel | 2019.00 | 648.52 | 658.33 | 14.09 | | | | | | 23.01 | 1.06 | | 10.47 | 0.67 |
| Soybean | Protéix | wang | T[air]^{ref} | Wang-Engel | 2020.00 | 507.81 | 516.92 | | | | | | | 24.35 | 1.40 | | 11.31 | 0.51 |
| Soybean | Tequila | asym | T[air]^{plot} | Asymptotic | 2017.00 | 615.36 | 627.08 | 14.05 | 7.96 | | | 0.00 | 3.05 | | | | 12.98 | 0.39 |
| Soybean | Tequila | asym | T[air]^{plot} | Asymptotic | 2018.00 | 574.27 | 585.58 | 14.69 | 7.18 | | | 0.00 | 2.80 | | | | 13.62 | 0.39 |
| Soybean | Tequila | asym | T[air]^{plot} | Asymptotic | 2019.00 | 621.03 | 633.42 | 11.52 | 9.38 | | | 2.78 | 0.89 | | | | 8.12 | 0.71 |
| Soybean | Tequila | asym | T[air]^{plot} | Asymptotic | 2020.00 | 592.23 | 603.68 | 14.66 | 2.52 | | | 0.00 | 2.31 | | | | 13.80 | 0.35 |
| Soybean | Tequila | asym | T[air]^{ref} | Asymptotic | 2017.00 | 615.36 | 627.08 | 14.05 | 7.96 | | | 0.00 | 3.05 | | | | 12.98 | 0.39 |
| Soybean | Tequila | asym | T[air]^{ref} | Asymptotic | 2018.00 | 574.27 | 585.58 | 14.69 | 7.18 | | | 0.00 | 2.80 | | | | 13.62 | 0.39 |
| Soybean | Tequila | asym | T[air]^{ref} | Asymptotic | 2019.00 | 621.03 | 633.42 | 11.52 | 9.38 | | | 2.78 | 0.89 | | | | 8.12 | 0.71 |
| Soybean | Tequila | asym | T[air]^{ref} | Asymptotic | 2020.00 | 592.23 | 603.68 | 14.66 | 2.52 | | | 0.00 | 2.31 | | | | 13.80 | 0.35 |
| Soybean | Tequila | bilinear | T[air]^{plot} | Bi-linear | 2017.00 | 613.20 | 624.92 | 13.97 | 14.77 | 0.08 | 0.56 | | | | | | 12.81 | 0.41 |
| Soybean | Tequila | bilinear | T[air]^{plot} | Bi-linear | 2018.00 | 577.52 | 588.84 | 15.40 | 30.00 | 0.20 | 1.38 | | | | | | 13.94 | 0.44 |
| Soybean | Tequila | bilinear | T[air]^{plot} | Bi-linear | 2019.00 | 629.06 | 641.45 | 12.65 | 8.67 | 0.01 | 0.68 | | | | | | 8.49 | 0.74 |
| Soybean | Tequila | bilinear | T[air]^{plot} | Bi-linear | 2020.00 | 591.87 | 603.32 | 14.62 | 29.81 | -0.02 | 1.49 | | | | | | 13.76 | 0.35 |
| Soybean | Tequila | bilinear | T[air]^{ref} | Bi-linear | 2017.00 | 613.20 | 624.92 | 13.97 | 14.77 | 0.08 | 0.56 | | | | | | 12.81 | 0.41 |
| Soybean | Tequila | bilinear | T[air]^{ref} | Bi-linear | 2018.00 | 577.52 | 588.84 | 15.40 | 30.00 | 0.20 | 1.38 | | | | | | 13.94 | 0.44 |
| Soybean | Tequila | bilinear | T[air]^{ref} | Bi-linear | 2019.00 | 629.06 | 641.45 | 12.65 | 8.67 | 0.01 | 0.68 | | | | | | 8.49 | 0.74 |
| Soybean | Tequila | bilinear | T[air]^{ref} | Bi-linear | 2020.00 | 591.87 | 603.32 | 14.62 | 29.81 | -0.02 | 1.49 | | | | | | 13.76 | 0.35 |
| Soybean | Tequila | gauss | ^{plot} | Gaussian | 2017.00 | 769.49 | 776.52 | 36.83 | | | | | | | | 64.48 | 36.74 | 0.09 |
| Soybean | Tequila | gauss | ^{plot} | Gaussian | 2018.00 | 702.00 | 708.79 | 34.83 | | | | | | | | 71.04 | 34.90 | 0.07 |
| Soybean | Tequila | gauss | ^{plot} | Gaussian | 2019.00 | 775.13 | 782.56 | 20.01 | | | | | | | | 56.39 | 20.11 | -0.01 |
| Soybean | Tequila | gauss | ^{plot} | Gaussian | 2020.00 | 726.29 | 733.16 | 36.03 | | | | | | | | 67.94 | 35.98 | 0.11 |
| Soybean | Tequila | gauss | ^{ref} | Gaussian | 2017.00 | 769.49 | 776.52 | 36.83 | | | | | | | | 64.48 | 36.74 | 0.09 |
| Soybean | Tequila | gauss | ^{ref} | Gaussian | 2018.00 | 702.00 | 708.79 | 34.83 | | | | | | | | 71.04 | 34.90 | 0.07 |
| Soybean | Tequila | gauss | ^{ref} | Gaussian | 2019.00 | 775.13 | 782.56 | 20.01 | | | | | | | | 56.39 | 20.11 | -0.01 |
| Soybean | Tequila | gauss | ^{ref} | Gaussian | 2020.00 | 726.29 | 733.16 | 36.03 | | | | | | | | 67.94 | 35.98 | 0.11 |
| Soybean | Tequila | linear | T[air]^{plot} | Linear | 2017.00 | 652.51 | 661.88 | 20.27 | -45.20 | 0.01 | | | | | | | 16.96 | 0.57 |
| Soybean | Tequila | linear | T[air]^{plot} | Linear | 2018.00 | 603.32 | 612.37 | 20.45 | -45.06 | 0.01 | | | | | | | 17.16 | 0.57 |
| Soybean | Tequila | linear | T[air]^{plot} | Linear | 2019.00 | 627.21 | 637.12 | 12.91 | -51.02 | 0.01 | | | | | | | 8.50 | 0.76 |
| Soybean | Tequila | linear | T[air]^{plot} | Linear | 2020.00 | 598.08 | 607.24 | 15.56 | -46.80 | 0.02 | | | | | | | 14.64 | 0.37 |
| Soybean | Tequila | linear | T[air]^{ref} | Linear | 2017.00 | 652.51 | 661.88 | 20.27 | -45.20 | 0.01 | | | | | | | 16.96 | 0.57 |
| Soybean | Tequila | linear | T[air]^{ref} | Linear | 2018.00 | 603.32 | 612.37 | 20.45 | -45.06 | 0.01 | | | | | | | 17.16 | 0.57 |
| Soybean | Tequila | linear | T[air]^{ref} | Linear | 2019.00 | 627.21 | 637.12 | 12.91 | -51.02 | 0.01 | | | | | | | 8.50 | 0.76 |
| Soybean | Tequila | linear | T[air]^{ref} | Linear | 2020.00 | 598.08 | 607.24 | 15.56 | -46.80 | 0.02 | | | | | | | 14.64 | 0.37 |
| Soybean | Tequila | thermal | T[air]^{plot} | Thermal time | 2017.00 | 617.94 | 624.98 | 15.42 | | | | | | | | | | |
| Soybean | Tequila | thermal | T[air]^{plot} | Thermal time | 2018.00 | 574.24 | 581.03 | 15.62 | | | | | | | | | | |
| Soybean | Tequila | thermal | T[air]^{plot} | Thermal time | 2019.00 | 642.98 | 650.41 | 11.11 | | | | | | | | | | |
| Soybean | Tequila | thermal | T[air]^{plot} | Thermal time | 2020.00 | 588.07 | 594.94 | 14.66 | | | | | | | | | | |
| Soybean | Tequila | thermal | T[air]^{ref} | Thermal time | 2017.00 | 617.94 | 624.98 | 15.42 | | | | | | | | | | |
| Soybean | Tequila | thermal | T[air]^{ref} | Thermal time | 2018.00 | 574.24 | 581.03 | 15.62 | | | | | | | | | | |
| Soybean | Tequila | thermal | T[air]^{ref} | Thermal time | 2019.00 | 642.98 | 650.41 | 11.11 | | | | | | | | | | |

| | | | | | | | | | | | | | | | | | | |
|---|---|---|---|---|---|---|---|---|---|---|---|---|---|---|---|---|---|---|
| Soybean | Tequila | thermal | T[air]^{ref} | Thermal time | 2020.00 | 588.07 | 594.94 | 14.66 | | | | | | | | | | |
| Soybean | Tequila | wang | T[air]^{plot} | Wang-Engel | 2017.00 | 619.95 | 629.33 | 14.45 | | | | | | 27.32 | 1.59 | | 13.56 | 0.36 |
| Soybean | Tequila | wang | T[air]^{plot} | Wang-Engel | 2018.00 | 574.38 | 583.43 | 14.88 | | | | | | 26.86 | 1.48 | | 13.83 | 0.38 |
| Soybean | Tequila | wang | T[air]^{plot} | Wang-Engel | 2019.00 | 618.35 | 628.26 | 12.02 | | | | | | 21.52 | 0.97 | | 8.08 | 0.74 |
| Soybean | Tequila | wang | T[air]^{plot} | Wang-Engel | 2020.00 | 596.49 | 605.65 | 15.11 | | | | | | 26.04 | 1.43 | | 14.41 | 0.32 |
| Soybean | Tequila | wang | T[air]^{ref} | Wang-Engel | 2017.00 | 619.95 | 629.33 | 14.45 | | | | | | 27.32 | 1.59 | | 13.56 | 0.36 |
| Soybean | Tequila | wang | T[air]^{ref} | Wang-Engel | 2018.00 | 574.38 | 583.43 | 14.88 | | | | | | 26.86 | 1.48 | | 13.83 | 0.38 |
| Soybean | Tequila | wang | T[air]^{ref} | Wang-Engel | 2019.00 | 618.35 | 628.26 | 12.02 | | | | | | 21.52 | 0.97 | | 8.08 | 0.74 |
| Soybean | Tequila | wang | T[air]^{ref} | Wang-Engel | 2020.00 | 596.49 | 605.65 | 15.11 | | | | | | 26.04 | 1.43 | | 14.41 | 0.32 |
| Soybean | Toutatis | asym | T[air]^{plot} | Asymptotic | 2017.00 | 590.66 | 602.38 | | 9.22 | | | 1.05 | 1.69 | | | | 13.65 | 0.52 |
| Soybean | Toutatis | asym | T[air]^{plot} | Asymptotic | 2018.00 | 538.98 | 550.29 | | 9.62 | | | 1.31 | 1.45 | | | | 13.31 | 0.47 |
| Soybean | Toutatis | asym | T[air]^{plot} | Asymptotic | 2019.00 | 649.01 | 661.29 | 13.80 | 10.13 | | | 2.73 | 0.96 | | | | 10.38 | 0.66 |
| Soybean | Toutatis | asym | T[air]^{plot} | Asymptotic | 2020.00 | 543.89 | 555.27 | | 0.00 | | | 0.44 | 1.68 | | | | 13.00 | 0.51 |
| Soybean | Toutatis | asym | T[air]^{ref} | Asymptotic | 2017.00 | 590.66 | 602.38 | | 9.22 | | | 1.05 | 1.69 | | | | 13.65 | 0.52 |
| Soybean | Toutatis | asym | T[air]^{ref} | Asymptotic | 2018.00 | 538.98 | 550.29 | | 9.62 | | | 1.31 | 1.45 | | | | 13.31 | 0.47 |
| Soybean | Toutatis | asym | T[air]^{ref} | Asymptotic | 2019.00 | 649.01 | 661.29 | 13.80 | 10.13 | | | 2.73 | 0.96 | | | | 10.38 | 0.66 |
| Soybean | Toutatis | asym | T[air]^{ref} | Asymptotic | 2020.00 | 543.89 | 555.27 | | 0.00 | | | 0.44 | 1.68 | | | | 13.00 | 0.51 |
| Soybean | Toutatis | bilinear | T[air]^{plot} | Bi-linear | 2017.00 | 592.55 | 604.27 | | 9.13 | 0.06 | 0.30 | | | | | | 13.82 | 0.53 |
| Soybean | Toutatis | bilinear | T[air]^{plot} | Bi-linear | 2018.00 | 544.38 | 555.69 | | 30.00 | 0.00 | 1.42 | | | | | | 13.87 | 0.51 |
| Soybean | Toutatis | bilinear | T[air]^{plot} | Bi-linear | 2019.00 | 654.31 | 666.58 | 14.58 | 17.57 | -0.00 | 0.94 | | | | | | 10.71 | 0.68 |
| Soybean | Toutatis | bilinear | T[air]^{plot} | Bi-linear | 2020.00 | 545.91 | 557.30 | | 29.91 | -0.30 | 1.62 | | | | | | 13.20 | 0.48 |
| Soybean | Toutatis | bilinear | T[air]^{ref} | Bi-linear | 2017.00 | 592.55 | 604.27 | | 9.13 | 0.06 | 0.30 | | | | | | 13.82 | 0.53 |
| Soybean | Toutatis | bilinear | T[air]^{ref} | Bi-linear | 2018.00 | 544.38 | 555.69 | | 30.00 | 0.00 | 1.42 | | | | | | 13.87 | 0.51 |
| Soybean | Toutatis | bilinear | T[air]^{ref} | Bi-linear | 2019.00 | 654.31 | 666.58 | 14.58 | 17.57 | -0.00 | 0.94 | | | | | | 10.71 | 0.68 |
| Soybean | Toutatis | bilinear | T[air]^{ref} | Bi-linear | 2020.00 | 545.91 | 557.30 | | 29.91 | -0.30 | 1.62 | | | | | | 13.20 | 0.48 |
| Soybean | Toutatis | gauss | ^{plot} | Gaussian | 2017.00 | 770.66 | 777.69 | 37.09 | | | | | | | | 66.85 | 37.03 | 0.11 |
| Soybean | Toutatis | gauss | ^{plot} | Gaussian | 2018.00 | 700.48 | 707.26 | 34.52 | | | | | | | | 72.32 | 34.52 | 0.09 |
| Soybean | Toutatis | gauss | ^{plot} | Gaussian | 2019.00 | 770.44 | 777.80 | 21.61 | | | | | | | | 58.54 | 21.71 | 0.04 |
| Soybean | Toutatis | gauss | ^{plot} | Gaussian | 2020.00 | 716.89 | 723.72 | 36.21 | | | | | | | | 71.22 | 36.14 | 0.12 |
| Soybean | Toutatis | gauss | ^{ref} | Gaussian | 2017.00 | 770.66 | 777.69 | 37.09 | | | | | | | | 66.85 | 37.03 | 0.11 |
| Soybean | Toutatis | gauss | ^{ref} | Gaussian | 2018.00 | 700.48 | 707.26 | 34.52 | | | | | | | | 72.32 | 34.52 | 0.09 |
| Soybean | Toutatis | gauss | ^{ref} | Gaussian | 2019.00 | 770.44 | 777.80 | 21.61 | | | | | | | | 58.54 | 21.71 | 0.04 |
| Soybean | Toutatis | gauss | ^{ref} | Gaussian | 2020.00 | 716.89 | 723.72 | 36.21 | | | | | | | | 71.22 | 36.14 | 0.12 |
| Soybean | Toutatis | linear | T[air]^{plot} | Linear | 2017.00 | 619.06 | 628.44 | | -45.17 | 0.01 | | | | | | | 17.00 | 0.63 |
| Soybean | Toutatis | linear | T[air]^{plot} | Linear | 2018.00 | 565.73 | 574.78 | | -45.38 | 0.01 | | | | | | | 16.70 | 0.61 |
| Soybean | Toutatis | linear | T[air]^{plot} | Linear | 2019.00 | 661.24 | 671.06 | 15.73 | -49.09 | 0.01 | | | | | | | 11.31 | 0.70 |
| Soybean | Toutatis | linear | T[air]^{plot} | Linear | 2020.00 | 543.92 | 553.02 | | -48.22 | 0.02 | | | | | | | 13.23 | 0.52 |
| Soybean | Toutatis | linear | T[air]^{ref} | Linear | 2017.00 | 619.06 | 628.44 | | -45.17 | 0.01 | | | | | | | 17.00 | 0.63 |
| Soybean | Toutatis | linear | T[air]^{ref} | Linear | 2018.00 | 565.73 | 574.78 | | -45.38 | 0.01 | | | | | | | 16.70 | 0.61 |
| Soybean | Toutatis | linear | T[air]^{ref} | Linear | 2019.00 | 661.24 | 671.06 | 15.73 | -49.09 | 0.01 | | | | | | | 11.31 | 0.70 |
| Soybean | Toutatis | linear | T[air]^{ref} | Linear | 2020.00 | 543.92 | 553.02 | | -48.22 | 0.02 | | | | | | | 13.23 | 0.52 |
| Soybean | Toutatis | thermal | T[air]^{plot} | Thermal time | 2017.00 | 591.58 | 598.62 | | | | | | | | | | | |
| Soybean | Toutatis | thermal | T[air]^{plot} | Thermal time | 2018.00 | 540.44 | 547.23 | | | | | | | | | | | |
| Soybean | Toutatis | thermal | T[air]^{plot} | Thermal time | 2019.00 | 661.74 | 669.10 | 14.10 | | | | | | | | | | |
| Soybean | Toutatis | thermal | T[air]^{plot} | Thermal time | 2020.00 | 546.23 | 553.06 | | | | | | | | | | | |
| Soybean | Toutatis | thermal | T[air]^{ref} | Thermal time | 2017.00 | 591.58 | 598.62 | | | | | | | | | | | |
| Soybean | Toutatis | thermal | T[air]^{ref} | Thermal time | 2018.00 | 540.44 | 547.23 | | | | | | | | | | | |
| Soybean | Toutatis | thermal | T[air]^{ref} | Thermal time | 2019.00 | 661.74 | 669.10 | 14.10 | | | | | | | | | | |
| Soybean | Toutatis | thermal | T[air]^{ref} | Thermal time | 2020.00 | 546.23 | 553.06 | | | | | | | | | | | |
| Soybean | Toutatis | wang | T[air]^{plot} | Wang-Engel | 2017.00 | 592.09 | 601.47 | | | | | | | 26.48 | 1.49 | | 13.97 | 0.50 |
| Soybean | Toutatis | wang | T[air]^{plot} | Wang-Engel | 2018.00 | 537.85 | 546.90 | | | | | | | 26.46 | 1.44 | | 13.40 | 0.48 |
| Soybean | Toutatis | wang | T[air]^{plot} | Wang-Engel | 2019.00 | 651.83 | 661.64 | 14.20 | | | | | | 22.91 | 1.05 | | 10.68 | 0.66 |

| | | | | | | | | | | | | | | | | | | |
|---|---|---|---|---|---|---|---|---|---|---|---|---|---|---|---|---|---|---|
| Soybean | Toutatis | wang | T[air]^{plot} | Wang-Engel | 2020.00 | 549.34 | 558.45 | | | | | | | 24.56 | 1.36 | | 13.75 | 0.44 |
| Soybean | Toutatis | wang | T[air]^{ref} | Wang-Engel | 2017.00 | 592.09 | 601.47 | | | | | | | 26.48 | 1.49 | | 13.97 | 0.50 |
| Soybean | Toutatis | wang | T[air]^{ref} | Wang-Engel | 2018.00 | 537.85 | 546.90 | | | | | | | 26.46 | 1.44 | | 13.40 | 0.48 |
| Soybean | Toutatis | wang | T[air]^{ref} | Wang-Engel | 2019.00 | 651.83 | 661.64 | 14.20 | | | | | | 22.91 | 1.05 | | 10.68 | 0.66 |
| Soybean | Toutatis | wang | T[air]^{ref} | Wang-Engel | 2020.00 | 549.34 | 558.45 | | | | | | | 24.56 | 1.36 | | 13.75 | 0.44 |
| Wheat | ALL | asym | T[air]^{plot} | Asymptotic | 2015.00 | 9930.90 | 9956.75 | | 4.30 | | | 2.28 | 0.68 | | | | 12.76 | 0.54 |
| Wheat | ALL | asym | T[air]^{plot} | Asymptotic | 2016.00 | 9484.15 | 9509.76 | | 5.60 | | | 4.09 | 0.58 | | | | 12.96 | 0.54 |
| Wheat | ALL | asym | T[air]^{plot} | Asymptotic | 2017.00 | 9662.51 | 9688.28 | 12.50 | 5.54 | | | 5.68 | 0.57 | | | | 10.57 | 0.54 |
| Wheat | ALL | asym | T[air]^{plot} | Asymptotic | 2018.00 | 9651.99 | 9677.61 | | 6.12 | | | 9.84 | 0.60 | | | | 13.73 | 0.51 |
| Wheat | ALL | asym | T[air]^{plot} | Asymptotic | 2019.00 | 8819.94 | 8845.14 | | 3.30 | | | 1.35 | 0.85 | | | | 13.67 | 0.52 |
| Wheat | ALL | asym | T[air]^{plot} | Asymptotic | 2021.00 | 9739.97 | 9765.63 | | 5.70 | | | 6.32 | 0.58 | | | | 13.76 | 0.52 |
| Wheat | ALL | asym | T[air]^{ref} | Asymptotic | 2015.00 | 10267.37 | 10293.21 | | 3.30 | | | 1.91 | 0.72 | | | | 12.70 | 0.53 |
| Wheat | ALL | asym | T[air]^{ref} | Asymptotic | 2016.00 | 9867.19 | 9892.80 | | 5.24 | | | 8.27 | 0.56 | | | | 13.13 | 0.55 |
| Wheat | ALL | asym | T[air]^{ref} | Asymptotic | 2017.00 | 9650.23 | 9676.00 | | 5.30 | | | 6.05 | 0.57 | | | | 10.62 | 0.52 |
| Wheat | ALL | asym | T[air]^{ref} | Asymptotic | 2018.00 | 10031.97 | 10057.59 | 16.05 | 5.70 | | | 10.15 | 0.59 | | | | 13.72 | 0.52 |
| Wheat | ALL | asym | T[air]^{ref} | Asymptotic | 2019.00 | 9237.88 | 9263.08 | | 6.80 | | | 4.86 | 0.62 | | | | 14.04 | 0.52 |
| Wheat | ALL | asym | T[air]^{ref} | Asymptotic | 2021.00 | 10058.32 | 10083.98 | | 2.84 | | | 1.65 | 0.73 | | | | 13.56 | 0.51 |
| Wheat | ALL | bilinear | T[air]^{plot} | Bi-linear | 2015.00 | 9929.92 | 9955.76 | | 10.21 | 0.02 | 0.55 | | | | | | 12.76 | 0.57 |
| Wheat | ALL | bilinear | T[air]^{plot} | Bi-linear | 2016.00 | 9490.10 | 9515.71 | | 9.96 | 0.02 | 0.51 | | | | | | 12.99 | 0.56 |
| Wheat | ALL | bilinear | T[air]^{plot} | Bi-linear | 2017.00 | 9619.08 | 9644.85 | 12.17 | 7.84 | 0.01 | 0.50 | | | | | | 10.39 | 0.52 |
| Wheat | ALL | bilinear | T[air]^{plot} | Bi-linear | 2018.00 | 9657.31 | 9682.93 | | 9.16 | 0.02 | 0.50 | | | | | | 13.75 | 0.53 |
| Wheat | ALL | bilinear | T[air]^{plot} | Bi-linear | 2019.00 | 8839.70 | 8864.90 | | 9.10 | 0.03 | 0.46 | | | | | | 13.80 | 0.53 |
| Wheat | ALL | bilinear | T[air]^{plot} | Bi-linear | 2021.00 | 9702.40 | 9728.06 | | 10.28 | 0.02 | 0.52 | | | | | | 13.52 | 0.55 |
| Wheat | ALL | bilinear | T[air]^{ref} | Bi-linear | 2015.00 | 10284.04 | 10309.89 | | 12.26 | 0.01 | 0.65 | | | | | | 12.78 | 0.54 |
| Wheat | ALL | bilinear | T[air]^{ref} | Bi-linear | 2016.00 | 9856.10 | 9881.71 | | 10.73 | 0.01 | 0.56 | | | | | | 13.08 | 0.54 |
| Wheat | ALL | bilinear | T[air]^{ref} | Bi-linear | 2017.00 | 9610.38 | 9636.15 | | 7.70 | 0.01 | 0.50 | | | | | | 10.44 | 0.52 |
| Wheat | ALL | bilinear | T[air]^{ref} | Bi-linear | 2018.00 | 10040.61 | 10066.23 | 16.01 | 11.21 | 0.01 | 0.60 | | | | | | 13.78 | 0.51 |
| Wheat | ALL | bilinear | T[air]^{ref} | Bi-linear | 2019.00 | 9188.95 | 9214.15 | | 12.80 | 0.02 | 0.62 | | | | | | 13.75 | 0.50 |
| Wheat | ALL | bilinear | T[air]^{ref} | Bi-linear | 2021.00 | 10066.30 | 10091.96 | | 11.10 | 0.01 | 0.57 | | | | | | 13.60 | 0.52 |
| Wheat | ALL | gauss | ^{plot} | Gaussian | 2015.00 | 12239.83 | 12255.34 | 28.91 | | | | | | | | 44.27 | 26.94 | 0.36 |
| Wheat | ALL | gauss | ^{plot} | Gaussian | 2016.00 | 11647.19 | 11662.56 | 27.98 | | | | | | | | 45.21 | 26.75 | 0.30 |
| Wheat | ALL | gauss | ^{plot} | Gaussian | 2017.00 | 11996.25 | 12011.71 | 27.41 | | | | | | | | 45.68 | 26.37 | 0.28 |
| Wheat | ALL | gauss | ^{plot} | Gaussian | 2018.00 | 11901.97 | 11917.34 | 30.87 | | | | | | | | 48.00 | 29.19 | 0.33 |
| Wheat | ALL | gauss | ^{plot} | Gaussian | 2019.00 | 10781.22 | 10796.34 | 29.01 | | | | | | | | 47.74 | 27.42 | 0.33 |
| Wheat | ALL | gauss | ^{plot} | Gaussian | 2021.00 | 11971.80 | 11987.20 | 30.44 | | | | | | | | 48.03 | 28.89 | 0.32 |
| Wheat | ALL | gauss | ^{ref} | Gaussian | 2015.00 | 12239.83 | 12255.34 | 28.91 | | | | | | | | 44.27 | 26.94 | 0.36 |
| Wheat | ALL | gauss | ^{ref} | Gaussian | 2016.00 | 11647.19 | 11662.56 | 27.98 | | | | | | | | 45.21 | 26.75 | 0.30 |
| Wheat | ALL | gauss | ^{ref} | Gaussian | 2017.00 | 11996.25 | 12011.71 | 27.41 | | | | | | | | 45.68 | 26.37 | 0.28 |
| Wheat | ALL | gauss | ^{ref} | Gaussian | 2018.00 | 11901.97 | 11917.34 | 30.87 | | | | | | | | 48.00 | 29.19 | 0.33 |
| Wheat | ALL | gauss | ^{ref} | Gaussian | 2019.00 | 10781.22 | 10796.34 | 29.01 | | | | | | | | 47.74 | 27.42 | 0.33 |
| Wheat | ALL | gauss | ^{ref} | Gaussian | 2021.00 | 11971.80 | 11987.20 | 30.44 | | | | | | | | 48.03 | 28.89 | 0.32 |
| Wheat | ALL | linear | T[air]^{plot} | Linear | 2015.00 | 10283.97 | 10304.64 | | -46.68 | 0.01 | | | | | | | 14.81 | 0.59 |
| Wheat | ALL | linear | T[air]^{plot} | Linear | 2016.00 | 9768.30 | 9788.78 | | -46.76 | 0.01 | | | | | | | 14.70 | 0.60 |
| Wheat | ALL | linear | T[air]^{plot} | Linear | 2017.00 | 9684.95 | 9705.56 | 12.84 | -49.51 | 0.01 | | | | | | | 10.69 | 0.56 |
| Wheat | ALL | linear | T[air]^{plot} | Linear | 2018.00 | 9961.79 | 9982.29 | | -46.04 | 0.01 | | | | | | | 15.74 | 0.57 |
| Wheat | ALL | linear | T[air]^{plot} | Linear | 2019.00 | 9171.79 | 9191.95 | | -45.73 | 0.01 | | | | | | | 16.19 | 0.56 |
| Wheat | ALL | linear | T[air]^{plot} | Linear | 2021.00 | 10012.74 | 10033.27 | | -46.18 | 0.01 | | | | | | | 15.53 | 0.57 |
| Wheat | ALL | linear | T[air]^{ref} | Linear | 2015.00 | 10650.31 | 10670.99 | | -46.66 | 0.01 | | | | | | | 14.83 | 0.58 |
| Wheat | ALL | linear | T[air]^{ref} | Linear | 2016.00 | 10140.44 | 10160.93 | | -46.71 | 0.01 | | | | | | | 14.76 | 0.58 |
| Wheat | ALL | linear | T[air]^{ref} | Linear | 2017.00 | 9664.89 | 9685.51 | | -49.51 | 0.01 | | | | | | | 10.71 | 0.56 |
| Wheat | ALL | linear | T[air]^{ref} | Linear | 2018.00 | 10353.76 | 10374.26 | 18.90 | -46.04 | 0.01 | | | | | | | 15.74 | 0.56 |
| Wheat | ALL | linear | T[air]^{ref} | Linear | 2019.00 | 9538.10 | 9558.25 | | -45.75 | 0.01 | | | | | | | 16.16 | 0.55 |

| | | | | | | | | | | | | | | | | | | |
|---|---|---|---|---|---|---|---|---|---|---|---|---|---|---|---|---|---|---|
| Wheat | ALL | linear | T[air]^{ref} | Linear | 2021.00 | 10380.91 | 10401.44 | | -46.17 | 0.01 | | | | | | | 15.54 | 0.57 |
| Wheat | ALL | thermal | T[air]^{plot} | Thermal time | 2015.00 | 10030.19 | 10045.70 | | | | | | | | | | | |
| Wheat | ALL | thermal | T[air]^{plot} | Thermal time | 2016.00 | 9604.03 | 9619.40 | | | | | | | | | | | |
| Wheat | ALL | thermal | T[air]^{plot} | Thermal time | 2017.00 | 9930.44 | 9945.91 | 13.59 | | | | | | | | | | |
| Wheat | ALL | thermal | T[air]^{plot} | Thermal time | 2018.00 | 9743.47 | 9758.84 | | | | | | | | | | | |
| Wheat | ALL | thermal | T[air]^{plot} | Thermal time | 2019.00 | 8871.13 | 8886.25 | | | | | | | | | | | |
| Wheat | ALL | thermal | T[air]^{plot} | Thermal time | 2021.00 | 9782.61 | 9798.01 | | | | | | | | | | | |
| Wheat | ALL | thermal | T[air]^{ref} | Thermal time | 2015.00 | 10448.21 | 10463.71 | | | | | | | | | | | |
| Wheat | ALL | thermal | T[air]^{ref} | Thermal time | 2016.00 | 10019.99 | 10035.36 | | | | | | | | | | | |
| Wheat | ALL | thermal | T[air]^{ref} | Thermal time | 2017.00 | 9956.52 | 9971.98 | | | | | | | | | | | |
| Wheat | ALL | thermal | T[air]^{ref} | Thermal time | 2018.00 | 10176.97 | 10192.35 | 16.49 | | | | | | | | | | |
| Wheat | ALL | thermal | T[air]^{ref} | Thermal time | 2019.00 | 9243.84 | 9258.95 | | | | | | | | | | | |
| Wheat | ALL | thermal | T[air]^{ref} | Thermal time | 2021.00 | 10190.36 | 10205.76 | | | | | | | | | | | |
| Wheat | ALL | wang | T[air]^{plot} | Wang-Engel | 2015.00 | 10106.91 | 10127.59 | | | | | | | 16.79 | 0.81 | | 13.70 | 0.50 |
| Wheat | ALL | wang | T[air]^{plot} | Wang-Engel | 2016.00 | 9696.76 | 9717.25 | | | | | | | 16.82 | 0.74 | | 14.18 | 0.53 |
| Wheat | ALL | wang | T[air]^{plot} | Wang-Engel | 2017.00 | 9932.34 | 9952.96 | 13.56 | | | | | | 15.36 | 0.68 | | 11.75 | 0.50 |
| Wheat | ALL | wang | T[air]^{plot} | Wang-Engel | 2018.00 | 9809.89 | 9830.39 | | | | | | | 16.83 | 0.78 | | 14.67 | 0.50 |
| Wheat | ALL | wang | T[air]^{plot} | Wang-Engel | 2019.00 | 9007.62 | 9027.78 | | | | | | | 17.98 | 0.86 | | 14.91 | 0.50 |
| Wheat | ALL | wang | T[air]^{plot} | Wang-Engel | 2021.00 | 9904.98 | 9925.51 | | | | | | | 17.11 | 0.78 | | 14.75 | 0.51 |
| Wheat | ALL | wang | T[air]^{ref} | Wang-Engel | 2015.00 | 10381.41 | 10402.08 | | | | | | | 16.71 | 0.81 | | 13.28 | 0.52 |
| Wheat | ALL | wang | T[air]^{ref} | Wang-Engel | 2016.00 | 9986.93 | 10007.42 | | | | | | | 16.72 | 0.74 | | 13.80 | 0.55 |
| Wheat | ALL | wang | T[air]^{ref} | Wang-Engel | 2017.00 | 9905.51 | 9926.12 | | | | | | | 15.15 | 0.68 | | 11.73 | 0.50 |
| Wheat | ALL | wang | T[air]^{ref} | Wang-Engel | 2018.00 | 10139.81 | 10160.31 | 16.74 | | | | | | 16.74 | 0.78 | | 14.34 | 0.52 |
| Wheat | ALL | wang | T[air]^{ref} | Wang-Engel | 2019.00 | 9278.99 | 9299.15 | | | | | | | 17.85 | 0.85 | | 14.32 | 0.50 |
| Wheat | ALL | wang | T[air]^{ref} | Wang-Engel | 2021.00 | 10198.94 | 10219.47 | | | | | | | 16.99 | 0.78 | | 14.35 | 0.52 |
| Wheat | CH CLARO | asym | T[air]^{plot} | Asymptotic | 2015.00 | 902.72 | 916.61 | | 5.20 | | | 4.71 | 0.54 | | | | 12.15 | 0.54 |
| Wheat | CH CLARO | asym | T[air]^{plot} | Asymptotic | 2016.00 | 873.76 | 887.48 | | 5.40 | | | 5.70 | 0.55 | | | | 12.27 | 0.52 |
| Wheat | CH CLARO | asym | T[air]^{plot} | Asymptotic | 2017.00 | 849.30 | 863.11 | 9.98 | 5.33 | | | 11.57 | 0.56 | | | | 9.01 | 0.45 |
| Wheat | CH CLARO | asym | T[air]^{plot} | Asymptotic | 2018.00 | 883.95 | 897.71 | | 6.13 | | | 11.26 | 0.59 | | | | 12.40 | 0.44 |
| Wheat | CH CLARO | asym | T[air]^{plot} | Asymptotic | 2019.00 | 779.22 | 792.34 | | 6.36 | | | 8.69 | 0.58 | | | | 12.76 | 0.52 |
| Wheat | CH CLARO | asym | T[air]^{plot} | Asymptotic | 2021.00 | 885.27 | 899.03 | | 5.61 | | | 9.58 | 0.57 | | | | 12.47 | 0.52 |
| Wheat | CH CLARO | asym | T[air]^{ref} | Asymptotic | 2015.00 | 932.37 | 946.27 | 14.42 | 5.12 | | | 6.65 | 0.54 | | | | 12.06 | 0.56 |
| Wheat | CH CLARO | asym | T[air]^{ref} | Asymptotic | 2016.00 | 914.86 | 928.59 | 14.97 | 6.79 | | | 7.47 | 0.59 | | | | 12.84 | 0.51 |
| Wheat | CH CLARO | asym | T[air]^{ref} | Asymptotic | 2017.00 | 848.46 | 862.27 | 9.90 | 5.20 | | | 6.97 | 0.56 | | | | 8.99 | 0.43 |
| Wheat | CH CLARO | asym | T[air]^{ref} | Asymptotic | 2018.00 | 910.33 | 924.10 | 14.01 | 5.10 | | | 4.82 | 0.58 | | | | 12.13 | 0.51 |
| Wheat | CH CLARO | asym | T[air]^{ref} | Asymptotic | 2019.00 | 805.94 | 819.07 | 15.07 | 5.40 | | | 4.65 | 0.58 | | | | 12.44 | 0.58 |
| Wheat | CH CLARO | asym | T[air]^{ref} | Asymptotic | 2021.00 | 915.19 | 928.96 | 14.69 | 5.25 | | | 6.99 | 0.57 | | | | 12.39 | 0.55 |
| Wheat | CH CLARO | bilinear | T[air]^{plot} | Bi-linear | 2015.00 | 905.44 | 919.33 | | 8.23 | 0.01 | 0.49 | | | | | | 12.29 | 0.55 |
| Wheat | CH CLARO | bilinear | T[air]^{plot} | Bi-linear | 2016.00 | 872.50 | 886.22 | | 7.31 | 0.02 | 0.45 | | | | | | 12.20 | 0.54 |
| Wheat | CH CLARO | bilinear | T[air]^{plot} | Bi-linear | 2017.00 | 834.64 | 848.45 | 9.39 | 6.60 | 0.00 | 0.52 | | | | | | 8.46 | 0.45 |
| Wheat | CH CLARO | bilinear | T[air]^{plot} | Bi-linear | 2018.00 | 885.33 | 899.10 | | 9.10 | 0.01 | 0.54 | | | | | | 12.48 | 0.48 |
| Wheat | CH CLARO | bilinear | T[air]^{plot} | Bi-linear | 2019.00 | 780.80 | 793.92 | | 8.80 | 0.02 | 0.49 | | | | | | 12.86 | 0.54 |
| Wheat | CH CLARO | bilinear | T[air]^{plot} | Bi-linear | 2021.00 | 887.31 | 901.08 | | 9.00 | 0.01 | 0.52 | | | | | | 12.59 | 0.55 |
| Wheat | CH CLARO | bilinear | T[air]^{ref} | Bi-linear | 2015.00 | 935.61 | 949.51 | 14.75 | 8.63 | 0.01 | 0.51 | | | | | | 12.22 | 0.56 |
| Wheat | CH CLARO | bilinear | T[air]^{ref} | Bi-linear | 2016.00 | 902.67 | 916.39 | 14.62 | 7.35 | 0.01 | 0.46 | | | | | | 12.14 | 0.55 |
| Wheat | CH CLARO | bilinear | T[air]^{ref} | Bi-linear | 2017.00 | 835.87 | 849.68 | 9.40 | 6.50 | 0.00 | 0.52 | | | | | | 8.51 | 0.44 |
| Wheat | CH CLARO | bilinear | T[air]^{ref} | Bi-linear | 2018.00 | 914.75 | 928.51 | 14.30 | 9.03 | 0.01 | 0.54 | | | | | | 12.36 | 0.51 |
| Wheat | CH CLARO | bilinear | T[air]^{ref} | Bi-linear | 2019.00 | 808.61 | 821.73 | 14.88 | 10.51 | 0.01 | 0.55 | | | | | | 12.61 | 0.54 |
| Wheat | CH CLARO | bilinear | T[air]^{ref} | Bi-linear | 2021.00 | 916.99 | 930.76 | 14.90 | 9.00 | 0.01 | 0.53 | | | | | | 12.48 | 0.56 |
| Wheat | CH CLARO | gauss | ^{plot} | Gaussian | 2015.00 | 1100.09 | 1108.43 | 27.65 | | | | | | | | 40.66 | 24.95 | 0.44 |
| Wheat | CH CLARO | gauss | ^{plot} | Gaussian | 2016.00 | 1080.43 | 1088.67 | 28.49 | | | | | | | | 42.22 | 26.94 | 0.33 |
| Wheat | CH CLARO | gauss | ^{plot} | Gaussian | 2017.00 | 1096.87 | 1105.16 | 27.54 | | | | | | | | 42.61 | 26.66 | 0.27 |

| | | | | | | | | | | | | | | | | | |
|---|---|---|---|---|---|---|---|---|---|---|---|---|---|---|---|---|---|
| Wheat | CH CLARO | gauss | ^{plot} | Gaussian | 2018.00 | 1108.66 | 1116.92 | 30.88 | | | | | | | 45.10 | 29.23 | 0.33 |
| Wheat | CH CLARO | gauss | ^{plot} | Gaussian | 2019.00 | 967.50 | 975.37 | 29.65 | | | | | | | 45.65 | 28.25 | 0.32 |
| Wheat | CH CLARO | gauss | ^{plot} | Gaussian | 2021.00 | 1109.13 | 1117.39 | 30.83 | | | | | | | 45.28 | 29.29 | 0.32 |
| Wheat | CH CLARO | gauss | ^{ref} | Gaussian | 2015.00 | 1100.09 | 1108.43 | 27.65 | | | | | | | 40.66 | 24.95 | 0.44 |
| Wheat | CH CLARO | gauss | ^{ref} | Gaussian | 2016.00 | 1080.43 | 1088.67 | 28.49 | | | | | | | 42.22 | 26.94 | 0.33 |
| Wheat | CH CLARO | gauss | ^{ref} | Gaussian | 2017.00 | 1096.87 | 1105.16 | 27.54 | | | | | | | 42.61 | 26.66 | 0.27 |
| Wheat | CH CLARO | gauss | ^{ref} | Gaussian | 2018.00 | 1108.66 | 1116.92 | 30.88 | | | | | | | 45.10 | 29.23 | 0.33 |
| Wheat | CH CLARO | gauss | ^{ref} | Gaussian | 2019.00 | 967.50 | 975.37 | 29.65 | | | | | | | 45.65 | 28.25 | 0.32 |
| Wheat | CH CLARO | gauss | ^{ref} | Gaussian | 2021.00 | 1109.13 | 1117.39 | 30.83 | | | | | | | 45.28 | 29.29 | 0.32 |
| Wheat | CH CLARO | linear | T[air]^{plot} | Linear | 2015.00 | 908.74 | 919.86 | | -7.82 | 0.02 | | | | | | 12.57 | 0.58 |
| Wheat | CH CLARO | linear | T[air]^{plot} | Linear | 2016.00 | 889.91 | 900.89 | | -47.66 | 0.01 | | | | | | 13.39 | 0.60 |
| Wheat | CH CLARO | linear | T[air]^{plot} | Linear | 2017.00 | 830.30 | 841.35 | 9.80 | -51.11 | 0.01 | | | | | | 8.38 | 0.53 |
| Wheat | CH CLARO | linear | T[air]^{plot} | Linear | 2018.00 | 909.67 | 920.69 | | -47.15 | 0.01 | | | | | | 14.13 | 0.56 |
| Wheat | CH CLARO | linear | T[air]^{plot} | Linear | 2019.00 | 804.00 | 814.50 | | -46.73 | 0.01 | | | | | | 14.73 | 0.59 |
| Wheat | CH CLARO | linear | T[air]^{plot} | Linear | 2021.00 | 907.64 | 918.66 | | -47.24 | 0.01 | | | | | | 14.00 | 0.60 |
| Wheat | CH CLARO | linear | T[air]^{ref} | Linear | 2015.00 | 940.27 | 951.38 | 15.19 | -8.62 | 0.02 | | | | | | 12.57 | 0.57 |
| Wheat | CH CLARO | linear | T[air]^{ref} | Linear | 2016.00 | 920.73 | 931.71 | 16.70 | -47.71 | 0.01 | | | | | | 13.31 | 0.61 |
| Wheat | CH CLARO | linear | T[air]^{ref} | Linear | 2017.00 | 830.01 | 841.06 | 9.78 | -51.11 | 0.01 | | | | | | 8.37 | 0.53 |
| Wheat | CH CLARO | linear | T[air]^{ref} | Linear | 2018.00 | 940.31 | 951.32 | 16.91 | -47.23 | 0.01 | | | | | | 14.01 | 0.57 |
| Wheat | CH CLARO | linear | T[air]^{ref} | Linear | 2019.00 | 814.07 | 824.57 | 15.07 | -3.81 | 0.03 | | | | | | 13.09 | 0.50 |
| Wheat | CH CLARO | linear | T[air]^{ref} | Linear | 2021.00 | 938.14 | 949.15 | 17.26 | -47.32 | 0.01 | | | | | | 13.88 | 0.61 |
| Wheat | CH CLARO | thermal | T[air]^{plot} | Thermal time | 2015.00 | 920.06 | 928.39 | | | | | | | | | | |
| Wheat | CH CLARO | thermal | T[air]^{plot} | Thermal time | 2016.00 | 884.47 | 892.70 | | | | | | | | | | |
| Wheat | CH CLARO | thermal | T[air]^{plot} | Thermal time | 2017.00 | 894.71 | 903.00 | 12.24 | | | | | | | | | |
| Wheat | CH CLARO | thermal | T[air]^{plot} | Thermal time | 2018.00 | 899.92 | 908.18 | | | | | | | | | | |
| Wheat | CH CLARO | thermal | T[air]^{plot} | Thermal time | 2019.00 | 785.94 | 793.82 | | | | | | | | | | |
| Wheat | CH CLARO | thermal | T[air]^{plot} | Thermal time | 2021.00 | 899.17 | 907.43 | | | | | | | | | | |
| Wheat | CH CLARO | thermal | T[air]^{ref} | Thermal time | 2015.00 | 954.33 | 962.67 | 15.49 | | | | | | | | | |
| Wheat | CH CLARO | thermal | T[air]^{ref} | Thermal time | 2016.00 | 918.14 | 926.37 | 15.01 | | | | | | | | | |
| Wheat | CH CLARO | thermal | T[air]^{ref} | Thermal time | 2017.00 | 898.59 | 906.88 | 12.37 | | | | | | | | | |
| Wheat | CH CLARO | thermal | T[air]^{ref} | Thermal time | 2018.00 | 933.70 | 941.96 | 14.99 | | | | | | | | | |
| Wheat | CH CLARO | thermal | T[air]^{ref} | Thermal time | 2019.00 | 815.46 | 823.34 | 14.72 | | | | | | | | | |
| Wheat | CH CLARO | thermal | T[air]^{ref} | Thermal time | 2021.00 | 932.70 | 940.96 | 15.55 | | | | | | | | | |
| Wheat | CH CLARO | wang | T[air]^{plot} | Wang-Engel | 2015.00 | 917.21 | 928.32 | | | | | | 15.02 | 0.64 | | 13.05 | 0.50 |
| Wheat | CH CLARO | wang | T[air]^{plot} | Wang-Engel | 2016.00 | 888.77 | 899.75 | | | | | | 15.70 | 0.66 | | 13.26 | 0.48 |
| Wheat | CH CLARO | wang | T[air]^{plot} | Wang-Engel | 2017.00 | 871.18 | 882.23 | 10.68 | | | | | 12.86 | 0.61 | | 9.99 | 0.37 |
| Wheat | CH CLARO | wang | T[air]^{plot} | Wang-Engel | 2018.00 | 891.66 | 902.67 | | | | | | 15.77 | 0.71 | | 12.96 | 0.41 |
| Wheat | CH CLARO | wang | T[air]^{plot} | Wang-Engel | 2019.00 | 790.59 | 801.09 | | | | | | 16.90 | 0.75 | | 13.67 | 0.46 |
| Wheat | CH CLARO | wang | T[air]^{plot} | Wang-Engel | 2021.00 | 899.65 | 910.66 | | | | | | 15.83 | 0.70 | | 13.43 | 0.47 |
| Wheat | CH CLARO | wang | T[air]^{ref} | Wang-Engel | 2015.00 | 943.02 | 954.13 | 15.31 | | | | | 15.22 | 0.65 | | 12.72 | 0.56 |
| Wheat | CH CLARO | wang | T[air]^{ref} | Wang-Engel | 2016.00 | 914.66 | 925.64 | 15.48 | | | | | 15.81 | 0.67 | | 12.91 | 0.55 |
| Wheat | CH CLARO | wang | T[air]^{ref} | Wang-Engel | 2017.00 | 870.50 | 881.55 | 10.66 | | | | | 12.87 | 0.61 | | 9.96 | 0.37 |
| Wheat | CH CLARO | wang | T[air]^{ref} | Wang-Engel | 2018.00 | 919.33 | 930.34 | 14.60 | | | | | 15.71 | 0.70 | | 12.72 | 0.50 |
| Wheat | CH CLARO | wang | T[air]^{ref} | Wang-Engel | 2019.00 | 812.82 | 823.32 | 15.09 | | | | | 17.02 | 0.76 | | 13.01 | 0.52 |
| Wheat | CH CLARO | wang | T[air]^{ref} | Wang-Engel | 2021.00 | 925.50 | 936.52 | 15.31 | | | | | 15.94 | 0.71 | | 13.07 | 0.53 |
| Wheat | CH NARA | asym | T[air]^{plot} | Asymptotic | 2015.00 | 713.10 | 726.02 | | 5.61 | | 9.33 | 0.48 | | | | 10.60 | 0.59 |
| Wheat | CH NARA | asym | T[air]^{plot} | Asymptotic | 2016.00 | 713.10 | 726.02 | | 5.61 | | 9.33 | 0.48 | | | | 10.60 | 0.59 |
| Wheat | CH NARA | asym | T[air]^{plot} | Asymptotic | 2017.00 | 531.81 | 543.47 | 9.25 | 4.49 | | 4.20 | 0.47 | | | | 7.84 | 0.56 |
| Wheat | CH NARA | asym | T[air]^{plot} | Asymptotic | 2018.00 | 571.73 | 583.51 | | 5.61 | | 9.58 | 0.50 | | | | 11.34 | 0.56 |
| Wheat | CH NARA | asym | T[air]^{plot} | Asymptotic | 2019.00 | 477.02 | 487.89 | | 5.61 | | 6.44 | 0.47 | | | | 11.86 | 0.62 |
| Wheat | CH NARA | asym | T[air]^{plot} | Asymptotic | 2021.00 | 544.54 | 556.13 | | 5.60 | | 9.59 | 0.45 | | | | 11.01 | 0.64 |
| Wheat | CH NARA | asym | T[air]^{ref} | Asymptotic | 2015.00 | 743.94 | 756.87 | 13.02 | 5.12 | | 4.79 | 0.48 | | | | 10.64 | 0.56 |

| | | | | | | | | | | | | | | | | |
|---|---|---|---|---|---|---|---|---|---|---|---|---|---|---|---|---|
| Wheat | CH NARA | asym | T[air]^{ref} | Asymptotic | 2016.00 | 743.94 | 756.87 | 13.02 | 5.12 | | | 4.79 | 0.48 | | 10.64 | 0.56 |
| Wheat | CH NARA | asym | T[air]^{ref} | Asymptotic | 2017.00 | 530.39 | 542.04 | 9.26 | 0.00 | | | 2.43 | 0.47 | | 7.77 | 0.58 |
| Wheat | CH NARA | asym | T[air]^{ref} | Asymptotic | 2018.00 | 600.79 | 612.58 | 13.59 | 5.71 | | | 10.40 | 0.52 | | 11.22 | 0.53 |
| Wheat | CH NARA | asym | T[air]^{ref} | Asymptotic | 2019.00 | 506.17 | 517.04 | 14.21 | 4.90 | | | 2.64 | 0.54 | | 11.59 | 0.54 |
| Wheat | CH NARA | asym | T[air]^{ref} | Asymptotic | 2021.00 | 575.82 | 587.41 | 13.80 | 5.12 | | | 4.68 | 0.46 | | 11.07 | 0.58 |
| Wheat | CH NARA | bilinear | T[air]^{plot} | Bi-linear | 2015.00 | 721.10 | 734.02 | | 8.80 | 0.00 | 0.45 | | | | 11.07 | 0.61 |
| Wheat | CH NARA | bilinear | T[air]^{plot} | Bi-linear | 2016.00 | 721.10 | 734.02 | | 8.80 | 0.00 | 0.45 | | | | 11.07 | 0.61 |
| Wheat | CH NARA | bilinear | T[air]^{plot} | Bi-linear | 2017.00 | 525.84 | 537.49 | 9.01 | 7.37 | -0.01 | 0.50 | | | | 7.54 | 0.59 |
| Wheat | CH NARA | bilinear | T[air]^{plot} | Bi-linear | 2018.00 | 579.77 | 591.55 | | 10.03 | 0.00 | 0.50 | | | | 11.98 | 0.58 |
| Wheat | CH NARA | bilinear | T[air]^{plot} | Bi-linear | 2019.00 | 480.91 | 491.79 | | 8.80 | 0.02 | 0.38 | | | | 12.25 | 0.63 |
| Wheat | CH NARA | bilinear | T[air]^{plot} | Bi-linear | 2021.00 | 547.06 | 558.65 | | 10.47 | 0.00 | 0.46 | | | | 11.21 | 0.66 |
| Wheat | CH NARA | bilinear | T[air]^{ref} | Bi-linear | 2015.00 | 750.55 | 763.48 | 13.67 | 10.51 | -0.00 | 0.51 | | | | 11.01 | 0.58 |
| Wheat | CH NARA | bilinear | T[air]^{ref} | Bi-linear | 2016.00 | 750.55 | 763.48 | 13.67 | 10.51 | -0.00 | 0.51 | | | | 11.01 | 0.58 |
| Wheat | CH NARA | bilinear | T[air]^{ref} | Bi-linear | 2017.00 | 525.64 | 537.30 | 8.99 | 7.18 | -0.01 | 0.51 | | | | 7.53 | 0.59 |
| Wheat | CH NARA | bilinear | T[air]^{ref} | Bi-linear | 2018.00 | 607.96 | 619.74 | 14.59 | 12.22 | -0.01 | 0.59 | | | | 11.75 | 0.57 |
| Wheat | CH NARA | bilinear | T[air]^{ref} | Bi-linear | 2019.00 | 510.08 | 520.95 | 14.97 | 12.60 | 0.00 | 0.54 | | | | 12.04 | 0.58 |
| Wheat | CH NARA | bilinear | T[air]^{ref} | Bi-linear | 2021.00 | 576.72 | 588.30 | 14.31 | 11.13 | -0.01 | 0.50 | | | | 11.14 | 0.62 |
| Wheat | CH NARA | gauss | ^{plot} | Gaussian | 2015.00 | 872.09 | 879.84 | 22.48 | | | | | | 35.59 | 21.02 | 0.37 |
| Wheat | CH NARA | gauss | ^{plot} | Gaussian | 2016.00 | 872.09 | 879.84 | 22.48 | | | | | | 35.59 | 21.02 | 0.37 |
| Wheat | CH NARA | gauss | ^{plot} | Gaussian | 2017.00 | 664.66 | 671.65 | 20.16 | | | | | | 34.42 | 19.53 | 0.27 |
| Wheat | CH NARA | gauss | ^{plot} | Gaussian | 2018.00 | 705.39 | 712.46 | 24.42 | | | | | | 36.28 | 22.70 | 0.38 |
| Wheat | CH NARA | gauss | ^{plot} | Gaussian | 2019.00 | 568.80 | 575.32 | 21.16 | | | | | | 36.11 | 19.65 | 0.39 |
| Wheat | CH NARA | gauss | ^{plot} | Gaussian | 2021.00 | 672.36 | 679.32 | 23.64 | | | | | | 35.41 | 21.84 | 0.40 |
| Wheat | CH NARA | gauss | ^{ref} | Gaussian | 2015.00 | 872.09 | 879.84 | 22.48 | | | | | | 35.59 | 21.02 | 0.37 |
| Wheat | CH NARA | gauss | ^{ref} | Gaussian | 2016.00 | 872.09 | 879.84 | 22.48 | | | | | | 35.59 | 21.02 | 0.37 |
| Wheat | CH NARA | gauss | ^{ref} | Gaussian | 2017.00 | 664.66 | 671.65 | 20.16 | | | | | | 34.42 | 19.53 | 0.27 |
| Wheat | CH NARA | gauss | ^{ref} | Gaussian | 2018.00 | 705.39 | 712.46 | 24.42 | | | | | | 36.28 | 22.70 | 0.38 |
| Wheat | CH NARA | gauss | ^{ref} | Gaussian | 2019.00 | 568.80 | 575.32 | 21.16 | | | | | | 36.11 | 19.65 | 0.39 |
| Wheat | CH NARA | gauss | ^{ref} | Gaussian | 2021.00 | 672.36 | 679.32 | 23.64 | | | | | | 35.41 | 21.84 | 0.40 |
| Wheat | CH NARA | linear | T[air]^{plot} | Linear | 2015.00 | 736.42 | 746.76 | | -48.48 | 0.01 | | | | | 12.19 | 0.66 |
| Wheat | CH NARA | linear | T[air]^{plot} | Linear | 2016.00 | 736.42 | 746.76 | | -48.48 | 0.01 | | | | | 12.19 | 0.66 |
| Wheat | CH NARA | linear | T[air]^{plot} | Linear | 2017.00 | 527.39 | 536.71 | 9.58 | -51.56 | 0.01 | | | | | 7.72 | 0.62 |
| Wheat | CH NARA | linear | T[air]^{plot} | Linear | 2018.00 | 595.17 | 604.60 | | -47.54 | 0.01 | | | | | 13.57 | 0.65 |
| Wheat | CH NARA | linear | T[air]^{plot} | Linear | 2019.00 | 481.37 | 490.06 | | -2.09 | 0.03 | | | | | 12.50 | 0.63 |
| Wheat | CH NARA | linear | T[air]^{plot} | Linear | 2021.00 | 560.38 | 569.65 | | -48.22 | 0.01 | | | | | 12.57 | 0.69 |
| Wheat | CH NARA | linear | T[air]^{ref} | Linear | 2015.00 | 767.46 | 777.80 | 15.62 | -48.49 | 0.01 | | | | | 12.17 | 0.63 |
| Wheat | CH NARA | linear | T[air]^{ref} | Linear | 2016.00 | 767.46 | 777.80 | 15.62 | -48.49 | 0.01 | | | | | 12.17 | 0.63 |
| Wheat | CH NARA | linear | T[air]^{ref} | Linear | 2017.00 | 527.51 | 536.83 | 9.59 | -51.56 | 0.01 | | | | | 7.72 | 0.62 |
| Wheat | CH NARA | linear | T[air]^{ref} | Linear | 2018.00 | 626.13 | 635.56 | 17.12 | -47.61 | 0.01 | | | | | 13.46 | 0.62 |
| Wheat | CH NARA | linear | T[air]^{ref} | Linear | 2019.00 | 526.34 | 535.04 | 18.19 | -47.27 | 0.01 | | | | | 13.95 | 0.66 |
| Wheat | CH NARA | linear | T[air]^{ref} | Linear | 2021.00 | 592.17 | 601.44 | 16.39 | -48.20 | 0.01 | | | | | 12.58 | 0.64 |
| Wheat | CH NARA | thermal | T[air]^{plot} | Thermal time | 2015.00 | 734.14 | 741.89 | | | | | | | | | |
| Wheat | CH NARA | thermal | T[air]^{plot} | Thermal time | 2016.00 | 734.14 | 741.89 | | | | | | | | | |
| Wheat | CH NARA | thermal | T[air]^{plot} | Thermal time | 2017.00 | 573.09 | 580.08 | 12.44 | | | | | | | | |
| Wheat | CH NARA | thermal | T[air]^{plot} | Thermal time | 2018.00 | 586.56 | 593.63 | | | | | | | | | |
| Wheat | CH NARA | thermal | T[air]^{plot} | Thermal time | 2019.00 | 479.83 | 486.35 | | | | | | | | | |
| Wheat | CH NARA | thermal | T[air]^{plot} | Thermal time | 2021.00 | 553.97 | 560.92 | | | | | | | | | |
| Wheat | CH NARA | thermal | T[air]^{ref} | Thermal time | 2015.00 | 769.48 | 777.23 | 14.53 | | | | | | | | |
| Wheat | CH NARA | thermal | T[air]^{ref} | Thermal time | 2016.00 | 769.48 | 777.23 | 14.53 | | | | | | | | |
| Wheat | CH NARA | thermal | T[air]^{ref} | Thermal time | 2017.00 | 577.57 | 584.56 | 12.73 | | | | | | | | |
| Wheat | CH NARA | thermal | T[air]^{ref} | Thermal time | 2018.00 | 621.68 | 628.75 | 15.37 | | | | | | | | |
| Wheat | CH NARA | thermal | T[air]^{ref} | Thermal time | 2019.00 | 511.75 | 518.27 | 14.75 | | | | | | | | |

| | | | | | | | | | | | | | | | | | | |
|---|---|---|---|---|---|---|---|---|---|---|---|---|---|---|---|---|---|---|
| Wheat | CH NARA | thermal | T[air]^{ref} | Thermal time | 2021.00 | 589.95 | 596.91 | 14.95 | | | | | | | | | | |
| Wheat | CH NARA | wang | T[air]^{plot} | Wang-Engel | 2015.00 | 728.60 | 738.94 | | | | | | | 14.79 | 0.54 | | 11.65 | 0.60 |
| Wheat | CH NARA | wang | T[air]^{plot} | Wang-Engel | 2016.00 | 728.60 | 738.94 | | | | | | | 14.79 | 0.54 | | 11.65 | 0.60 |
| Wheat | CH NARA | wang | T[air]^{plot} | Wang-Engel | 2017.00 | 538.75 | 548.07 | 9.81 | | | | | | 11.70 | 0.51 | | 8.33 | 0.57 |
| Wheat | CH NARA | wang | T[air]^{plot} | Wang-Engel | 2018.00 | 581.15 | 590.58 | | | | | | | 15.45 | 0.60 | | 12.27 | 0.57 |
| Wheat | CH NARA | wang | T[air]^{plot} | Wang-Engel | 2019.00 | 484.56 | 493.26 | | | | | | | 16.66 | 0.58 | | 12.84 | 0.65 |
| Wheat | CH NARA | wang | T[air]^{plot} | Wang-Engel | 2021.00 | 551.52 | 560.79 | | | | | | | 15.24 | 0.52 | | 11.74 | 0.65 |
| Wheat | CH NARA | wang | T[air]^{ref} | Wang-Engel | 2015.00 | 753.40 | 763.74 | 14.21 | | | | | | 14.90 | 0.55 | | 11.28 | 0.61 |
| Wheat | CH NARA | wang | T[air]^{ref} | Wang-Engel | 2016.00 | 753.40 | 763.74 | 14.21 | | | | | | 14.90 | 0.55 | | 11.28 | 0.61 |
| Wheat | CH NARA | wang | T[air]^{ref} | Wang-Engel | 2017.00 | 537.99 | 547.31 | 9.79 | | | | | | 11.53 | 0.50 | | 8.28 | 0.58 |
| Wheat | CH NARA | wang | T[air]^{ref} | Wang-Engel | 2018.00 | 606.49 | 615.91 | 14.74 | | | | | | 15.40 | 0.61 | | 11.79 | 0.59 |
| Wheat | CH NARA | wang | T[air]^{ref} | Wang-Engel | 2019.00 | 508.73 | 517.43 | 15.32 | | | | | | 16.68 | 0.61 | | 12.10 | 0.61 |
| Wheat | CH NARA | wang | T[air]^{ref} | Wang-Engel | 2021.00 | 576.89 | 586.16 | 14.75 | | | | | | 15.30 | 0.54 | | 11.30 | 0.64 |
| Wheat | FASTNET | asym | T[air]^{plot} | Asymptotic | 2015.00 | 848.45 | 862.34 | | 5.21 | | | 3.22 | 0.54 | | | | 9.57 | 0.47 |
| Wheat | FASTNET | asym | T[air]^{plot} | Asymptotic | 2016.00 | 805.42 | 819.06 | | 6.07 | | | 6.34 | 0.50 | | | | 9.62 | 0.49 |
| Wheat | FASTNET | asym | T[air]^{plot} | Asymptotic | 2017.00 | 819.39 | 833.25 | 8.86 | 5.59 | | | 4.70 | 0.49 | | | | 7.69 | 0.49 |
| Wheat | FASTNET | asym | T[air]^{plot} | Asymptotic | 2018.00 | 808.03 | 821.63 | | 6.10 | | | 6.44 | 0.51 | | | | 10.08 | 0.47 |
| Wheat | FASTNET | asym | T[air]^{plot} | Asymptotic | 2019.00 | 772.29 | 785.65 | | 5.20 | | | 2.68 | 0.54 | | | | 10.15 | 0.50 |
| Wheat | FASTNET | asym | T[air]^{plot} | Asymptotic | 2021.00 | 828.37 | 842.13 | | 5.97 | | | 3.55 | 0.52 | | | | 9.65 | 0.56 |
| Wheat | FASTNET | asym | T[air]^{ref} | Asymptotic | 2015.00 | 857.65 | 871.55 | | 5.48 | | | 4.35 | 0.52 | | | | 9.65 | 0.48 |
| Wheat | FASTNET | asym | T[air]^{ref} | Asymptotic | 2016.00 | 816.38 | 830.02 | | 5.48 | | | 4.18 | 0.50 | | | | 9.78 | 0.50 |
| Wheat | FASTNET | asym | T[air]^{ref} | Asymptotic | 2017.00 | 802.12 | 815.97 | | 5.30 | | | 4.49 | 0.49 | | | | 7.81 | 0.47 |
| Wheat | FASTNET | asym | T[air]^{ref} | Asymptotic | 2018.00 | 838.56 | 852.15 | 11.59 | 5.74 | | | 6.73 | 0.51 | | | | 10.11 | 0.48 |
| Wheat | FASTNET | asym | T[air]^{ref} | Asymptotic | 2019.00 | 779.91 | 793.28 | | 5.10 | | | 2.57 | 0.56 | | | | 10.17 | 0.47 |
| Wheat | FASTNET | asym | T[air]^{ref} | Asymptotic | 2021.00 | 839.81 | 853.58 | | 6.10 | | | 7.82 | 0.50 | | | | 9.83 | 0.53 |
| Wheat | FASTNET | bilinear | T[air]^{plot} | Bi-linear | 2015.00 | 854.98 | 868.87 | | 11.24 | 0.01 | 0.53 | | | | | | 9.85 | 0.50 |
| Wheat | FASTNET | bilinear | T[air]^{plot} | Bi-linear | 2016.00 | 813.15 | 826.79 | | 10.48 | 0.00 | 0.49 | | | | | | 9.97 | 0.55 |
| Wheat | FASTNET | bilinear | T[air]^{plot} | Bi-linear | 2017.00 | 826.74 | 840.60 | 8.89 | 9.00 | 0.00 | 0.47 | | | | | | 7.93 | 0.45 |
| Wheat | FASTNET | bilinear | T[air]^{plot} | Bi-linear | 2018.00 | 816.84 | 830.43 | | 10.72 | 0.01 | 0.49 | | | | | | 10.50 | 0.49 |
| Wheat | FASTNET | bilinear | T[air]^{plot} | Bi-linear | 2019.00 | 777.10 | 790.46 | | 10.46 | 0.02 | 0.45 | | | | | | 10.40 | 0.50 |
| Wheat | FASTNET | bilinear | T[air]^{plot} | Bi-linear | 2021.00 | 831.53 | 845.29 | | 10.96 | 0.01 | 0.48 | | | | | | 9.79 | 0.59 |
| Wheat | FASTNET | bilinear | T[air]^{ref} | Bi-linear | 2015.00 | 860.64 | 874.53 | | 14.23 | -0.02 | 0.66 | | | | | | 9.77 | 0.50 |
| Wheat | FASTNET | bilinear | T[air]^{ref} | Bi-linear | 2016.00 | 820.58 | 834.22 | | 12.60 | -0.01 | 0.57 | | | | | | 9.97 | 0.55 |
| Wheat | FASTNET | bilinear | T[air]^{ref} | Bi-linear | 2017.00 | 807.85 | 821.71 | | 8.98 | 0.00 | 0.48 | | | | | | 8.01 | 0.45 |
| Wheat | FASTNET | bilinear | T[air]^{ref} | Bi-linear | 2018.00 | 844.83 | 858.42 | 11.88 | 13.09 | -0.01 | 0.59 | | | | | | 10.40 | 0.49 |
| Wheat | FASTNET | bilinear | T[air]^{ref} | Bi-linear | 2019.00 | 785.27 | 798.64 | | 12.80 | 0.00 | 0.55 | | | | | | 10.43 | 0.48 |
| Wheat | FASTNET | bilinear | T[air]^{ref} | Bi-linear | 2021.00 | 840.71 | 854.48 | | 12.60 | -0.00 | 0.55 | | | | | | 9.87 | 0.56 |
| Wheat | FASTNET | gauss | ^{plot} | Gaussian | 2015.00 | 1067.38 | 1075.72 | 23.13 | | | | | | | | 37.11 | 21.72 | 0.35 |
| Wheat | FASTNET | gauss | ^{plot} | Gaussian | 2016.00 | 1014.09 | 1022.27 | 22.74 | | | | | | | | 39.12 | 21.79 | 0.29 |
| Wheat | FASTNET | gauss | ^{plot} | Gaussian | 2017.00 | 1054.65 | 1062.96 | 22.05 | | | | | | | | 38.74 | 21.38 | 0.25 |
| Wheat | FASTNET | gauss | ^{plot} | Gaussian | 2018.00 | 1026.09 | 1034.24 | 25.13 | | | | | | | | 40.77 | 23.95 | 0.31 |
| Wheat | FASTNET | gauss | ^{plot} | Gaussian | 2019.00 | 962.72 | 970.74 | 23.32 | | | | | | | | 40.38 | 22.06 | 0.33 |
| Wheat | FASTNET | gauss | ^{plot} | Gaussian | 2021.00 | 1054.23 | 1062.49 | 24.07 | | | | | | | | 40.84 | 23.07 | 0.29 |
| Wheat | FASTNET | gauss | ^{ref} | Gaussian | 2015.00 | 1067.38 | 1075.72 | 23.13 | | | | | | | | 37.11 | 21.72 | 0.35 |
| Wheat | FASTNET | gauss | ^{ref} | Gaussian | 2016.00 | 1014.09 | 1022.27 | 22.74 | | | | | | | | 39.12 | 21.79 | 0.29 |
| Wheat | FASTNET | gauss | ^{ref} | Gaussian | 2017.00 | 1054.65 | 1062.96 | 22.05 | | | | | | | | 38.74 | 21.38 | 0.25 |
| Wheat | FASTNET | gauss | ^{ref} | Gaussian | 2018.00 | 1026.09 | 1034.24 | 25.13 | | | | | | | | 40.77 | 23.95 | 0.31 |
| Wheat | FASTNET | gauss | ^{ref} | Gaussian | 2019.00 | 962.72 | 970.74 | 23.32 | | | | | | | | 40.38 | 22.06 | 0.33 |
| Wheat | FASTNET | gauss | ^{ref} | Gaussian | 2021.00 | 1054.23 | 1062.49 | 24.07 | | | | | | | | 40.84 | 23.07 | 0.29 |
| Wheat | FASTNET | linear | T[air]^{plot} | Linear | 2015.00 | 891.23 | 902.34 | | -48.82 | 0.01 | | | | | | | 11.71 | 0.57 |
| Wheat | FASTNET | linear | T[air]^{plot} | Linear | 2016.00 | 840.05 | 850.96 | | -49.00 | 0.01 | | | | | | | 11.44 | 0.62 |
| Wheat | FASTNET | linear | T[air]^{plot} | Linear | 2017.00 | 839.46 | 850.54 | 9.85 | -51.05 | 0.01 | | | | | | | 8.47 | 0.51 |

| | | | | | | | | | | | | | | | | | |
|---|---|---|---|---|---|---|---|---|---|---|---|---|---|---|---|---|---|
| Wheat | FASTNET | linear | T[air]^{plot} | Linear | 2018.00 | 848.11 | 858.98 | | -48.39 | 0.01 | | | | | | 12.33 | 0.56 |
| Wheat | FASTNET | linear | T[air]^{plot} | Linear | 2019.00 | 808.76 | 819.46 | | -48.39 | 0.01 | | | | | | 12.33 | 0.58 |
| Wheat | FASTNET | linear | T[air]^{plot} | Linear | 2021.00 | 866.67 | 877.68 | | -48.86 | 0.01 | | | | | | 11.64 | 0.62 |
| Wheat | FASTNET | linear | T[air]^{ref} | Linear | 2015.00 | 900.68 | 911.80 | | -48.76 | 0.01 | | | | | | 11.79 | 0.56 |
| Wheat | FASTNET | linear | T[air]^{ref} | Linear | 2016.00 | 850.25 | 861.16 | | -48.91 | 0.01 | | | | | | 11.57 | 0.60 |
| Wheat | FASTNET | linear | T[air]^{ref} | Linear | 2017.00 | 819.63 | 830.71 | | -51.01 | 0.01 | | | | | | 8.52 | 0.51 |
| Wheat | FASTNET | linear | T[air]^{ref} | Linear | 2018.00 | 855.91 | 866.79 | 12.35 | -5.21 | 0.03 | | | | | | 11.03 | 0.45 |
| Wheat | FASTNET | linear | T[air]^{ref} | Linear | 2019.00 | 791.25 | 801.94 | | -2.64 | 0.03 | | | | | | 10.84 | 0.42 |
| Wheat | FASTNET | linear | T[air]^{ref} | Linear | 2021.00 | 876.51 | 887.53 | | -48.79 | 0.01 | | | | | | 11.75 | 0.60 |
| Wheat | FASTNET | thermal | T[air]^{plot} | Thermal time | 2015.00 | 868.02 | 876.36 | | | | | | | | | | |
| Wheat | FASTNET | thermal | T[air]^{plot} | Thermal time | 2016.00 | 830.79 | 838.97 | | | | | | | | | | |
| Wheat | FASTNET | thermal | T[air]^{plot} | Thermal time | 2017.00 | 862.22 | 870.53 | 10.27 | | | | | | | | | |
| Wheat | FASTNET | thermal | T[air]^{plot} | Thermal time | 2018.00 | 826.88 | 835.04 | | | | | | | | | | |
| Wheat | FASTNET | thermal | T[air]^{plot} | Thermal time | 2019.00 | 779.87 | 787.89 | | | | | | | | | | |
| Wheat | FASTNET | thermal | T[air]^{plot} | Thermal time | 2021.00 | 841.85 | 850.11 | | | | | | | | | | |
| Wheat | FASTNET | thermal | T[air]^{ref} | Thermal time | 2015.00 | 884.27 | 892.61 | | | | | | | | | | |
| Wheat | FASTNET | thermal | T[air]^{ref} | Thermal time | 2016.00 | 846.22 | 854.41 | | | | | | | | | | |
| Wheat | FASTNET | thermal | T[air]^{ref} | Thermal time | 2017.00 | 849.18 | 857.49 | | | | | | | | | | |
| Wheat | FASTNET | thermal | T[air]^{ref} | Thermal time | 2018.00 | 862.90 | 871.05 | 12.43 | | | | | | | | | |
| Wheat | FASTNET | thermal | T[air]^{ref} | Thermal time | 2019.00 | 791.43 | 799.45 | | | | | | | | | | |
| Wheat | FASTNET | thermal | T[air]^{ref} | Thermal time | 2021.00 | 858.59 | 866.85 | | | | | | | | | | |
| Wheat | FASTNET | wang | T[air]^{plot} | Wang-Engel | 2015.00 | 859.45 | 870.56 | | | | | | | 16.50 | 0.69 | 10.13 | 0.46 |
| Wheat | FASTNET | wang | T[air]^{plot} | Wang-Engel | 2016.00 | 821.00 | 831.91 | | | | | | | 16.06 | 0.59 | 10.43 | 0.59 |
| Wheat | FASTNET | wang | T[air]^{plot} | Wang-Engel | 2017.00 | 848.98 | 860.06 | 9.99 | | | | | | 15.24 | 0.57 | 8.80 | 0.47 |
| Wheat | FASTNET | wang | T[air]^{plot} | Wang-Engel | 2018.00 | 821.84 | 832.71 | | | | | | | 16.58 | 0.65 | 10.85 | 0.49 |
| Wheat | FASTNET | wang | T[air]^{plot} | Wang-Engel | 2019.00 | 786.51 | 797.20 | | | | | | | 17.37 | 0.67 | 10.99 | 0.53 |
| Wheat | FASTNET | wang | T[air]^{plot} | Wang-Engel | 2021.00 | 843.67 | 854.69 | | | | | | | 16.83 | 0.64 | 10.44 | 0.59 |
| Wheat | FASTNET | wang | T[air]^{ref} | Wang-Engel | 2015.00 | 860.26 | 871.38 | | | | | | | 16.29 | 0.67 | 9.84 | 0.48 |
| Wheat | FASTNET | wang | T[air]^{ref} | Wang-Engel | 2016.00 | 822.07 | 832.98 | | | | | | | 15.77 | 0.58 | 10.13 | 0.59 |
| Wheat | FASTNET | wang | T[air]^{ref} | Wang-Engel | 2017.00 | 824.74 | 835.83 | | | | | | | 14.89 | 0.56 | 8.70 | 0.47 |
| Wheat | FASTNET | wang | T[air]^{ref} | Wang-Engel | 2018.00 | 847.42 | 858.30 | 12.27 | | | | | | 16.39 | 0.64 | 10.61 | 0.50 |
| Wheat | FASTNET | wang | T[air]^{ref} | Wang-Engel | 2019.00 | 787.96 | 798.65 | | | | | | | 17.10 | 0.66 | 10.67 | 0.52 |
| Wheat | FASTNET | wang | T[air]^{ref} | Wang-Engel | 2021.00 | 846.09 | 857.10 | | | | | | | 16.51 | 0.62 | 10.20 | 0.57 |
| Wheat | MARKSMAN | asym | T[air]^{plot} | Asymptotic | 2015.00 | 745.14 | 758.26 | | 5.31 | | | 4.44 | 0.54 | | | 10.70 | 0.47 |
| Wheat | MARKSMAN | asym | T[air]^{plot} | Asymptotic | 2016.00 | 699.95 | 712.77 | | 5.61 | | | 10.76 | 0.51 | | | 10.29 | 0.50 |
| Wheat | MARKSMAN | asym | T[air]^{plot} | Asymptotic | 2017.00 | 754.40 | 767.62 | 9.86 | 5.30 | | | 5.17 | 0.51 | | | 8.99 | 0.41 |
| Wheat | MARKSMAN | asym | T[air]^{plot} | Asymptotic | 2018.00 | 730.37 | 743.34 | | 5.61 | | | 9.27 | 0.52 | | | 11.17 | 0.45 |
| Wheat | MARKSMAN | asym | T[air]^{plot} | Asymptotic | 2019.00 | 621.23 | 633.39 | | 5.30 | | | 4.62 | 0.53 | | | 11.59 | 0.41 |
| Wheat | MARKSMAN | asym | T[air]^{plot} | Asymptotic | 2021.00 | 727.79 | 740.81 | | 5.30 | | | 3.94 | 0.51 | | | 10.58 | 0.52 |
| Wheat | MARKSMAN | asym | T[air]^{ref} | Asymptotic | 2015.00 | 777.94 | 791.06 | 12.26 | 3.57 | | | 2.51 | 0.58 | | | 10.83 | 0.47 |
| Wheat | MARKSMAN | asym | T[air]^{ref} | Asymptotic | 2016.00 | 723.05 | 735.87 | 11.98 | 5.30 | | | 4.86 | 0.52 | | | 10.32 | 0.48 |
| Wheat | MARKSMAN | asym | T[air]^{ref} | Asymptotic | 2017.00 | 762.88 | 776.10 | 10.14 | 0.00 | | | 1.87 | 0.54 | | | 9.35 | 0.39 |
| Wheat | MARKSMAN | asym | T[air]^{ref} | Asymptotic | 2018.00 | 761.16 | 774.13 | 12.50 | 5.34 | | | 10.81 | 0.52 | | | 11.17 | 0.45 |
| Wheat | MARKSMAN | asym | T[air]^{ref} | Asymptotic | 2019.00 | 650.91 | 663.07 | 12.64 | 5.34 | | | 8.57 | 0.53 | | | 11.50 | 0.42 |
| Wheat | MARKSMAN | asym | T[air]^{ref} | Asymptotic | 2021.00 | 760.77 | 773.80 | 12.32 | 3.90 | | | 2.62 | 0.54 | | | 10.73 | 0.50 |
| Wheat | MARKSMAN | bilinear | T[air]^{plot} | Bi-linear | 2015.00 | 750.96 | 764.08 | | 9.33 | 0.01 | 0.51 | | | | | 11.03 | 0.51 |
| Wheat | MARKSMAN | bilinear | T[air]^{plot} | Bi-linear | 2016.00 | 710.21 | 723.03 | | 8.77 | 0.00 | 0.50 | | | | | 10.88 | 0.50 |
| Wheat | MARKSMAN | bilinear | T[air]^{plot} | Bi-linear | 2017.00 | 758.19 | 771.42 | 9.93 | 7.70 | 0.00 | 0.49 | | | | | 9.15 | 0.39 |
| Wheat | MARKSMAN | bilinear | T[air]^{plot} | Bi-linear | 2018.00 | 737.17 | 750.14 | | 7.99 | 0.01 | 0.45 | | | | | 11.58 | 0.47 |
| Wheat | MARKSMAN | bilinear | T[air]^{plot} | Bi-linear | 2019.00 | 626.26 | 638.41 | | 8.16 | 0.01 | 0.45 | | | | | 11.96 | 0.42 |
| Wheat | MARKSMAN | bilinear | T[air]^{plot} | Bi-linear | 2021.00 | 731.37 | 744.40 | | 10.08 | 0.00 | 0.50 | | | | | 10.78 | 0.53 |
| Wheat | MARKSMAN | bilinear | T[air]^{ref} | Bi-linear | 2015.00 | 780.36 | 793.48 | 12.56 | 11.60 | -0.00 | 0.60 | | | | | 10.97 | 0.49 |

| | | | | | | | | | | | | | | | | |
|---|---|---|---|---|---|---|---|---|---|---|---|---|---|---|---|---|
| Wheat | MARKSMAN | bilinear | T[air]^{ref} | Bi-linear | 2016.00 | 731.44 | 744.26 | 12.61 | 8.73 | 0.00 | 0.51 | | | | 10.78 | 0.50 |
| Wheat | MARKSMAN | bilinear | T[air]^{ref} | Bi-linear | 2017.00 | 758.70 | 771.92 | 9.95 | 7.70 | 0.00 | 0.50 | | | | 9.17 | 0.39 |
| Wheat | MARKSMAN | bilinear | T[air]^{ref} | Bi-linear | 2018.00 | 767.72 | 780.69 | 13.01 | 7.98 | 0.01 | 0.46 | | | | 11.56 | 0.47 |
| Wheat | MARKSMAN | bilinear | T[air]^{ref} | Bi-linear | 2019.00 | 655.55 | 667.71 | 12.94 | 8.56 | 0.01 | 0.47 | | | | 11.82 | 0.41 |
| Wheat | MARKSMAN | bilinear | T[air]^{ref} | Bi-linear | 2021.00 | 761.89 | 774.91 | 12.53 | 10.84 | 0.00 | 0.53 | | | | 10.79 | 0.52 |
| Wheat | MARKSMAN | gauss | ^{plot} | Gaussian | 2015.00 | 938.93 | 946.80 | 26.12 | | | | | | 39.13 | 24.52 | 0.35 |
| Wheat | MARKSMAN | gauss | ^{plot} | Gaussian | 2016.00 | 872.48 | 880.17 | 24.28 | | | | | | 39.45 | 23.14 | 0.31 |
| Wheat | MARKSMAN | gauss | ^{plot} | Gaussian | 2017.00 | 946.12 | 954.06 | 24.37 | | | | | | 39.98 | 23.21 | 0.31 |
| Wheat | MARKSMAN | gauss | ^{plot} | Gaussian | 2018.00 | 914.29 | 922.07 | 26.58 | | | | | | 41.83 | 24.91 | 0.36 |
| Wheat | MARKSMAN | gauss | ^{plot} | Gaussian | 2019.00 | 766.24 | 773.54 | 24.81 | | | | | | 43.02 | 23.59 | 0.32 |
| Wheat | MARKSMAN | gauss | ^{plot} | Gaussian | 2021.00 | 916.45 | 924.26 | 25.56 | | | | | | 41.91 | 24.03 | 0.35 |
| Wheat | MARKSMAN | gauss | ^{ref} | Gaussian | 2015.00 | 938.93 | 946.80 | 26.12 | | | | | | 39.13 | 24.52 | 0.35 |
| Wheat | MARKSMAN | gauss | ^{ref} | Gaussian | 2016.00 | 872.48 | 880.17 | 24.28 | | | | | | 39.45 | 23.14 | 0.31 |
| Wheat | MARKSMAN | gauss | ^{ref} | Gaussian | 2017.00 | 946.12 | 954.06 | 24.37 | | | | | | 39.98 | 23.21 | 0.31 |
| Wheat | MARKSMAN | gauss | ^{ref} | Gaussian | 2018.00 | 914.29 | 922.07 | 26.58 | | | | | | 41.83 | 24.91 | 0.36 |
| Wheat | MARKSMAN | gauss | ^{ref} | Gaussian | 2019.00 | 766.24 | 773.54 | 24.81 | | | | | | 43.02 | 23.59 | 0.32 |
| Wheat | MARKSMAN | gauss | ^{ref} | Gaussian | 2021.00 | 916.45 | 924.26 | 25.56 | | | | | | 41.91 | 24.03 | 0.35 |
| Wheat | MARKSMAN | linear | T[air]^{plot} | Linear | 2015.00 | 755.26 | 765.76 | | -8.26 | 0.02 | | | | | 11.39 | 0.56 |
| Wheat | MARKSMAN | linear | T[air]^{plot} | Linear | 2016.00 | 724.45 | 734.70 | | -48.68 | 0.01 | | | | | 11.91 | 0.56 |
| Wheat | MARKSMAN | linear | T[air]^{plot} | Linear | 2017.00 | 765.90 | 776.48 | 10.63 | -50.27 | 0.01 | | | | | 9.60 | 0.44 |
| Wheat | MARKSMAN | linear | T[air]^{plot} | Linear | 2018.00 | 752.15 | 762.53 | | -48.12 | 0.01 | | | | | 12.72 | 0.52 |
| Wheat | MARKSMAN | linear | T[air]^{plot} | Linear | 2019.00 | 641.02 | 650.74 | | -47.68 | 0.01 | | | | | 13.35 | 0.48 |
| Wheat | MARKSMAN | linear | T[air]^{plot} | Linear | 2021.00 | 748.68 | 759.10 | | -48.63 | 0.01 | | | | | 11.98 | 0.56 |
| Wheat | MARKSMAN | linear | T[air]^{ref} | Linear | 2015.00 | 786.69 | 797.19 | 13.42 | -9.38 | 0.02 | | | | | 11.43 | 0.53 |
| Wheat | MARKSMAN | linear | T[air]^{ref} | Linear | 2016.00 | 746.99 | 757.24 | 14.23 | -48.71 | 0.01 | | | | | 11.86 | 0.54 |
| Wheat | MARKSMAN | linear | T[air]^{ref} | Linear | 2017.00 | 763.77 | 774.34 | 10.44 | -22.33 | 0.01 | | | | | 9.47 | 0.42 |
| Wheat | MARKSMAN | linear | T[air]^{ref} | Linear | 2018.00 | 783.31 | 793.70 | 14.69 | -48.14 | 0.01 | | | | | 12.69 | 0.52 |
| Wheat | MARKSMAN | linear | T[air]^{ref} | Linear | 2019.00 | 672.02 | 681.75 | 15.03 | -47.74 | 0.01 | | | | | 13.27 | 0.48 |
| Wheat | MARKSMAN | linear | T[air]^{ref} | Linear | 2021.00 | 780.08 | 790.50 | 14.23 | -48.62 | 0.01 | | | | | 12.00 | 0.55 |
| Wheat | MARKSMAN | thermal | T[air]^{plot} | Thermal time | 2015.00 | 767.29 | 775.17 | | | | | | | | | |
| Wheat | MARKSMAN | thermal | T[air]^{plot} | Thermal time | 2016.00 | 735.56 | 743.25 | | | | | | | | | |
| Wheat | MARKSMAN | thermal | T[air]^{plot} | Thermal time | 2017.00 | 792.96 | 800.89 | 12.30 | | | | | | | | |
| Wheat | MARKSMAN | thermal | T[air]^{plot} | Thermal time | 2018.00 | 752.08 | 759.86 | | | | | | | | | |
| Wheat | MARKSMAN | thermal | T[air]^{plot} | Thermal time | 2019.00 | 634.23 | 641.52 | | | | | | | | | |
| Wheat | MARKSMAN | thermal | T[air]^{plot} | Thermal time | 2021.00 | 750.32 | 758.13 | | | | | | | | | |
| Wheat | MARKSMAN | thermal | T[air]^{ref} | Thermal time | 2015.00 | 802.55 | 810.42 | 14.33 | | | | | | | | |
| Wheat | MARKSMAN | thermal | T[air]^{ref} | Thermal time | 2016.00 | 763.50 | 771.20 | 14.68 | | | | | | | | |
| Wheat | MARKSMAN | thermal | T[air]^{ref} | Thermal time | 2017.00 | 797.86 | 805.79 | 12.61 | | | | | | | | |
| Wheat | MARKSMAN | thermal | T[air]^{ref} | Thermal time | 2018.00 | 787.19 | 794.97 | 14.65 | | | | | | | | |
| Wheat | MARKSMAN | thermal | T[air]^{ref} | Thermal time | 2019.00 | 665.33 | 672.62 | 13.76 | | | | | | | | |
| Wheat | MARKSMAN | thermal | T[air]^{ref} | Thermal time | 2021.00 | 785.44 | 793.26 | 14.05 | | | | | | | | |
| Wheat | MARKSMAN | wang | T[air]^{plot} | Wang-Engel | 2015.00 | 758.28 | 768.78 | | | | | 16.02 | 0.70 | | 11.57 | 0.41 |
| Wheat | MARKSMAN | wang | T[air]^{plot} | Wang-Engel | 2016.00 | 723.49 | 733.75 | | | | | 15.17 | 0.61 | | 11.82 | 0.44 |
| Wheat | MARKSMAN | wang | T[air]^{plot} | Wang-Engel | 2017.00 | 777.43 | 788.01 | 10.73 | | | | 14.05 | 0.59 | | 10.14 | 0.34 |
| Wheat | MARKSMAN | wang | T[air]^{plot} | Wang-Engel | 2018.00 | 750.47 | 760.85 | | | | | 15.86 | 0.65 | | 12.56 | 0.38 |
| Wheat | MARKSMAN | wang | T[air]^{plot} | Wang-Engel | 2019.00 | 635.51 | 645.23 | | | | | 17.03 | 0.73 | | 12.84 | 0.34 |
| Wheat | MARKSMAN | wang | T[air]^{plot} | Wang-Engel | 2021.00 | 742.92 | 753.34 | | | | | 16.08 | 0.64 | | 11.58 | 0.47 |
| Wheat | MARKSMAN | wang | T[air]^{ref} | Wang-Engel | 2015.00 | 785.58 | 796.08 | 12.65 | | | | 16.02 | 0.70 | | 11.36 | 0.44 |
| Wheat | MARKSMAN | wang | T[air]^{ref} | Wang-Engel | 2016.00 | 741.83 | 752.08 | 13.37 | | | | 15.04 | 0.60 | | 11.51 | 0.47 |
| Wheat | MARKSMAN | wang | T[air]^{ref} | Wang-Engel | 2017.00 | 777.76 | 788.33 | 10.76 | | | | 13.71 | 0.58 | | 10.15 | 0.34 |
| Wheat | MARKSMAN | wang | T[air]^{ref} | Wang-Engel | 2018.00 | 779.42 | 789.80 | 13.64 | | | | 15.91 | 0.66 | | 12.39 | 0.43 |
| Wheat | MARKSMAN | wang | T[air]^{ref} | Wang-Engel | 2019.00 | 662.49 | 672.21 | 13.37 | | | | 17.04 | 0.73 | | 12.48 | 0.37 |

| | | | | | | | | | | | | | | | | | | |
|---|---|---|---|---|---|---|---|---|---|---|---|---|---|---|---|---|---|---|
| Wheat | MARKSMAN | wang | T[air]^{ref} | Wang-Engel | 2021.00 | 770.62 | 781.04 | 13.12 | | | | | | 16.06 | 0.64 | | 11.39 | 0.51 |
| Wheat | OSTKA STRZELECKA | asym | T[air]^{plot} | Asymptotic | 2015.00 | 792.07 | 805.53 | | 3.48 | | | 1.66 | 0.99 | | | | 10.39 | 0.54 |
| Wheat | OSTKA STRZELECKA | asym | T[air]^{plot} | Asymptotic | 2016.00 | 786.84 | 800.06 | | 2.80 | | | 1.21 | 1.01 | | | | 11.77 | 0.62 |
| Wheat | OSTKA STRZELECKA | asym | T[air]^{plot} | Asymptotic | 2017.00 | 868.98 | 882.66 | 12.66 | 5.70 | | | 3.86 | 0.70 | | | | 10.82 | 0.53 |
| Wheat | OSTKA STRZELECKA | asym | T[air]^{plot} | Asymptotic | 2018.00 | 812.48 | 825.80 | | 4.20 | | | 1.78 | 0.93 | | | | 12.85 | 0.52 |
| Wheat | OSTKA STRZELECKA | asym | T[air]^{plot} | Asymptotic | 2019.00 | 792.10 | 805.32 | | 2.80 | | | 0.98 | 1.18 | | | | 12.57 | 0.53 |
| Wheat | OSTKA STRZELECKA | asym | T[air]^{plot} | Asymptotic | 2021.00 | 785.72 | 798.90 | | 2.80 | | | 1.14 | 1.09 | | | | 12.66 | 0.56 |
| Wheat | OSTKA STRZELECKA | asym | T[air]^{ref} | Asymptotic | 2015.00 | 821.27 | 834.73 | 11.80 | 3.44 | | | 1.78 | 0.96 | | | | 10.35 | 0.47 |
| Wheat | OSTKA STRZELECKA | asym | T[air]^{ref} | Asymptotic | 2016.00 | 814.07 | 827.29 | 13.99 | 2.80 | | | 1.45 | 0.92 | | | | 11.99 | 0.53 |
| Wheat | OSTKA STRZELECKA | asym | T[air]^{ref} | Asymptotic | 2017.00 | 871.19 | 884.87 | 12.65 | 5.70 | | | 4.18 | 0.69 | | | | 10.93 | 0.51 |
| Wheat | OSTKA STRZELECKA | asym | T[air]^{ref} | Asymptotic | 2018.00 | 844.87 | 858.18 | 14.21 | 3.30 | | | 1.59 | 0.96 | | | | 12.89 | 0.44 |
| Wheat | OSTKA STRZELECKA | asym | T[air]^{ref} | Asymptotic | 2019.00 | 824.62 | 837.84 | 14.00 | 2.92 | | | 1.20 | 1.08 | | | | 12.62 | 0.45 |
| Wheat | OSTKA STRZELECKA | asym | T[air]^{ref} | Asymptotic | 2021.00 | 818.73 | 831.90 | 14.50 | 2.80 | | | 1.34 | 1.00 | | | | 12.75 | 0.49 |
| Wheat | OSTKA STRZELECKA | bilinear | T[air]^{plot} | Bi-linear | 2015.00 | 797.15 | 810.60 | | 13.06 | 0.02 | 0.83 | | | | | | 10.65 | 0.54 |
| Wheat | OSTKA STRZELECKA | bilinear | T[air]^{plot} | Bi-linear | 2016.00 | 787.66 | 800.88 | | 9.64 | 0.03 | 0.55 | | | | | | 11.81 | 0.59 |
| Wheat | OSTKA STRZELECKA | bilinear | T[air]^{plot} | Bi-linear | 2017.00 | 862.53 | 876.21 | 11.73 | 8.70 | 0.02 | 0.57 | | | | | | 10.52 | 0.46 |
| Wheat | OSTKA STRZELECKA | bilinear | T[air]^{plot} | Bi-linear | 2018.00 | 814.78 | 828.10 | | 12.84 | 0.03 | 0.74 | | | | | | 13.00 | 0.49 |
| Wheat | OSTKA STRZELECKA | bilinear | T[air]^{plot} | Bi-linear | 2019.00 | 793.28 | 806.50 | | 8.95 | 0.04 | 0.52 | | | | | | 12.64 | 0.51 |
| Wheat | OSTKA STRZELECKA | bilinear | T[air]^{plot} | Bi-linear | 2021.00 | 786.00 | 799.17 | | 10.15 | 0.03 | 0.59 | | | | | | 12.69 | 0.55 |
| Wheat | OSTKA STRZELECKA | bilinear | T[air]^{ref} | Bi-linear | 2015.00 | 826.93 | 840.39 | 12.10 | 14.10 | 0.00 | 0.91 | | | | | | 10.62 | 0.47 |
| Wheat | OSTKA STRZELECKA | bilinear | T[air]^{ref} | Bi-linear | 2016.00 | 816.00 | 829.23 | 13.97 | 9.62 | 0.03 | 0.59 | | | | | | 12.11 | 0.51 |
| Wheat | OSTKA STRZELECKA | bilinear | T[air]^{ref} | Bi-linear | 2017.00 | 863.76 | 877.44 | 11.76 | 8.36 | 0.02 | 0.56 | | | | | | 10.57 | 0.45 |
| Wheat | OSTKA STRZELECKA | bilinear | T[air]^{ref} | Bi-linear | 2018.00 | 848.54 | 861.86 | 14.38 | 11.52 | 0.02 | 0.70 | | | | | | 13.12 | 0.42 |
| Wheat | OSTKA STRZELECKA | bilinear | T[air]^{ref} | Bi-linear | 2019.00 | 827.12 | 840.35 | 14.16 | 13.40 | 0.02 | 0.75 | | | | | | 12.78 | 0.45 |
| Wheat | OSTKA STRZELECKA | bilinear | T[air]^{ref} | Bi-linear | 2021.00 | 820.05 | 833.22 | 14.49 | 10.17 | 0.03 | 0.62 | | | | | | 12.83 | 0.48 |
| Wheat | OSTKA STRZELECKA | gauss | ^{plot} | Gaussian | 2015.00 | 1061.42 | 1069.49 | 32.66 | | | | | | | | 55.68 | 32.05 | 0.20 |
| Wheat | OSTKA STRZELECKA | gauss | ^{plot} | Gaussian | 2016.00 | 997.90 | 1005.83 | 29.92 | | | | | | | | 56.35 | 29.85 | 0.11 |
| Wheat | OSTKA STRZELECKA | gauss | ^{plot} | Gaussian | 2017.00 | 1094.91 | 1103.12 | 30.66 | | | | | | | | 56.19 | 29.94 | 0.22 |
| Wheat | OSTKA STRZELECKA | gauss | ^{plot} | Gaussian | 2018.00 | 1038.50 | 1046.49 | 33.90 | | | | | | | | 60.15 | 33.04 | 0.23 |
| Wheat | OSTKA STRZELECKA | gauss | ^{plot} | Gaussian | 2019.00 | 1005.67 | 1013.61 | 31.58 | | | | | | | | 57.99 | 31.00 | 0.20 |
| Wheat | OSTKA STRZELECKA | gauss | ^{plot} | Gaussian | 2021.00 | 1015.37 | 1023.27 | 34.36 | | | | | | | | 59.92 | 34.09 | 0.14 |
| Wheat | OSTKA STRZELECKA | gauss | ^{ref} | Gaussian | 2015.00 | 1061.42 | 1069.49 | 32.66 | | | | | | | | 55.68 | 32.05 | 0.20 |
| Wheat | OSTKA STRZELECKA | gauss | ^{ref} | Gaussian | 2016.00 | 997.90 | 1005.83 | 29.92 | | | | | | | | 56.35 | 29.85 | 0.11 |
| Wheat | OSTKA STRZELECKA | gauss | ^{ref} | Gaussian | 2017.00 | 1094.91 | 1103.12 | 30.66 | | | | | | | | 56.19 | 29.94 | 0.22 |
| Wheat | OSTKA STRZELECKA | gauss | ^{ref} | Gaussian | 2018.00 | 1038.50 | 1046.49 | 33.90 | | | | | | | | 60.15 | 33.04 | 0.23 |
| Wheat | OSTKA STRZELECKA | gauss | ^{ref} | Gaussian | 2019.00 | 1005.67 | 1013.61 | 31.58 | | | | | | | | 57.99 | 31.00 | 0.20 |
| Wheat | OSTKA STRZELECKA | gauss | ^{ref} | Gaussian | 2021.00 | 1015.37 | 1023.27 | 34.36 | | | | | | | | 59.92 | 34.09 | 0.14 |
| Wheat | OSTKA STRZELECKA | linear | T[air]^{plot} | Linear | 2015.00 | 806.23 | 816.99 | | -3.32 | 0.04 | | | | | | | 11.23 | 0.56 |
| Wheat | OSTKA STRZELECKA | linear | T[air]^{plot} | Linear | 2016.00 | 790.75 | 801.32 | | -3.03 | 0.04 | | | | | | | 12.12 | 0.58 |
| Wheat | OSTKA STRZELECKA | linear | T[air]^{plot} | Linear | 2017.00 | | | | | | | | | | | | | |
| Wheat | OSTKA STRZELECKA | linear | T[air]^{plot} | Linear | 2018.00 | 861.73 | 872.39 | | -45.33 | 0.01 | | | | | | | 16.76 | 0.43 |
| Wheat | OSTKA STRZELECKA | linear | T[air]^{plot} | Linear | 2019.00 | 837.33 | 847.90 | | -45.77 | 0.01 | | | | | | | 16.13 | 0.47 |
| Wheat | OSTKA STRZELECKA | linear | T[air]^{plot} | Linear | 2021.00 | 832.11 | 842.65 | | -45.58 | 0.01 | | | | | | | 16.41 | 0.47 |
| Wheat | OSTKA STRZELECKA | linear | T[air]^{ref} | Linear | 2015.00 | 893.77 | 904.54 | 16.75 | -46.69 | 0.01 | | | | | | | 14.79 | 0.49 |
| Wheat | OSTKA STRZELECKA | linear | T[air]^{ref} | Linear | 2016.00 | 821.36 | 831.93 | 14.26 | -4.35 | 0.04 | | | | | | | 12.55 | 0.49 |
| Wheat | OSTKA STRZELECKA | linear | T[air]^{ref} | Linear | 2017.00 | 888.53 | 899.48 | 13.29 | -48.63 | 0.01 | | | | | | | 11.98 | 0.45 |
| Wheat | OSTKA STRZELECKA | linear | T[air]^{ref} | Linear | 2018.00 | 894.88 | 905.54 | 18.21 | -45.37 | 0.01 | | | | | | | 16.71 | 0.43 |
| Wheat | OSTKA STRZELECKA | linear | T[air]^{ref} | Linear | 2019.00 | 829.20 | 839.78 | 14.29 | -2.59 | 0.04 | | | | | | | 13.03 | 0.42 |
| Wheat | OSTKA STRZELECKA | linear | T[air]^{ref} | Linear | 2021.00 | 823.64 | 834.18 | 14.76 | -3.51 | 0.04 | | | | | | | 13.19 | 0.46 |
| Wheat | OSTKA STRZELECKA | thermal | T[air]^{plot} | Thermal time | 2015.00 | 812.53 | 820.60 | | | | | | | | | | | |
| Wheat | OSTKA STRZELECKA | thermal | T[air]^{plot} | Thermal time | 2016.00 | 793.60 | 801.54 | | | | | | | | | | | |
| Wheat | OSTKA STRZELECKA | thermal | T[air]^{plot} | Thermal time | 2017.00 | 878.00 | 886.21 | 13.15 | | | | | | | | | | |

| | | | | | | | | | | | | | | | | | | |
|---|---|---|---|---|---|---|---|---|---|---|---|---|---|---|---|---|---|---|
| Wheat | OSTKA STRZELECKA | thermal | T[air]^{plot} | Thermal time | 2018.00 | 817.47 | 825.46 | | | | | | | | | | | |
| Wheat | OSTKA STRZELECKA | thermal | T[air]^{plot} | Thermal time | 2019.00 | 794.10 | 802.03 | | | | | | | | | | | |
| Wheat | OSTKA STRZELECKA | thermal | T[air]^{plot} | Thermal time | 2021.00 | 789.71 | 797.61 | | | | | | | | | | | |
| Wheat | OSTKA STRZELECKA | thermal | T[air]^{ref} | Thermal time | 2015.00 | 853.13 | 861.20 | 13.74 | | | | | | | | | | |
| Wheat | OSTKA STRZELECKA | thermal | T[air]^{ref} | Thermal time | 2016.00 | 828.28 | 836.21 | 14.96 | | | | | | | | | | |
| Wheat | OSTKA STRZELECKA | thermal | T[air]^{ref} | Thermal time | 2017.00 | 882.16 | 890.37 | 13.39 | | | | | | | | | | |
| Wheat | OSTKA STRZELECKA | thermal | T[air]^{ref} | Thermal time | 2018.00 | 856.67 | 864.66 | 15.31 | | | | | | | | | | |
| Wheat | OSTKA STRZELECKA | thermal | T[air]^{ref} | Thermal time | 2019.00 | 830.48 | 838.41 | 14.44 | | | | | | | | | | |
| Wheat | OSTKA STRZELECKA | thermal | T[air]^{ref} | Thermal time | 2021.00 | 828.40 | 836.30 | 15.27 | | | | | | | | | | |
| Wheat | OSTKA STRZELECKA | wang | T[air]^{plot} | Wang-Engel | 2015.00 | 830.13 | 840.90 | | | | | | | 17.62 | 1.16 | | 12.60 | 0.40 |
| Wheat | OSTKA STRZELECKA | wang | T[air]^{plot} | Wang-Engel | 2016.00 | 829.67 | 840.25 | | | | | | | 18.16 | 1.04 | | 14.72 | 0.57 |
| Wheat | OSTKA STRZELECKA | wang | T[air]^{plot} | Wang-Engel | 2017.00 | 898.49 | 909.43 | 14.15 | | | | | | 17.05 | 0.92 | | 12.44 | 0.50 |
| Wheat | OSTKA STRZELECKA | wang | T[air]^{plot} | Wang-Engel | 2018.00 | 844.45 | 855.10 | | | | | | | 17.94 | 1.07 | | 15.21 | 0.51 |
| Wheat | OSTKA STRZELECKA | wang | T[air]^{plot} | Wang-Engel | 2019.00 | 827.42 | 838.00 | | | | | | | 18.78 | 1.16 | | 15.17 | 0.50 |
| Wheat | OSTKA STRZELECKA | wang | T[air]^{plot} | Wang-Engel | 2021.00 | 826.33 | 836.87 | | | | | | | 18.20 | 1.09 | | 15.74 | 0.50 |
| Wheat | OSTKA STRZELECKA | wang | T[air]^{ref} | Wang-Engel | 2015.00 | 844.90 | 855.66 | 12.61 | | | | | | 17.43 | 1.13 | | 11.65 | 0.37 |
| Wheat | OSTKA STRZELECKA | wang | T[air]^{ref} | Wang-Engel | 2016.00 | 843.72 | 854.30 | 16.32 | | | | | | 17.79 | 1.00 | | 13.98 | 0.53 |
| Wheat | OSTKA STRZELECKA | wang | T[air]^{ref} | Wang-Engel | 2017.00 | 898.11 | 909.05 | 14.01 | | | | | | 16.77 | 0.91 | | 12.42 | 0.48 |
| Wheat | OSTKA STRZELECKA | wang | T[air]^{ref} | Wang-Engel | 2018.00 | 868.67 | 879.32 | 16.31 | | | | | | 17.67 | 1.05 | | 14.58 | 0.47 |
| Wheat | OSTKA STRZELECKA | wang | T[air]^{ref} | Wang-Engel | 2019.00 | 847.19 | 857.76 | 15.79 | | | | | | 18.49 | 1.12 | | 14.22 | 0.45 |
| Wheat | OSTKA STRZELECKA | wang | T[air]^{ref} | Wang-Engel | 2021.00 | 848.78 | 859.32 | 16.56 | | | | | | 17.94 | 1.06 | | 14.92 | 0.45 |
| Wheat | ROMANUS | asym | T[air]^{plot} | Asymptotic | 2015.00 | 778.10 | 791.41 | | 5.13 | | | 2.56 | 0.82 | | | | 10.84 | 0.44 |
| Wheat | ROMANUS | asym | T[air]^{plot} | Asymptotic | 2016.00 | 785.43 | 798.40 | | 6.50 | | | 5.11 | 0.65 | | | | 14.33 | 0.47 |
| Wheat | ROMANUS | asym | T[air]^{plot} | Asymptotic | 2017.00 | 881.59 | 895.14 | 14.22 | 6.80 | | | 8.99 | 0.67 | | | | 12.80 | 0.43 |
| Wheat | ROMANUS | asym | T[air]^{plot} | Asymptotic | 2018.00 | 816.09 | 829.22 | | 5.70 | | | 3.12 | 0.72 | | | | 15.43 | 0.40 |
| Wheat | ROMANUS | asym | T[air]^{plot} | Asymptotic | 2019.00 | 784.78 | 797.70 | | 6.70 | | | 4.43 | 0.69 | | | | 15.59 | 0.42 |
| Wheat | ROMANUS | asym | T[air]^{plot} | Asymptotic | 2021.00 | 816.38 | 829.61 | | 6.50 | | | 3.97 | 0.69 | | | | 14.17 | 0.49 |
| Wheat | ROMANUS | asym | T[air]^{ref} | Asymptotic | 2015.00 | 818.48 | 831.80 | 13.23 | 4.90 | | | 2.45 | 0.85 | | | | 11.37 | 0.51 |
| Wheat | ROMANUS | asym | T[air]^{ref} | Asymptotic | 2016.00 | 815.74 | 828.72 | 18.54 | 6.01 | | | 9.81 | 0.61 | | | | 14.76 | 0.58 |
| Wheat | ROMANUS | asym | T[air]^{ref} | Asymptotic | 2017.00 | 882.47 | 896.02 | 14.01 | 5.80 | | | 4.58 | 0.65 | | | | 12.77 | 0.41 |
| Wheat | ROMANUS | asym | T[air]^{ref} | Asymptotic | 2018.00 | 857.23 | 870.36 | 18.22 | 6.00 | | | 3.74 | 0.70 | | | | 16.04 | 0.47 |
| Wheat | ROMANUS | asym | T[air]^{ref} | Asymptotic | 2019.00 | 823.75 | 836.67 | 17.90 | 5.20 | | | 2.15 | 0.81 | | | | 16.05 | 0.45 |
| Wheat | ROMANUS | asym | T[air]^{ref} | Asymptotic | 2021.00 | 857.98 | 871.20 | 17.87 | 6.50 | | | 4.12 | 0.69 | | | | 14.84 | 0.56 |
| Wheat | ROMANUS | bilinear | T[air]^{plot} | Bi-linear | 2015.00 | 787.46 | 800.77 | | 13.12 | 0.01 | 0.80 | | | | | | 11.36 | 0.45 |
| Wheat | ROMANUS | bilinear | T[air]^{plot} | Bi-linear | 2016.00 | 790.56 | 803.53 | | 12.06 | 0.02 | 0.62 | | | | | | 14.72 | 0.46 |
| Wheat | ROMANUS | bilinear | T[air]^{plot} | Bi-linear | 2017.00 | 886.98 | 900.53 | 13.73 | 8.90 | 0.02 | 0.54 | | | | | | 13.03 | 0.32 |
| Wheat | ROMANUS | bilinear | T[air]^{plot} | Bi-linear | 2018.00 | 819.42 | 832.55 | | 9.07 | 0.03 | 0.52 | | | | | | 15.69 | 0.35 |
| Wheat | ROMANUS | bilinear | T[air]^{plot} | Bi-linear | 2019.00 | 785.56 | 798.48 | | 8.99 | 0.03 | 0.49 | | | | | | 15.65 | 0.35 |
| Wheat | ROMANUS | bilinear | T[air]^{plot} | Bi-linear | 2021.00 | 820.59 | 833.81 | | 10.81 | 0.03 | 0.59 | | | | | | 14.51 | 0.44 |
| Wheat | ROMANUS | bilinear | T[air]^{ref} | Bi-linear | 2015.00 | 826.32 | 839.64 | 13.58 | 13.95 | 0.00 | 0.87 | | | | | | 11.80 | 0.49 |
| Wheat | ROMANUS | bilinear | T[air]^{ref} | Bi-linear | 2016.00 | 822.39 | 835.37 | 18.39 | 12.19 | 0.01 | 0.65 | | | | | | 15.27 | 0.54 |
| Wheat | ROMANUS | bilinear | T[air]^{ref} | Bi-linear | 2017.00 | 887.98 | 901.53 | 13.81 | 8.97 | 0.02 | 0.56 | | | | | | 13.09 | 0.32 |
| Wheat | ROMANUS | bilinear | T[air]^{ref} | Bi-linear | 2018.00 | 860.36 | 873.49 | 17.72 | 12.22 | 0.02 | 0.68 | | | | | | 16.29 | 0.40 |
| Wheat | ROMANUS | bilinear | T[air]^{ref} | Bi-linear | 2019.00 | 826.28 | 839.20 | 17.64 | 12.11 | 0.03 | 0.65 | | | | | | 16.26 | 0.40 |
| Wheat | ROMANUS | bilinear | T[air]^{ref} | Bi-linear | 2021.00 | 861.50 | 874.72 | 17.25 | 12.23 | 0.02 | 0.67 | | | | | | 15.10 | 0.49 |
| Wheat | ROMANUS | gauss | ^{plot} | Gaussian | 2015.00 | 1023.59 | 1031.58 | 31.88 | | | | | | | | 53.58 | 30.78 | 0.27 |
| Wheat | ROMANUS | gauss | ^{plot} | Gaussian | 2016.00 | 937.02 | 944.80 | 28.63 | | | | | | | | 54.03 | 27.97 | 0.22 |
| Wheat | ROMANUS | gauss | ^{plot} | Gaussian | 2017.00 | 1049.57 | 1057.70 | 28.61 | | | | | | | | 53.98 | 27.79 | 0.24 |
| Wheat | ROMANUS | gauss | ^{plot} | Gaussian | 2018.00 | 991.54 | 999.42 | 32.82 | | | | | | | | 58.05 | 31.82 | 0.25 |
| Wheat | ROMANUS | gauss | ^{plot} | Gaussian | 2019.00 | 938.71 | 946.46 | 30.90 | | | | | | | | 56.69 | 29.63 | 0.29 |
| Wheat | ROMANUS | gauss | ^{plot} | Gaussian | 2021.00 | 1005.23 | 1013.17 | 31.69 | | | | | | | | 57.79 | 30.93 | 0.22 |
| Wheat | ROMANUS | gauss | ^{ref} | Gaussian | 2015.00 | 1023.59 | 1031.58 | 31.88 | | | | | | | | 53.58 | 30.78 | 0.27 |

| | | | | | | | | | | | | | | | | | |
|---|---|---|---|---|---|---|---|---|---|---|---|---|---|---|---|---|---|
| Wheat | ROMANUS | gauss | ^{ref} | Gaussian | 2016.00 | 937.02 | 944.80 | 28.63 | | | | | | | 54.03 | 27.97 | 0.22 |
| Wheat | ROMANUS | gauss | ^{ref} | Gaussian | 2017.00 | 1049.57 | 1057.70 | 28.61 | | | | | | | 53.98 | 27.79 | 0.24 |
| Wheat | ROMANUS | gauss | ^{ref} | Gaussian | 2018.00 | 991.54 | 999.42 | 32.82 | | | | | | | 58.05 | 31.82 | 0.25 |
| Wheat | ROMANUS | gauss | ^{ref} | Gaussian | 2019.00 | 938.71 | 946.46 | 30.90 | | | | | | | 56.69 | 29.63 | 0.29 |
| Wheat | ROMANUS | gauss | ^{ref} | Gaussian | 2021.00 | 1005.23 | 1013.17 | 31.69 | | | | | | | 57.79 | 30.93 | 0.22 |
| Wheat | ROMANUS | linear | T[air]^{plot} | Linear | 2015.00 | 841.36 | 852.01 | | -46.46 | 0.01 | | | | | | 15.13 | 0.48 |
| Wheat | ROMANUS | linear | T[air]^{plot} | Linear | 2016.00 | 813.46 | 823.84 | | -45.24 | 0.01 | | | | | | 16.90 | 0.50 |
| Wheat | ROMANUS | linear | T[air]^{plot} | Linear | 2017.00 | 905.14 | 915.97 | 15.62 | -47.00 | 0.01 | | | | | | 14.35 | 0.41 |
| Wheat | ROMANUS | linear | T[air]^{plot} | Linear | 2018.00 | 849.18 | 859.68 | | -44.03 | 0.01 | | | | | | 18.65 | 0.42 |
| Wheat | ROMANUS | linear | T[air]^{plot} | Linear | 2019.00 | 811.53 | 821.87 | | -44.25 | 0.01 | | | | | | 18.34 | 0.42 |
| Wheat | ROMANUS | linear | T[air]^{plot} | Linear | 2021.00 | 852.24 | 862.82 | | -44.93 | 0.01 | | | | | | 17.35 | 0.48 |
| Wheat | ROMANUS | linear | T[air]^{ref} | Linear | 2015.00 | 879.85 | 890.51 | 18.16 | -46.18 | 0.01 | | | | | | 15.54 | 0.54 |
| Wheat | ROMANUS | linear | T[air]^{ref} | Linear | 2016.00 | 843.54 | 853.92 | 21.14 | -44.97 | 0.01 | | | | | | 17.29 | 0.56 |
| Wheat | ROMANUS | linear | T[air]^{ref} | Linear | 2017.00 | 905.31 | 916.15 | 15.63 | -46.99 | 0.01 | | | | | | 14.36 | 0.41 |
| Wheat | ROMANUS | linear | T[air]^{ref} | Linear | 2018.00 | 888.67 | 899.17 | 21.54 | -43.72 | 0.01 | | | | | | 19.10 | 0.48 |
| Wheat | ROMANUS | linear | T[air]^{ref} | Linear | 2019.00 | 850.96 | 861.30 | 21.23 | -43.93 | 0.01 | | | | | | 18.81 | 0.48 |
| Wheat | ROMANUS | linear | T[air]^{ref} | Linear | 2021.00 | 891.78 | 902.36 | 20.86 | -44.61 | 0.01 | | | | | | 17.81 | 0.54 |
| Wheat | ROMANUS | thermal | T[air]^{plot} | Thermal time | 2015.00 | 797.57 | 805.56 | | | | | | | | | | |
| Wheat | ROMANUS | thermal | T[air]^{plot} | Thermal time | 2016.00 | 792.33 | 800.12 | | | | | | | | | | |
| Wheat | ROMANUS | thermal | T[air]^{plot} | Thermal time | 2017.00 | 893.69 | 901.82 | 14.51 | | | | | | | | | |
| Wheat | ROMANUS | thermal | T[air]^{plot} | Thermal time | 2018.00 | 818.37 | 826.25 | | | | | | | | | | |
| Wheat | ROMANUS | thermal | T[air]^{plot} | Thermal time | 2019.00 | 783.49 | 791.24 | | | | | | | | | | |
| Wheat | ROMANUS | thermal | T[air]^{plot} | Thermal time | 2021.00 | 821.53 | 829.46 | | | | | | | | | | |
| Wheat | ROMANUS | thermal | T[air]^{ref} | Thermal time | 2015.00 | 842.25 | 850.24 | 14.41 | | | | | | | | | |
| Wheat | ROMANUS | thermal | T[air]^{ref} | Thermal time | 2016.00 | 828.40 | 836.18 | 18.08 | | | | | | | | | |
| Wheat | ROMANUS | thermal | T[air]^{ref} | Thermal time | 2017.00 | 897.08 | 905.20 | 14.76 | | | | | | | | | |
| Wheat | ROMANUS | thermal | T[air]^{ref} | Thermal time | 2018.00 | 860.87 | 868.75 | 17.87 | | | | | | | | | |
| Wheat | ROMANUS | thermal | T[air]^{ref} | Thermal time | 2019.00 | 824.57 | 832.33 | 17.50 | | | | | | | | | |
| Wheat | ROMANUS | thermal | T[air]^{ref} | Thermal time | 2021.00 | 864.59 | 872.52 | 17.25 | | | | | | | | | |
| Wheat | ROMANUS | wang | T[air]^{plot} | Wang-Engel | 2015.00 | 803.30 | 813.96 | | | | | | 17.35 | 1.06 | | 12.41 | 0.40 |
| Wheat | ROMANUS | wang | T[air]^{plot} | Wang-Engel | 2016.00 | 808.75 | 819.13 | | | | | | 17.24 | 0.82 | | 16.37 | 0.55 |
| Wheat | ROMANUS | wang | T[air]^{plot} | Wang-Engel | 2017.00 | 910.21 | 921.05 | 16.06 | | | | | 16.46 | 0.81 | | 14.61 | 0.42 |
| Wheat | ROMANUS | wang | T[air]^{plot} | Wang-Engel | 2018.00 | 838.46 | 848.96 | | | | | | 17.59 | 0.95 | | 17.49 | 0.41 |
| Wheat | ROMANUS | wang | T[air]^{plot} | Wang-Engel | 2019.00 | 804.02 | 814.36 | | | | | | 18.12 | 0.97 | | 17.47 | 0.42 |
| Wheat | ROMANUS | wang | T[air]^{plot} | Wang-Engel | 2021.00 | 840.17 | 850.75 | | | | | | 17.61 | 0.91 | | 16.18 | 0.49 |
| Wheat | ROMANUS | wang | T[air]^{ref} | Wang-Engel | 2015.00 | 836.95 | 847.60 | 13.92 | | | | | 17.25 | 1.06 | | 12.53 | 0.44 |
| Wheat | ROMANUS | wang | T[air]^{ref} | Wang-Engel | 2016.00 | 833.72 | 844.10 | 21.12 | | | | | 16.80 | 0.78 | | 16.35 | 0.61 |
| Wheat | ROMANUS | wang | T[air]^{ref} | Wang-Engel | 2017.00 | 910.38 | 921.22 | 16.05 | | | | | 16.15 | 0.80 | | 14.63 | 0.42 |
| Wheat | ROMANUS | wang | T[air]^{ref} | Wang-Engel | 2018.00 | 875.12 | 885.62 | 19.64 | | | | | 17.52 | 0.95 | | 17.70 | 0.44 |
| Wheat | ROMANUS | wang | T[air]^{ref} | Wang-Engel | 2019.00 | 839.80 | 850.14 | 19.54 | | | | | 18.06 | 0.98 | | 17.61 | 0.44 |
| Wheat | ROMANUS | wang | T[air]^{ref} | Wang-Engel | 2021.00 | 876.66 | 887.23 | 19.21 | | | | | 17.48 | 0.91 | | 16.41 | 0.53 |
| Wheat | RUNAL | asym | T[air]^{plot} | Asymptotic | 2015.00 | 853.02 | 866.57 | | 2.80 | | 2.03 | 0.66 | | | | 12.90 | 0.55 |
| Wheat | RUNAL | asym | T[air]^{plot} | Asymptotic | 2016.00 | 809.47 | 822.69 | | 5.20 | | 7.06 | 0.55 | | | | 13.72 | 0.61 |
| Wheat | RUNAL | asym | T[air]^{plot} | Asymptotic | 2017.00 | 762.11 | 775.38 | 10.74 | 3.59 | | 6.39 | 0.54 | | | | 9.05 | 0.57 |
| Wheat | RUNAL | asym | T[air]^{plot} | Asymptotic | 2018.00 | 851.64 | 865.00 | | 7.90 | | 10.86 | 0.66 | | | | 15.00 | 0.51 |
| Wheat | RUNAL | asym | T[air]^{plot} | Asymptotic | 2019.00 | 764.10 | 777.03 | | 2.60 | | 1.20 | 0.86 | | | | 13.94 | 0.57 |
| Wheat | RUNAL | asym | T[air]^{plot} | Asymptotic | 2021.00 | 820.89 | 834.16 | | 5.40 | | 5.70 | 0.58 | | | | 13.95 | 0.59 |
| Wheat | RUNAL | asym | T[air]^{ref} | Asymptotic | 2015.00 | 881.71 | 895.26 | 15.37 | 5.12 | | 7.94 | 0.58 | | | | 12.88 | 0.55 |
| Wheat | RUNAL | asym | T[air]^{ref} | Asymptotic | 2016.00 | 839.55 | 852.77 | 17.07 | 5.12 | | 11.36 | 0.56 | | | | 13.57 | 0.61 |
| Wheat | RUNAL | asym | T[air]^{ref} | Asymptotic | 2017.00 | 762.53 | 775.80 | 10.77 | 3.34 | | 8.29 | 0.54 | | | | 9.12 | 0.56 |
| Wheat | RUNAL | asym | T[air]^{ref} | Asymptotic | 2018.00 | 862.86 | 876.22 | 16.42 | 5.10 | | 6.36 | 0.58 | | | | 13.52 | 0.58 |
| Wheat | RUNAL | asym | T[air]^{ref} | Asymptotic | 2019.00 | 806.88 | 819.81 | 17.61 | 7.86 | | 4.05 | 0.65 | | | | 14.71 | 0.56 |

| | | | | | | | | | | | | | | | | | | |
|---|---|---|---|---|---|---|---|---|---|---|---|---|---|---|---|---|---|---|
| Wheat | RUNAL | asym | T[air]^{ref} | Asymptotic | 2021.00 | 852.05 | 865.32 | 17.04 | 5.10 | | | 5.14 | 0.58 | | | | 13.85 | 0.59 |
| Wheat | RUNAL | bilinear | T[air]^{plot} | Bi-linear | 2015.00 | 852.09 | 865.63 | | 8.58 | 0.01 | 0.51 | | | | | | 12.85 | 0.57 |
| Wheat | RUNAL | bilinear | T[air]^{plot} | Bi-linear | 2016.00 | 808.34 | 821.56 | | 7.44 | 0.02 | 0.45 | | | | | | 13.64 | 0.59 |
| Wheat | RUNAL | bilinear | T[air]^{plot} | Bi-linear | 2017.00 | 751.46 | 764.73 | 10.32 | 5.99 | 0.01 | 0.50 | | | | | | 8.55 | 0.57 |
| Wheat | RUNAL | bilinear | T[air]^{plot} | Bi-linear | 2018.00 | 832.50 | 845.86 | | 8.75 | 0.01 | 0.51 | | | | | | 13.64 | 0.57 |
| Wheat | RUNAL | bilinear | T[air]^{plot} | Bi-linear | 2019.00 | 764.01 | 776.94 | | 8.30 | 0.03 | 0.42 | | | | | | 13.94 | 0.58 |
| Wheat | RUNAL | bilinear | T[air]^{plot} | Bi-linear | 2021.00 | 820.03 | 833.30 | | 8.62 | 0.02 | 0.49 | | | | | | 13.89 | 0.59 |
| Wheat | RUNAL | bilinear | T[air]^{ref} | Bi-linear | 2015.00 | 882.87 | 896.42 | 15.54 | 11.10 | 0.00 | 0.63 | | | | | | 12.79 | 0.57 |
| Wheat | RUNAL | bilinear | T[air]^{ref} | Bi-linear | 2016.00 | 840.94 | 854.16 | 16.99 | 10.86 | 0.01 | 0.58 | | | | | | 13.66 | 0.59 |
| Wheat | RUNAL | bilinear | T[air]^{ref} | Bi-linear | 2017.00 | 753.91 | 767.18 | 10.40 | 5.89 | 0.01 | 0.50 | | | | | | 8.65 | 0.57 |
| Wheat | RUNAL | bilinear | T[air]^{ref} | Bi-linear | 2018.00 | 864.22 | 877.59 | 16.58 | 11.07 | 0.00 | 0.61 | | | | | | 13.60 | 0.58 |
| Wheat | RUNAL | bilinear | T[air]^{ref} | Bi-linear | 2019.00 | 794.71 | 807.64 | 16.65 | 11.10 | 0.02 | 0.55 | | | | | | 13.81 | 0.57 |
| Wheat | RUNAL | bilinear | T[air]^{ref} | Bi-linear | 2021.00 | 852.36 | 865.63 | 16.94 | 10.37 | 0.01 | 0.57 | | | | | | 13.90 | 0.58 |
| Wheat | RUNAL | gauss | ^{plot} | Gaussian | 2015.00 | 1041.17 | 1049.30 | 29.23 | | | | | | | | 45.67 | 26.75 | 0.41 |
| Wheat | RUNAL | gauss | ^{plot} | Gaussian | 2016.00 | 984.87 | 992.80 | 29.16 | | | | | | | | 45.90 | 28.02 | 0.29 |
| Wheat | RUNAL | gauss | ^{plot} | Gaussian | 2017.00 | 985.03 | 993.00 | 27.32 | | | | | | | | 46.46 | 26.79 | 0.21 |
| Wheat | RUNAL | gauss | ^{plot} | Gaussian | 2018.00 | 1025.84 | 1033.86 | 31.20 | | | | | | | | 49.17 | 29.71 | 0.32 |
| Wheat | RUNAL | gauss | ^{plot} | Gaussian | 2019.00 | 927.71 | 935.46 | 29.43 | | | | | | | | 47.94 | 28.00 | 0.32 |
| Wheat | RUNAL | gauss | ^{plot} | Gaussian | 2021.00 | 1011.43 | 1019.39 | 31.76 | | | | | | | | 49.29 | 30.41 | 0.30 |
| Wheat | RUNAL | gauss | ^{ref} | Gaussian | 2015.00 | 1041.17 | 1049.30 | 29.23 | | | | | | | | 45.67 | 26.75 | 0.41 |
| Wheat | RUNAL | gauss | ^{ref} | Gaussian | 2016.00 | 984.87 | 992.80 | 29.16 | | | | | | | | 45.90 | 28.02 | 0.29 |
| Wheat | RUNAL | gauss | ^{ref} | Gaussian | 2017.00 | 985.03 | 993.00 | 27.32 | | | | | | | | 46.46 | 26.79 | 0.21 |
| Wheat | RUNAL | gauss | ^{ref} | Gaussian | 2018.00 | 1025.84 | 1033.86 | 31.20 | | | | | | | | 49.17 | 29.71 | 0.32 |
| Wheat | RUNAL | gauss | ^{ref} | Gaussian | 2019.00 | 927.71 | 935.46 | 29.43 | | | | | | | | 47.94 | 28.00 | 0.32 |
| Wheat | RUNAL | gauss | ^{ref} | Gaussian | 2021.00 | 1011.43 | 1019.39 | 31.76 | | | | | | | | 49.29 | 30.41 | 0.30 |
| Wheat | RUNAL | linear | T[air]^{plot} | Linear | 2015.00 | 871.27 | 882.11 | | -47.05 | 0.01 | | | | | | | 14.28 | 0.63 |
| Wheat | RUNAL | linear | T[air]^{plot} | Linear | 2016.00 | 822.72 | 833.30 | | -46.62 | 0.01 | | | | | | | 14.89 | 0.64 |
| Wheat | RUNAL | linear | T[air]^{plot} | Linear | 2017.00 | 742.10 | 752.71 | 10.32 | -51.19 | 0.01 | | | | | | | 8.26 | 0.62 |
| Wheat | RUNAL | linear | T[air]^{plot} | Linear | 2018.00 | 849.65 | 860.35 | | -46.50 | 0.01 | | | | | | | 15.07 | 0.62 |
| Wheat | RUNAL | linear | T[air]^{plot} | Linear | 2019.00 | 786.14 | 796.48 | | -45.88 | 0.01 | | | | | | | 15.98 | 0.63 |
| Wheat | RUNAL | linear | T[air]^{plot} | Linear | 2021.00 | 837.86 | 848.48 | | -46.25 | 0.01 | | | | | | | 15.43 | 0.62 |
| Wheat | RUNAL | linear | T[air]^{ref} | Linear | 2015.00 | 903.56 | 914.40 | 18.00 | -47.06 | 0.01 | | | | | | | 14.26 | 0.62 |
| Wheat | RUNAL | linear | T[air]^{ref} | Linear | 2016.00 | 855.39 | 865.96 | 19.00 | -46.64 | 0.01 | | | | | | | 14.87 | 0.63 |
| Wheat | RUNAL | linear | T[air]^{ref} | Linear | 2017.00 | 742.52 | 753.13 | 10.33 | -51.18 | 0.01 | | | | | | | 8.28 | 0.61 |
| Wheat | RUNAL | linear | T[air]^{ref} | Linear | 2018.00 | 882.00 | 892.69 | 18.84 | -46.54 | 0.01 | | | | | | | 15.02 | 0.62 |
| Wheat | RUNAL | linear | T[air]^{ref} | Linear | 2019.00 | 818.62 | 828.96 | 20.09 | -45.93 | 0.01 | | | | | | | 15.89 | 0.63 |
| Wheat | RUNAL | linear | T[air]^{ref} | Linear | 2021.00 | 870.27 | 880.89 | 19.32 | -46.29 | 0.01 | | | | | | | 15.37 | 0.62 |
| Wheat | RUNAL | thermal | T[air]^{plot} | Thermal time | 2015.00 | 862.30 | 870.43 | | | | | | | | | | | |
| Wheat | RUNAL | thermal | T[air]^{plot} | Thermal time | 2016.00 | 815.54 | 823.47 | | | | | | | | | | | |
| Wheat | RUNAL | thermal | T[air]^{plot} | Thermal time | 2017.00 | 800.93 | 808.89 | 12.58 | | | | | | | | | | |
| Wheat | RUNAL | thermal | T[air]^{plot} | Thermal time | 2018.00 | 841.76 | 849.78 | | | | | | | | | | | |
| Wheat | RUNAL | thermal | T[air]^{plot} | Thermal time | 2019.00 | 762.90 | 770.66 | | | | | | | | | | | |
| Wheat | RUNAL | thermal | T[air]^{plot} | Thermal time | 2021.00 | 824.94 | 832.90 | | | | | | | | | | | |
| Wheat | RUNAL | thermal | T[air]^{ref} | Thermal time | 2015.00 | 898.27 | 906.40 | 15.68 | | | | | | | | | | |
| Wheat | RUNAL | thermal | T[air]^{ref} | Thermal time | 2016.00 | 851.78 | 859.72 | 16.89 | | | | | | | | | | |
| Wheat | RUNAL | thermal | T[air]^{ref} | Thermal time | 2017.00 | 806.88 | 814.84 | 12.81 | | | | | | | | | | |
| Wheat | RUNAL | thermal | T[air]^{ref} | Thermal time | 2018.00 | 877.84 | 885.85 | 16.63 | | | | | | | | | | |
| Wheat | RUNAL | thermal | T[air]^{ref} | Thermal time | 2019.00 | 795.43 | 803.18 | 16.33 | | | | | | | | | | |
| Wheat | RUNAL | thermal | T[air]^{ref} | Thermal time | 2021.00 | 861.14 | 869.10 | 17.00 | | | | | | | | | | |
| Wheat | RUNAL | wang | T[air]^{plot} | Wang-Engel | 2015.00 | 860.41 | 871.25 | | | | | | | 15.92 | 0.74 | | 13.49 | 0.54 |
| Wheat | RUNAL | wang | T[air]^{plot} | Wang-Engel | 2016.00 | 820.24 | 830.82 | | | | | | | 16.08 | 0.70 | | 14.63 | 0.60 |
| Wheat | RUNAL | wang | T[air]^{plot} | Wang-Engel | 2017.00 | 791.88 | 802.50 | 12.71 | | | | | | 13.49 | 0.63 | | 10.48 | 0.60 |

| | | | | | | | | | | | | | | | | | | |
|---|---|---|---|---|---|---|---|---|---|---|---|---|---|---|---|---|---|---|
| Wheat | RUNAL | wang | T[air]^{plot} | Wang-Engel | 2018.00 | 840.41 | 851.10 | | | | | | | 15.91 | 0.71 | | 14.32 | 0.58 |
| Wheat | RUNAL | wang | T[air]^{plot} | Wang-Engel | 2019.00 | 776.28 | 786.62 | | | | | | | 17.53 | 0.79 | | 15.05 | 0.56 |
| Wheat | RUNAL | wang | T[air]^{plot} | Wang-Engel | 2021.00 | 833.73 | 844.35 | | | | | | | 16.22 | 0.72 | | 15.02 | 0.58 |
| Wheat | RUNAL | wang | T[air]^{ref} | Wang-Engel | 2015.00 | 885.53 | 896.37 | 16.01 | | | | | | 15.92 | 0.74 | | 13.06 | 0.58 |
| Wheat | RUNAL | wang | T[air]^{ref} | Wang-Engel | 2016.00 | 846.82 | 857.40 | 18.17 | | | | | | 16.00 | 0.70 | | 14.20 | 0.63 |
| Wheat | RUNAL | wang | T[air]^{ref} | Wang-Engel | 2017.00 | 791.69 | 802.31 | 12.70 | | | | | | 13.40 | 0.63 | | 10.47 | 0.60 |
| Wheat | RUNAL | wang | T[air]^{ref} | Wang-Engel | 2018.00 | 867.94 | 878.63 | 17.56 | | | | | | 15.84 | 0.71 | | 13.98 | 0.62 |
| Wheat | RUNAL | wang | T[air]^{ref} | Wang-Engel | 2019.00 | 799.92 | 810.26 | 17.55 | | | | | | 17.47 | 0.79 | | 14.34 | 0.59 |
| Wheat | RUNAL | wang | T[air]^{ref} | Wang-Engel | 2021.00 | 860.64 | 871.26 | 18.12 | | | | | | 16.11 | 0.72 | | 14.59 | 0.61 |
| Wheat | RYWALKA | asym | T[air]^{plot} | Asymptotic | 2015.00 | 847.25 | 860.97 | | 4.38 | | | 1.73 | 0.95 | | | | 10.88 | 0.58 |
| Wheat | RYWALKA | asym | T[air]^{plot} | Asymptotic | 2016.00 | 835.98 | 849.39 | | 6.74 | | | 5.21 | 0.67 | | | | 12.83 | 0.50 |
| Wheat | RYWALKA | asym | T[air]^{plot} | Asymptotic | 2017.00 | 942.73 | 956.71 | 13.85 | 6.21 | | | 10.91 | 0.64 | | | | 11.79 | 0.54 |
| Wheat | RYWALKA | asym | T[air]^{plot} | Asymptotic | 2018.00 | 862.35 | 875.94 | | 3.54 | | | 1.21 | 1.05 | | | | 12.99 | 0.54 |
| Wheat | RYWALKA | asym | T[air]^{plot} | Asymptotic | 2019.00 | 795.82 | 809.04 | | 2.80 | | | 0.57 | 1.46 | | | | 12.80 | 0.51 |
| Wheat | RYWALKA | asym | T[air]^{plot} | Asymptotic | 2021.00 | 839.54 | 853.04 | | 3.33 | | | 1.03 | 1.12 | | | | 12.57 | 0.58 |
| Wheat | RYWALKA | asym | T[air]^{ref} | Asymptotic | 2015.00 | 875.56 | 889.28 | 12.89 | 4.30 | | | 1.89 | 0.90 | | | | 10.78 | 0.55 |
| Wheat | RYWALKA | asym | T[air]^{ref} | Asymptotic | 2016.00 | 853.71 | 867.12 | 14.32 | 3.79 | | | 1.56 | 0.88 | | | | 12.47 | 0.50 |
| Wheat | RYWALKA | asym | T[air]^{ref} | Asymptotic | 2017.00 | 942.71 | 956.69 | 13.83 | 6.10 | | | 5.42 | 0.65 | | | | 11.79 | 0.53 |
| Wheat | RYWALKA | asym | T[air]^{ref} | Asymptotic | 2018.00 | 894.92 | 908.51 | 14.88 | 3.56 | | | 1.47 | 0.94 | | | | 13.03 | 0.50 |
| Wheat | RYWALKA | asym | T[air]^{ref} | Asymptotic | 2019.00 | 829.23 | 842.45 | 14.35 | 2.80 | | | 0.84 | 1.24 | | | | 12.91 | 0.45 |
| Wheat | RYWALKA | asym | T[air]^{ref} | Asymptotic | 2021.00 | 871.94 | 885.44 | 14.78 | 3.35 | | | 1.30 | 0.99 | | | | 12.61 | 0.54 |
| Wheat | RYWALKA | bilinear | T[air]^{plot} | Bi-linear | 2015.00 | 855.27 | 868.99 | | 14.30 | 0.02 | 0.80 | | | | | | 11.28 | 0.56 |
| Wheat | RYWALKA | bilinear | T[air]^{plot} | Bi-linear | 2016.00 | 831.43 | 844.84 | | 10.61 | 0.03 | 0.53 | | | | | | 12.56 | 0.51 |
| Wheat | RYWALKA | bilinear | T[air]^{plot} | Bi-linear | 2017.00 | 932.41 | 946.39 | 12.81 | 9.64 | 0.03 | 0.52 | | | | | | 11.29 | 0.49 |
| Wheat | RYWALKA | bilinear | T[air]^{plot} | Bi-linear | 2018.00 | 866.54 | 880.13 | | 14.20 | 0.03 | 0.72 | | | | | | 13.24 | 0.51 |
| Wheat | RYWALKA | bilinear | T[air]^{plot} | Bi-linear | 2019.00 | 797.69 | 810.91 | | 10.93 | 0.04 | 0.54 | | | | | | 12.92 | 0.50 |
| Wheat | RYWALKA | bilinear | T[air]^{plot} | Bi-linear | 2021.00 | 842.86 | 856.36 | | 11.39 | 0.04 | 0.58 | | | | | | 12.77 | 0.55 |
| Wheat | RYWALKA | bilinear | T[air]^{ref} | Bi-linear | 2015.00 | 884.46 | 898.18 | 13.16 | 15.10 | -0.00 | 0.88 | | | | | | 11.20 | 0.53 |
| Wheat | RYWALKA | bilinear | T[air]^{ref} | Bi-linear | 2016.00 | 859.05 | 872.46 | 14.49 | 11.24 | 0.03 | 0.59 | | | | | | 12.79 | 0.48 |
| Wheat | RYWALKA | bilinear | T[air]^{ref} | Bi-linear | 2017.00 | 933.73 | 947.71 | 12.92 | 11.64 | 0.03 | 0.61 | | | | | | 11.36 | 0.49 |
| Wheat | RYWALKA | bilinear | T[air]^{ref} | Bi-linear | 2018.00 | 899.89 | 913.48 | 15.07 | 14.56 | 0.01 | 0.77 | | | | | | 13.33 | 0.48 |
| Wheat | RYWALKA | bilinear | T[air]^{ref} | Bi-linear | 2019.00 | 831.45 | 844.67 | 14.47 | 13.94 | 0.03 | 0.71 | | | | | | 13.05 | 0.45 |
| Wheat | RYWALKA | bilinear | T[air]^{ref} | Bi-linear | 2021.00 | 876.86 | 890.36 | 14.87 | 11.28 | 0.03 | 0.60 | | | | | | 12.91 | 0.51 |
| Wheat | RYWALKA | gauss | ^{plot} | Gaussian | 2015.00 | 1100.30 | 1108.53 | 30.68 | | | | | | | | 51.00 | 29.39 | 0.30 |
| Wheat | RYWALKA | gauss | ^{plot} | Gaussian | 2016.00 | 1023.60 | 1031.64 | 28.60 | | | | | | | | 51.06 | 28.11 | 0.20 |
| Wheat | RYWALKA | gauss | ^{plot} | Gaussian | 2017.00 | 1149.27 | 1157.65 | 29.45 | | | | | | | | 52.25 | 28.35 | 0.28 |
| Wheat | RYWALKA | gauss | ^{plot} | Gaussian | 2018.00 | 1082.29 | 1090.44 | 32.11 | | | | | | | | 54.82 | 30.85 | 0.29 |
| Wheat | RYWALKA | gauss | ^{plot} | Gaussian | 2019.00 | 997.08 | 1005.02 | 30.77 | | | | | | | | 55.28 | 29.73 | 0.27 |
| Wheat | RYWALKA | gauss | ^{plot} | Gaussian | 2021.00 | 1066.57 | 1074.67 | 32.35 | | | | | | | | 54.86 | 31.38 | 0.25 |
| Wheat | RYWALKA | gauss | ^{ref} | Gaussian | 2015.00 | 1100.30 | 1108.53 | 30.68 | | | | | | | | 51.00 | 29.39 | 0.30 |
| Wheat | RYWALKA | gauss | ^{ref} | Gaussian | 2016.00 | 1023.60 | 1031.64 | 28.60 | | | | | | | | 51.06 | 28.11 | 0.20 |
| Wheat | RYWALKA | gauss | ^{ref} | Gaussian | 2017.00 | 1149.27 | 1157.65 | 29.45 | | | | | | | | 52.25 | 28.35 | 0.28 |
| Wheat | RYWALKA | gauss | ^{ref} | Gaussian | 2018.00 | 1082.29 | 1090.44 | 32.11 | | | | | | | | 54.82 | 30.85 | 0.29 |
| Wheat | RYWALKA | gauss | ^{ref} | Gaussian | 2019.00 | 997.08 | 1005.02 | 30.77 | | | | | | | | 55.28 | 29.73 | 0.27 |
| Wheat | RYWALKA | gauss | ^{ref} | Gaussian | 2021.00 | 1066.57 | 1074.67 | 32.35 | | | | | | | | 54.86 | 31.38 | 0.25 |
| Wheat | RYWALKA | linear | T[air]^{plot} | Linear | 2015.00 | 919.63 | 930.61 | | -46.26 | 0.01 | | | | | | | 15.42 | 0.51 |
| Wheat | RYWALKA | linear | T[air]^{plot} | Linear | 2016.00 | 873.97 | 884.70 | | -46.07 | 0.01 | | | | | | | 15.69 | 0.49 |
| Wheat | RYWALKA | linear | T[air]^{plot} | Linear | 2017.00 | 965.62 | 976.80 | 14.75 | -47.81 | 0.01 | | | | | | | 13.16 | 0.47 |
| Wheat | RYWALKA | linear | T[air]^{plot} | Linear | 2018.00 | 912.99 | 923.87 | | -45.22 | 0.01 | | | | | | | 16.75 | 0.49 |
| Wheat | RYWALKA | linear | T[air]^{plot} | Linear | 2019.00 | 843.22 | 853.80 | | -45.44 | 0.01 | | | | | | | 16.60 | 0.48 |
| Wheat | RYWALKA | linear | T[air]^{plot} | Linear | 2021.00 | 842.74 | 853.55 | | -0.65 | 0.04 | | | | | | | 12.88 | 0.55 |
| Wheat | RYWALKA | linear | T[air]^{ref} | Linear | 2015.00 | 951.78 | 962.76 | 17.68 | -46.31 | 0.01 | | | | | | | 15.35 | 0.52 |

| | | | | | | | | | | | | | | | | | |
|---|---|---|---|---|---|---|---|---|---|---|---|---|---|---|---|---|---|
| Wheat | RYWALKA | linear | T[air]^{ref} | Linear | 2016.00 | 899.64 | 910.37 | 17.99 | -46.04 | 0.01 | | | | | | 15.74 | 0.50 |
| Wheat | RYWALKA | linear | T[air]^{ref} | Linear | 2017.00 | 965.73 | 976.91 | 14.74 | -47.81 | 0.01 | | | | | | 13.17 | 0.47 |
| Wheat | RYWALKA | linear | T[air]^{ref} | Linear | 2018.00 | 904.08 | 914.96 | 15.33 | -1.51 | 0.04 | | | | | | 13.70 | 0.46 |
| Wheat | RYWALKA | linear | T[air]^{ref} | Linear | 2019.00 | 875.98 | 886.56 | 18.59 | -45.50 | 0.01 | | | | | | 16.52 | 0.49 |
| Wheat | RYWALKA | linear | T[air]^{ref} | Linear | 2021.00 | 925.62 | 936.42 | 18.60 | -45.54 | 0.01 | | | | | | 16.46 | 0.49 |
| Wheat | RYWALKA | thermal | T[air]^{plot} | Thermal time | 2015.00 | 859.92 | 868.16 | | | | | | | | | | |
| Wheat | RYWALKA | thermal | T[air]^{plot} | Thermal time | 2016.00 | 830.72 | 838.77 | | | | | | | | | | |
| Wheat | RYWALKA | thermal | T[air]^{plot} | Thermal time | 2017.00 | 933.41 | 941.80 | 13.30 | | | | | | | | | |
| Wheat | RYWALKA | thermal | T[air]^{plot} | Thermal time | 2018.00 | 865.12 | 873.28 | | | | | | | | | | |
| Wheat | RYWALKA | thermal | T[air]^{plot} | Thermal time | 2019.00 | 794.11 | 802.05 | | | | | | | | | | |
| Wheat | RYWALKA | thermal | T[air]^{plot} | Thermal time | 2021.00 | 840.95 | 849.05 | | | | | | | | | | |
| Wheat | RYWALKA | thermal | T[air]^{ref} | Thermal time | 2015.00 | 898.71 | 906.94 | 14.09 | | | | | | | | | |
| Wheat | RYWALKA | thermal | T[air]^{ref} | Thermal time | 2016.00 | 862.49 | 870.54 | 14.86 | | | | | | | | | |
| Wheat | RYWALKA | thermal | T[air]^{ref} | Thermal time | 2017.00 | 936.64 | 945.03 | 13.49 | | | | | | | | | |
| Wheat | RYWALKA | thermal | T[air]^{ref} | Thermal time | 2018.00 | 903.17 | 911.32 | 15.49 | | | | | | | | | |
| Wheat | RYWALKA | thermal | T[air]^{ref} | Thermal time | 2019.00 | 829.60 | 837.53 | 14.47 | | | | | | | | | |
| Wheat | RYWALKA | thermal | T[air]^{ref} | Thermal time | 2021.00 | 878.41 | 886.51 | 15.20 | | | | | | | | | |
| Wheat | RYWALKA | wang | T[air]^{plot} | Wang-Engel | 2015.00 | 878.12 | 889.10 | | | | | | | 17.96 | 1.09 | 12.63 | 0.47 |
| Wheat | RYWALKA | wang | T[air]^{plot} | Wang-Engel | 2016.00 | 858.78 | 869.51 | | | | | | | 18.60 | 1.02 | 14.46 | 0.52 |
| Wheat | RYWALKA | wang | T[air]^{plot} | Wang-Engel | 2017.00 | 966.44 | 977.63 | 15.09 | | | | | | 17.95 | 0.95 | 13.13 | 0.51 |
| Wheat | RYWALKA | wang | T[air]^{plot} | Wang-Engel | 2018.00 | 885.10 | 895.98 | | | | | | | 18.45 | 1.04 | 14.58 | 0.54 |
| Wheat | RYWALKA | wang | T[air]^{plot} | Wang-Engel | 2019.00 | 822.88 | 833.46 | | | | | | | 19.25 | 1.17 | 14.83 | 0.51 |
| Wheat | RYWALKA | wang | T[air]^{plot} | Wang-Engel | 2021.00 | 871.25 | 882.05 | | | | | | | 18.65 | 1.07 | 14.76 | 0.54 |
| Wheat | RYWALKA | wang | T[air]^{ref} | Wang-Engel | 2015.00 | 895.88 | 906.86 | 13.44 | | | | | | 17.72 | 1.07 | 11.88 | 0.47 |
| Wheat | RYWALKA | wang | T[air]^{ref} | Wang-Engel | 2016.00 | 875.48 | 886.21 | 16.00 | | | | | | 18.15 | 0.97 | 13.94 | 0.50 |
| Wheat | RYWALKA | wang | T[air]^{ref} | Wang-Engel | 2017.00 | 963.40 | 974.58 | 14.89 | | | | | | 17.72 | 0.93 | 12.96 | 0.51 |
| Wheat | RYWALKA | wang | T[air]^{ref} | Wang-Engel | 2018.00 | 911.66 | 922.53 | 16.42 | | | | | | 18.08 | 1.00 | 14.18 | 0.52 |
| Wheat | RYWALKA | wang | T[air]^{ref} | Wang-Engel | 2019.00 | 848.13 | 858.71 | 16.03 | | | | | | 18.84 | 1.11 | 14.29 | 0.47 |
| Wheat | RYWALKA | wang | T[air]^{ref} | Wang-Engel | 2021.00 | 895.86 | 906.66 | 16.41 | | | | | | 18.27 | 1.02 | 14.21 | 0.52 |
| Wheat | SEMAFOR | asym | T[air]^{plot} | Asymptotic | 2015.00 | 757.75 | 770.97 | | 5.20 | | | 5.21 | 0.53 | | | 10.59 | 0.61 |
| Wheat | SEMAFOR | asym | T[air]^{plot} | Asymptotic | 2016.00 | 730.12 | 743.05 | | 5.60 | | | 5.81 | 0.48 | | | 11.62 | 0.72 |
| Wheat | SEMAFOR | asym | T[air]^{plot} | Asymptotic | 2017.00 | 729.25 | 742.47 | 8.74 | 4.50 | | | 4.13 | 0.49 | | | 7.94 | 0.42 |
| Wheat | SEMAFOR | asym | T[air]^{plot} | Asymptotic | 2018.00 | 761.96 | 775.08 | | 5.40 | | | 5.08 | 0.49 | | | 11.68 | 0.71 |
| Wheat | SEMAFOR | asym | T[air]^{plot} | Asymptotic | 2019.00 | 690.68 | 703.35 | | 5.30 | | | 3.58 | 0.46 | | | 11.57 | 0.85 |
| Wheat | SEMAFOR | asym | T[air]^{plot} | Asymptotic | 2021.00 | 732.37 | 745.35 | | 5.35 | | | 4.93 | 0.47 | | | 11.29 | 0.80 |
| Wheat | SEMAFOR | asym | T[air]^{ref} | Asymptotic | 2015.00 | 788.82 | 802.04 | 11.94 | 5.12 | | | 5.42 | 0.54 | | | 10.61 | 0.46 |
| Wheat | SEMAFOR | asym | T[air]^{ref} | Asymptotic | 2016.00 | 776.92 | 789.84 | 13.62 | 7.80 | | | 11.74 | 0.60 | | | 12.61 | 0.39 |
| Wheat | SEMAFOR | asym | T[air]^{ref} | Asymptotic | 2017.00 | 728.76 | 741.99 | 8.63 | 3.14 | | | 3.32 | 0.49 | | | 7.93 | 0.40 |
| Wheat | SEMAFOR | asym | T[air]^{ref} | Asymptotic | 2018.00 | 793.98 | 807.11 | 13.28 | 5.30 | | | 8.50 | 0.51 | | | 11.72 | 0.47 |
| Wheat | SEMAFOR | asym | T[air]^{ref} | Asymptotic | 2019.00 | 722.50 | 735.17 | 13.27 | 4.40 | | | 2.27 | 0.58 | | | 11.63 | 0.49 |
| Wheat | SEMAFOR | asym | T[air]^{ref} | Asymptotic | 2021.00 | 766.14 | 779.12 | 13.25 | 5.30 | | | 8.46 | 0.51 | | | 11.47 | 0.51 |
| Wheat | SEMAFOR | bilinear | T[air]^{plot} | Bi-linear | 2015.00 | 758.99 | 772.21 | | 10.46 | -0.00 | 0.56 | | | | | 10.63 | 0.63 |
| Wheat | SEMAFOR | bilinear | T[air]^{plot} | Bi-linear | 2016.00 | 734.98 | 747.91 | | 9.66 | -0.00 | 0.48 | | | | | 11.93 | 0.73 |
| Wheat | SEMAFOR | bilinear | T[air]^{plot} | Bi-linear | 2017.00 | 719.85 | 733.07 | 8.59 | 7.26 | -0.00 | 0.49 | | | | | 7.59 | 0.47 |
| Wheat | SEMAFOR | bilinear | T[air]^{plot} | Bi-linear | 2018.00 | 766.03 | 779.16 | | 9.05 | 0.00 | 0.47 | | | | | 11.92 | 0.69 |
| Wheat | SEMAFOR | bilinear | T[air]^{plot} | Bi-linear | 2019.00 | 693.99 | 706.65 | | 8.92 | 0.01 | 0.40 | | | | | 11.79 | 0.83 |
| Wheat | SEMAFOR | bilinear | T[air]^{plot} | Bi-linear | 2021.00 | 732.11 | 745.09 | | 10.47 | -0.00 | 0.49 | | | | | 11.27 | 0.82 |
| Wheat | SEMAFOR | bilinear | T[air]^{ref} | Bi-linear | 2015.00 | 789.63 | 802.85 | 12.16 | 12.60 | -0.01 | 0.64 | | | | | 10.65 | 0.49 |
| Wheat | SEMAFOR | bilinear | T[air]^{ref} | Bi-linear | 2016.00 | 766.98 | 779.91 | 13.56 | 11.30 | -0.00 | 0.55 | | | | | 11.98 | 0.48 |
| Wheat | SEMAFOR | bilinear | T[air]^{ref} | Bi-linear | 2017.00 | 721.09 | 734.31 | 8.63 | 7.06 | -0.00 | 0.49 | | | | | 7.64 | 0.47 |
| Wheat | SEMAFOR | bilinear | T[air]^{ref} | Bi-linear | 2018.00 | 798.38 | 811.51 | 13.68 | 11.10 | 0.00 | 0.55 | | | | | 11.99 | 0.49 |
| Wheat | SEMAFOR | bilinear | T[air]^{ref} | Bi-linear | 2019.00 | 725.65 | 738.32 | 13.70 | 12.28 | 0.01 | 0.54 | | | | | 11.83 | 0.51 |

| | | | | | | | | | | | | | | | | | | |
|---|---|---|---|---|---|---|---|---|---|---|---|---|---|---|---|---|---|---|
| Wheat | SEMAFOR | bilinear | T[air]^{ref} | Bi-linear | 2021.00 | 766.49 | 779.47 | 13.62 | 11.60 | -0.00 | 0.55 | | | | | | 11.49 | 0.54 |
| Wheat | SEMAFOR | gauss | ^{plot} | Gaussian | 2015.00 | 950.63 | 958.56 | 25.31 | | | | | | | | 40.66 | 23.73 | 0.36 |
| Wheat | SEMAFOR | gauss | ^{plot} | Gaussian | 2016.00 | 900.44 | 908.20 | 25.75 | | | | | | | | 42.70 | 24.33 | 0.31 |
| Wheat | SEMAFOR | gauss | ^{plot} | Gaussian | 2017.00 | 937.58 | 945.51 | 22.88 | | | | | | | | 42.62 | 22.27 | 0.24 |
| Wheat | SEMAFOR | gauss | ^{plot} | Gaussian | 2018.00 | 945.85 | 953.72 | 26.73 | | | | | | | | 44.45 | 25.38 | 0.32 |
| Wheat | SEMAFOR | gauss | ^{plot} | Gaussian | 2019.00 | 846.79 | 854.39 | 24.85 | | | | | | | | 43.83 | 23.35 | 0.35 |
| Wheat | SEMAFOR | gauss | ^{plot} | Gaussian | 2021.00 | 916.81 | 924.59 | 26.31 | | | | | | | | 45.18 | 25.23 | 0.29 |
| Wheat | SEMAFOR | gauss | ^{ref} | Gaussian | 2015.00 | 950.63 | 958.56 | 25.31 | | | | | | | | 40.66 | 23.73 | 0.36 |
| Wheat | SEMAFOR | gauss | ^{ref} | Gaussian | 2016.00 | 900.44 | 908.20 | 25.75 | | | | | | | | 42.70 | 24.33 | 0.31 |
| Wheat | SEMAFOR | gauss | ^{ref} | Gaussian | 2017.00 | 937.58 | 945.51 | 22.88 | | | | | | | | 42.62 | 22.27 | 0.24 |
| Wheat | SEMAFOR | gauss | ^{ref} | Gaussian | 2018.00 | 945.85 | 953.72 | 26.73 | | | | | | | | 44.45 | 25.38 | 0.32 |
| Wheat | SEMAFOR | gauss | ^{ref} | Gaussian | 2019.00 | 846.79 | 854.39 | 24.85 | | | | | | | | 43.83 | 23.35 | 0.35 |
| Wheat | SEMAFOR | gauss | ^{ref} | Gaussian | 2021.00 | 916.81 | 924.59 | 26.31 | | | | | | | | 45.18 | 25.23 | 0.29 |
| Wheat | SEMAFOR | linear | T[air]^{plot} | Linear | 2015.00 | 785.34 | 795.92 | | -48.38 | 0.01 | | | | | | | 12.33 | 0.66 |
| Wheat | SEMAFOR | linear | T[air]^{plot} | Linear | 2016.00 | 751.69 | 762.03 | | -47.76 | 0.01 | | | | | | | 13.24 | 0.71 |
| Wheat | SEMAFOR | linear | T[air]^{plot} | Linear | 2017.00 | 717.30 | 727.88 | 8.58 | -51.66 | 0.01 | | | | | | | 7.57 | 0.48 |
| Wheat | SEMAFOR | linear | T[air]^{plot} | Linear | 2018.00 | 786.43 | 796.93 | | -47.62 | 0.01 | | | | | | | 13.44 | 0.67 |
| Wheat | SEMAFOR | linear | T[air]^{plot} | Linear | 2019.00 | 698.08 | 708.21 | | -3.93 | 0.03 | | | | | | | 12.21 | 0.80 |
| Wheat | SEMAFOR | linear | T[air]^{plot} | Linear | 2021.00 | 753.82 | 764.20 | | -48.03 | 0.01 | | | | | | | 12.85 | 0.74 |
| Wheat | SEMAFOR | linear | T[air]^{ref} | Linear | 2015.00 | 820.82 | 831.39 | 14.84 | -48.22 | 0.01 | | | | | | | 12.58 | 0.55 |
| Wheat | SEMAFOR | linear | T[air]^{ref} | Linear | 2016.00 | 787.04 | 797.38 | 16.31 | -47.60 | 0.01 | | | | | | | 13.47 | 0.56 |
| Wheat | SEMAFOR | linear | T[air]^{ref} | Linear | 2017.00 | 717.99 | 728.57 | 8.59 | -51.65 | 0.01 | | | | | | | 7.60 | 0.47 |
| Wheat | SEMAFOR | linear | T[air]^{ref} | Linear | 2018.00 | 804.10 | 814.60 | 13.97 | -6.14 | 0.03 | | | | | | | 12.46 | 0.46 |
| Wheat | SEMAFOR | linear | T[air]^{ref} | Linear | 2019.00 | 730.04 | 740.17 | 13.81 | -3.00 | 0.03 | | | | | | | 12.25 | 0.47 |
| Wheat | SEMAFOR | linear | T[air]^{ref} | Linear | 2021.00 | 772.66 | 783.04 | 13.74 | -6.31 | 0.03 | | | | | | | 11.97 | 0.50 |
| Wheat | SEMAFOR | thermal | T[air]^{plot} | Thermal time | 2015.00 | 775.91 | 783.84 | | | | | | | | | | | |
| Wheat | SEMAFOR | thermal | T[air]^{plot} | Thermal time | 2016.00 | 748.94 | 756.70 | | | | | | | | | | | |
| Wheat | SEMAFOR | thermal | T[air]^{plot} | Thermal time | 2017.00 | 775.42 | 783.36 | 10.75 | | | | | | | | | | |
| Wheat | SEMAFOR | thermal | T[air]^{plot} | Thermal time | 2018.00 | 780.07 | 787.94 | | | | | | | | | | | |
| Wheat | SEMAFOR | thermal | T[air]^{plot} | Thermal time | 2019.00 | 699.14 | 706.73 | | | | | | | | | | | |
| Wheat | SEMAFOR | thermal | T[air]^{plot} | Thermal time | 2021.00 | 748.72 | 756.51 | | | | | | | | | | | |
| Wheat | SEMAFOR | thermal | T[air]^{ref} | Thermal time | 2015.00 | 807.39 | 815.32 | 12.94 | | | | | | | | | | |
| Wheat | SEMAFOR | thermal | T[air]^{ref} | Thermal time | 2016.00 | 779.48 | 787.23 | 13.93 | | | | | | | | | | |
| Wheat | SEMAFOR | thermal | T[air]^{ref} | Thermal time | 2017.00 | 781.04 | 788.97 | 10.98 | | | | | | | | | | |
| Wheat | SEMAFOR | thermal | T[air]^{ref} | Thermal time | 2018.00 | 811.85 | 819.73 | 14.16 | | | | | | | | | | |
| Wheat | SEMAFOR | thermal | T[air]^{ref} | Thermal time | 2019.00 | 730.34 | 737.94 | 13.68 | | | | | | | | | | |
| Wheat | SEMAFOR | thermal | T[air]^{ref} | Thermal time | 2021.00 | 780.83 | 788.62 | 13.81 | | | | | | | | | | |
| Wheat | SEMAFOR | wang | T[air]^{plot} | Wang-Engel | 2015.00 | 759.69 | 770.27 | | | | | | | 15.48 | 0.66 | | 10.78 | 0.59 |
| Wheat | SEMAFOR | wang | T[air]^{plot} | Wang-Engel | 2016.00 | 739.27 | 749.61 | | | | | | | 15.06 | 0.53 | | 12.34 | 0.79 |
| Wheat | SEMAFOR | wang | T[air]^{plot} | Wang-Engel | 2017.00 | 741.15 | 751.72 | 10.72 | | | | | | 12.59 | 0.51 | | 8.50 | 0.61 |
| Wheat | SEMAFOR | wang | T[air]^{plot} | Wang-Engel | 2018.00 | 772.25 | 782.75 | | | | | | | 15.02 | 0.55 | | 12.44 | 0.73 |
| Wheat | SEMAFOR | wang | T[air]^{plot} | Wang-Engel | 2019.00 | 699.10 | 709.23 | | | | | | | 16.73 | 0.56 | | 12.28 | 0.87 |
| Wheat | SEMAFOR | wang | T[air]^{plot} | Wang-Engel | 2021.00 | 737.82 | 748.20 | | | | | | | 15.32 | 0.54 | | 11.74 | 0.82 |
| Wheat | SEMAFOR | wang | T[air]^{ref} | Wang-Engel | 2015.00 | 788.47 | 799.05 | 12.16 | | | | | | 15.94 | 0.69 | | 10.69 | 0.48 |
| Wheat | SEMAFOR | wang | T[air]^{ref} | Wang-Engel | 2016.00 | 770.54 | 780.87 | 14.55 | | | | | | 15.82 | 0.61 | | 12.33 | 0.54 |
| Wheat | SEMAFOR | wang | T[air]^{ref} | Wang-Engel | 2017.00 | 740.56 | 751.13 | 10.67 | | | | | | 12.21 | 0.51 | | 8.48 | 0.61 |
| Wheat | SEMAFOR | wang | T[air]^{ref} | Wang-Engel | 2018.00 | 803.47 | 813.97 | 14.63 | | | | | | 15.55 | 0.61 | | 12.42 | 0.54 |
| Wheat | SEMAFOR | wang | T[air]^{ref} | Wang-Engel | 2019.00 | 727.93 | 738.06 | 15.24 | | | | | | 17.04 | 0.64 | | 12.11 | 0.62 |
| Wheat | SEMAFOR | wang | T[air]^{ref} | Wang-Engel | 2021.00 | 769.84 | 780.22 | 14.54 | | | | | | 15.85 | 0.61 | | 11.80 | 0.59 |
| Wheat | TAMARO | asym | T[air]^{plot} | Asymptotic | 2015.00 | 922.39 | 936.37 | | 4.60 | | | 2.17 | 0.75 | | | | 12.35 | 0.47 |
| Wheat | TAMARO | asym | T[air]^{plot} | Asymptotic | 2016.00 | 872.97 | 886.74 | | 6.02 | | | 12.22 | 0.61 | | | | 11.80 | 0.54 |
| Wheat | TAMARO | asym | T[air]^{plot} | Asymptotic | 2017.00 | 855.72 | 869.57 | 10.03 | 5.50 | | | 5.00 | 0.61 | | | | 8.98 | 0.46 |

| | | | | | | | | | | | | | | | | |
|---|---|---|---|---|---|---|---|---|---|---|---|---|---|---|---|---|
| Wheat | TAMARO | asym | T[air]^{plot} | Asymptotic | 2018.00 | 905.49 | 919.34 | | 4.30 | | | 1.97 | 0.77 | | 12.72 | 0.47 |
| Wheat | TAMARO | asym | T[air]^{plot} | Asymptotic | 2019.00 | 866.72 | 880.40 | | 4.30 | | | 1.76 | 0.81 | | 12.32 | 0.49 |
| Wheat | TAMARO | asym | T[air]^{plot} | Asymptotic | 2021.00 | 879.22 | 892.85 | | 7.60 | | | 4.57 | 0.69 | | 13.53 | 0.52 |
| Wheat | TAMARO | asym | T[air]^{ref} | Asymptotic | 2015.00 | 953.26 | 967.24 | 13.94 | 4.30 | | | 2.11 | 0.76 | | 12.33 | 0.47 |
| Wheat | TAMARO | asym | T[air]^{ref} | Asymptotic | 2016.00 | 902.89 | 916.66 | 14.66 | 5.80 | | | 9.56 | 0.61 | | 11.43 | 0.60 |
| Wheat | TAMARO | asym | T[air]^{ref} | Asymptotic | 2017.00 | 854.19 | 868.04 | 9.83 | 5.22 | | | 4.99 | 0.61 | | 8.92 | 0.43 |
| Wheat | TAMARO | asym | T[air]^{ref} | Asymptotic | 2018.00 | 936.99 | 950.84 | 14.39 | 4.30 | | | 2.07 | 0.76 | | 12.71 | 0.48 |
| Wheat | TAMARO | asym | T[air]^{ref} | Asymptotic | 2019.00 | 896.62 | 910.30 | 13.98 | 4.30 | | | 1.83 | 0.80 | | 12.23 | 0.50 |
| Wheat | TAMARO | asym | T[air]^{ref} | Asymptotic | 2021.00 | 900.16 | 913.79 | 14.95 | 4.30 | | | 2.07 | 0.75 | | 12.87 | 0.52 |
| Wheat | TAMARO | bilinear | T[air]^{plot} | Bi-linear | 2015.00 | 922.50 | 936.48 | | 17.20 | -0.03 | 0.88 | | | | 12.36 | 0.49 |
| Wheat | TAMARO | bilinear | T[air]^{plot} | Bi-linear | 2016.00 | 875.93 | 889.70 | | 10.37 | 0.01 | 0.58 | | | | 11.96 | 0.56 |
| Wheat | TAMARO | bilinear | T[air]^{plot} | Bi-linear | 2017.00 | 844.67 | 858.52 | 9.27 | 8.21 | 0.01 | 0.56 | | | | 8.57 | 0.40 |
| Wheat | TAMARO | bilinear | T[air]^{plot} | Bi-linear | 2018.00 | 906.29 | 920.14 | | 18.31 | -0.03 | 0.90 | | | | 12.77 | 0.50 |
| Wheat | TAMARO | bilinear | T[air]^{plot} | Bi-linear | 2019.00 | 866.99 | 880.67 | | 21.20 | -0.09 | 0.99 | | | | 12.33 | 0.51 |
| Wheat | TAMARO | bilinear | T[air]^{plot} | Bi-linear | 2021.00 | 868.73 | 882.37 | | 20.76 | -0.08 | 0.98 | | | | 12.89 | 0.53 |
| Wheat | TAMARO | bilinear | T[air]^{ref} | Bi-linear | 2015.00 | 949.97 | 963.95 | 14.00 | 18.27 | -0.08 | 0.95 | | | | 12.15 | 0.50 |
| Wheat | TAMARO | bilinear | T[air]^{ref} | Bi-linear | 2016.00 | 907.02 | 920.79 | 14.86 | 12.80 | -0.00 | 0.69 | | | | 11.96 | 0.58 |
| Wheat | TAMARO | bilinear | T[air]^{ref} | Bi-linear | 2017.00 | 845.05 | 858.90 | 9.27 | 7.99 | 0.01 | 0.56 | | | | 8.58 | 0.39 |
| Wheat | TAMARO | bilinear | T[air]^{ref} | Bi-linear | 2018.00 | 935.82 | 949.68 | 14.59 | 18.50 | -0.06 | 0.93 | | | | 12.65 | 0.51 |
| Wheat | TAMARO | bilinear | T[air]^{ref} | Bi-linear | 2019.00 | 896.43 | 910.12 | 14.13 | 20.10 | -0.09 | 0.96 | | | | 12.22 | 0.51 |
| Wheat | TAMARO | bilinear | T[air]^{ref} | Bi-linear | 2021.00 | 898.62 | 912.26 | 15.01 | 18.60 | -0.06 | 0.92 | | | | 12.78 | 0.54 |
| Wheat | TAMARO | gauss | ^{plot} | Gaussian | 2015.00 | 1142.27 | 1150.66 | 28.92 | | | | | | 47.27 | 27.54 | 0.32 |
| Wheat | TAMARO | gauss | ^{plot} | Gaussian | 2016.00 | 1098.21 | 1106.47 | 28.65 | | | | | | 48.11 | 27.93 | 0.23 |
| Wheat | TAMARO | gauss | ^{plot} | Gaussian | 2017.00 | 1108.03 | 1116.34 | 27.06 | | | | | | 48.52 | 26.86 | 0.15 |
| Wheat | TAMARO | gauss | ^{plot} | Gaussian | 2018.00 | 1134.03 | 1142.34 | 31.01 | | | | | | 50.93 | 30.01 | 0.26 |
| Wheat | TAMARO | gauss | ^{plot} | Gaussian | 2019.00 | 1082.36 | 1090.56 | 29.11 | | | | | | 49.61 | 28.32 | 0.24 |
| Wheat | TAMARO | gauss | ^{plot} | Gaussian | 2021.00 | 1091.42 | 1099.60 | 31.43 | | | | | | 51.86 | 30.77 | 0.22 |
| Wheat | TAMARO | gauss | ^{ref} | Gaussian | 2015.00 | 1142.27 | 1150.66 | 28.92 | | | | | | 47.27 | 27.54 | 0.32 |
| Wheat | TAMARO | gauss | ^{ref} | Gaussian | 2016.00 | 1098.21 | 1106.47 | 28.65 | | | | | | 48.11 | 27.93 | 0.23 |
| Wheat | TAMARO | gauss | ^{ref} | Gaussian | 2017.00 | 1108.03 | 1116.34 | 27.06 | | | | | | 48.52 | 26.86 | 0.15 |
| Wheat | TAMARO | gauss | ^{ref} | Gaussian | 2018.00 | 1134.03 | 1142.34 | 31.01 | | | | | | 50.93 | 30.01 | 0.26 |
| Wheat | TAMARO | gauss | ^{ref} | Gaussian | 2019.00 | 1082.36 | 1090.56 | 29.11 | | | | | | 49.61 | 28.32 | 0.24 |
| Wheat | TAMARO | gauss | ^{ref} | Gaussian | 2021.00 | 1091.42 | 1099.60 | 31.43 | | | | | | 51.86 | 30.77 | 0.22 |
| Wheat | TAMARO | linear | T[air]^{plot} | Linear | 2015.00 | 967.74 | 978.93 | | -46.36 | 0.01 | | | | | 15.27 | 0.53 |
| Wheat | TAMARO | linear | T[air]^{plot} | Linear | 2016.00 | 885.19 | 896.20 | | -6.11 | 0.03 | | | | | 12.58 | 0.55 |
| Wheat | TAMARO | linear | T[air]^{plot} | Linear | 2017.00 | 850.24 | 861.32 | 9.77 | -50.77 | 0.01 | | | | | 8.88 | 0.44 |
| Wheat | TAMARO | linear | T[air]^{plot} | Linear | 2018.00 | 950.95 | 962.04 | | -45.97 | 0.01 | | | | | 15.84 | 0.50 |
| Wheat | TAMARO | linear | T[air]^{plot} | Linear | 2019.00 | 911.84 | 922.79 | | -46.26 | 0.01 | | | | | 15.42 | 0.54 |
| Wheat | TAMARO | linear | T[air]^{plot} | Linear | 2021.00 | 909.59 | 920.50 | | -45.96 | 0.01 | | | | | 15.86 | 0.53 |
| Wheat | TAMARO | linear | T[air]^{ref} | Linear | 2015.00 | 968.08 | 979.26 | 14.88 | -4.08 | 0.04 | | | | | 13.21 | 0.47 |
| Wheat | TAMARO | linear | T[air]^{ref} | Linear | 2016.00 | 940.22 | 951.24 | 17.88 | -47.23 | 0.01 | | | | | 14.02 | 0.61 |
| Wheat | TAMARO | linear | T[air]^{ref} | Linear | 2017.00 | 850.26 | 861.34 | 9.77 | -50.77 | 0.01 | | | | | 8.88 | 0.44 |
| Wheat | TAMARO | linear | T[air]^{ref} | Linear | 2018.00 | 984.19 | 995.28 | 18.29 | -45.98 | 0.01 | | | | | 15.83 | 0.52 |
| Wheat | TAMARO | linear | T[air]^{ref} | Linear | 2019.00 | 944.76 | 955.71 | 18.26 | -46.27 | 0.01 | | | | | 15.40 | 0.56 |
| Wheat | TAMARO | linear | T[air]^{ref} | Linear | 2021.00 | 942.89 | 953.80 | 18.65 | -45.96 | 0.01 | | | | | 15.85 | 0.55 |
| Wheat | TAMARO | thermal | T[air]^{plot} | Thermal time | 2015.00 | 935.41 | 943.79 | | | | | | | | | |
| Wheat | TAMARO | thermal | T[air]^{plot} | Thermal time | 2016.00 | 894.39 | 902.65 | | | | | | | | | |
| Wheat | TAMARO | thermal | T[air]^{plot} | Thermal time | 2017.00 | 894.97 | 903.29 | 11.77 | | | | | | | | |
| Wheat | TAMARO | thermal | T[air]^{plot} | Thermal time | 2018.00 | 914.79 | 923.10 | | | | | | | | | |
| Wheat | TAMARO | thermal | T[air]^{plot} | Thermal time | 2019.00 | 875.69 | 883.90 | | | | | | | | | |
| Wheat | TAMARO | thermal | T[air]^{plot} | Thermal time | 2021.00 | 879.17 | 887.35 | | | | | | | | | |
| Wheat | TAMARO | thermal | T[air]^{ref} | Thermal time | 2015.00 | 973.41 | 981.80 | 15.01 | | | | | | | | |

| | | | | | | | | | | | | | | | | | | |
|---|---|---|---|---|---|---|---|---|---|---|---|---|---|---|---|---|---|---|
| Wheat | TAMARO | thermal | T[air]^{ref} | Thermal time | 2016.00 | 933.07 | 941.33 | 15.25 | | | | | | | | | | |
| Wheat | TAMARO | thermal | T[air]^{ref} | Thermal time | 2017.00 | 899.95 | 908.26 | 11.97 | | | | | | | | | | |
| Wheat | TAMARO | thermal | T[air]^{ref} | Thermal time | 2018.00 | 952.42 | 960.74 | 15.16 | | | | | | | | | | |
| Wheat | TAMARO | thermal | T[air]^{ref} | Thermal time | 2019.00 | 910.22 | 918.43 | 14.56 | | | | | | | | | | |
| Wheat | TAMARO | thermal | T[air]^{ref} | Thermal time | 2021.00 | 917.08 | 925.26 | 15.68 | | | | | | | | | | |
| Wheat | TAMARO | wang | T[air]^{plot} | Wang-Engel | 2015.00 | 927.40 | 938.58 | | | | | | | 17.47 | 0.93 | | 12.73 | 0.48 |
| Wheat | TAMARO | wang | T[air]^{plot} | Wang-Engel | 2016.00 | 888.84 | 899.85 | | | | | | | 17.10 | 0.81 | | 12.79 | 0.59 |
| Wheat | TAMARO | wang | T[air]^{plot} | Wang-Engel | 2017.00 | 869.90 | 880.98 | 10.80 | | | | | | 15.92 | 0.76 | | 9.63 | 0.47 |
| Wheat | TAMARO | wang | T[air]^{plot} | Wang-Engel | 2018.00 | 913.93 | 925.01 | | | | | | | 17.66 | 0.90 | | 13.32 | 0.53 |
| Wheat | TAMARO | wang | T[air]^{plot} | Wang-Engel | 2019.00 | 877.94 | 888.88 | | | | | | | 18.21 | 0.94 | | 13.09 | 0.51 |
| Wheat | TAMARO | wang | T[air]^{plot} | Wang-Engel | 2021.00 | 878.97 | 889.88 | | | | | | | 17.72 | 0.90 | | 13.64 | 0.54 |
| Wheat | TAMARO | wang | T[air]^{ref} | Wang-Engel | 2015.00 | 949.65 | 960.84 | 14.12 | | | | | | 17.30 | 0.91 | | 12.24 | 0.50 |
| Wheat | TAMARO | wang | T[air]^{ref} | Wang-Engel | 2016.00 | 911.39 | 922.40 | 15.52 | | | | | | 16.90 | 0.80 | | 12.29 | 0.61 |
| Wheat | TAMARO | wang | T[air]^{ref} | Wang-Engel | 2017.00 | 868.01 | 879.09 | 10.68 | | | | | | 15.71 | 0.75 | | 9.55 | 0.47 |
| Wheat | TAMARO | wang | T[air]^{ref} | Wang-Engel | 2018.00 | 939.03 | 950.11 | 15.33 | | | | | | 17.42 | 0.88 | | 12.93 | 0.55 |
| Wheat | TAMARO | wang | T[air]^{ref} | Wang-Engel | 2019.00 | 899.79 | 910.73 | 14.54 | | | | | | 18.01 | 0.92 | | 12.52 | 0.52 |
| Wheat | TAMARO | wang | T[air]^{ref} | Wang-Engel | 2021.00 | 903.15 | 914.06 | 15.68 | | | | | | 17.49 | 0.88 | | 13.16 | 0.56 |
| Wheat | TORONTO | asym | T[air]^{plot} | Asymptotic | 2015.00 | 753.45 | 766.62 | | 5.61 | | | 9.21 | 0.53 | | | | 10.74 | 0.60 |
| Wheat | TORONTO | asym | T[air]^{plot} | Asymptotic | 2016.00 | 741.94 | 755.02 | | 5.21 | | | 4.18 | 0.51 | | | | 10.95 | 0.58 |
| Wheat | TORONTO | asym | T[air]^{plot} | Asymptotic | 2017.00 | 728.21 | 741.38 | 9.18 | 4.54 | | | 6.60 | 0.49 | | | | 8.18 | 0.45 |
| Wheat | TORONTO | asym | T[air]^{plot} | Asymptotic | 2018.00 | 746.82 | 759.89 | | 6.12 | | | 10.39 | 0.54 | | | | 11.23 | 0.56 |
| Wheat | TORONTO | asym | T[air]^{plot} | Asymptotic | 2019.00 | 631.31 | 643.59 | | 5.60 | | | 4.16 | 0.53 | | | | 11.20 | 0.63 |
| Wheat | TORONTO | asym | T[air]^{plot} | Asymptotic | 2021.00 | 745.10 | 758.17 | | 5.62 | | | 9.72 | 0.52 | | | | 11.13 | 0.59 |
| Wheat | TORONTO | asym | T[air]^{ref} | Asymptotic | 2015.00 | 784.66 | 797.84 | 12.50 | 5.30 | | | 8.38 | 0.52 | | | | 10.79 | 0.50 |
| Wheat | TORONTO | asym | T[air]^{ref} | Asymptotic | 2016.00 | 772.48 | 785.56 | 12.41 | 5.70 | | | 6.02 | 0.51 | | | | 11.00 | 0.48 |
| Wheat | TORONTO | asym | T[air]^{ref} | Asymptotic | 2017.00 | 727.34 | 740.51 | 9.18 | 3.68 | | | 7.15 | 0.48 | | | | 8.15 | 0.46 |
| Wheat | TORONTO | asym | T[air]^{ref} | Asymptotic | 2018.00 | 775.90 | 788.98 | 12.81 | 5.20 | | | 5.21 | 0.51 | | | | 11.14 | 0.49 |
| Wheat | TORONTO | asym | T[air]^{ref} | Asymptotic | 2019.00 | 668.59 | 680.86 | 13.03 | 8.10 | | | 5.97 | 0.60 | | | | 11.64 | 0.46 |
| Wheat | TORONTO | asym | T[air]^{ref} | Asymptotic | 2021.00 | 777.26 | 790.34 | 12.88 | 3.30 | | | 2.17 | 0.58 | | | | 11.22 | 0.49 |
| Wheat | TORONTO | bilinear | T[air]^{plot} | Bi-linear | 2015.00 | 759.65 | 772.83 | | 8.92 | 0.01 | 0.49 | | | | | | 11.09 | 0.61 |
| Wheat | TORONTO | bilinear | T[air]^{plot} | Bi-linear | 2016.00 | 745.25 | 758.33 | | 8.88 | 0.01 | 0.47 | | | | | | 11.14 | 0.59 |
| Wheat | TORONTO | bilinear | T[air]^{plot} | Bi-linear | 2017.00 | 721.12 | 734.29 | 9.11 | 7.33 | -0.00 | 0.49 | | | | | | 7.90 | 0.49 |
| Wheat | TORONTO | bilinear | T[air]^{plot} | Bi-linear | 2018.00 | 750.47 | 763.55 | | 8.73 | 0.01 | 0.47 | | | | | | 11.45 | 0.58 |
| Wheat | TORONTO | bilinear | T[air]^{plot} | Bi-linear | 2019.00 | 635.45 | 647.72 | | 10.44 | 0.01 | 0.50 | | | | | | 11.49 | 0.61 |
| Wheat | TORONTO | bilinear | T[air]^{plot} | Bi-linear | 2021.00 | 747.47 | 760.54 | | 10.31 | 0.01 | 0.51 | | | | | | 11.27 | 0.61 |
| Wheat | TORONTO | bilinear | T[air]^{ref} | Bi-linear | 2015.00 | 790.69 | 803.86 | 12.93 | 10.78 | 0.00 | 0.54 | | | | | | 11.11 | 0.51 |
| Wheat | TORONTO | bilinear | T[air]^{ref} | Bi-linear | 2016.00 | 774.84 | 787.91 | 12.71 | 8.80 | 0.01 | 0.46 | | | | | | 11.08 | 0.50 |
| Wheat | TORONTO | bilinear | T[air]^{ref} | Bi-linear | 2017.00 | 722.17 | 735.35 | 9.14 | 7.34 | -0.00 | 0.50 | | | | | | 7.95 | 0.49 |
| Wheat | TORONTO | bilinear | T[air]^{ref} | Bi-linear | 2018.00 | 780.41 | 793.49 | 13.08 | 10.68 | 0.01 | 0.52 | | | | | | 11.40 | 0.50 |
| Wheat | TORONTO | bilinear | T[air]^{ref} | Bi-linear | 2019.00 | 662.95 | 675.23 | 12.97 | 11.70 | 0.01 | 0.53 | | | | | | 11.27 | 0.50 |
| Wheat | TORONTO | bilinear | T[air]^{ref} | Bi-linear | 2021.00 | 777.54 | 790.61 | 13.08 | 11.10 | 0.01 | 0.53 | | | | | | 11.23 | 0.52 |
| Wheat | TORONTO | gauss | ^{plot} | Gaussian | 2015.00 | 946.65 | 954.55 | 26.84 | | | | | | | | 37.59 | 24.34 | 0.43 |
| Wheat | TORONTO | gauss | ^{plot} | Gaussian | 2016.00 | 931.92 | 939.77 | 26.01 | | | | | | | | 38.77 | 24.80 | 0.30 |
| Wheat | TORONTO | gauss | ^{plot} | Gaussian | 2017.00 | 950.84 | 958.74 | 25.87 | | | | | | | | 39.11 | 24.84 | 0.29 |
| Wheat | TORONTO | gauss | ^{plot} | Gaussian | 2018.00 | 951.19 | 959.03 | 29.05 | | | | | | | | 40.49 | 27.30 | 0.35 |
| Wheat | TORONTO | gauss | ^{plot} | Gaussian | 2019.00 | 803.47 | 810.84 | 27.71 | | | | | | | | 41.78 | 26.37 | 0.32 |
| Wheat | TORONTO | gauss | ^{plot} | Gaussian | 2021.00 | 948.78 | 956.63 | 28.35 | | | | | | | | 41.45 | 26.98 | 0.32 |
| Wheat | TORONTO | gauss | ^{ref} | Gaussian | 2015.00 | 946.65 | 954.55 | 26.84 | | | | | | | | 37.59 | 24.34 | 0.43 |
| Wheat | TORONTO | gauss | ^{ref} | Gaussian | 2016.00 | 931.92 | 939.77 | 26.01 | | | | | | | | 38.77 | 24.80 | 0.30 |
| Wheat | TORONTO | gauss | ^{ref} | Gaussian | 2017.00 | 950.84 | 958.74 | 25.87 | | | | | | | | 39.11 | 24.84 | 0.29 |
| Wheat | TORONTO | gauss | ^{ref} | Gaussian | 2018.00 | 951.19 | 959.03 | 29.05 | | | | | | | | 40.49 | 27.30 | 0.35 |
| Wheat | TORONTO | gauss | ^{ref} | Gaussian | 2019.00 | 803.47 | 810.84 | 27.71 | | | | | | | | 41.78 | 26.37 | 0.32 |

| | | | | | | | | | | | | | | | | | | |
|---|---|---|---|---|---|---|---|---|---|---|---|---|---|---|---|---|---|---|
| Wheat | TORONTO | gauss | ^{ref} | Gaussian | 2021.00 | 948.78 | 956.63 | 28.35 | | | | | | | | 41.45 | 26.98 | 0.32 |
| Wheat | TORONTO | linear | T[air]^{plot} | Linear | 2015.00 | 784.94 | 795.48 | | -48.06 | 0.01 | | | | | | | 12.81 | 0.63 |
| Wheat | TORONTO | linear | T[air]^{plot} | Linear | 2016.00 | 763.70 | 774.16 | | -48.31 | 0.01 | | | | | | | 12.44 | 0.62 |
| Wheat | TORONTO | linear | T[air]^{plot} | Linear | 2017.00 | 724.05 | 734.58 | 9.51 | -51.30 | 0.01 | | | | | | | 8.10 | 0.53 |
| Wheat | TORONTO | linear | T[air]^{plot} | Linear | 2018.00 | 772.83 | 783.29 | | -47.88 | 0.01 | | | | | | | 13.06 | 0.61 |
| Wheat | TORONTO | linear | T[air]^{plot} | Linear | 2019.00 | 663.44 | 673.25 | | -47.29 | 0.01 | | | | | | | 13.92 | 0.63 |
| Wheat | TORONTO | linear | T[air]^{plot} | Linear | 2021.00 | 773.48 | 783.94 | | -47.85 | 0.01 | | | | | | | 13.11 | 0.62 |
| Wheat | TORONTO | linear | T[air]^{ref} | Linear | 2015.00 | 815.59 | 826.13 | 15.22 | -48.10 | 0.01 | | | | | | | 12.74 | 0.56 |
| Wheat | TORONTO | linear | T[air]^{ref} | Linear | 2016.00 | 793.69 | 804.16 | 14.72 | -48.38 | 0.01 | | | | | | | 12.35 | 0.54 |
| Wheat | TORONTO | linear | T[air]^{ref} | Linear | 2017.00 | 724.28 | 734.82 | 9.53 | -51.30 | 0.01 | | | | | | | 8.11 | 0.53 |
| Wheat | TORONTO | linear | T[air]^{ref} | Linear | 2018.00 | 805.08 | 815.54 | 15.40 | -47.87 | 0.01 | | | | | | | 13.09 | 0.54 |
| Wheat | TORONTO | linear | T[air]^{ref} | Linear | 2019.00 | 667.01 | 676.83 | 13.14 | -1.53 | 0.03 | | | | | | | 11.68 | 0.47 |
| Wheat | TORONTO | linear | T[air]^{ref} | Linear | 2021.00 | 804.03 | 814.49 | 15.48 | -47.91 | 0.01 | | | | | | | 13.03 | 0.55 |
| Wheat | TORONTO | thermal | T[air]^{plot} | Thermal time | 2015.00 | 771.67 | 779.58 | | | | | | | | | | | |
| Wheat | TORONTO | thermal | T[air]^{plot} | Thermal time | 2016.00 | 760.23 | 768.08 | | | | | | | | | | | |
| Wheat | TORONTO | thermal | T[air]^{plot} | Thermal time | 2017.00 | 773.19 | 781.09 | 11.64 | | | | | | | | | | |
| Wheat | TORONTO | thermal | T[air]^{plot} | Thermal time | 2018.00 | 760.74 | 768.59 | | | | | | | | | | | |
| Wheat | TORONTO | thermal | T[air]^{plot} | Thermal time | 2019.00 | 638.64 | 646.01 | | | | | | | | | | | |
| Wheat | TORONTO | thermal | T[air]^{plot} | Thermal time | 2021.00 | 758.23 | 766.08 | | | | | | | | | | | |
| Wheat | TORONTO | thermal | T[air]^{ref} | Thermal time | 2015.00 | 803.36 | 811.26 | 13.82 | | | | | | | | | | |
| Wheat | TORONTO | thermal | T[air]^{ref} | Thermal time | 2016.00 | 790.28 | 798.13 | 13.72 | | | | | | | | | | |
| Wheat | TORONTO | thermal | T[air]^{ref} | Thermal time | 2017.00 | 777.71 | 785.61 | 11.92 | | | | | | | | | | |
| Wheat | TORONTO | thermal | T[air]^{ref} | Thermal time | 2018.00 | 791.92 | 799.76 | 13.82 | | | | | | | | | | |
| Wheat | TORONTO | thermal | T[air]^{ref} | Thermal time | 2019.00 | 665.66 | 673.02 | 13.02 | | | | | | | | | | |
| Wheat | TORONTO | thermal | T[air]^{ref} | Thermal time | 2021.00 | 789.12 | 796.96 | 13.87 | | | | | | | | | | |
| Wheat | TORONTO | wang | T[air]^{plot} | Wang-Engel | 2015.00 | 765.68 | 776.21 | | | | | | | 15.42 | 0.64 | | 11.55 | 0.52 |
| Wheat | TORONTO | wang | T[air]^{plot} | Wang-Engel | 2016.00 | 754.75 | 765.21 | | | | | | | 15.30 | 0.59 | | 11.83 | 0.53 |
| Wheat | TORONTO | wang | T[air]^{plot} | Wang-Engel | 2017.00 | 743.18 | 753.72 | 9.89 | | | | | | 13.13 | 0.55 | | 8.89 | 0.44 |
| Wheat | TORONTO | wang | T[air]^{plot} | Wang-Engel | 2018.00 | 758.69 | 769.15 | | | | | | | 15.64 | 0.63 | | 12.07 | 0.51 |
| Wheat | TORONTO | wang | T[air]^{plot} | Wang-Engel | 2019.00 | 640.56 | 650.38 | | | | | | | 17.04 | 0.70 | | 12.01 | 0.56 |
| Wheat | TORONTO | wang | T[air]^{plot} | Wang-Engel | 2021.00 | 755.54 | 766.00 | | | | | | | 16.02 | 0.65 | | 11.88 | 0.54 |
| Wheat | TORONTO | wang | T[air]^{ref} | Wang-Engel | 2015.00 | 795.44 | 805.97 | 13.07 | | | | | | 15.52 | 0.64 | | 11.48 | 0.48 |
| Wheat | TORONTO | wang | T[air]^{ref} | Wang-Engel | 2016.00 | 783.01 | 793.47 | 13.28 | | | | | | 15.50 | 0.60 | | 11.66 | 0.49 |
| Wheat | TORONTO | wang | T[air]^{ref} | Wang-Engel | 2017.00 | 743.42 | 753.95 | 9.92 | | | | | | 12.93 | 0.55 | | 8.90 | 0.44 |
| Wheat | TORONTO | wang | T[air]^{ref} | Wang-Engel | 2018.00 | 787.81 | 798.27 | 13.60 | | | | | | 15.78 | 0.63 | | 11.94 | 0.48 |
| Wheat | TORONTO | wang | T[air]^{ref} | Wang-Engel | 2019.00 | 665.71 | 675.52 | 13.22 | | | | | | 17.28 | 0.71 | | 11.59 | 0.49 |
| Wheat | TORONTO | wang | T[air]^{ref} | Wang-Engel | 2021.00 | 784.92 | 795.38 | 13.50 | | | | | | 16.09 | 0.65 | | 11.77 | 0.49 |
| Wheat | WINNETOU | asym | T[air]^{plot} | Asymptotic | 2015.00 | 727.46 | 740.07 | | 5.00 | | | 2.99 | 0.64 | | | | 14.95 | 0.55 |
| Wheat | WINNETOU | asym | T[air]^{plot} | Asymptotic | 2016.00 | 664.78 | 677.05 | | 5.20 | | | 3.20 | 0.62 | | | | 13.78 | 0.62 |
| Wheat | WINNETOU | asym | T[air]^{plot} | Asymptotic | 2017.00 | 675.35 | 687.74 | 11.84 | 5.33 | | | 9.61 | 0.61 | | | | 11.08 | 0.36 |
| Wheat | WINNETOU | asym | T[air]^{plot} | Asymptotic | 2018.00 | 708.06 | 720.50 | | 5.10 | | | 3.36 | 0.64 | | | | 15.43 | 0.50 |
| Wheat | WINNETOU | asym | T[air]^{plot} | Asymptotic | 2019.00 | 680.65 | 692.87 | | 7.20 | | | 10.01 | 0.63 | | | | 16.00 | 0.56 |
| Wheat | WINNETOU | asym | T[air]^{plot} | Asymptotic | 2021.00 | 868.42 | 881.92 | | 5.10 | | | 3.07 | 0.64 | | | | 14.42 | 0.54 |
| Wheat | WINNETOU | asym | T[air]^{ref} | Asymptotic | 2015.00 | 754.92 | 767.53 | 16.31 | 4.30 | | | 2.52 | 0.67 | | | | 14.50 | 0.47 |
| Wheat | WINNETOU | asym | T[air]^{ref} | Asymptotic | 2016.00 | 694.42 | 706.69 | 16.06 | 5.60 | | | 4.79 | 0.61 | | | | 13.55 | 0.51 |
| Wheat | WINNETOU | asym | T[air]^{ref} | Asymptotic | 2017.00 | 674.46 | 686.85 | 11.80 | 5.34 | | | 8.99 | 0.61 | | | | 11.03 | 0.37 |
| Wheat | WINNETOU | asym | T[air]^{ref} | Asymptotic | 2018.00 | 736.76 | 749.21 | 16.64 | 5.20 | | | 4.46 | 0.63 | | | | 15.02 | 0.44 |
| Wheat | WINNETOU | asym | T[air]^{ref} | Asymptotic | 2019.00 | 708.43 | 720.64 | 17.46 | 6.70 | | | 3.22 | 0.67 | | | | 15.46 | 0.48 |
| Wheat | WINNETOU | asym | T[air]^{ref} | Asymptotic | 2021.00 | 895.55 | 909.06 | 15.83 | 4.60 | | | 2.68 | 0.66 | | | | 14.06 | 0.47 |
| Wheat | WINNETOU | bilinear | T[air]^{plot} | Bi-linear | 2015.00 | 729.96 | 742.57 | | 11.68 | 0.00 | 0.64 | | | | | | 15.16 | 0.57 |
| Wheat | WINNETOU | bilinear | T[air]^{plot} | Bi-linear | 2016.00 | 660.20 | 672.47 | | 21.80 | -0.53 | 0.99 | | | | | | 13.39 | 0.66 |
| Wheat | WINNETOU | bilinear | T[air]^{plot} | Bi-linear | 2017.00 | 671.53 | 683.91 | 11.44 | 8.00 | 0.01 | 0.54 | | | | | | 10.84 | 0.34 |

| | | | | | | | | | | | | | | | | |
|---|---|---|---|---|---|---|---|---|---|---|---|---|---|---|---|---|
| Wheat | WINNETOU | bilinear | T[air]^{plot} | Bi-linear | 2018.00 | 700.88 | 713.32 | | 21.91 | -0.34 | 1.05 | | | | 14.79 | 0.55 |
| Wheat | WINNETOU | bilinear | T[air]^{plot} | Bi-linear | 2019.00 | 674.30 | 686.51 | | 22.80 | -0.52 | 1.02 | | | | 15.38 | 0.58 |
| Wheat | WINNETOU | bilinear | T[air]^{plot} | Bi-linear | 2021.00 | 870.87 | 884.38 | | 13.40 | -0.01 | 0.71 | | | | 14.59 | 0.56 |
| Wheat | WINNETOU | bilinear | T[air]^{ref} | Bi-linear | 2015.00 | 754.88 | 767.49 | 16.65 | 14.50 | -0.03 | 0.79 | | | | 14.50 | 0.50 |
| Wheat | WINNETOU | bilinear | T[air]^{ref} | Bi-linear | 2016.00 | 688.54 | 700.82 | 16.13 | 19.00 | -0.12 | 0.91 | | | | 13.10 | 0.56 |
| Wheat | WINNETOU | bilinear | T[air]^{ref} | Bi-linear | 2017.00 | 671.48 | 683.87 | 11.42 | 8.00 | 0.01 | 0.54 | | | | 10.83 | 0.33 |
| Wheat | WINNETOU | bilinear | T[air]^{ref} | Bi-linear | 2018.00 | 735.88 | 748.33 | 16.99 | 14.28 | -0.03 | 0.79 | | | | 14.96 | 0.49 |
| Wheat | WINNETOU | bilinear | T[air]^{ref} | Bi-linear | 2019.00 | 704.01 | 716.23 | 17.04 | 21.50 | -0.23 | 0.98 | | | | 15.04 | 0.48 |
| Wheat | WINNETOU | bilinear | T[air]^{ref} | Bi-linear | 2021.00 | 895.99 | 909.49 | 16.16 | 14.10 | -0.02 | 0.76 | | | | 14.09 | 0.50 |
| Wheat | WINNETOU | gauss | ^{plot} | Gaussian | 2015.00 | 879.35 | 886.91 | 31.03 | | | | | | 48.34 | 29.36 | 0.34 |
| Wheat | WINNETOU | gauss | ^{plot} | Gaussian | 2016.00 | 826.94 | 834.31 | 30.72 | | | | | | 49.55 | 30.27 | 0.20 |
| Wheat | WINNETOU | gauss | ^{plot} | Gaussian | 2017.00 | 842.08 | 849.51 | 29.64 | | | | | | 51.59 | 29.55 | 0.12 |
| Wheat | WINNETOU | gauss | ^{plot} | Gaussian | 2018.00 | 871.11 | 878.57 | 33.60 | | | | | | 53.77 | 33.00 | 0.21 |
| Wheat | WINNETOU | gauss | ^{plot} | Gaussian | 2019.00 | 817.53 | 824.86 | 31.23 | | | | | | 51.60 | 30.31 | 0.26 |
| Wheat | WINNETOU | gauss | ^{plot} | Gaussian | 2021.00 | 1061.36 | 1069.46 | 31.33 | | | | | | 51.00 | 30.64 | 0.23 |
| Wheat | WINNETOU | gauss | ^{ref} | Gaussian | 2015.00 | 879.35 | 886.91 | 31.03 | | | | | | 48.34 | 29.36 | 0.34 |
| Wheat | WINNETOU | gauss | ^{ref} | Gaussian | 2016.00 | 826.94 | 834.31 | 30.72 | | | | | | 49.55 | 30.27 | 0.20 |
| Wheat | WINNETOU | gauss | ^{ref} | Gaussian | 2017.00 | 842.08 | 849.51 | 29.64 | | | | | | 51.59 | 29.55 | 0.12 |
| Wheat | WINNETOU | gauss | ^{ref} | Gaussian | 2018.00 | 871.11 | 878.57 | 33.60 | | | | | | 53.77 | 33.00 | 0.21 |
| Wheat | WINNETOU | gauss | ^{ref} | Gaussian | 2019.00 | 817.53 | 824.86 | 31.23 | | | | | | 51.60 | 30.31 | 0.26 |
| Wheat | WINNETOU | gauss | ^{ref} | Gaussian | 2021.00 | 1061.36 | 1069.46 | 31.33 | | | | | | 51.00 | 30.64 | 0.23 |
| Wheat | WINNETOU | linear | T[air]^{plot} | Linear | 2015.00 | 750.35 | 760.44 | | -44.92 | 0.01 | | | | | 17.36 | 0.59 |
| Wheat | WINNETOU | linear | T[air]^{plot} | Linear | 2016.00 | 691.28 | 701.09 | | -45.50 | 0.01 | | | | | 16.52 | 0.65 |
| Wheat | WINNETOU | linear | T[air]^{plot} | Linear | 2017.00 | 678.87 | 688.78 | 12.08 | -48.98 | 0.01 | | | | | 11.48 | 0.34 |
| Wheat | WINNETOU | linear | T[air]^{plot} | Linear | 2018.00 | 729.94 | 739.89 | | -44.54 | 0.01 | | | | | 17.91 | 0.57 |
| Wheat | WINNETOU | linear | T[air]^{plot} | Linear | 2019.00 | 700.00 | 709.78 | | -44.20 | 0.01 | | | | | 18.41 | 0.58 |
| Wheat | WINNETOU | linear | T[air]^{plot} | Linear | 2021.00 | 895.36 | 906.16 | | -45.41 | 0.01 | | | | | 16.65 | 0.56 |
| Wheat | WINNETOU | linear | T[air]^{ref} | Linear | 2015.00 | 781.51 | 791.60 | 19.76 | -45.12 | 0.01 | | | | | 17.08 | 0.52 |
| Wheat | WINNETOU | linear | T[air]^{ref} | Linear | 2016.00 | 723.07 | 732.88 | 19.64 | -45.63 | 0.01 | | | | | 16.33 | 0.56 |
| Wheat | WINNETOU | linear | T[air]^{ref} | Linear | 2017.00 | 678.86 | 688.77 | 12.07 | -48.98 | 0.01 | | | | | 11.48 | 0.34 |
| Wheat | WINNETOU | linear | T[air]^{ref} | Linear | 2018.00 | 761.11 | 771.06 | 20.25 | -44.77 | 0.01 | | | | | 17.59 | 0.51 |
| Wheat | WINNETOU | linear | T[air]^{ref} | Linear | 2019.00 | 731.36 | 741.13 | 20.85 | -44.45 | 0.01 | | | | | 18.05 | 0.52 |
| Wheat | WINNETOU | linear | T[air]^{ref} | Linear | 2021.00 | 925.98 | 936.78 | 18.81 | -45.58 | 0.01 | | | | | 16.41 | 0.50 |
| Wheat | WINNETOU | thermal | T[air]^{plot} | Thermal time | 2015.00 | 738.02 | 745.58 | | | | | | | | | |
| Wheat | WINNETOU | thermal | T[air]^{plot} | Thermal time | 2016.00 | 685.41 | 692.77 | | | | | | | | | |
| Wheat | WINNETOU | thermal | T[air]^{plot} | Thermal time | 2017.00 | 690.29 | 697.72 | 13.28 | | | | | | | | |
| Wheat | WINNETOU | thermal | T[air]^{plot} | Thermal time | 2018.00 | 722.05 | 729.51 | | | | | | | | | |
| Wheat | WINNETOU | thermal | T[air]^{plot} | Thermal time | 2019.00 | 687.74 | 695.07 | | | | | | | | | |
| Wheat | WINNETOU | thermal | T[air]^{plot} | Thermal time | 2021.00 | 885.30 | 893.40 | | | | | | | | | |
| Wheat | WINNETOU | thermal | T[air]^{ref} | Thermal time | 2015.00 | 772.01 | 779.57 | 18.34 | | | | | | | | |
| Wheat | WINNETOU | thermal | T[air]^{ref} | Thermal time | 2016.00 | 718.27 | 725.63 | 17.77 | | | | | | | | |
| Wheat | WINNETOU | thermal | T[air]^{ref} | Thermal time | 2017.00 | 693.00 | 700.44 | 13.44 | | | | | | | | |
| Wheat | WINNETOU | thermal | T[air]^{ref} | Thermal time | 2018.00 | 755.37 | 762.84 | 18.58 | | | | | | | | |
| Wheat | WINNETOU | thermal | T[air]^{ref} | Thermal time | 2019.00 | 718.47 | 725.80 | 18.35 | | | | | | | | |
| Wheat | WINNETOU | thermal | T[air]^{ref} | Thermal time | 2021.00 | 918.11 | 926.21 | 17.44 | | | | | | | | |
| Wheat | WINNETOU | wang | T[air]^{plot} | Wang-Engel | 2015.00 | 726.19 | 736.28 | | | | | 17.00 | 0.85 | | 15.01 | 0.55 |
| Wheat | WINNETOU | wang | T[air]^{plot} | Wang-Engel | 2016.00 | 662.18 | 672.00 | | | | | 17.14 | 0.80 | | 13.73 | 0.68 |
| Wheat | WINNETOU | wang | T[air]^{plot} | Wang-Engel | 2017.00 | 689.23 | 699.14 | 13.02 | | | | 15.46 | 0.73 | | 12.14 | 0.38 |
| Wheat | WINNETOU | wang | T[air]^{plot} | Wang-Engel | 2018.00 | 705.76 | 715.72 | | | | | 16.90 | 0.83 | | 15.40 | 0.54 |
| Wheat | WINNETOU | wang | T[air]^{plot} | Wang-Engel | 2019.00 | 679.11 | 688.88 | | | | | 17.94 | 0.87 | | 16.05 | 0.57 |
| Wheat | WINNETOU | wang | T[air]^{plot} | Wang-Engel | 2021.00 | 869.44 | 880.24 | | | | | 17.02 | 0.82 | | 14.63 | 0.57 |
| Wheat | WINNETOU | wang | T[air]^{ref} | Wang-Engel | 2015.00 | 751.33 | 761.41 | 16.42 | | | | 16.59 | 0.82 | | 14.37 | 0.49 |

| | | | | | | | | | | | | |
|---|---|---|---|---|---|---|---|---|---|---|---|---|
| Wheat | WINNETOU | wang | T[air]^{ref} | Wang-Engel | 2016.00 | 687.77 | 697.59 | 16.38 | 16.66 | 0.78 | 13.19 | 0.57 |
| Wheat | WINNETOU | wang | T[air]^{ref} | Wang-Engel | 2017.00 | 688.09 | 698.00 | 12.89 | 15.23 | 0.73 | 12.06 | 0.37 |
| Wheat | WINNETOU | wang | T[air]^{ref} | Wang-Engel | 2018.00 | 733.20 | 743.15 | 16.89 | 16.47 | 0.80 | 14.90 | 0.48 |
| Wheat | WINNETOU | wang | T[air]^{ref} | Wang-Engel | 2019.00 | 704.76 | 714.53 | 17.46 | 17.45 | 0.83 | 15.31 | 0.49 |
| Wheat | WINNETOU | wang | T[air]^{ref} | Wang-Engel | 2021.00 | 894.38 | 905.18 | 16.28 | 16.63 | 0.79 | 14.11 | 0.51 |
